# Mechanistic insights into transcriptional regulation of ARHGAP36 expression identify a factor predictive of neuroblastoma survival

**DOI:** 10.1101/2025.06.13.659594

**Authors:** Serhiy Havrylov, Armin M Gamper, Ordan J Lehmann

## Abstract

Cancer repeatedly exploits attributes fundamental for morphogenesis to advance malignancy and metastasis. This is illustrated by lineage specific transcription factors that regulate neural crest migration representing frequent drivers of malignancy. One such example is the *forkhead* transcription factor FOXC1 where gain of function is a feature of diverse cancers that is associated with an unfavourable prognosis. Using RNA-, ChIP-sequencing and CRISPR interference, we show that Foxc1 binds a locus in a region of closed chromatin to induce expression of Arhgap36, a tissue-specific inhibitor of Protein Kinase A. Because PKA is a core Hedgehog (Hh) pathway inhibitor, Foxc1’s induction of Arhgap36 expression increases Hh activity. The function of Sufu, a PKA substrate and a second essential Hh pathway inhibitor, is likewise impaired. The resulting increased Hh pathway output is resistant to pharmacological inhibition of *Smoothened*, a phenotype of more aggressive cancers. The Foxc1-Arhgap36 relationship identified in murine cells was further evaluated in neuroblastoma, a neural crest derived pediatric malignancy. This demonstrated in a cohort of 1348 patients that high levels of ARHGAP36 are predictive of improved five-year survival. Accordingly, this study has identified as a novel transcription factor which enhances ARHGAP36 expression, one that induces Hh activity in multiple tissues during development. It also establishes a model by which increased levels of FOXC1 via ARHGAP36 and PKA inhibition dysregulate multiple facets of Hh signaling, and provides evidence demonstrating relevance to a common neural-crest derived malignancy.

## Introduction

The neural crest comprises a multi-potent stem cell population that contributes to formation of diverse tissues. Through an epithelial to mesenchymal transition (EMT) that permits delamination from the neural plate, the acquisition of mesenchymal characteristics enables neural crest cells to transition towards a migratory phenotype. During a frequently lengthy path, neural crest cells proliferate, generating precursor cells that populate targets in multiple tissues, and so contribute to the formation of diverse organs. Such attributes essential to embryonic morphogenesis, are reminiscent of the invasive steps efficiently co-opted by cancer cells to drive malignancy and metastasis.

The lineage-specific transcription factors that regulate neural crest development represent frequent drivers of malignancy. An example of this paradigm is the *forkhead* box transcription factor *FOXC1*, that influences the migration and differentiation of neural crest cells. In addition to key roles in brain, ocular, cardiac, renal and skeletal development [1], FOXC1 is aberrantly expressed in diverse cancers. This overexpression was first identified in basal like breast cancer [2, 3], the subtype with the worst prognosis, and subsequently in at least fifteen additional tumor types, encompassing both solid tumors and hematological malignancies, such as AML [4, 5]. FOXC1 is also a prognostic factor for metastasis and survival, with its expression increasing tumour proliferation, invasion and dissemination [6]. Roles that include EMT induction, cell cycle control, angiogenesis, and regulation of stem cell populations are thought to promote FOXC1’s adverse oncological effects. However, despite such insights, the precise mechanisms behind FOXC1’s contributions to diverse malignancies remain incompletely defined.

Dysregulation of Hedgehog (Hh) signaling occurs frequently in sporadic and inherited cancers [7–10] and is estimated to underlie some 30% of malignancies. Hedgehog’s critical role in malignancy is linked to the pathway’s regulation of the Glioma (Gli) transcription factors and the requirement of Hh activity to maintain normal and cancer cell stemness. Notably, *FOXC1* and several of its paralogs are able to induce Hh expression and Hh pathway activity [11–15]. The output of the Hh pathway is tightly regulated by three equipotent inhibitors: the Patched transmembrane receptors, Suppressor of Fused (Sufu), and Protein Kinase A (PKA). Of these, PKA is positioned comparatively late in the signal transduction cascade and, by phosphorylating the GLI oncoproteins, directs their conversion from full length active to truncated repressor forms [8, 16, 17]. As a result, PKA controls the transcriptional outcome of Hh signaling. In recent years, a tissue-specific PKA antagonist has been identified. This Rho GTPase activating protein, Arhgap36, alters the balance between activator and repressor categories of GLI by inhibiting PKA, thereby inducing strong Hh pathway activation [18–20].

In this study, RNA- and ChIP-sequencing coupled with CRISPR interference were used to demonstrate that Foxc1 binds a locus located in a region of closed chromatin to induce Arhgap36 expression. In turn, Arhgap36 depletes the levels of PKA and its catalytic subunit PKAC, both strongly activating Hh signaling and making signal transduction less dependent on regulation via *Smoothened*. The oncological relevance of the Foxc1-Arhgap36 relationship is demonstrated in neuroblastoma, a neural crest derived pediatric malignancy where ARHGAP36 levels predict five-year mortality. Consequently, our study identifies a mechanism through which overexpression of FOXC1 dysregulates the Hedgehog pathway and contributes to malignancy.

## Results

To discern pathways by which Foxc1 may contribute to tumorigenesis, *Foxc1* was stably overexpressed in NIH3T3 fibroblasts. A retroviral construct was used that recapitulated the high levels of FOXC1 expression observed in a range of malignancies [15]. After RNA-sequencing, analysis using a stringent false discovery rate (*q* ≤ 0.01; log_2_fold change ≥ 1.0) defined a set of differentially expressed genes that were each distinguished by two independent bioinformatic workflows (DEseq2, and Cufdiff; Table S1). The majority were up-regulated by Foxc1 expression (n=192 genes; total 285; Fig. 1*A*). Gene-set enrichment analysis demonstrated significant enrichment for cancer invasiveness phenotypes, and malignancies, that importantly included neural crest associated malignancies such as melanoma and neuroblastoma (EMT *P* = 4.8 × 10^-11^; Malignancies *P* = 10^-3^ - 10^-4^; Table S2). Also enriched were phenotypes either induced by *FOXC1* mutation, (glaucoma and abnormal intraocular pressure, *P* ≤ 4.7 × 10^-4^) [21, 22] or present in *Foxc1^-/-^* murine mutants (abnormal skeletal morphology and renal anomalies, *P* = 0.05 - 10^-3^) [1, 23, 24]. Finally, genes relevant to Foxc1’s biological functions (angiogenesis, endochondral ossification, and hair follicle development) [13, 14, 25–30] were also enriched in relevant cell types (vasculature *P* = 2.1 × 10^-5^; neural crest *P* = 3.0 × 10^-4^; Table S2). Together, these results support the differentially expressed genes being representative of Foxc1 functions in development and disease.

**Figure 1.**
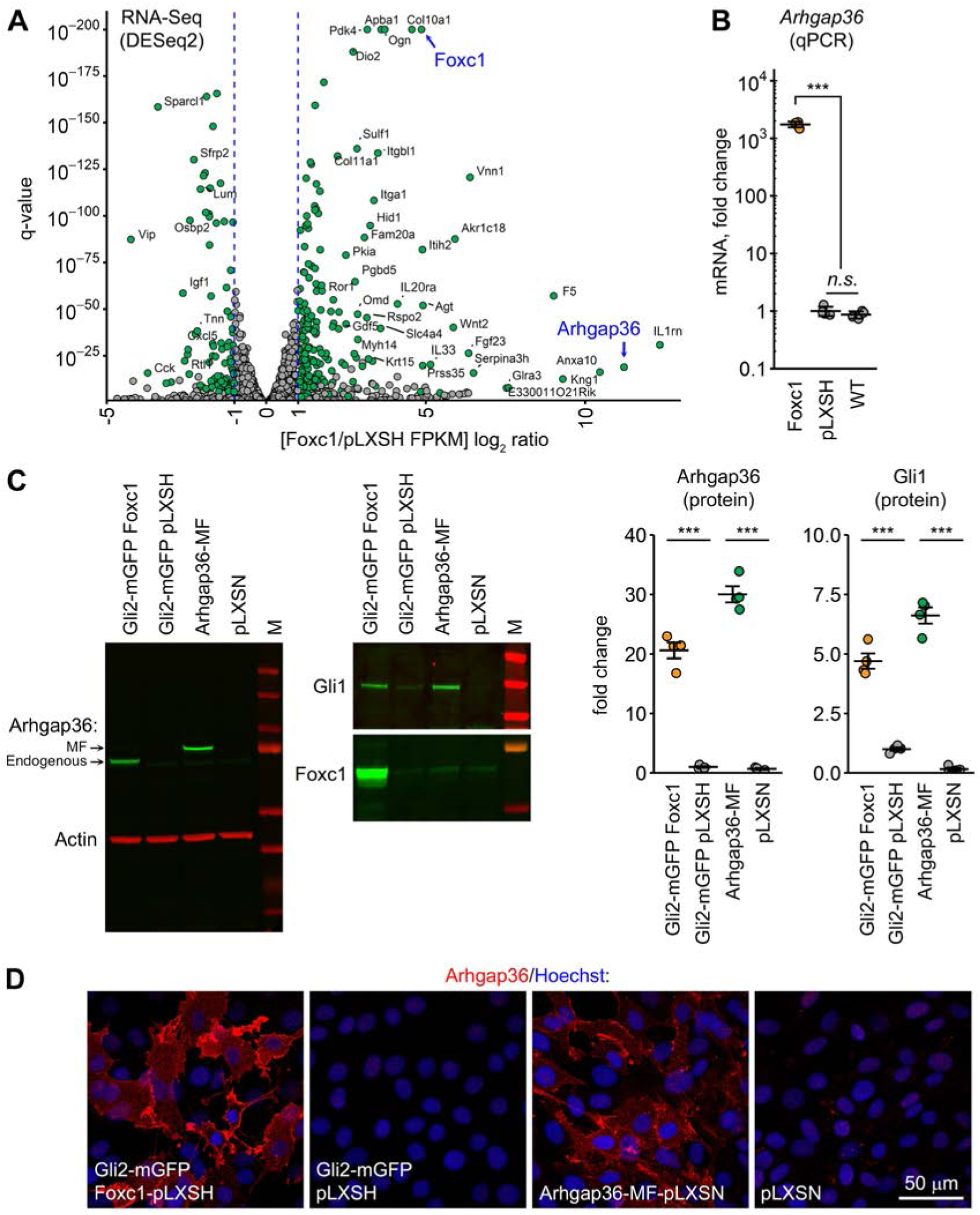
Foxc1 induces expression of Arhgap36 and activates Hedgehog signalling. (**A**) Volcano plot depicting in green differentially expressed genes with Foxc1 expression in NIH3T3 cells. (**B**) Confirmation of the robust Foxc1-induced increase in Arhgap36 mRNA in NIH3T3 cells by qPCR. (**C**) Foxc1 drives endogenous Arhgap36 protein expression, and that of Gli1, in NIH3T3-Gli2-mGFP cells to comparable levels of Myc-FLAG Arhgap36 ectopically expressed in parental NIH3T3 cells. (**D**) Immunofluorescence imaging demonstrates strong endogenous Arhgap36 expression, with the membrane staining in Foxc1-expressing cells recapitulating that of ectopically expressed Arhgap36-MF protein. [RNA-seq: n=3; quantitative Western blots: n=4 replicates; MF denotes Myc-FLAG tagged Arhgap36]

20% of the significantly dysregulated genes are implicated in Hh signalling (62 of 285) either as regulators or targets of the pathway (Table S1). Besides a cytokine that regulates inflammation and immunity (Interleukin Receptor 1 antagonist), Arhgap36 mRNA was the most upregulated in the RNA-sequencing dataset (Fig. 1A). To test whether Foxc1 consistently induces Arhgap36 expression in mesenchymal-derived cells, qPCR measurements were performed and strong induction of Arhgap36 mRNA observed after Foxc1 overexpression in several relevant cell lines [C2C12 (murine myoblast) 10^5^-fold, ATDC5 (chondrogenic) 10^2^-fold, NIH3T3 10^3^-fold increase; Fig. 1*B* and Fig. S1]. Moreover, in a Hh signalling reporter cell line (NIH3T3-Gli2-mGFP) [15, 31], Foxc1 overexpression induced Arhgap36 protein expression at levels comparable to ectopically expressed Myc-FLAG tagged Arhgap36 (Fig. 1*C*). Both Foxc1 overexpression and ectopically expressed Arhgap36 elevated protein expression of Gli1, a terminal effector and major readout of Hh pathway activity (Fig. 1*C*). Notably, the cellular localisation patterns of Foxc1-induced endogenous and ectopically expressed Arhgap36 are similar (Fig. 1*D*, Fig. S2*A*). Collectively, these data demonstrate Foxc1’s ability to induce Arhgap36 expression and activate Hedgehog signalling.

To characterize the functional interaction between the transcription factor Foxc1 and Arhgap36, whole genome ChIP-sequencing (ChIP-seq) was performed on Gli2-mGFP NIH3T3 cells stably expressing *Foxc1*. Two independent Foxc1 antibodies were used, together with an isotype-matched control immunoglobulin as control for Foxc1 peak specificity. The number of uniquely mapped 60 bp reads (Input > 5.3 × 10^8^, Foxc1-ChIP 4.8 × 10^8^) corresponds with deep sequencing coverage. Examination of chromatin accessibility demonstrates that, with the exception of embryonic stem cells, the Arhagp36 locus is inactive (Fig. S3), an observation consistent with Arhgap36’s highly restricted tissue expression. Within a 200 kb interval encompassing the Arhgap36 locus, five regions displayed strongly overlapping ChIP-seq peaks with both anti-Foxc1 antibodies (Fig. 2*A*). Three (Prox-1 to Prox-3) were 0.2 - 2.5 kb from the transcription start sites, while two (Dist-1, Dist-2) were 55 and 71 kb upstream (Fig. 2*A*). Two amplicons were selected within each peak, and ChIP-qPCR demonstrated strongly increased Foxc1 signal at each peak, relative to DNA precipitated using a non-specific normal IgG control (Fig. S4). Two groups of position weight matrices were significantly enriched in the bulk ChIP-seq peak data (STREME sequence motif discovery algorithm) (Fig. 2*B*, Fig. S4). The first comprised consensus sequences closely resembling known Foxc1-binding motifs (*P* = 10^-41^ - 10^-65^), while the second corresponded closely with that of Fos-Jun transcription factor dimers (*P* = 10^-13^ - 10^-20^, Fig. 2*B*, Fig. S5). Each of the five ChIP-seq peaks contains the identified Foxc1 motifs, supporting Arhgap36 being a direct transcriptional target of Foxc1.

**Figure 2.**
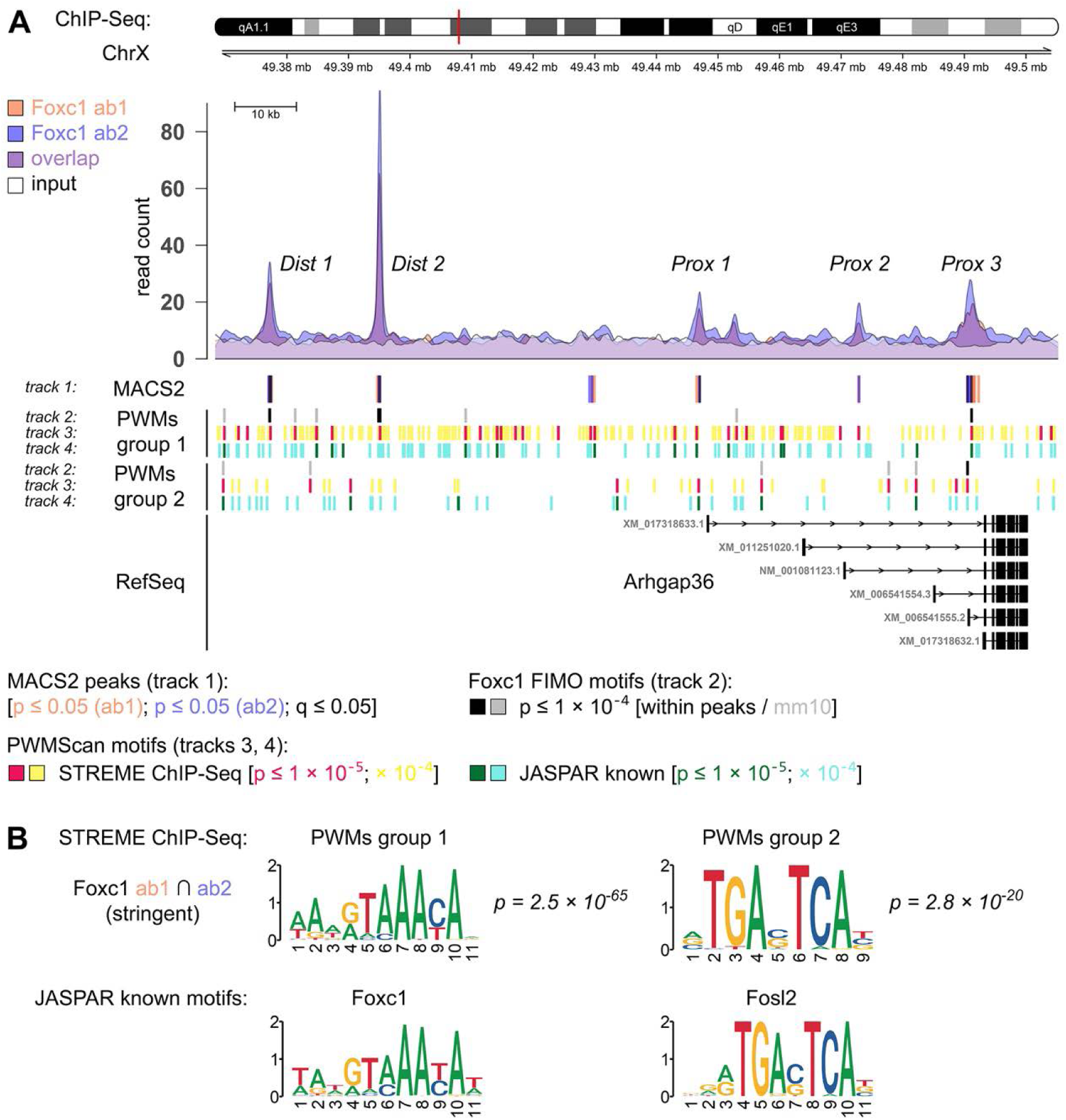
ChIP-seq identification of Foxc1-binding sites at the Arhgap36 locus. (**A**) ChIP sequencing with independent anti-Foxc1 antibodies revealed substantial peak overlap for both ChIP samples, consistent with high antibody specificity. Peak calling identified five significant ChIP signal regions within ± 100 kb of *Arhgap36* [2 distal, 3 proximal; q-value ≤ 0.05]. (**B**) Within these Foxc1 ChIP peaks, the discovery algorithm STREME identified two major groups of significantly enriched motifs [*p* = 2.8 × 10^-20^, 2.5 × 10^-65^]. The group 1 position weight matrices are highly similar to known Foxc1 motifs, while the group 2 PWMs very closely resemble the heptanucleotide recognition sequence bound by Fos-Jun transcription factor dimers [most prominently Fosl2]. The distribution of both PWM groups in the vicinity of Arhgap36 is shown on plot **A.**

We next investigated in a Hh reporter line [15] if Foxc1 overexpression recapitulated Arhgap36’s augmentation of Hh signaling, via down-regulation of PKA. As expected, ectopic expression of Foxc1 depleted the level of PKAC, PKA’s catalytic subunit (63% reduction, *P* = 1.4 × 10^-7^; Fig. 3*A*). Foxc1 overexpression also reduced phosphorylation of threonine 197 in PKAC’s activation loop, a residue essential to PKA enzymatic function and protein stability [32, 33]. This *∼*70% depletion of pT197 PKAC (*P* = 1.3 × 10^-8^) is analogous to the effect of ectopic Arhgap36 expression in parental NIH3T3 cells (Fig. 3*A*). Immunofluorescent staining demonstrated in cells with ectopic Foxc1 or Arhgap36 expression the near complete absence of PKAC and pT197 PKAC in the cytoplasm, especially at the basal body, where compartmentalized PKA-dependent control of Hh pathway occurs (Fig. 3*B*). Notably, in a transduced cell population with heterogeneous Foxc1 expression, PKAC signal is almost entirely depleted from cells with a high Foxc1 nuclear signal, demonstrating an inverse correlation between levels of PKAC and nuclear Foxc1 immunostaining (Fig. S2*B*). Similar results were observed in a second cell model, pre-adipocyte cells (3T3-L1) where expression of Foxc1 induced expression of Arhgap36 (albeit at lower levels) and a 40-50% reduction in protein levels of PKAC and pT197 PKAC (Fig. S2*C*,*D*). Direct overexpression of Arhgap36 led to strong downregulation of PKAC. These data, from two cell-based systems, demonstrate that Foxc1 expression depletes PKAC, the enzyme essential for promoting repressor forms of Gli, which regulate Hh target gene expression.

**Figure 3.**
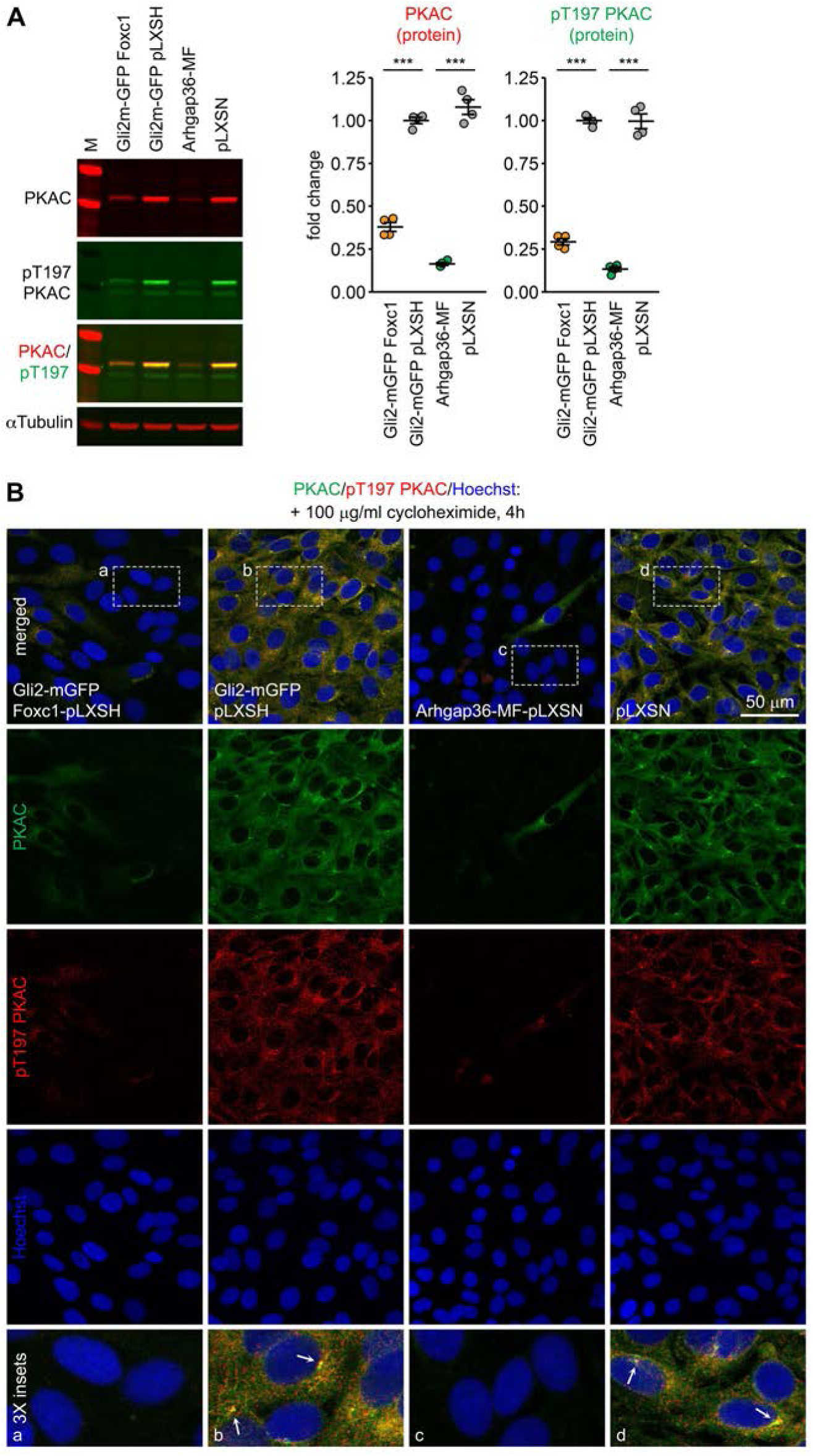
Foxc1-induced Arhgap36 reduces levels of protein kinase A catalytic subunit (PKAC). (**A**) Foxc1 expression in Gli2-mGFP NIH3T3 cells strongly reduces PKAC, and catalytically active pT197 PKAC, to comparable levels to those observed with ectopic expression of Arhgap36-MF. Quantification shows > 2-fold reduction of PKAC/pT197 PKAC protein levels [Western blots: n=4 replicates]. (**B**) Immunofluorescent staining demonstrates equivalent reductions in PKAC/pT197 PKAC signal in the cytoplasm and at the basal bodies of cells expressing either Foxc1 or ectopic Arhgap36-MF. [Dashed box: 3x insets, basal body: white arrows].

We next examined using CRIPSR interference whether steric repression of the five Foxc1 binding loci affected Arhgap36 transcription. Two clones of NIH3T3-Gli2-mGFP cells that overexpress Foxc1 were created that each stably expressed an inactive form of Cas9 fused with the Krüppel-associated box (KRAB) repressor domain of ZIM3 protein. Recruitment of ZIM3 KRAB-dCas9 to each of the five Foxc1-binding regions by retroviral delivery of five separate pools each comprising 3-4 single guide RNAs, altered Arhgap36 expression, although to varying degrees (Fig. 4*A*, Fig. S6). The sgRNA pools targeting Prox-3 and Dist-2 reduced Arghap36 mRNA expression by 98-99% and 67-76% respectively (Figure 4*A*). In both cases, reduced expression of Arhgap36 was associated with lower levels of Gli1 mRNA, and the degree of Arhgap36 mRNA reduction correlated with the decrease in Gli1 mRNA levels. Western immunoblot analysis confirmed that silencing either the Prox-3 or Dist-2 regions decreased Arhgap36 protein expression. This in turn correlated with an increase in PKAC and pT197 PKAC levels and a reduction in Gli1 protein levels (Fig. 4*B*). Since these data support Prox-3 and Dist-2 functioning, respectively, as promoter and distal enhancer for murine *Arhgap36*, evolutionary conservation of the Foxc1-binding loci was assessed. Multiple alignment across 60 vertebrates (Multiz) and measurements of evolutionary conservation (PhastCons) demonstrated that Prox-3 and Dist-2 are both conserved in placental, but not in marsupial mammals, nor in other vertebrates (Fig. S7). Multiple potential binding motifs for Foxc1 and Fos-Jun family transcription factors were identified in both regions; as illustrated by 16 Foxc1-binding motifs in both Prox-3 (8 motifs) and Dist-2 (8 motifs) of the murine locus, that were conserved or partially conserved in human (Fig. S7; Table S3). Consistent with this finding, luciferase reporter assays demonstrate that mutation of predicted Foxc1-binding motifs in the Prox-3 and Dist-2 regions, abrogates Foxc1-dependent transcriptional activity (Fig. S8). Taken together, the data presented support the Prox-3 and Dist-2 regions functioning as cis-regulatory elements for murine *Arhgap36*, and other placental mammals, including human.

**Figure 4.**
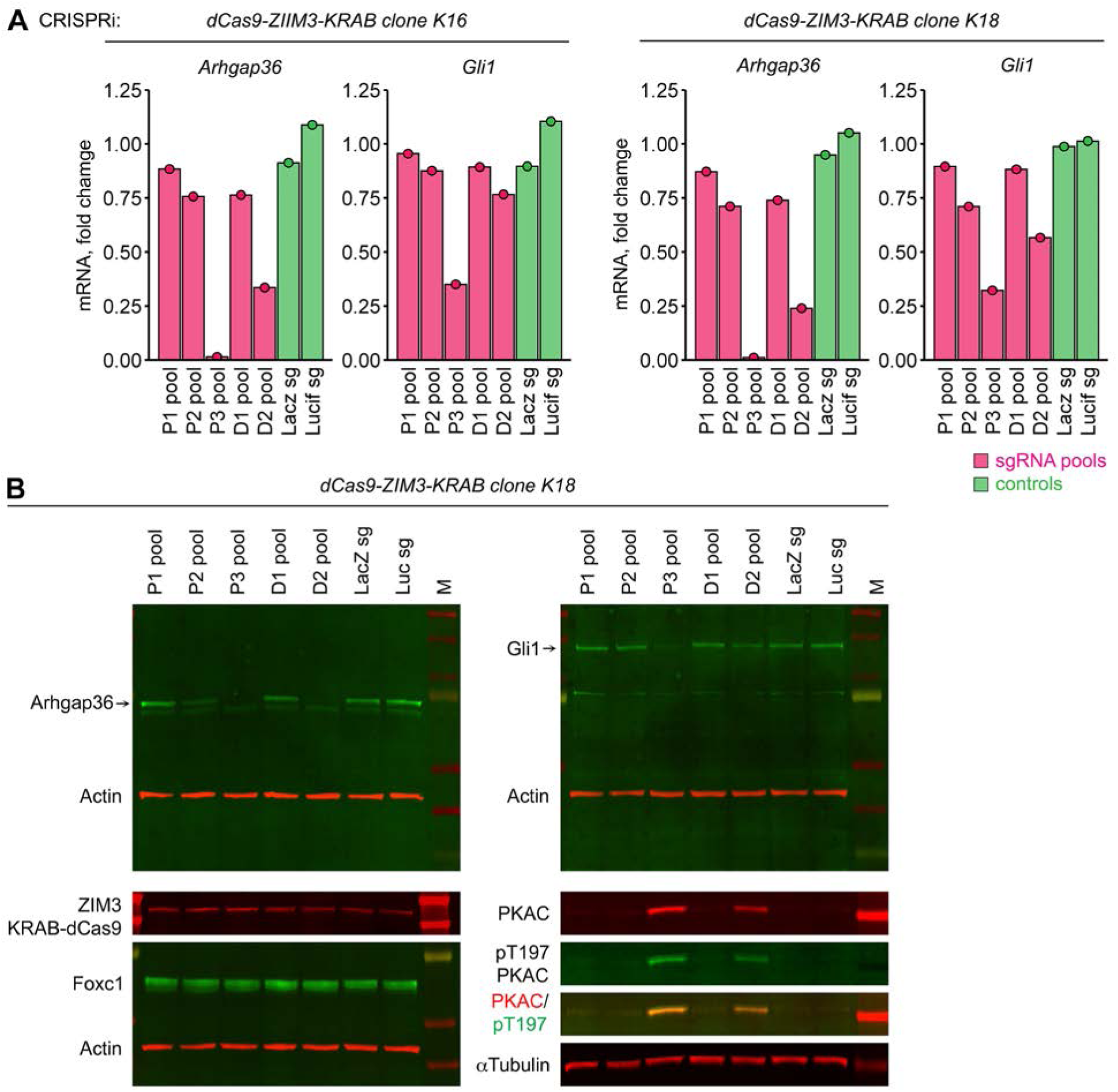
CRISPRi at potential Foxc1-binding sites diminishes Arhgap36 expression and Hedgehog activity. (**A**) In independent CRISPRi-competent clonal cell lines, pools of guide RNAs targeting the Prox-3 and Dist-2 regions, robustly reduced Arhgap36 and Gli1mRNA expression. (**B**) The reduced Arhgap36 and Gli1 protein expression with CRISPRi sgRNA targeting, implicates the same two ChIP-seq identified peaks in Foxc1’s transcriptional control of the Arhgap36 locus.

Since PKA depletion de-represses Hh signaling, the pathway activation is predicted to become less dependent on Hh ligand, which pharmacologically would manifest as tumour resistance to inhibition of *Smoothened* (*Smo*). To test whether Foxc1 expression induces such ligand-independent Hh signal transduction, Gli1 levels were measured with and without two *Smoothened*-specific antagonists (Sonidegib and Cyclopamine). The results revealed that the Foxc1-induced increase of Gli1 mRNA and protein levels is resistant to inhibition by either antagonist (Fig. 5*A*,*B*). Furthermore, the Foxc1-induced resistance to Sonidegib is phenocopied by the ectopic expression of Arhgap36 (Fig. S9*A*). In both cases Gli1 upregulation, whether by Foxc1 or by Arhgap36 overexpression, is not significantly reduced by co-treatment with Sonidegib. Indeed, Foxc1 overexpression leads to a similar level of Gli1 upregulation in NIH3T3 cells as treatment with a *Smoothened* agonist (SAG), and co-treatment of cells with SAG does not significantly enhance Gli1 levels beyond Foxc1-induced upregulation alone (Fig. 5*B*). Consequently, these data demonstrate that Foxc1 induces strong non-canonical activation of Hh signaling, a characteristic of multiple malignancies, including advanced and metastatic tumors.

**Figure 5.**
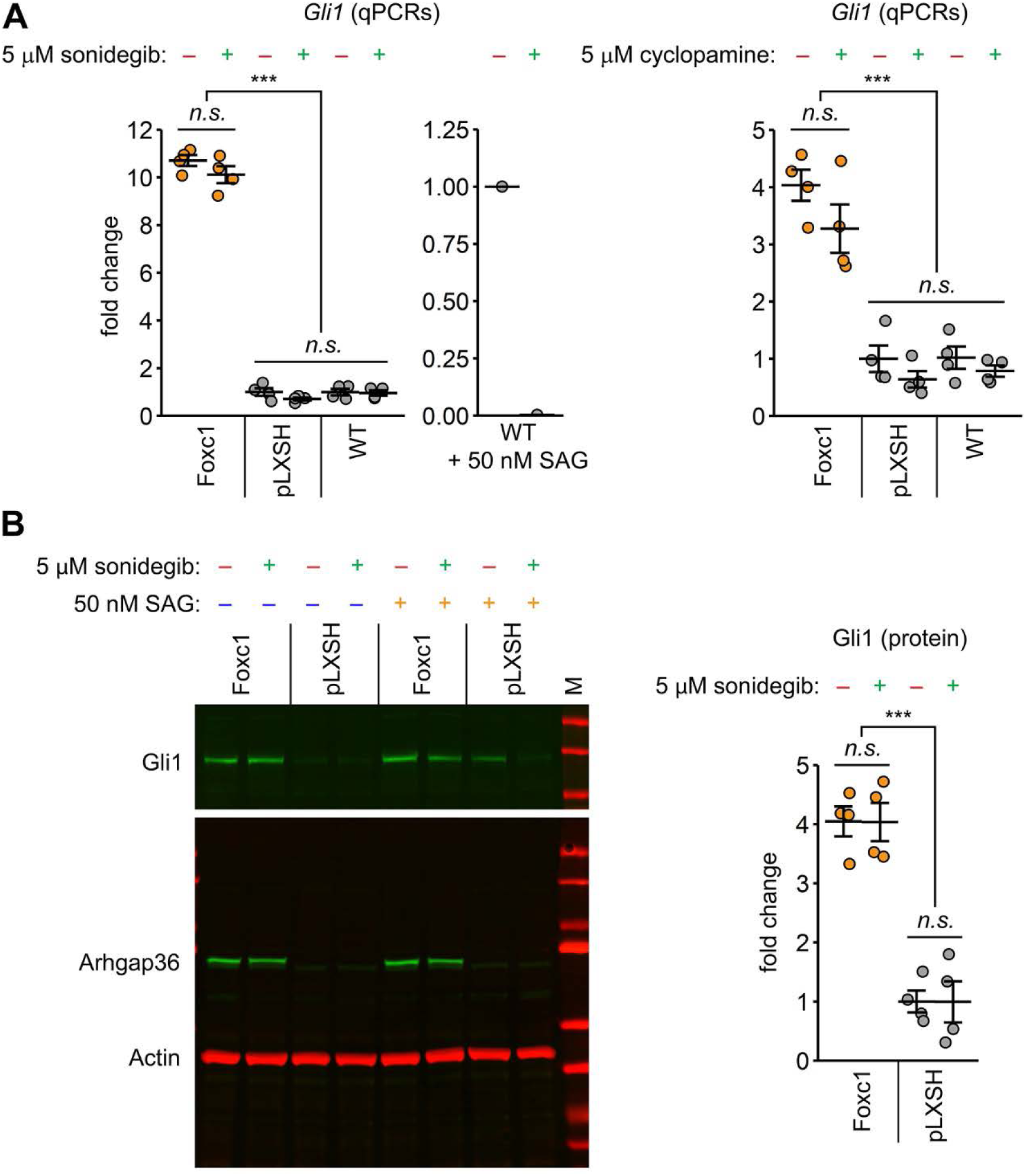
Foxc1-induced Hh signalling has reduced dependence on *Smoothened*. (**A**) Elevated levels of Gli1 mRNA in Foxc1-expressing NIH3T3 cells are resistant to inhibition by the *Smoothened* antagonists sonidegib and cyclopamine. Wild-type NIH3T3 cells stimulated with *Smoothened* agonist (SAG) and treated with sonidegib, provide a control for inhibitor efficiency. (**B**) Resistance to *Smoothened* inhibition is supported by the elevated Gli1 protein levels in Foxc1-expressing Gli2-mGFP NIH3T3 cells treated with sonidegib. Note that expression of Foxc1 induces comparable Gli1 protein levels to vector control cells [pLXSH] treated with SAG; and that levels of Arhgap36 protein itself are unaffected by either sonidegib, or SAG treatment. [qPCRs, quantitative Western blots: n=4 replicates].

PKA’s roles regulating the Hh pathway extend beyond repressing Gli activity. PKA also phosphorylates key serine and threonine residues within Sufu. This inhibits ciliary trafficking of Sufu-Gli complexes, resulting in retention of Sufu at the tip of the primary cilium [34, 35]. We found that in addition to substantially increased axonemal tip accumulation of Gli2-mGFP (Fig. 6*A*, Fig. S9*B*,*C*), expression of Foxc1 also increases Sufu levels at the tip of the axoneme 3-fold (*P* ≤ 2 × 10^-10^) while a similar 2-fold increase is induced by ectopic expression of Arhgap36 (Fig. 6*A-C*). The status of Sufu’s PKA-dependent phosphorylation sites is known to influence Sufu activity. Two of these adjacent serine residues comprise a classical dual phosphorylation site for PKA (pS346) and GSK3β (pS342) [34]. Consequently, the ∼50% reduction in levels of Sufu pS342 after Foxc1 (*P* = 6.4 × 10^-5^) or Arhgap36 expression (Fig. 6*D*), provides further support for Hh pathway regulation by Foxc1. Together these data demonstrate that the effects of Foxc1 expression on Hh signal transduction involve several core regulators of Hh pathway activity. The presented data support a model by which Foxc1-dependent Arhgap36 expression attenuates PKA activity and Sufu function. To verify that the observed changes in Sufu and Gli2 ciliary accumulation induced by Foxc1 overexpression are mediated by Arhgap36, shRNA silencing of Arhgap36 was performed in Foxc1-expressing Gli2-mGFP cells (Fig. 7). Two different Arhgap36-targeting shRNAs each substantially diminished ciliary Gli2-mGFP accumulation (shRNA1: 35%, shRNA2: 64% reduction; *P* = 4.9 × 10^-7^ and 4.5 × 10^-10^; Fig. 7*A*-*C*) while Gli2-mGFP levels in control cells that express very low levels of Arhgap36 were unchanged. The decrease in axonemal Gli2-mGFP correlated with the reduction in Gli1 mRNA (Fig. 7D), and the shRNA that more efficiently inhibited Arhgap36 induced a greater decrease in both Gli2-mGFP accumulation and Gli1 expression. The specificity of Arhgap36 inhibition is apparent from marked reductions in Gli1 and Arhgap36 mRNA expression, while levels of Foxc1 were unaltered (Fig. 7D). Analysis of Gli1 expression in parental Foxc1-expressing NIH3T3 cells treated with the same Arhgap36-targeting shRNA recapitulated the prior findings (Fig. S10). Taken together, these data demonstrate that the effects of Foxc1 expression on Hh pathway activity are Arhgap36-mediated.

**Figure 6.**
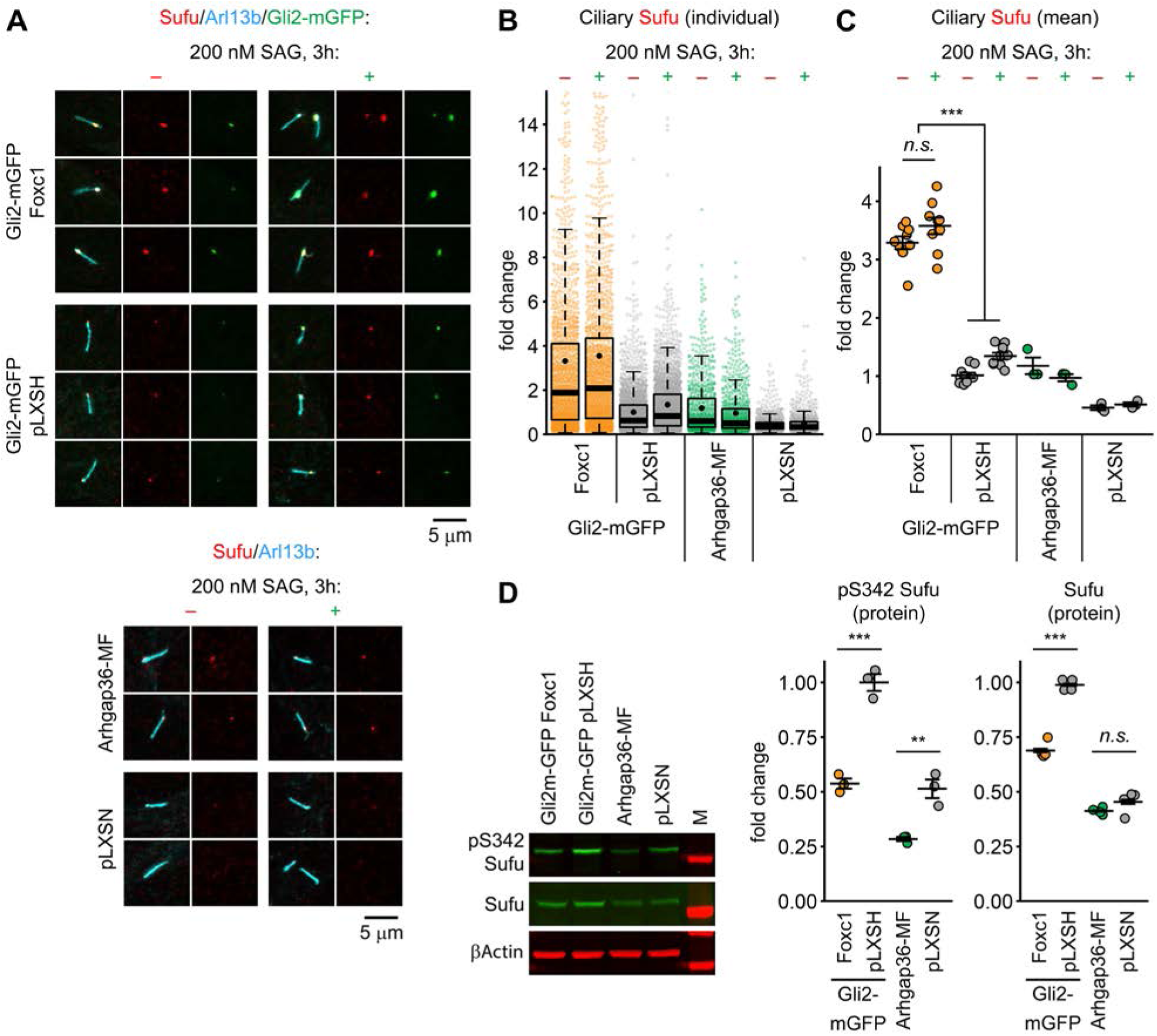
Foxc1 promotes ciliary accumulation and decreases phosphorylation of Sufu. (**A**) Representative immunofluorescence images demonstrate increased Sufu accumulation at axonemal tips of Foxc1-expressing cells, and NIH3T3 cells that express Myc-FLAG Arhgap36. Note, SAG stimulation *per se* does not substantially affect ciliary accumulation of Sufu. (**B**) Distribution of Sufu intensity in individual cilia (**C**) Mean ciliary Sufu intensity values. These demonstrate that the ciliary Sufu signal in cells expressing Foxc1, and separately Arhgap36, is substantially increased relative to empty vector controls [pLXSH and pLXSN; n = 9 combined experiments]. (**D**) Decreased phosphorylation of Sufu at the S342 residue in Gli2-mGFP NIH3T3s expressing Foxc1, and NIH3T3 cells expressing Myc-FLAG Arhgap36, relative to vector controls. Expression of Foxc1 also significantly impacts the total protein levels of Sufu, in contrast to the non-significant effect of Arhgap36 [quantitative Western blots: n=4 replicates].

**Figure 7.**
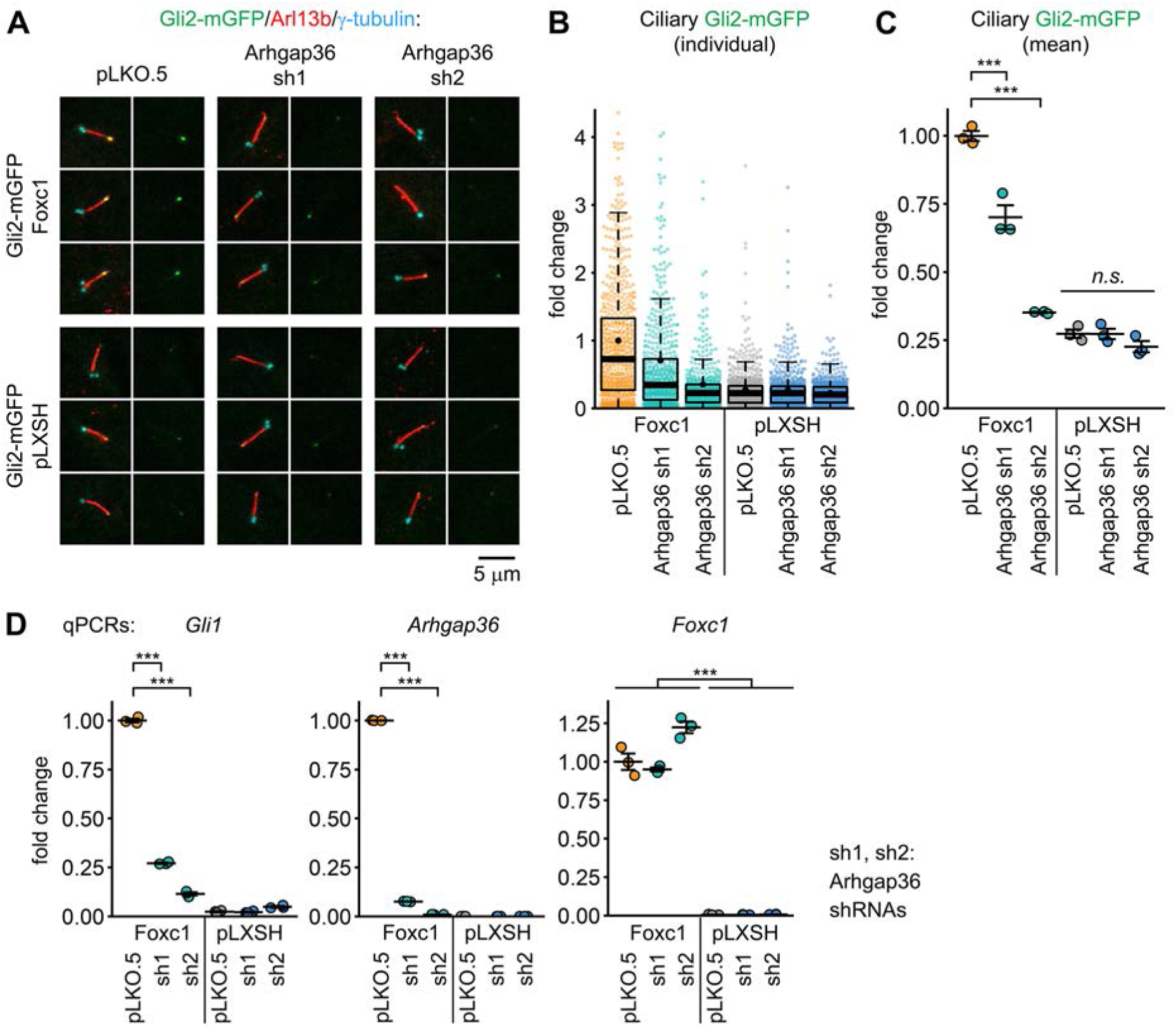
Arhgap36 is required for Foxc1-induced activation of Hh signalling. (**A**) Immunofluorescence images demonstrate that two Arhgap36-targeting shRNAs each substantially reduce axonemal tip accumulation of Gli2 in cells that express Foxc1. (**B**, **C**) Individual and mean ciliary Gli2 intensity values demonstrate the decrease of ciliary Gli2 signal with Arhgap36 shRNA inhibition [n = 3 combined experiments]. (**D**) qPCR analyses demonstrate a strong decrease in Gli1 mRNA levels in Foxc1-expressing cells treated with Arhgap36 shRNAs, relative to pLKO.5 control. Note, concordant Gli1 and Arhgap36 expression levels across all conditions.

To test the oncologic relevance of the Foxc1 - Arhgap36 relationship, *ARHGAP36* mRNA levels were first surveyed in cancer expression datasets [CCLE, TCGA, TARGET, PCAWG (Table S4)]. This revealed high expression in a common pediatric malignancy, neuroblastoma, as well as specific CNS, breast, lung, and neuroendocrine tumors. Due to neuroblastoma’s neural crest origin and Foxc1’s roles in this stem cell population, we focused on neuroblastoma data. Survival was evaluated in three independent neuroblastoma patient cohorts using Kaplan-Meier analyses after stratification into terciles of high, medium and low ARHGAP36 mRNA expression. The highest tercile of *ARHGAP36* expression correlated with a favorable overall survival compared to the lowest: <u>GSE49711</u>, 91 vs 63% 5-year survival; <u>E-MTAB-178191</u>, 91 vs 61%; <u>TARGET</u> study, 60 vs 31%, (Fig. 8*A-C*) [36–38]. The effect was consistent across all three cohorts, despite differences in cancer staging - the first two cohorts primarily comprise patients with early disease, while TARGET mainly consists of advanced or metastatic neuroblastoma cases. Merging the data into a single dataset of 1348 patients narrowed the confidence intervals to provide more precise estimates. The highest tercile of *ARHGAP36* expression was associated with 87% five-year survival (range 86-92%) in contrast to 58% (53-63%) for the lowest tercile (*P* = 1.7 × 10^-19^). Illustrating these findings in the context of *MYCN*, whose amplification represents the primary genetic marker for high-risk and prognostically poor neuroblastoma, high ARHGAP36 expression was associated with an 89% five-year survival compared to 37% with *MYCN* amplification (Fig. S11).

**Figure 8.**
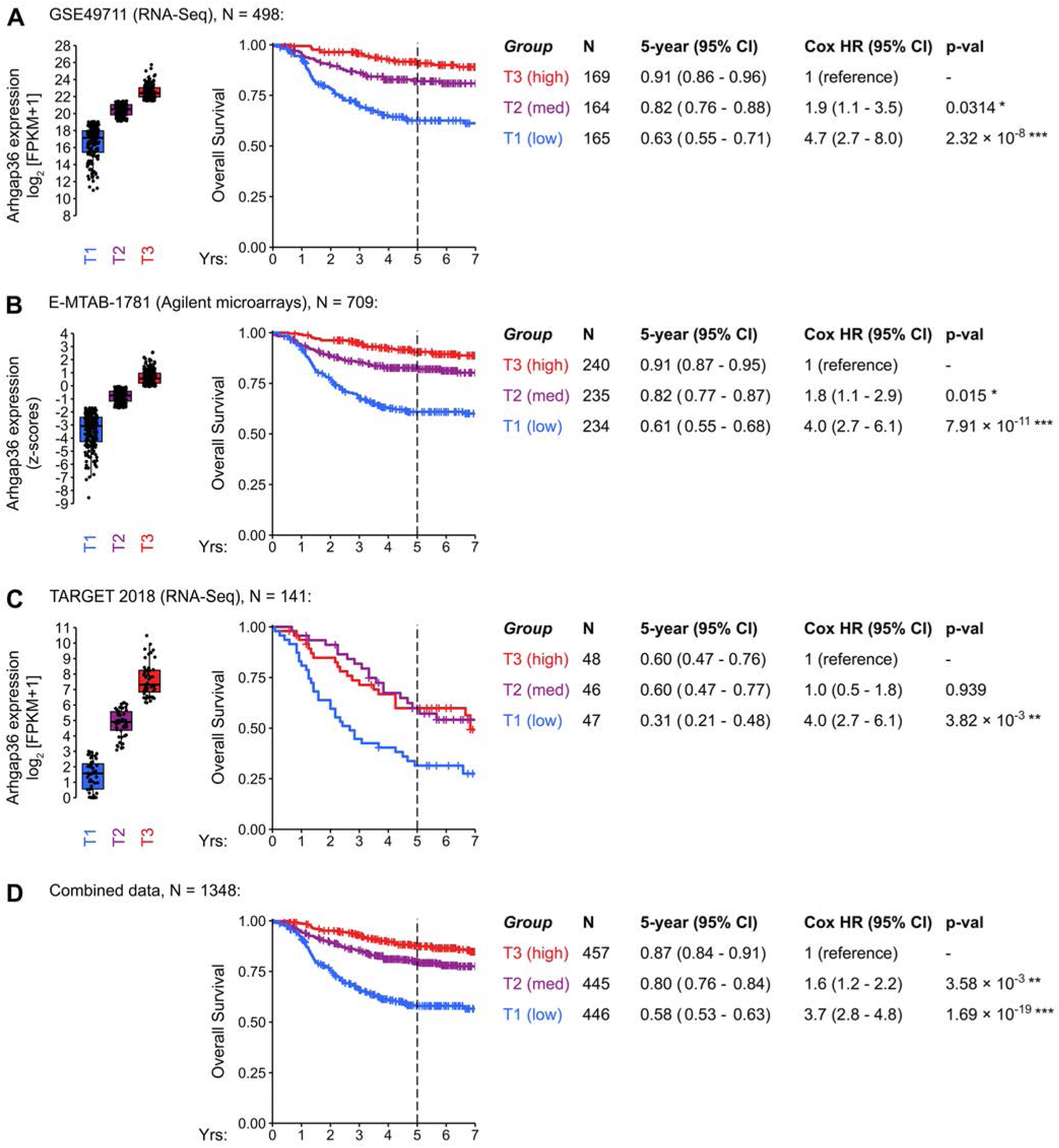
Overall survival of neuroblastoma patients stratified by Arhgap36 expression levels. (**A-C**) Kaplan-Meier plots show poor overall survival with low Arhgap36 expression across three independent neuroblastoma datasets. The significant reduced 5-year survival for cases with low levels of Arhgap36 expression is evident from the range of Hazard Ratios (2.7 – 8.0). Panel (**D**) shows the same analysis once the data are merged into a single dataset comprising n=1348 individuals (HR 2.8 – 4.8). [Cases stratified into terciles of Arhgap36 expression as shown in boxplots: first tercile (T1) “low”; second (T2) “medium”; third (T3) “high”].

## Discussion

Here we show that the PKA antagonist Arhgap36 is a central gene dysregulated by Foxc1 overexpression, and that this linkage provides new perspectives in Foxc1’s contributions to malignancy. Foxc1’s transcriptional regulation of Arhgap36, and the consequent inhibition of PKA, alters the equilibrium between full length transcriptional GLI activators and truncated transcriptional repressors, weakening regulation and increasing activity of a pathway fundamental to development. It is therefore logical that a transcription factor identified to enhance Arhgap36 expression is one that induces Hh activity in multiple tissues during development [13–15]. In some contexts, Hedgehog functions as a morphogen that confers positional information, with a gradient of activity specifying individual cell types and behaviors required for establishment of complex tissue patterns [39–41]. Increased gene dosage from *FOXC1* copy number variation causes phenotypes indicative of impaired Hedgehog signaling, including cerebellar hypoplasia and anomalous facial skeletal development [42–45]. Consequently, Foxc1’s ability to induce Arhgap36 expression establishes a new mechanism by which the transcription factor may modulate Hh pathway activity. This has particular relevance for oncology, where dysregulated Hh expression makes major contributions and larger fold increases in FOXC1 expression occur.

A second finding concerns the breadth of alteration to Hh signaling. Foxc1’s induction of Arhgap36 depletes an evolutionary conserved negative regulator of Hh signaling (PKA) from the cytoplasm and base of the primary cilium. Such wide loss of PKA activity should release Gli transcription factors from phosphorylation-induced direct inhibition (in case of Gli2) and proteolytic processing (in case of Gli2 and Gli3). In parallel, Foxc1-induced depletion of PKAC predictably dysregulates the PKA substrate Sufu: a second major negative regulator of the Hh pathway, whose roles encompass development, disease, and notably tumour suppression. Sufu acts as a scaffold between PKA and Gli proteins, that facilitates their phosphorylation and proteolytic processing, while also directly inhibiting transcriptional activity of active Gli forms [35]. The reduced phosphorylation of Sufu and increased translocation of Sufu and Gli2 to the tip of the cilium represent alterations to fundamental features of Hh signal transduction. As Arhgap36 acts independently of Smo [18], another consequence of Foxc1-driven Arhgap36 expression observed in our experiments is acquired resistance of Hh pathway activity to Smo antagonists. Consequently, by altering activity of PKA, Foxc1 overexpression impairs central elements of the Hh pathway resulting in increased transcription of Hh-target genes and, via augmented Smo-independent activity, tumor resistance to *Smoothened* antagonists. Collectively, such properties are consistent with the overexpression of FOXC1 in multiple tumors being associated with more aggressive cancer phenotypes, drug resistance and metastasis.

Forkhead box transcription factors’ ability to access areas of closed chromatin is mediated by a conserved winged-helix DNA binding domain that is structurally similar to histones H1 and H5 [46]. This capacity to directly bind DNA targets in condensed chromatin enables pioneer factors to activate enhancers and, by opening condensed chromatin, provide access to transcription factors that lack pioneering ability. The closely correlated ChIP-seq peaks identified in this study, which contain consensus sequences for Foxc1 and Fos-Jun dimers, support coordinated transcription factor binding at the Arhgap36 locus. The consistent co-localisation of these transcription factors’ binding motifs, including in other placental mammals, accords with enrichment of FOSL2 and JUNB motifs at the FOXC1 ChIP-seq peaks noted in a previous study [3]. Together, such data support a model in which the overexpression of FOXC1, by initiating local opening of chromatin [47], makes the ARHAGP36 locus accessible to other transcription factors to induce gene expression. Such combinatorial control of Arhgap36’s transcriptional regulation is predicted to have heterogeneous effects on Hh pathway activity (Fig. S12), varying in tissue and cell-specific manners and being determined by levels of FOXC1 and FOS-JUN dimers.

FOXC1’s induction of Hh signaling via ARHGAP36 and PKA inhibition is expected to modify processes in both development and disease. Since our data were derived with retroviral constructs, where 10-20 fold increases in Foxc1 protein levels reiterate overexpression in FOXC1-associated tumours [2], we evaluated the relevance of the Foxc1-Arhgap36 interaction in human malignancy. Analyses in neuroblastoma, a heterogeneous, life-threatening and common pediatric malignancy that originates from sympathoadrenal neural crest progenitors, revealed higher levels of ARHGAP36 expression were comparatively protective. The 2.8 to 8-fold greater five-year survival identified in three independent patient cohorts suggests ARHAGP36 may represent an informative prognostic factor. Because neuroblastoma treatment regimens frequently impact the development of surviving children [48], the ability to stratify by prognosis may facilitate treatment choice and minimize morbidity for a portion of patients. Although the mechanism(s) mediating this effect remain to be explored, this observation is consistent with the requirement of Hh activity for many tumours to proliferate [49, 50] and for maintenance of stem cell self-renewal, including that of cancer stem cells [51].

In summary, dysregulated expression of *FOXC1* is well-implicated in malignancy, with expanding evidence that elevated *FOXC1* levels are associated with increased mortality in breast and hepatic cancer and malignancies as diverse as AML and low grade glioma [4, 52–54]. In contrast with such progress, determining how FOXC1 promotes malignancy has proved more elusive. This work establishes that overexpression of FOXC1, at levels consistent with FOXC1-associated malignancies, transcriptionally activates Arhgap36 expression which in turn dysregulates multiple fundamental aspects of Hh pathway activity. The presented data further demonstrate that ARHGAP36 levels are predictive of five-year survival in a common pediatric malignancy and identify a biological characteristic that merits further investigation. Overall, this study provides additional mechanistic insight into FOXC1’s capacity for stimulating malignancy.

## Data Availability Statement

All data generated and/or analysed in this study are available from the corresponding author upon reasonable request. The RNA-sequencing and ChIP-sequencing data have been deposited to the NCBI GEO database with the following accession numbers: GSE297719 (RNA-seq), GSE297865 (ChIP-seq).

## Additional Information

Competing Interest Declaration: The authors declare no competing interests.

## Supporting information

Experimental proceedures and supplemental methods

## Acknowledgements

The authors are grateful to the patients who have contributed to this study. The authors thank Dr. Fred Berry for numerous fruitful discussions that facilitated and helped guide this research. We also thank multiple colleagues for critically reviewing earlier versions of the manuscript. Funding was provided by the Canadian Institutes of Health Research (CIHR) (MOP-133658). Experiments were performed at the Cell Imaging Core (RRID:SCR_019200), which is supported by the University of Alberta Faculty of Medicine & Dentistry, University Hospital Foundation, Striving for Pandemic Preparedness – The Alberta Research Consortium, and Canada Foundation for Innovation.

---

[See attached .xlsx file]

**File S1. List of differentially expressed genes in NIH3T3 cells expressing Foxc1 or pLXSH empty vector control.**

Complete list of differentially expressed genes identified in NIH3T3 cells expressing Foxc1 or empty vector control [pLXSH] by RNA-sequencing (n = 3) and used for volcano plot (Figure 1A). n = 292 differentially expressed genes; 195 upregulated, 97 downregulated. Selection criteria: q-value (FDR-adjusted p-value) ≤ 0.01, absolute log2 ratio ≥ 1, signal cut-off ≥ 1 FPKM in either condition].

[See attached .xlsx file]

**File S2. Functional profiling of the list of differentially expressed genes from File S1.**

Functional profiling of the list of genes differentially expressed in Foxc1-expressing NIH3T3 cells vs empty vector control [pLXSH] was performed using gene set enrichment analysis (GSEA) platform Enrichr [https://maayanlab.cloud/Enrichr/]. The list was analysed against multiple gene set libraries and databases. Results demonstrate significant associations between differentially expressed genes and molecular pathways / biological processes / pathological conditions with previous implication of Foxc1, as well as Foxc1-specific cell and tissue expression. Overall, these results indicate that ectopic expression of Foxc1 in NIH3T3 affects functionally relevant targets. Examples include association with ocular and vascular disease [glaucoma p = 5.8 × 10^-4^; intraocular pressure p = 5.9 × 10^-3^; hypertension p = 5.7 × 10^-4^; myocardial ischemia p = 2.4 × 10^-4^; coronary artery disease p = 0.01], vascular development [regulation of angiogenesis p = 9.4 × 10^-5^], bone development [endochondral ossification 2.8 × 10^-4^], a number of mouse skeletal and renal phenotypes [MGI phenotypes]; multiple cancers and cancer invasiveness phenotypes [epithelial mesenchymal transition p = 7.7 × 10^-11^]. Relevant cell and tissue expression signatures [mesenchyme p = 3.4 × 10^-8^; mesenchymal stem cells p = 5.8 × 10^-4^; (mesenchymal) stromal cells p = 1.8 × 10^-6^; vasculature p = 2.7 × 10^-5^; neural crest p = 3.8 × 10^-4^; several skeletal tissues; differentiating osteoblasts p ≤ 1.4 × 10^-6^]. p-values are FDR-adjusted.

**Figure S1.**
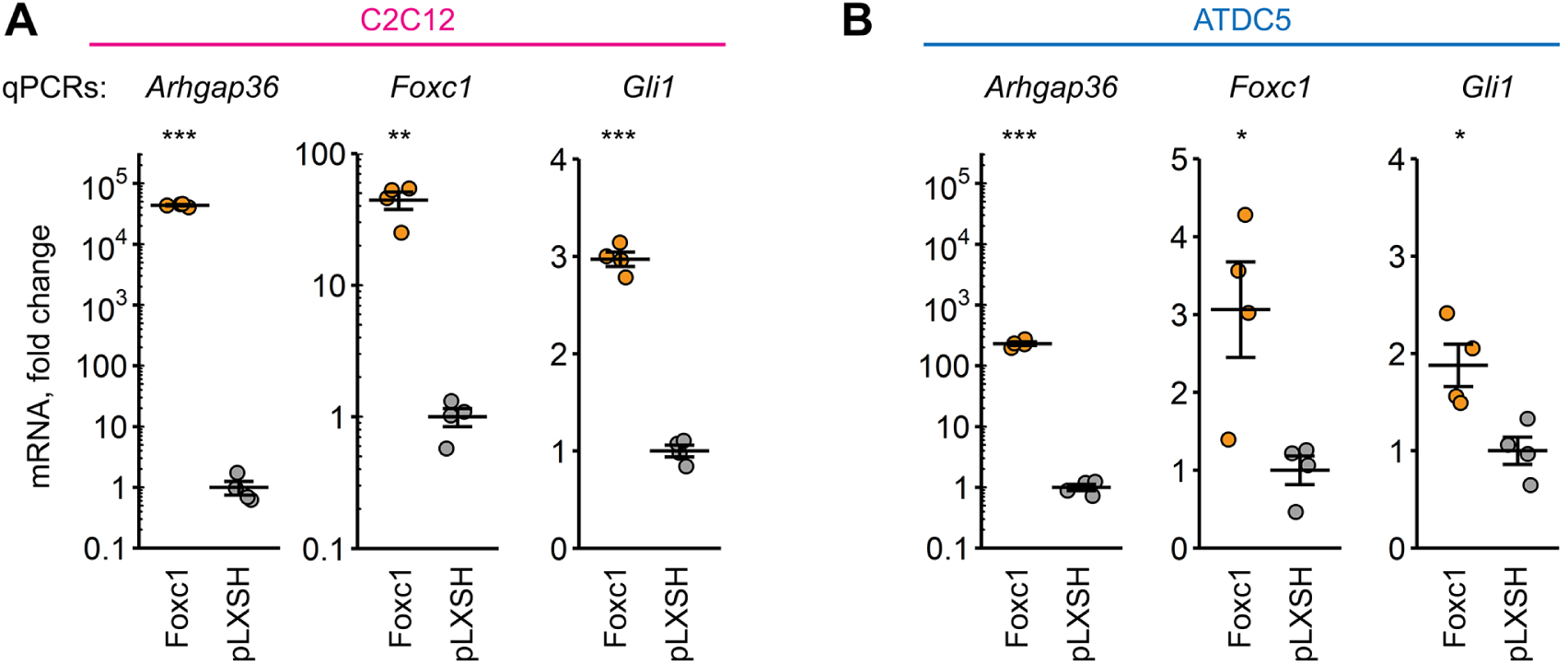
Foxc1 induces expression of Arhgap36 and activates Hedgehog signalling. (**A**,**B**) Stable overexpression of Foxc1 in C2C12 and ATDC5 cells induces strong increases in Arhgap36, and Gli1, mRNA levels.

**Figure S2.**
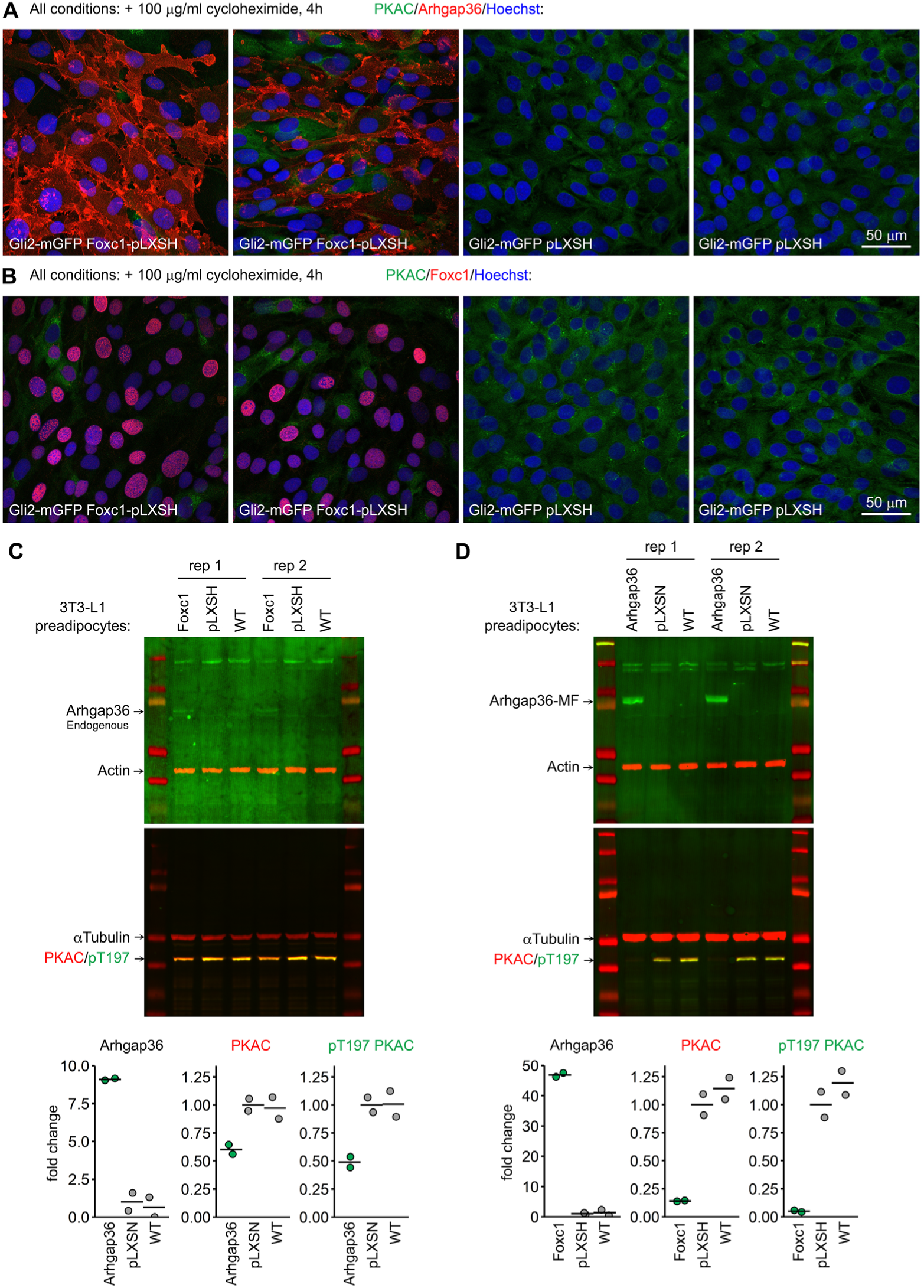
Foxc1-driven Arhgap36 reduces levels of protein kinase A catalytic subunit (PKAC) in Gli2-mGFP NIH3T3 cells and 3T3-L1 preadipocytes. (**A**) Immunofluorescent staining confirms substantial reduction of PKAC signal in Foxc1-expressing Gli2-mGFP NIH3T3 cells. Note correlation between positive Arhgap36 staining and PKAC loss in individual cells. (**B**) Immunofluorescent staining of Foxc1 and PKAC under the same conditions. Note nearly complete loss of PKAC signal in cells with high Foxc1 nuclear signal. (**C**) Foxc1 induced expression of endogenous Arhgap36 in 3T3-L1 cells is accompanied by 40-50% reduction in PKAC/pT197 PKAC protein levels. (**D**) Arhgap36-MycFLAG expression in 3T3-L1 cells strongly reduces PKAC, and catalytically active pT197 PKAC.

**Figure S3.**
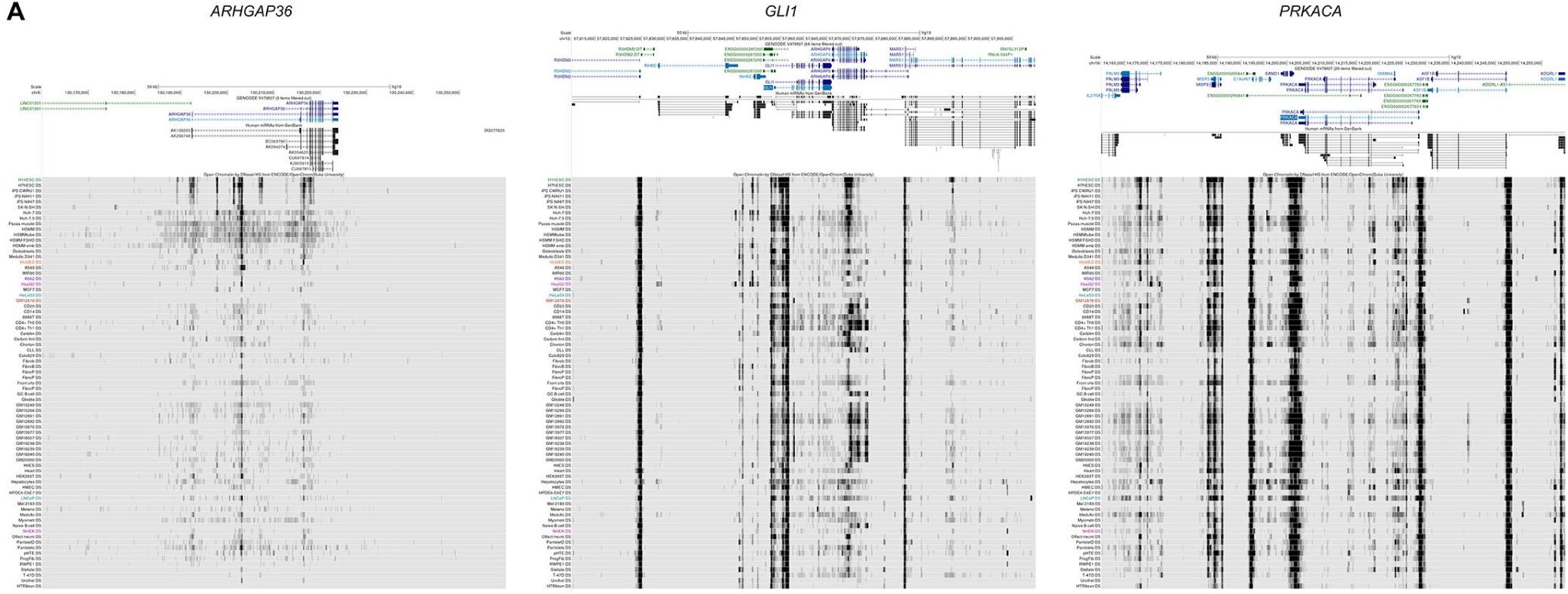
The *ARHGAP36* locus is located in a region of predominantly closed chromatin. (**A**) DNAse I hypersensistivity data from the UCSC genome browser for the 100 kb regions surrounding human *ARHGAP36*, and *GLI1* and *PRKACA* (for comparison), from diverse human cell lines (ENCODE). DNAse I HS is a marker for open chromatin that is commonly found at active cis-regulatory sequences including promoters and enhancers. At the *GLI1* and *PRKACA* loci, note the multiple, relatively uniform density signal peaks (appearing as dark vertical lines) for the majority of cell lines particularly immediately adjacent to transcription start sites (TSSs), illustrating “open chromatin” that is consistent with expression of both genes in a wide range of tissues. In contrast, strong DNAse I HS signal at the *ARHGAP36* locus is observed in the subset of the 75 cell lines of developmental origin marked with a purple vertical bar [Embryonic stem cells (H1hESC, H7hESC) and induced pluripotent stem (iPS CWRU1, iPS NIHi11, iPS NIHi7) cell lines]. The signal density for these is highest adjacent to several *ARHGAP36* TSSs, corresponding to the multiple known transcripts, with weak signal levels observed for most of the other cell lines.

**Figure S4.**
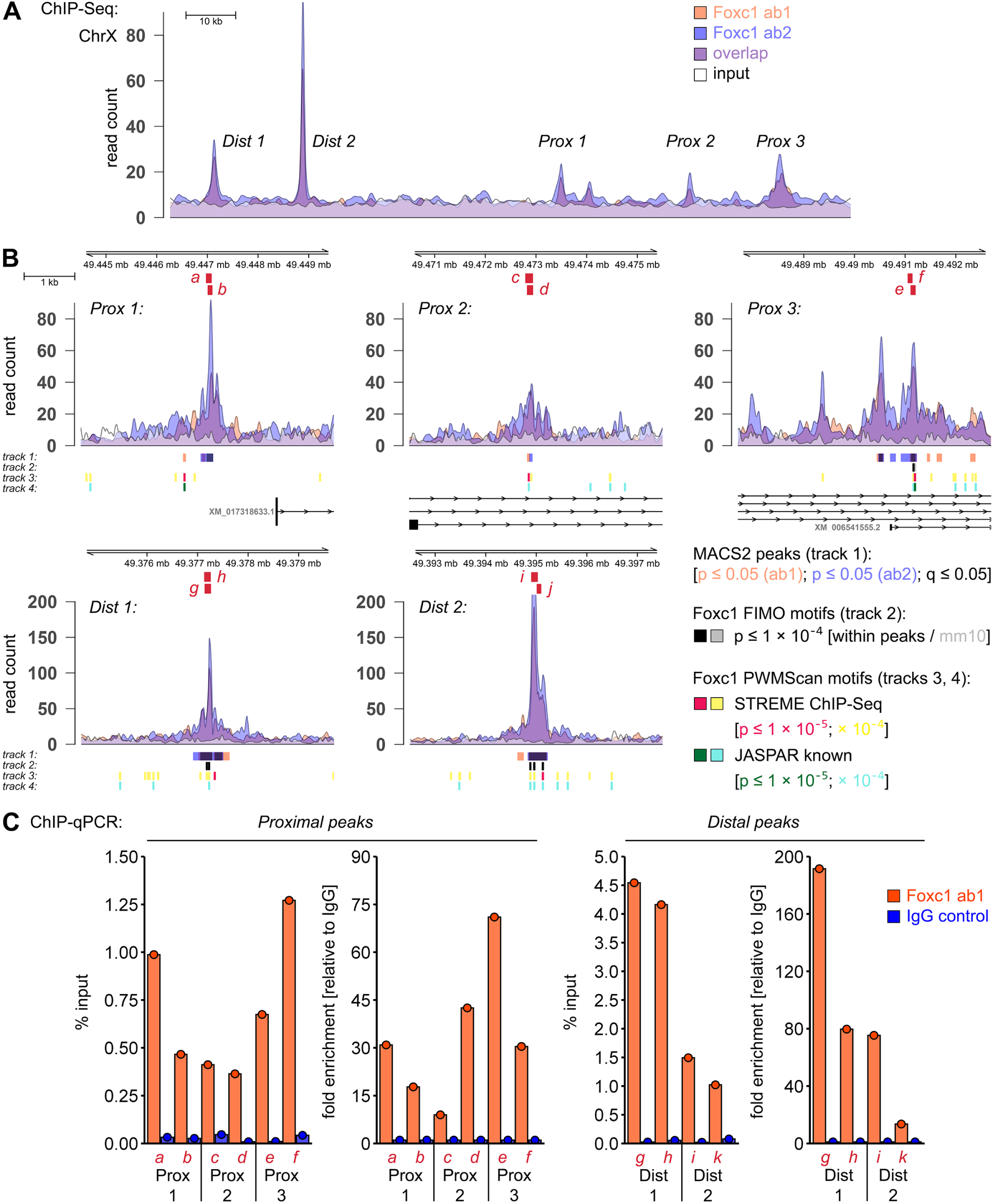
ChIP-qPCR validation of Foxc1 ChIP-Seq peaks in the vicinity of Arhgap36 gene. (**A**,**B**) Plots demonstrate read distribution for Foxc1 ChIP-Seq data from Figure 2 [**A**, entire locus (within ± 100 kb of Arhgap36 ORFs); **B**, individual peaks (5 kb regions); location of amplicons (a-j) that were analysed by ChIP-qPCR is indicated in red]. (**C**) ChIP-qPCR data confirm substantial enrichment of signal in all proximal and distal peaks identified by ChIP-Seq (N = 1).

**Figure S5.**
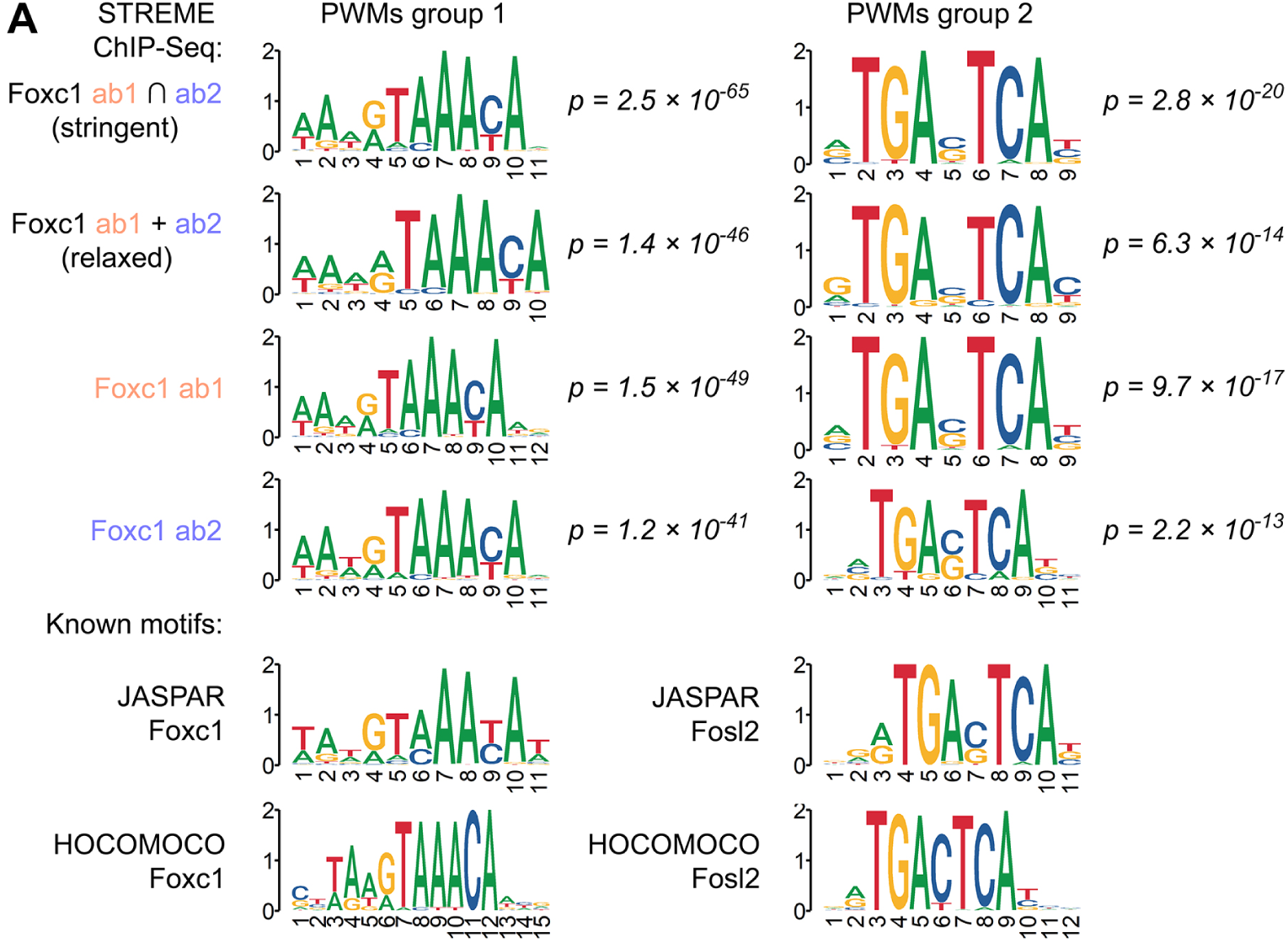
*De novo* motifs detected in Foxc1 ChIP-Seq peaks using STREME closely resemble known binding motifs of Foxc1 and Fosl2. (**A**) *De novo* motif analysis conducted using STREME tool identified two major groups of significantly enriched motifs (PWMs groups 1 and 2). Group 1 PWMs (ranked top #1 for intersected peaks) share high degree of similarity with known Foxc1 motifs, while Group 2 PWMs (ranked top #2 for intersected peaks) are highly similar to motifs of Fos-Jun family TFs, most prominently Fosl2. Notably, when analysed separately, ChIP-Seq data obtained using each of two anti-Foxc1 antibodies produce highly similar *de novo* PWMs. Known motifs of Foxc1 and Fosl2 transcription factors were retrieved from JASPAR and HOCOMOCO databases for comparison.

**Figure S6.**
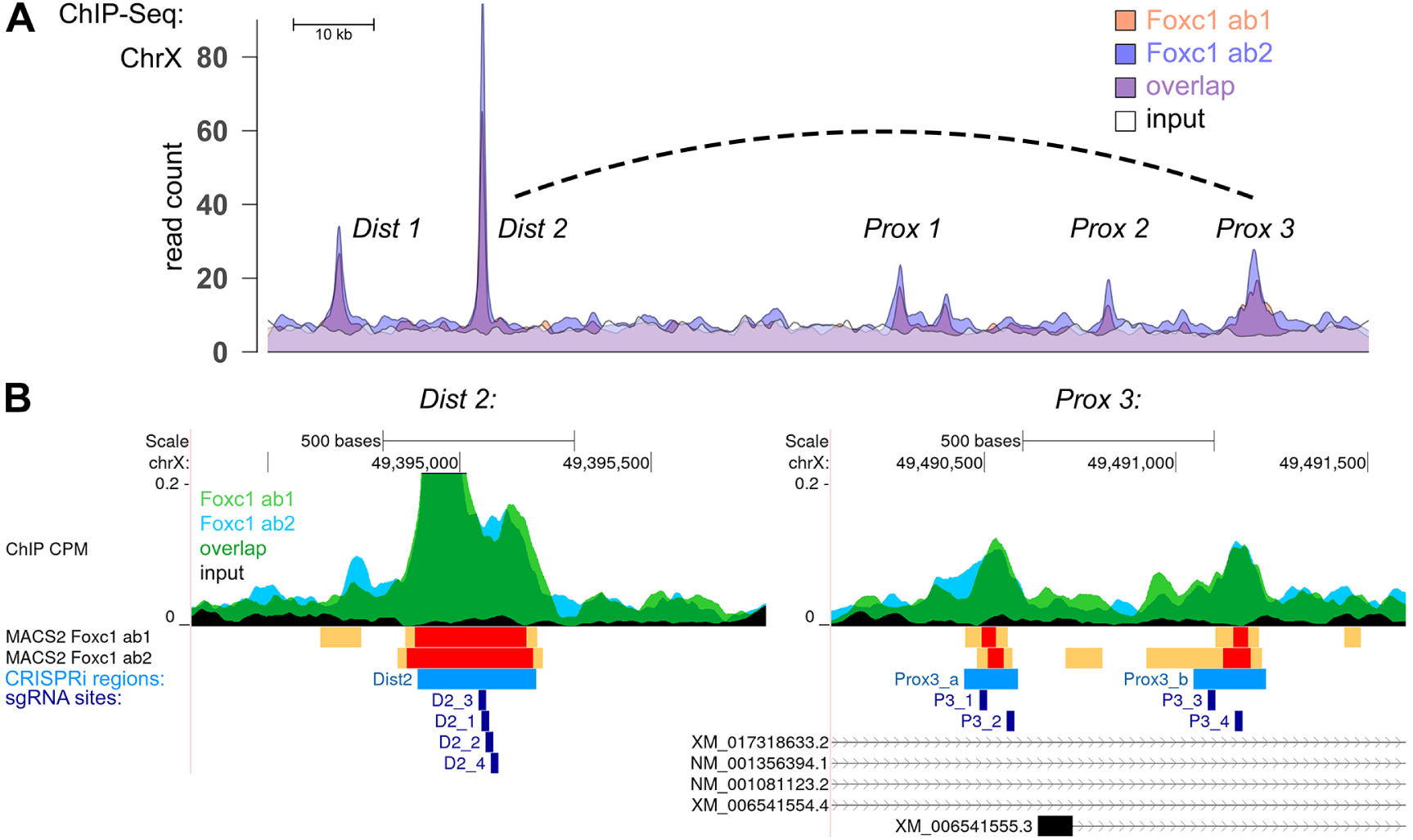
CRISPRi analysis of potential Foxc1-binding sites in the vicinity of Arhgap36 gene. (**A**) Plot shows read distribution for the ChIP samples and input chromatin control with five identified peaks (Prox-1 to Prox-3, Dist-1 and Dist-2). Dashed line demonstrates potential relationship between Prox-3 and Dist-2, as promoter and enhancer regions for murine *Arhgap36*. Panel (**B**) shows 1) ChIP-Seq coverage at both affected peaks; 2) nearby location of TSS for a major Arhgap36 isoform within the wide Prox-3 peak (approximately 80 – 110 bp downstream of narrow peak Prox-3a, and 500 bp upstream narrow peak Prox-3b); 3) location of individual sgRNA sites used for CRISPRi targeting.

**Figure S7.**
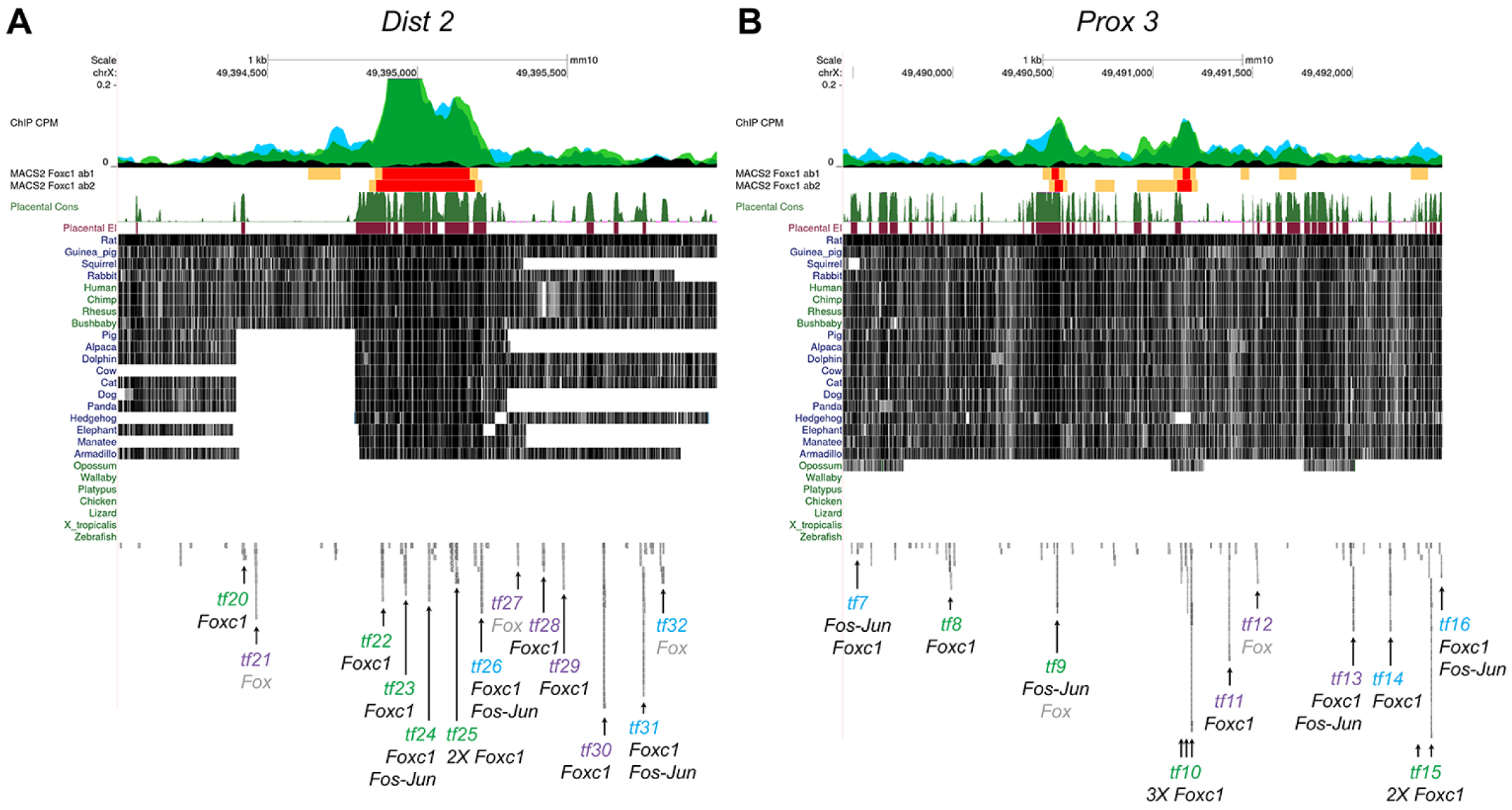
Conservation in Prox 3 and Dist 2 regions of Arhgap36 locus. (**A,B**) Representation of portions of the Arhagp36 locus corresponding to ChIP-seq peaks Prox-3 and Dist-2. Depicted in order are: ChIP-seq tracks with read coverage for two Foxc1 antibodies, UCSC-derived placental and non-placental mammal conservation data [Multiz alignment for representative vertebrate species, black lines], and distribution of predicted Fox and Fos-Jun transcription factor binding sites. Note, the heavy conservation in the Prox-3 region across all placental mammals, and in the Dist-2 region, that this is confined to the short interval detected by both Foxc1 antibodies. In addition, Prox-3 contains a cluster of 3 predicted binding sites for Foxc1, while Dist-2 includes a cluster of Foxc1 and Fos-Jun predicted binding motifs. [Placental mammal conservation by PhastCons (dark green “Placental Cons” track); placental mammal conserved elements (cherry “Placental El” track”)].

**Figure S8.**
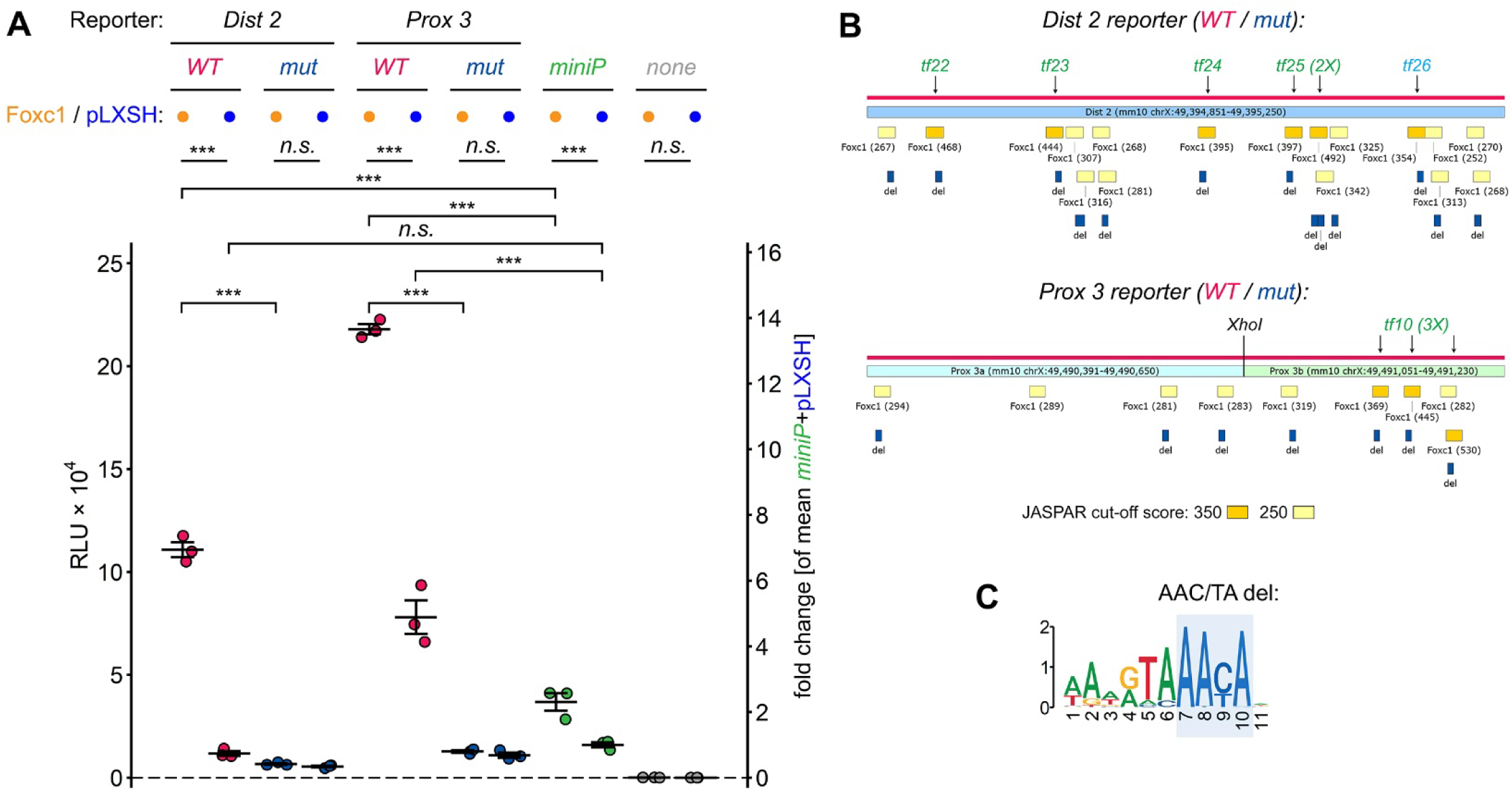
Two regions of the Arhgap36 locus sustain Foxc1-dependent transcription *in vitro*. (**A**) Luciferase reporter assays using wild-type or mutated Dist 2 and Prox 3 sequences upstream of a minimal promoter demonstrate that both wild-type reporters sustain significantly elevated transcriptional activity compared to the minimal reporter. Transcriptional activity is diminished by the absence of Foxc1 and abrogated by mutation of predicted Foxc1-binding motifs. (**B**) Depicts for each region locations of: (i) Foxc1-binding motifs predicted using stringent (orange) or relaxed (yellow) JASPAR criteria, (ii) the stringent Foxc1-binding motifs from Figure S7, and (iii) in blue, the core AAC/TA sequences in Foxc1’s binding motif (**C**) that were deleted in the mutated reporter constructs. [Wild-type (*WT*), mutated (*mut*), minimal promoter (*miniP*)]

**Figure S9.**
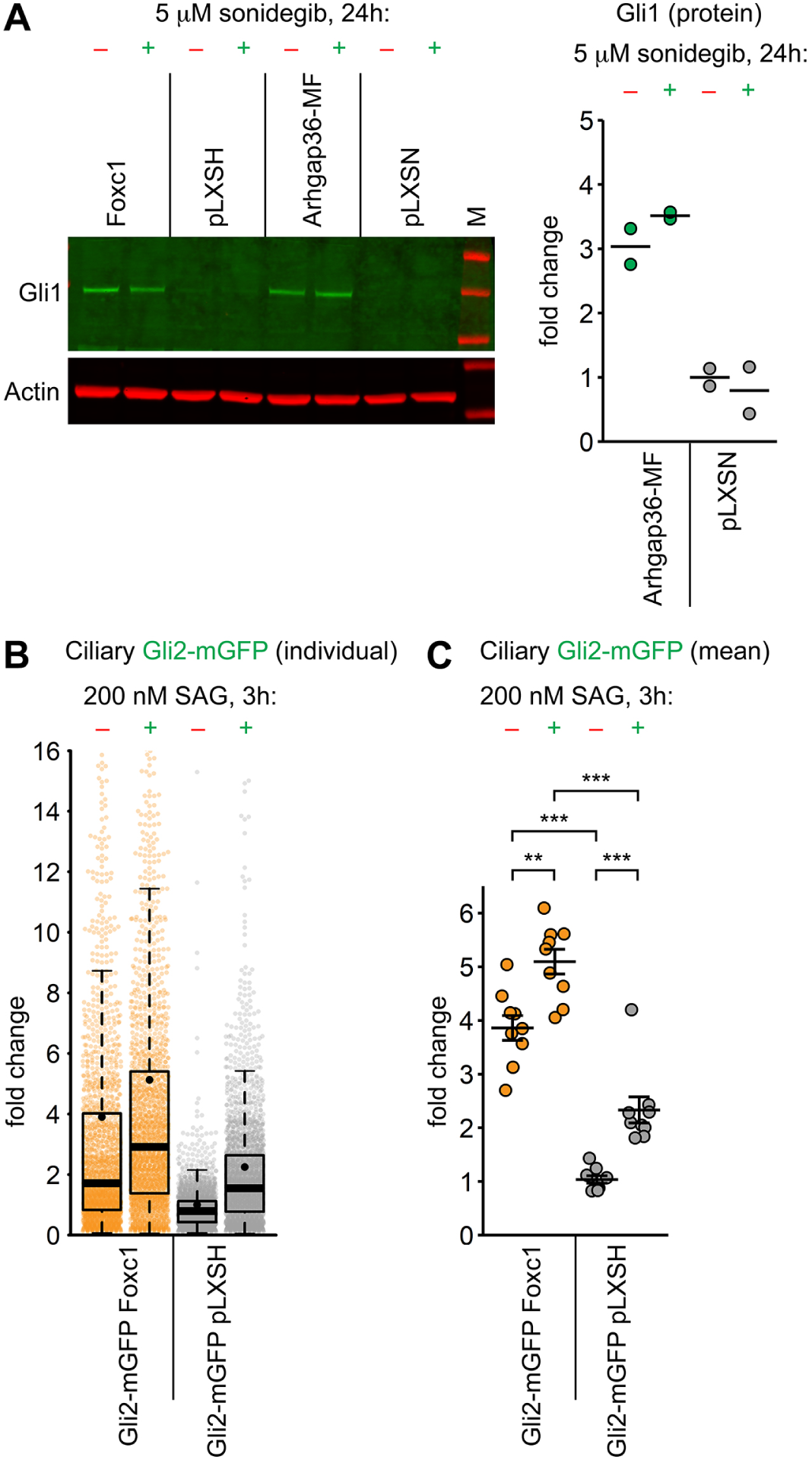
Foxc1 expression phenocopies Arhgap36-induced resistance to *Smoothened* inhibition and facilitates ciliary accumulation of Gli2. (**A**) Quantitative western blotting demonstrates that resistance to sonidegib inhibition observed in Foxc1-expressing Gli2-mGFP NIH3T3 cells recapitulates observations in NIH3T3 cells with ectopic expression of Myc-FLAG Arhgap36 [Arhgap36-MF]. (**B**,**C**) Accumulation of Gli2-mGFP at the tips of primary cilia in Gli2-mGFP NIH3T3 cells that express Foxc1, relative to relevant empty vector control [pLXSH]. Cells serum-starved for 20 h in media containing 0% FBS, 0.5% BSA, and next to stimulated with either vehicle or *Smoothened* agonist (SAG). Beeswarm/box plot **B** shows distribution of Gli2-mGFP intensity in individual cilia across all conditions tested [boxplot: box/whiskers – quartiles; black bar – median; black dot – mean; N = 9 combined experiments]. Mean ciliary Gli2-mGFP intensity values are provided in plot **C**.

**Figure S10.**
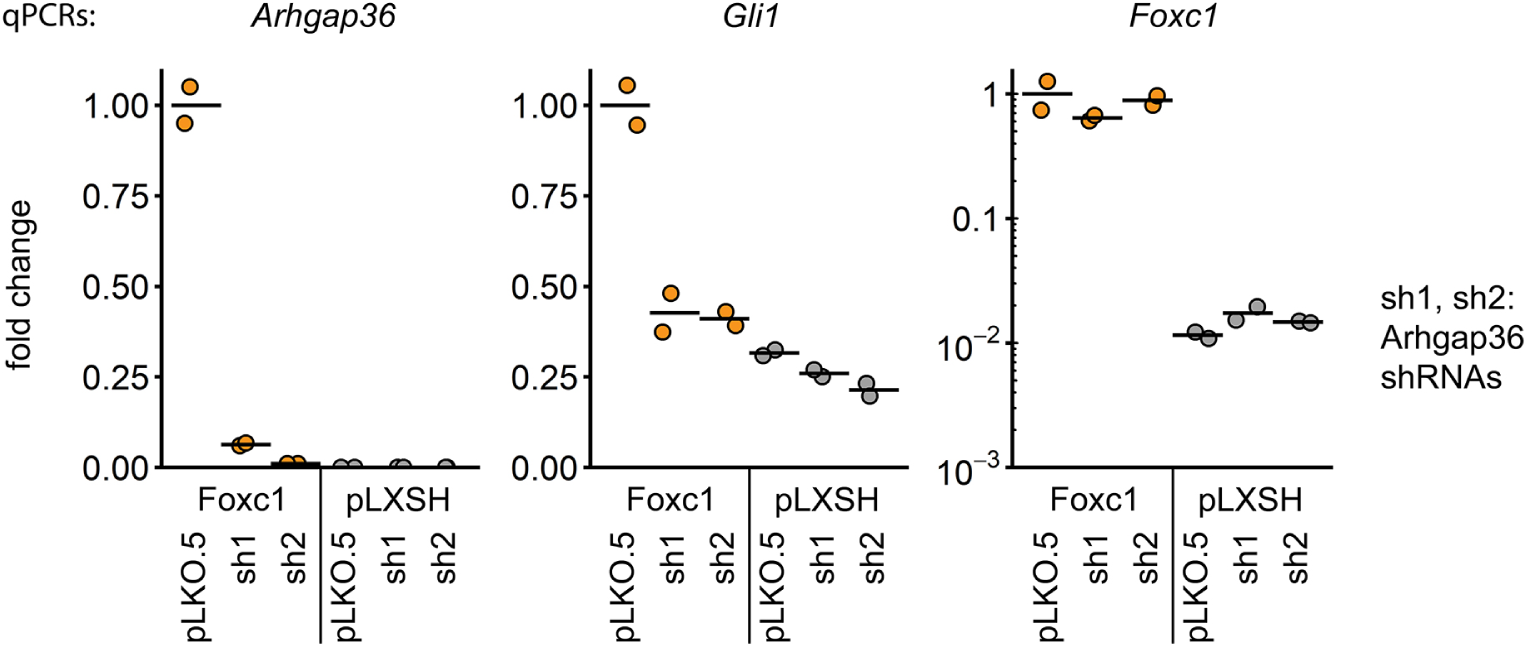
Foxc1-induced Gli1 expression is attributable to Arhgap36. qPCR analyses demonstrate substantial increase in Gli1 mRNA levels in NIH3T3 cells expressing Foxc1. This effect is reversed by shRNA-mediated knock-down of Arhgap36 expression.

**Figure S11.**
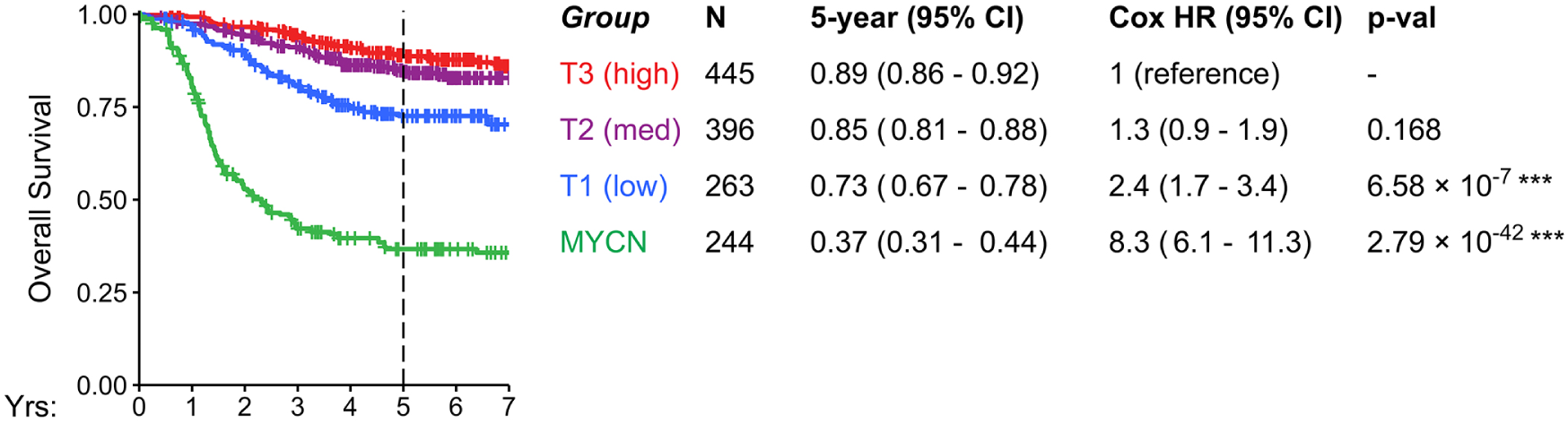
Overall survival of neuroblastoma patients stratified by Arhgap36 expression levels and MYCN amplification status. Analysis was performed on the GSE49711, E-MTAB-1781 and TARGET 2018 data merged into a single dataset comprising n=1348 individuals (Figure 8D). MYCN-amplified cases were segregated into a separate group. Kaplan-Meier plots illustrate three features: (i) best overall survival for patients with high Arhgap36 expression and without MYCN amplification, (ii) preserved lower overall survival rate among patients with low Arhgap36 expression and without MYCN amplification (HR 1.7 – 3.4), and (iii) predictably poor survival among high-risk MYCN-amplified cases (HR 6.1 – 11.3 compared to Arhgap36 high reference).

**Figure S12.**
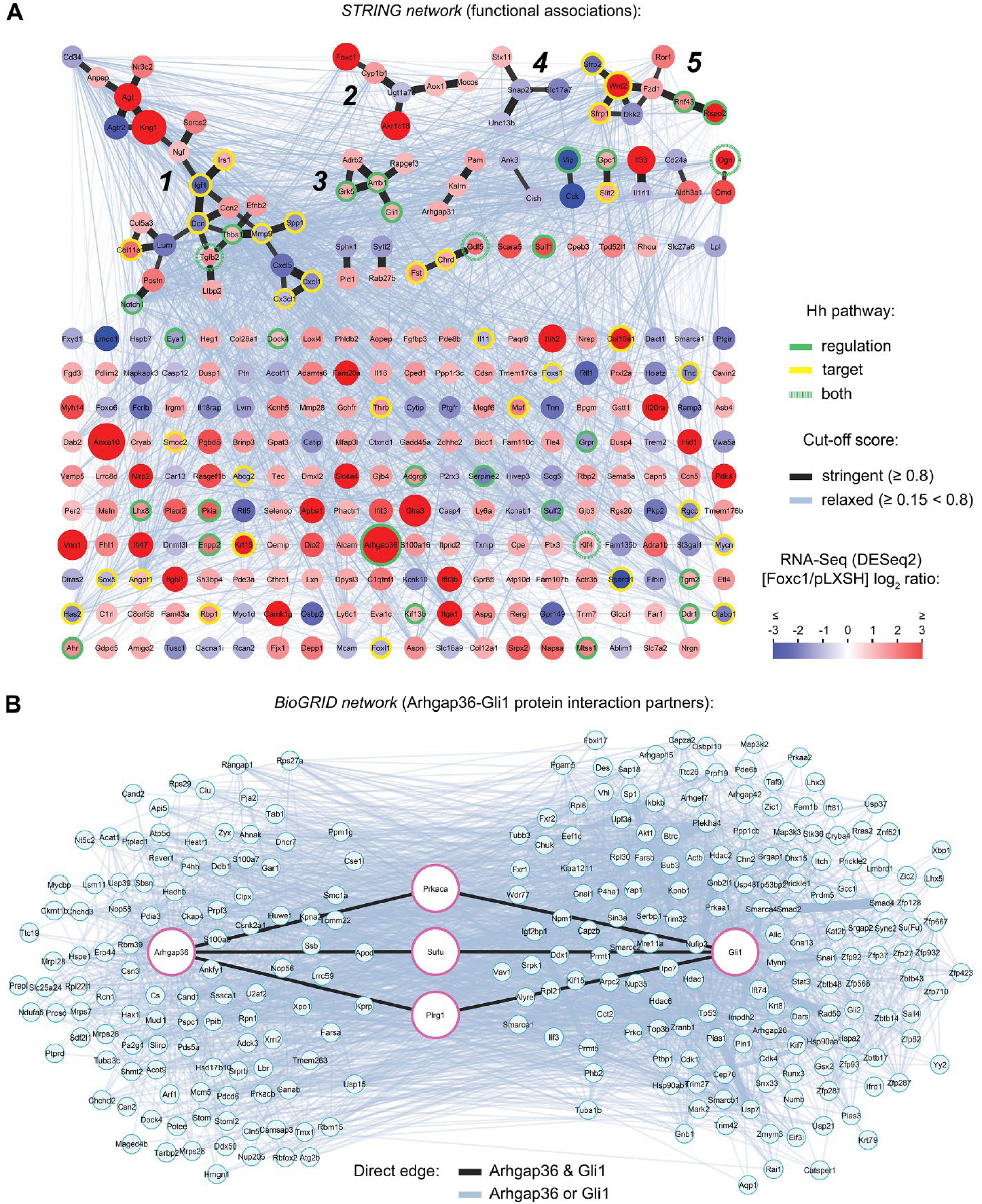
Network analysis of Foxc1-driven differential gene expression patterns. (**A**) Five highly functionally-associated clusters of 4 or more genes were identified using the STRING database [black lines; cutoff score ≥ 0.8]. Notably, cluster #3 contains Hh components Gli1, Grk5 and Arrb1. The individual node colour reflects differential gene expression (Fig. 1A); genes regulating or targeted by Hh signalling, depicted in green / yellow circles. (**B**) Protein interaction partners of Arhgap36 and Gli1 retrieved from BioGRID database reveals multiple interactions between those binding Gli1 or Arhgap36, however PKAC, Sufu and Plrg1 are the only proteins to directly interact with both.

## Notes

### Competing Interest Statement

The authors have declared no competing interest.

### Summary of Updates

the helpful reviews provided by eLife

