## Supplementary material for "Mechanistic insights into transcriptional regulation of ARHGAP36 expression identify a factor predictive of neuroblastoma survival": Experimental proceedures and supplemental methods

### Experimental procedures

*Plasmids, antibodies and other reagents*

Retroviral vector for stable expression of MycFLAG-tagged mouse *Arhgap36* was created by InFusion HD subcloning into pLXSN backbone. pLX303-ZIM3-KRAB-dCas9 and pLCKO plasmids for ZIM3 KRAB-dCas9 CRISPRi platform (Alerasool et al., 2020; PMID: 33020655) were obtained from Addgene. The sgRNA sequences targeting various regions in within Arhgap36 locus were generated by cloning of annealed primer duplexes into pLCKO vector using InFusion HD. All lentiviral shRNA plasmids from TRC mouse MISSION shRNA libraries were purchased from Millipore Sigma. Retroviral pLXSH vector for stable expression of *Foxc1* ORF was previously described [1]. Comprehensive lists of plasmids, antibodies, primers, as well as other reagents used in the study are provided in Supplemental methods.

*Cell lines*

NIH3T3 mouse fibroblasts and Gli2-mGFP NIH3T3 reporter cells stably expressing *Foxc1* ORF or pLXSH vector control were generated and maintained in presence of Geneticin and/or Hygromycin [1]. NIH3T3 cells stably expressing Arhgap36 MycFLAG-tagged protein or empty pLXSN vector control were generated by transduction with retroviral particles, followed by selection in 1200 µg/ml Geneticin. Resulting cell populations were maintained in media containing 400 µg/ml Geneticin. NIH3T3 and Gli2-mGFP NIH3T3 Foxc1/pLXSH cell lines stably expressing *Arhgap36* shRNAs were generated via lentiviral transduction and next selection in 2.5 ug/ml Puromycin. All NIH3T3 derivative cell lines were cultured in DMEM supplemented with 10% FBS. All cells were kept at 37ºC in 5% $CO_2$, in growth media containing 100 U/ml penicillin and 100 µg/ml streptomycin.

*RNA Sequencing*

RNA was isolated from NIH3T3 cells stably expressing *Foxc1* ORF or empty vector control using RNeasy Plus Mini Kit (Qiagen). Sample quality was checked using the Agilent 2100 Bioanalyzer to ensure high RNA integrity. Samples were then prepped following the standard protocol for the NEBnext Ultra II Stranded mRNA (New England Biolabs). Sequencing was performed at the BRC Sequencing Core (University of British Columbia, Vancouver, BC) on the Illumina NextSeq500 with paired end 43 × 43 bp reads. Obtained data were de-multiplexed using Illumina's bcl2fastq2. De-multiplexed read sequences were then aligned to the Mus musculus (mm10) reference sequence, and analysed for differential expression using DESeq2 and Cufflinks, through bioinformatics apps available on Illumina Sequence Hub (basespace.illumina.com).

*ChIP-sequencing*

Cross-linked chromatin was prepared and immunoprecipitated from Gli2-mGFP NIH3T3 reporter cells stably expressing *Foxc1* ORF using Simple ChIP plus Enzymatic Chromatin IP Kit (New England Biolabs), with two different anti-Foxc1 antibodies. Enriched chromatin was reverse-crosslinked, purified and sequenced together with input control on Illumina MiSeq 300 with paired end 61 × 61 bp reads. Sequencing was performed at the BRC Sequencing Core (University of British Columbia, Vancouver, BC). Peak calling was done with MACS2 software on Galaxy platform (usegalaxy.org). De novo motif detection in the ChIP peaks was conducted using STREME workflow from MEME Suite (meme-suite.org). Detailed ChIP protocol is provided in Supplemental methods.

*CRISPRi experiments*

K16 and K18 cell lines expressing ZIM3 KRAB domain fused with N terminus of dCas9 (clones K16 and K18) used in CRISPRi experiments were derived from Gli2-mGFP *Foxc1*-expressing NIH3T3 cells via lentiviral delivery of ZIM3-KRAB-dCas9 construct, followed by clonal selection in media containing 1.25 µg/ml Blasticidin. Resulting clonal cell lines were maintained in the presence of µg/ml Blasticidin, 80 µg/ml Hygromycin and 500 µg/ml Geneticin. To achieve CRISPR interference at various regions of *Arhgap36* locus, pools of 3 to 4 sgRNAs per each targeted peak region identified in ChIP-seq experiments, alongside with non-targeting *Lacz* and *Luc* sgRNA controls, were packaged into lentiviral particles and delivered via transduction into K16 and K18 clonal cell lines expressing ZIM3-KRAB-dCas9. Following a week of transient selection for sgRNA expression in 2 µg/ml Puromycin, cells were processed for either mRNA isolation and subsequent qPCR analyses, or preparation of cell lysates for western blotting.

*RT-qPCR analyses*

Total RNA was isolated with RNeasy Plus Mini Kit (Qiagen), quantified and used for cDNA synthesis with Primescript RT Master Mix (Clontech). qPCR reactions were run with TB Green Premix Ex Taq (Tli RNAse H Plus) master mix (Clontech) on LightCycler 96 Instrument and analysed using LightCycler 96 Application (Roche Life Science). Primer sets are provided in Supplemental methods.

*Quantitative western blotting*

Cells were lysed in 1.5% SDS lysis buffer (50 mM Tris pH 7.5, 150 mM NaCl, 1 mM EDTA, 1.5% SDS) supplemented with either Halt Protease Inhibitor Cocktail or Pierce Protease Inhibitor Mini Tablets (Thermo), as well as phosphatase inhibitors (0.5 mM $Na_3VO_4$, 5 mM NaF, 10 mM β-glycerophosphate). Resulting lysates were passed through QIAshredder columns (Qiagen). Obtained protein samples were normalised using BCA Protein Assay Kit (Thermo), resolved by SDS-PAGE (NuPage 4-12% Bis-Tris gels, Invitrogen), transferred to Immobilon-FL PVDF membranes (EMD Millipore) and blocked with Intercept Blocking Buffer, TBS (Li-Cor). Membranes were next incubated with relevant primary antibodies, followed by IRDye-conjugated secondary antibodies. Resulting membranes were scanned with Odyssey Imaging System (Li-Cor). Protein levels were quantified using Odyssey Application Software (Li-Cor).

*Immunofluorescent staining and quantification*

Cells seeded on optic bottom 24-well plates or 8-well slides (Ibidi; Cellvis) were treated as indicated in each experiment and fixed in Dent’s fixative for 30 min at room temperature. Following one wash with 1 ml D-PBS (Thermo-Fisher), blocking was performed in a blocking solution [TBS, 0.02% or 0.2% Triton X-100, 2 - 5% horse serum] for 1 hour at room temperature. Cells were next incubated with primary antibodies diluted in 100 - 150 ul blocking solution (2% horse serum) per well (see Supplemental methods for dilutions), overnight at 4C. Next day, cells were washed in TBSTx [TBS with 0.02% or 0.2% Triton X-100] on a rocker platform, and incubated with secondary antibodies (1:1000 for all secondary antibodies used) and Hoechst 33258 (2 µg/ml) diluted in 100 - 150 ul blocking solution, in the darkness, for 1 h at room temperature. Afterwards, cells were washed TBSTx, then in TBS. Finally, TBS was added to each well and cells were visualised by confocal microscopy. Images were collected using Zeiss LSM 700 laser scanning confocal microscope. Ciliary accumulation of Gli2-mGFP and Sufu, as well as numbers of cleaved-PARP1 positive cells were quantified by automated image analysis using Cell Profiler software [2].

*Omics datasets and expression dataset analyses*

RNA-seq and ChIP-seq data generated by this study have been deposited in NCBI's Gene Expression Omnibus (GEO) and are accessible through GEO Series accession numbers GSE297719 (RNA-seq), GSE297865 (ChIP-seq). Publicly available gene expression datasets for three neuroblastoma patient cohorts were retrieved from NCBI's GEO (GSE49711; part of GSE47792) [3], EMBL-EBI ArrayExpress (E-MTAB-1781) [4] and cBioPortal for Cancer Genomics (Pediatric Neuroblastoma TARGET, 2018) [5]. To define risk groups for each gene, normalized gene expression values within each dataset were ranked and split at 33.3% and 66.6% percentiles into three groups representing "low" (tercile T1), "medium" (tercile T2), and "high" (tericle T3) expression levels. Subsequent Kaplan-Meyer survival analyses and Cox proportional hazards analyses of the resulting data were performed using R language version 4.4.1 ("survival", "survminer" packages).

*Statistics*

Statistical analyses and plotting were performed using R language version 4.4.1 (The R Project for Statistical Computing). Analyses of statistical significance ($P < 0.05$) of differences between multiple groups were performed using one-way or two-way ANOVA followed up by Tukey's post-hoc test, differences between two groups – using Student's t-test, as indicated for specific experiments. Barplots and line plots show: individual data points for independent experimental replicates (circles), mean values (horizontal lines) ± SEM (errorbars) where applicable. Box-whisker plots (Figs. 6B, 7B) show quartiles, median (black horizontal lines) and mean (black dots) values. Boxplots (Figs. 8, S11) show quartiles, median (black horizontal limes) and individual expression data points for each sample. Significance codes ***$P < 0.001$, **$P < 0.01$, *$P < 0.05$.

### Supplemental methods

#### *ChIP-qPCR analyses*

Preparation of enriched chromatin samples for ChIP-PCR analyses was performed the same way, except immunuprocepitation was performed with anti-Foxc1 antibody alongside non-specific normal IgG control. Resulting samples were analysed by qPCR using pairs of primers designed to individual peak regions identified by ChIP-sequencig.

#### *Detailed ChIP protocol*

Cross-linked chromatin was prepared and immunoprecipitated from Gli2-mGFP NIH3T3 reporter cells stably expressing *Foxc1* ORF using Simple ChIP plus Enzymatic Chromatin IP Kit (New England Biolabs). Unless indicated otherwise, formulation of all reagents used was according to the manufacturers instructions. In detail, 3 million cells were seeded per 15 cm dish and grown for 72 h in DMEM media supplemented with 10% FBS, 80 µg/ml Hygromycin and 500 µg/ml Geneticin. 24 h prior to cross-linking media were replaced with 30 ml fresh DMEM supplemented with 10% FBS. To crosslink proteins to DNA, 1873 ul of 16% paraformaldehyde (PFA) solution was added to each plate. After 10 min incubation at room temperature, 3 ml of 10X Glycine solution was added to each plate and incubated for additional 5 min at room temperature to quench the remaining PFA. Following two washes with 20 ml ice-cold PBS per plate, cells were scraped in 2 ml ice-cold PBS solution supplemented with 10 µl of 200X Protease Inhibitor Cocktail (PIC) and centrifuged at 2000 G for 5 min, at 4°C. For preparation of nuclei and chromatin digestion, the cell pellets were re-suspended in 2 ml of 1X buffer A supplemented with 1 µl 1M DTT and 10 µl of 200X PIC and incubated for 10 min on ice, with mixing every 3 min. Nuclei were next pelleted by centrifugation at 2000 G for 5 min, at 4°C. Resulting nuclear pellet was resuspended in 500 µl of 1X buffer B with 0.25 ul of 1M DTT, and 2.5 µl of Micrococcal Nuclease (MNase) was added to each tube to fragment DNA, and samples were incubated at for 20 min at 37°C with frequent mixing [optimisation of MNase quantities was performed beforehand according to the manufacturers protocol for optimisation of chromatin digestion, to achieve DNA fragmentation range of 200 – 800 bp]. Digestion was stopped by addition of 50 ul of 0.5 M EDTA to each sample following 2 min incubation on ice. Next, nuclei were pelleted by centrifugation at 16000 G for 1 min, at 4°C, and re-suspended in 325 ul of ChIP buffer per sample. A total of 650 ul sample lysates were combined, kept for 10 min on ice, and further sonicated with 4 sets of 10 second pulses, separated by 20 second breaks, using Model 60 Sonic Dismembrator (Fisher Scientific), set at ~ 10 Watt power output. Lysates were clarified by centrifugation 9400 G for 10 min, at 4°C, and chromatin-containing supernatants were transferred to new tubes. After analysis of digestion, chromatin concentration was normalised to 150 µg/ml, in ChIP buffer with PIC. Chromatin immunoprecipitation was performed by mixing 100 ul chromatin sample, 400 ul of 1X ChIP buffer with PIC and antibodies (Foxc1 CST – 3 ul/sample; Foxc1 Abcam – 2 ul [2 ug], Normal IgG – 2 ul/sample [2 ug], H3 positive control – 10 ul), and incubation overnight at 4°C with rotation. Next day, 30 ul of ChIP-grade Protein G Magnetic Beads were added to each tube, and samples were incubated for additional 2 h at 4°C with rotation. Beads were pelleted using magnetic separation rack, washed three times for 5 min at 4°C in 1 ml of 1X ChIP buffer (low salt wash), and then one time for 5 min at 4°C in 1 ml of high salt wash (1X ChIP buffer with extra 350 mM NaCl). Following the washes, chromatin was eluted in 150 ul of 1X ChIP Elution buffer per sample for 30 min at 65°C, with occasional gentle vortexing. After pelleting the beads, resulting immunoprecipitated supernatant containing eluted chromatin was transferred into new tubes. Reversal of cross-links in immunoprecipitated chromatin, as well as equal volumes of 2% chromatin input samples, was done by addition of 6 µl of 5M NaCl and 2 µl Proteinase K, followed by 2 h digestion at 65°C and DNA purification using spin columns according to kit manufacturers instructions.

*List of plasmids*

| Name | Notes | Source | Figures |
|---|---|---|---|
| Arhgap36-MF-pLXSN | Retroviral vector for expression of Myc-FLAG-tagged mouse Arhgap36. In-Fusion® HD cloning from mouse Arhgap36-Myc-DDK ORF clone (Origene MR222487); pLXSN linearized by EcoRI/BamHI digestion; PCR cloning primers:<br>F: TTCTCTAGGCGCCGGAATTCATGGCGTGGATGCTGGACTG<br>R: ACATTCCACAGCCGGATCCTTAAACCTTATCGTCGTCATCC | This study | |
| Foxc1-pLXSH (mouse) | Retroviral vector for expression of mouse Foxc1. Previously generated in our laboratory. | PMID: 39217245 | |
| FOXC1-pLXSH (human) | Retroviral vector for expression of human FOXC1. In-Fusion® HD cloning of from human pENTR221-FOXC1 WT ORF (generous gift from Dr. Mike Walter, University of Alberta, Edmonton, AB); pLXSH linearized by BamHI digestion; PCR cloning pairs:<br>F: TCGTTAACTCGAGGATCCATGCAGGCGCGCTACTCCGTG<br>R: CTTGTCGACAGATCTGGATCCTCAAAACTTGCTACAGTCGTAGACG | This study | |
| pLXSN,<br>pLXSH,<br>vPak-VGV, | Empty pLXSN and pLXSH vector backbones, vPak-VGV helper plasmid (and 293-VSV-G packaging cell line) were gifts from Dr. Morag Park (McGill University, Montreal, QC). | McGill University | |
| pLX303-ZIM3-KRAB-dCas9 | Lentiviral vector for mammalian expression of ZIM3 KRAB domain fused to the N terminus of dCas9. Used in CRISPRi. | Addgene #154472 | |
| pLCKO,<br>pLCKO-LacZ-sgRNA,<br>pLCKO-Luc-sgRNA | Empty pLCKO lentiviral backbone and pLCKO vectors for expression of non-targeting LacZ and Luciferase sgRNAs under U6 promoter. Used for CRISPRi. pLCKO, pLCKO-LacZ-sgRNA and pLCKO-Luciferase-sgRNA vectors were gifts from Dr. Jason Moffat (University of Toronto, Toronto, ON). | Addgene #73311 | |
| pLCKO-Arhgap36-sgRNA vectors | Lentiviral pLCKO vectors for expression sgRNA pools targeting various regions in vicinity of Arhgap36 locus: Dist 1 (D1 vector pool), Dist 2 (D2 vector pool), Prox l (P1 vector pool), Prox 2 (P2 vector pool), Prox 3 (P3 vector pool). Used for CRISPRi. In-Fusion® HD cloning of annealed primer duplexes; pLCKO vector was linearized by BfuAI digestion. Primer duplex pairs (F/R):<br>D1_1F: aaggacgaggtaccg ACATACCCTAGGATGCAATA gttttagagctagaa<br>D1_1R: ttctagctctaaaac TATTGCATCCTAGGGTATGT cggtacctcgtcctt<br>D1_2F: aaggacgaggtaccg GTGGTACAGCAAGTAAATAA gttttagagctagaa<br>D1_2R: ttctagctctaaaac TTATTTACTTGCTGTACCAC cggtacctcgtcctt<br>D1_3F: aaggacgaggtaccg AGCGCCCTTATTGCATCCTA gttttagagctagaa | This study | |

| | | |
|---|---|---|
| | D1_3R: ttctagctctaaaac TAGGATGCAATAAGGGCGCT cggtacctcgtcctt<br>D1_4F: aaggacgaggtaccg GTACCACTCTATCCACAAGG gttttagagctagaa<br>D1_4R: ttctagctctaaaac CCTTGTGGATAGAGTGGTAC cggtacctcgtcctt<br>D2_1F: aaggacgaggtaccg TTTCTCTAGTGGTATTCGGA gttttagagctagaa<br>D2_1R: ttctagctctaaaac TCCGAATACCACTAGAGAAA cggtacctcgtcctt<br>D2_2F: aaggacgaggtaccg GTATTCGGATGGTTGTATTT gttttagagctagaa<br>D2_2R: ttctagctctaaaac AAATACAACCATCCGAATAC cggtacctcgtcctt<br>D2_3F: aaggacgaggtaccg CCACTAGAGAAAATATTGTT gttttagagctagaa<br>D2_3R: ttctagctctaaaac AACAATATTTTCTCTAGTGG cggtacctcgtcctt<br>D2_4F: aaggacgaggtaccg TGTATTTGGGGGACTTGATT gttttagagctagaa<br>D2_4R: ttctagctctaaaac AATCAAGTCCCCCAAATACA cggtacctcgtcctt<br>P1_1F: aaggacgaggtaccg AGCCAAGAGAGTGTGTGTTT gttttagagctagaa<br>P1_1R: ttctagctctaaaac AAACACACACTCTCTTGGCT cggtacctcgtcctt<br>P1_2F: aaggacgaggtaccg GATACTCCAGAACTGTGCTT gttttagagctagaa<br>P1_2R: ttctagctctaaaac AAGCACAGTTCTGGAGTATC cggtacctcgtcctt<br>P1_3F: aaggacgaggtaccg TTGACATGCTCCTAATTTAA gttttagagctagaa<br>P1_3R: ttctagctctaaaac TTAAATTAGGAGCATGTCAA cggtacctcgtcctt<br>P2_1F: aaggacgaggtaccg CAACTGTGTCGAATTTGCAT gttttagagctagaa<br>P2_1R: ttctagctctaaaac ATGCAAATTCGACACAGTTG cggtacctcgtcctt<br>P2_2F: aaggacgaggtaccg ACACTGACTAAACAGAAAAT gttttagagctagaa<br>P2_2R: ttctagctctaaaac ATTTTCTGTTTAGTCAGTGT cggtacctcgtcctt<br>P2_3F: aaggacgaggtaccg TCGAATTTGCATTGGTGTGT gttttagagctagaa<br>P2_3R: ttctagctctaaaac ACACACCAATGCAAATTCGA cggtacctcgtcctt<br>P3_1F: aaggacgaggtaccg ACAATTCCGAGGGAAATCAA gttttagagctagaa<br>P3_1R: ttctagctctaaaac TTGATTTCCCTCGGAATTGT cggtacctcgtcctt<br>P3_2F: aaggacgaggtaccg TCATCAGCCCAAAAAGGAGA gttttagagctagaa<br>P3_2R: ttctagctctaaaac TCTCCTTTTTGGGCTGATGA cggtacctcgtcctt<br>P3_3F: aaggacgaggtaccg TAGATATAAGAAATGCTTGT gttttagagctagaa<br>P3_3R: ttctagctctaaaac ACAAGCATTTCTTATATCTA cggtacctcgtcctt<br>P3_4F: aaggacgaggtaccg AAAAAGAAAATATACTTCTT gttttagagctagaa<br>P3_4R: ttctagctctaaaac AAGAAGTATATTTTCTTTTT cggtacctcgtcctt | |
| psPAX2,<br>pMD2.G | Retroviral packaging plasmids used to package pLCKO, pLKO.1, pLKO.5 vectors. These plasmids were obtained through RNAi Screening Core, Li Ka Shing Institute of Virology, University of Alberta, Edmonton, AB). | University of Alberta |
| pLKO.1 | Empty shRNA vector control. Obtained through RNAi Screening Core, Li Ka Shing Institute of Virology, University of Alberta, Edmonton, AB). | University of Alberta |
| pLKO.5 | Empty shRNA vector control. Cat. #SHC201. | Millipore-Sigma |
| Arhgap36 sh1 (mouse) | Mouse Arhgap36 shRNA targeting vector (pLKO.5 backbone).<br>TRC clone ID: TRCN 0000 268750 | Millipore-Sigma |

| | | |
|---|---|---|
| Arhgap36 sh2 (mouse) | Mouse Arhgap36 shRNA targeting vector (pLKO.5 backbone). TRC clone ID: TRCN 0000 268695 | Millipore-Sigma |
| Arhgap36 sh1 (human) | Human Arhgap36 shRNA targeting vector (pLKO.5 backbone). TRC clone ID: TRCN 0000 421375 | Millipore-Sigma |
| Arhgap36 sh2 (human) | Human Arhgap36 shRNA targeting vector (pLKO.5 backbone). TRC clone ID: TRCN 0000 421905 | Millipore-Sigma |
| Arhgap36 sh3 (human) | Human Arhgap36 shRNA targeting vector (pLKO.5 backbone). TRC clone ID: TRCN 0000 416368 | Millipore-Sigma |
| Arhgap36 sh4 (human) | Human Arhgap36 shRNA targeting vector (pLKO.1 backbone). TRC clone ID: TRCN 0000 147647 | Millipore-Sigma |
| Arhgap36 sh5 (human) | Human Arhgap36 shRNA targeting vector (pLKO.1 backbone). TRC clone ID: TRCN 0000 148948 | Millipore-Sigma |

*List of antibodies*

| Target | Cat. # | Description | Source | Figure |
|---|---|---|---|---|
| Arhgap36 / ARHGAP36 | HPA002064 | Anti-ARHGAP36 rabbit polyclonal antibody; WB dilution 1:1000-1:5000 (recommended 1:5000; non-specific binding in western blots can be reduced by addition of 0.25% Triton X-100 to TBST); IF dilution 1:500 – 1:1200. | Millipore-Sigma, Atlas Antibodies | 1, 4, 5, S4, S9, S11 |
| β-actin | # 3700 | β-Actin (8H10D10) mouse monoclonal antibody; WB dilution 1:10000 – 1:20000. | Cell Signaling | 1, 4, 5, 6, S4, S7, S9, S11 |
| Gli1/ GLI1 | # 2534 | Gli1(V812) rabbit polyclonal antibody; WB dilution 1:500 – 1:1000. | Cell Signaling | 1, 5, S7 |
| Foxc1 / FOXC1 | # 8758 | Foxc1 (D8A6) rabbit monoclonal antibody; WB dilution 1:500-1:1000; IF dilution 1:250 – 1:500. | Cell Signaling | 1, 2, S3, S4, S5, S9 |
| Foxc1 | Ab227977 | Foxc1 (EPR20685) rabbit monoclonal antibody. | Abcam | 2, S3, S5 |
| IgG control | # 2729 | Normal rabbit IgG | Cell Signaling | S3 |
| PKAC | 610980 | Purified Mouse Anti-PKA[C]; WB dilution 1:2000; IF dilution: 1:500 – 1:2000. | BD Trans-duction | 3, S4, S9, S11 |
| pT197 PKAC | # 5661 | Phospho-PKA C (Thr197) (D45D3) rabbit mAb; WB dilution 1:2000; IF dilution: 1:250. | Cell Signaling | 3, S4, S9 |
| α-tubulin | # 3873 | α-tubulin (DM1A) mouse mAb; WB dilution 1:10000 – 1:20000. | Cell Signaling | 3, 4, S4, S9, S11 |
| dCas9 | # 14697 | Cas9 (*S.pyogenes*) (7A9-3A3) mouse mAb. WB dilution 1:1000. | Cell Signaling | 4 |
| Sufu | # 2520 | SUFU (C54G2) rabbit mAb; WB dilution 1:1000; IF dilution 1:250. | Cell Signaling | 6 |
| pS342 Sufu | # 11552 | Sufu (Phospho-Ser342) rabbit polyclonal antibody. WB dilution 1:1000. | SAB Signalway | 6 |
| Arl13b | Ab136648 | Anti-ARL13B [N295B/66] mouse monoclonal antibody. IF dilution 1:250. | Abcam, Neuromab | 6 |
| Arl13b / ARL13b | 17711-1-AP | Arl13b rabbit polyclonal antibody; IF dilution 1:500; same antibody conjugated with CF555 dye using Mix-n-Stain antibody labeling kit (Biotium, Cat. # 92254) can be used at IF dilution 1:200. | Proteintech | 7 |

| | | | | |
|---|---|---|---|---|
| γ-tubulin | T5326 | γ-tubulin mouse monoclonal antibody, clone GTU-88; IF dilution 1:500. | Millipore-Sigma | 7 |
| AF555 donkey anti-rabbit | A-31572 | Donkey anti-rabbit highly cross-adsorbed secondary antibody, Alexa Fluor 555; IF dilution 1:500 – 1:1000. | Thermo-Fisher | 1, 3, 6, 9, S4 |
| AF647 goat anti-mouse | A-21237 | F(ab')2-Goat anti-mouse cross-adsorbed secondary antibody, Alexa Fluor 647; IF dilution 1:500 - 1:1000. | Thermo-Fisher | 3, 6, S4 |
| 800CW goat anti-rabbit | 926-32211 | IRDye® 800CW goat anti-rabbit secondary antibody; WB dilution 1:15000 | Li-Cor | 1, 3, 4, 5, 6, S4, S7, S9, S11 |
| 680RD donkey anti-mouse | 925-68072 | IRDye® 680RD donkey anti-mouse secondary antibody; WB dilution 1:15000 | Li-Cor | 1, 3, 4, 5, 6, S4, S7, S9, S11 |

*List of ChIP-PCR primers*

| **Target** | **Pair** | **Sequence** | **Figure** |
|---|---|---|---|
| *Prox 1 (a)* | F<br>R | CCAAAGCACAGTTCTGGAGTA<br>GAGCCAAGAGAGTGTGTGTT | S3 |
| *Prox 1 (b)* | F<br>R | CCATTATTACAAGGCAATCGCTAAA<br>AAATGTTACCAGGCCAGAGC | S3 |
| *Prox 2 (c)* | F<br>R | CGACACAGTTGACAGATATCCA<br>GGAAATGCAGACTTTGTGTTTGT | S3 |
| *Prox 2 (d)* | F<br>R | TTCAGGGAAAGCTACTTAAA<br>GCTGGAAATGCAGACTT | S3 |
| *Prox 3 (e)* | F<br>R | AAGGTATGTGAGGAGAAGAGAA<br>GTAAACAAATGAGCAATGCTGT | S3 |
| *Prox 3 (f)* | F<br>R | CTCATGCTGTGCCTCAGTTA<br>TTCTCTTCTCCTCACATACCTTAAT | S3 |
| *Dist 1 (g)* | F<br>R | GCACAGAAATGTGGAAGGAATC<br>CCTTGTGGATAGAGTGGTACAG | S3 |
| *Dist 1 (h)* | F<br>R | GAGCTTGCAGCACAGAAATG<br>GGATAGAGTGGTACAGCAAGTAAA | S3 |
| *Dist 2 (i)* | F<br>R | CATTCTGCTTCCTACTGTTCAAA<br>GCTGATTCATCTCACTAATCATCAC | S3 |
| *Dist 2 (j)* | F | AGCTGTGATGATTAGTGAGATGA | S3 |

| | | |
|---|---|---|
| | R | CCAAATACAACCATCCGAATACC |

All primers were from IDT.

*List of qPCR primers*

| **Target** | **Pair** | **Sequence** | **Figure** |
|---|---|---|---|
| *Arhgap36* (mouse) | F<br>R | GAT CCA GAG TGC TCG CAT AAA<br>TTC AAG ACC TCG TGC ACA TC | 1, 4, 7, S1 |
| *Gli1* (mouse) | F<br>R | GACTTTCTGGTCTGCCCTTT<br>AGGAGGAAAGAGAGATCCTTCA | 4, 5, 7, S1 |
| *Foxc1* (mouse) | F<br>R | CTCAACGAGTGCTTCGTCAA<br>CTTCACTGCGTCCTTCTTCTT | 7, S1 |
| *ARHGAP36* *(human)* | F1<br>R1 | GATCCAGAGTGCACGCATAA<br>CTTCATCAGAAGGCTTGCTTTG | S9 |
| *ARHGAP36* *(human)* | F2<br>R2 | ATTCCTCCTGACAGCAACTTTA<br>CCTCGCAGTTCTCAGTGATT | S9 (SK-N-FI) |

All primers were from IDT.

*Other reagents*

| **Reagent** | **Cat. #** | **Description** | **Source** | **Figure** |
|---|---|---|---|---|
| Sonidegib | S2151 | Specific antagonist of *Smoothened* (Smo) | Selleckchem | 5, S7 |
| Cyclopamine | S1146 | Specific antagonist of *Smoothened* (Smo) | Selleckchem | 5 |
| SAG | S7779 | Specific agonist of *Smoothened* (Smo) | Selleckchem | 5, 6, S7 |
| Cycloheximide | S7418 | Eukaryote protein synthesis inhibitor | Selleckchem | S5 |
| Doxycycline hyclate | D5207 | Tetracycline group antibiotic, used for Tet-inducible shRNA expression | Millipore-Sigma | S11 |
