## Supplemental Table 1 for "Mechanistic insights into transcriptional regulation of ARHGAP36 expression identify a factor predictive of neuroblastoma survival"

**Table S1. List of differentially expressed genes in NIH3T3 cells expressing Foxc1 or pLXSH empty vector control.**

**Differentially expressed genes with known roles as either targets (t) or regulators (r) of the Hh signalling pathway:**

| | | | DESeq2: | | CuffDiff: | | Role | References / Pubmed IDs |
|---|---|---|---|---|---|---|---|---|
| **#** | **Gene** | **Locus** | **log2 ratio** | **q value** | **log2 ratio** | **q value** | | |
| 1 | Arhgap36 | chrX:49470449-49500250 | 11.22 | 1.31E-19 | 12.78 | 0.000483 | r | Rack et al., 2014 (PMID: 25024229) |
| 2 | Wnt2 | chr6:17988939-18030445 | 5.86 | 6.45E-41 | 5.82 | 0.000483 | t | Rankin et al., 2016 (PMID: 27320915) |
| 3 | Col10a1 | chr10:34303611-34418528 | 4.56 | 0.00E+00 | 4.59 | 0.000483 | t | Koyama et al., 2007 (PMID: 17507416); Yoshida et al., 2015 (PMID: 25808752); Amano et al., 2014 (PMID: 25028519) |
| 4 | Ogn | chr13:49464058-49652731 | 3.72 | 0.00E+00 | 3.71 | 0.000483 | t,r | Lin et al., 2023 (PMID: 36515316); Huang et al., 2013 (PMID: 23645682); Lu et al., 2015 (PMID: 26597826) |
| 5 | Krt15 | chr11:100131758-100135949 | 3.20 | 5.07E-24 | 3.18 | 0.000483 | t | Rittie et al., 2009 (PMID: 20050020); Garcia-Zaragoza et al., 2012 (PMID: 23000969). |
| 6 | Rspo2 | chr15:43020794-43170818 | 3.15 | 4.43E-46 | 3.19 | 0.000483 | r | Sam et al., 2007 (PMID: 17904116); Aoki et al., 2008 (PMID: 18067586); Wang et al., 2021 (PMID: 34273374) |
| 7 | Sulf1 | chr1:12692429-12860372 | 2.85 | 1.04E-136 | 2.85 | 0.000483 | r | Danesin et al., 2006 (PMID: 16687495); Wojcinski et al., 2011 (PMID: 21806980); Jiang et al., 2017 (PMID: 28490013) |
| 8 | Pkia | chr3:7366603-7445365 | 2.50 | 9.69E-80 | 2.52 | 0.000483 | r | Via regulation of PKA. Hoy et al., 2020 (PMID: 32830375); Kawakami and Nakanishi, 2001 (PMID: 11493567); Hammerschmidt et al., 1996 (PMID: 8598293) |
| 9 | Gdf5 | chr2:155941024-155945364 | 2.27 | 3.45E-40 | 2.28 | 0.000483 | t,r | Koyama et al., 2007 (PMID: 18083924); Niedermaier et al., 2005 (PMID: 15841179); Craft et al., 2013 (PMID: 23715552) |
| 10 | Col11a1 | chr3:114030539-114220326 | 2.24 | 9.04E-133 | 2.23 | 0.000483 | t | Delaine-Smith et al., 2021 (PMID: 34189438); Feldman et al., 2008 (PMID: 18515410) |
| 11 | Enpp2 | chr15:54838678-54920146 | 2.06 | 1.45E-17 | 2.06 | 0.000483 | r | Frisca et al., 2016 (PMID: 27883058) |
| 12 | Maf | chr8:115703252-115706894 | 1.88 | 5.68E-27 | 1.87 | 0.000483 | t | Kerr et al., 2012 (PMID: 22491411). Essential for lens formation and osteoblast dfifferentiation (PMID: 33762508). |
| 13 | Mtss1 | chr15:58941233-59082026 | 1.83 | 1.72E-11 | 1.86 | 0.000483 | r | Drummond et al., 2018 (PMID: 29945904); Bershteyn et al., 2010 (PMID: 20708589) |
| 14 | Tgfb2 | chr1:186623185-186705992 | 1.71 | 3.13E-50 | 1.71 | 0.000483 | t,r | Rowan et al., 2018 (PMID: 29945868); Alvarez et al., 2002 (PMID: 11934857); Tang et al., 2018 (PMID: 29891662) |
| 15 | Sfrp1 | chr8:23411501-23449632 | 1.69 | 1.53E-82 | 1.69 | 0.000483 | t | Borday et al., 2012 (PMID: 22899850); Gomes et al., 2014 (PMID: 24712566) |
| 16 | Ahr | chr12:35497978-35534989 | 1.68 | 6.54E-114 | 1.68 | 0.000483 | r | Flegel et al., 2022 (PMID: 36459434) |
| 17 | Rnf43 | chr11:87663086-87735539 | 1.68 | 2.23E-29 | 1.61 | 0.000483 | r | Wang et al., 2021 (PMID: 34273374) |
| 18 | Lhx8 | chr3:154306293-154330560 | 1.65 | 4.48E-16 | 1.65 | 0.000483 | r | Flandin et al., 2011 (PMID: 21658586) |
| 19 | Fst | chr13:114452261-114458951 | 1.59 | 5.16E-16 | 1.6 | 0.000483 | t | Eichberger et al., 2008 (PMID: 18319260); Son et al., 2015 (PMID: 25775517); Muthu et al., 2019 (PMID: 31488567) |
| 20 | Adgrg6 | chr10:14402584-14545036 | 1.58 | 1.59E-20 | 1.62 | 0.000483 | r | Cui et al., 2021 (PMID: 33629464 ); Bian et al., 2024 (PMID: 39236220) |
| 21 | Klf4 | chr4:55527136-55532475 | 1.57 | 1.96E-104 | 1.57 | 0.000483 | r,t | Takeuchi et al., 2018 (PMID: 30193838); Zeng et al., 2016 (PMID: 27237094) |
| 22 | Rbp1 | chr9:98422960-98446550 | 1.52 | 6.85E-104 | 1.51 | 0.000483 | t | Hager-Theodorides et al., 2009 (PMID: 19667090) |
| 23 | Thrb | chr14:17660959-18038088 | 1.49 | 7.78E-29 | 1.48 | 0.000483 | t | Ostasov et al., 2020 (PMID: 32456161) |
| 24 | Smoc2 | chr17:14279505-14404790 | 1.32 | 1.04E-45 | 1.32 | 0.000483 | t | Pazin and Albrecht, 2009 (PMID: 19842175); Lu et al., 2015 (PMID: 26597826) |
| 25 | Gpc1 | chr1:92831685-92860196 | 1.27 | 1.10E-100 | 1.27 | 0.000483 | r | Wilson and Stoecki, 2013 (PMID: 23931997); Aguirre-Tamaral et al., 2022 (PMID: 36163184) |
| 26 | Grk5 | chr19:60889748-61140840 | 1.27 | 1.94E-23 | 1.28 | 0.000483 | r | Franco et al., 2018 (PMID: 29969579) |
| 27 | Tgm2 | chr2:158116404-158146392 | 1.22 | 1.81E-23 | 1.21 | 0.000483 | r | Shi et al., 2021 (PMID: 33245424) |
| 28 | Chrd | chr16:20724453-20742384 | 1.21 | 1.76E-13 | 1.2 | 0.000483 | t | Mouri et al., 2018 (PMID: 29255029) |
| 29 | Slit2 | chr5:47983154-48306282 | 1.20 | 1.05E-53 | 1.2 | 0.000483 | t | Barresi et al., 2005 (PMID: 16033800); Farmer et al., 2008 (PMID: 18842816); |
| 30 | Gli1 | chr10:127321963-127341579 | 1.20 | 2.11E-05 | 1.23 | 0.000483 | r | Kopinke et al., 2021 (PMID: 32540122) |
| 31 | Irs1 | chr1:82233104-82291439 | 1.16 | 2.35E-49 | 1.16 | 0.000483 | t | Sui et al., 2024 (PMID: 38775809); Parathath et al., 2008 (PMID: 18755774) |
| 32 | Kif13b | chr14:64652530-64806296 | 1.14 | 1.77E-23 | 1.15 | 0.000483 | r | Schou et al., 2017 (PMID: 28134340); Morthorst et al., 2022 (PMID: 35673984) |
| 33 | Arrb1 | chr7:99535485-99606771 | 1.13 | 2.94E-18 | 1.14 | 0.000483 | r | Kovacs et al., 2008 (PMID: 18497258); Parathath et al., 2010 (PMID: 20935513); Miele et al., 2017 (PMID: 28716052) |
| 34 | Angpt1 | chr15:42424666-42676977 | 1.09 | 2.06E-13 | 1.09 | 0.000483 | t | Nakamura et al., 2010 (PMID: 20098680); Kim et al., 2014 (PMID: 24807580) |
| 35 | Ddr1 | chr17:35681566-35704139 | 1.06 | 2.62E-42 | 1.07 | 0.000483 | r | Villegas Villarroel et al., 2024 (PMID: 39714220) |
| 36 | Thbs1 | chr2:118111921-118127133 | 1.05 | 6.62E-55 | 1.05 | 0.000483 | r | Wang et al., 2013 (PMID: 24148348) |
| 37 | Dock4 | chr12:40446052-40846488 | 1.02 | 8.46E-11 | 1.03 | 0.000483 | r | Makihara et al., 2018 (PMID: 30078728) |
| 38 | Cx3cl1 | chr8:94772179-94782426 | -1.05 | 6.29E-38 | -1.05 | 0.000483 | t | Giammona et al., 2024 (PMID: 39090759) |
| 39 | Notch1 | chr2:26457901-26503822 | -1.06 | 6.67E-18 | -1.05 | 0.000483 | r | Stasiulewicz et al., 2015 (PMID: 25995356) |
| 40 | Foxs1 | chr2:152931897-152933208 | -1.08 | 0.000269378 | -1.07 | 0.000483 | t | Diao et al., 2018 (PMID: 30098229); Wang et al., 2019 (PMID: 30918291) |
| 41 | Il11 | chr7:4772377-4782857 | -1.09 | 2.30E-19 | -1.08 | 0.000483 | t | Li et al., 2023 (PMID: 36853961) |
| 42 | Sox5 | chr6:143828424-144209568 | -1.11 | 2.60E-22 | -1.13 | 0.000483 | t | Hojo et al., 2013 (PMID: 23423383) |
| 43 | Abcg2 | chr6:58596671-58692451 | -1.16 | 6.59E-33 | -1.18 | 0.000483 | t | Xu et al., 2015 (PMID: 26276718); Yu et al., 2017 (PMID: 28847472); Kukal et al., 2021 (PMID: 34586444) |
| 44 | Mmp9 | chr2:164948218-164955849 | -1.19 | 4.11E-08 | -1.2 | 0.000483 | t | Faiao-Flores et al., 2016 (PMID: 27748762) |
| 45 | Foxl1 | chr8:121127684-121130644 | -1.21 | 1.18E-07 | -1.19 | 0.000483 | t | Madison et al., 2009 (PMID: 19049965) |
| 46 | Mycn | chr12:12936092-12941836 | -1.29 | 6.85E-11 | -1.28 | 0.000483 | t | Singh et al., 2018 (PMID: 30315164); Wickstrom et al., 2013 (PMID: 22949014); Oliver et al., 2003 (PMID: 12777630) |
| 47 | Eya1 | chr1:14168957-14310199 | -1.44 | 1.07E-14 | -1.44 | 0.000483 | r | Eisner et al., 2015 (PMID: 25816987) |
| 48 | Grpr | chrX:163513903-163549736 | -1.52 | 7.19E-10 | -1.51 | 0.000483 | r | Castellone et al., 2015 (PMID: 24747971) |
| 49 | Spp1 | chr5:104435110-104441053 | -1.55 | 2.62E-166 | -1.54 | 0.000483 | t | Iwaya et al., 2024 (PMID: 38086515); Hou et al., 2021 (PMID: 33314622) |
| 50 | Sulf2 | chr2:166073898-166155683 | -1.65 | 8.78E-31 | -1.65 | 0.000483 | r | Nakamura et al., 2013 (PMID: 23740243); Jiang et al., 2017 (PMID: 28490013) |
| 51 | Rgcc | chr14:79288749-79301635 | -1.73 | 3.58E-12 | -1.74 | 0.000483 | t | Zhao et al., 2023 (PMID: 37552059) |
| 52 | H19 | chr7:142575529-142578146 | -1.73 | 1.09E-57 | -1.74 | 0.000483 | t | Chan et al., 2014 (PMID: 24141783) |
| 53 | Cxcl1 | chr5:90891244-90893115 | -1.74 | 7.80E-05 | -1.71 | 0.000483 | t | Zaki et al., 2022 (PMID: 35388502) |
| 54 | Tnc | chr4:63959784-64047015 | -1.76 | 1.50E-115 | -1.76 | 0.000483 | t | Ohki et al., 2020 (PMID: 32581832) |
| 55 | Has2 | chr15:56665626-56694546 | -1.80 | 8.75E-102 | -1.8 | 0.000483 | t | Liu et al., 2013 (PMID: 23313125) |
| 56 | Dcn | chr10:97479499-97518162 | -1.87 | 1.22E-164 | -1.87 | 0.000483 | t | Montano et al., 2018 (PMID: 29368411) |
| 57 | Crabp1 | chr9:54764747-54773110 | -1.92 | 9.81E-124 | -1.92 | 0.000483 | t | Lin et al., 2020 (PMID: 32527063) |
| 58 | Serpine2 | chr1:79794320-79858665 | -1.97 | 3.54E-122 | -1.97 | 0.000483 | r | Vaillant et al., 2007 (PMID: 17409116); Vaillant et al., 2015 (PMID: 25901736) |
| 59 | Sfrp2 | chr3:83766320-83774314 | -2.27 | 7.73E-131 | -2.27 | 0.000483 | t | Ingram et al., 2002 (PMID: 12444557); Tavella et al., 2006 (PMID: 16962305); Lu et al., 2015 (PMID: 26597826) |
| 60 | Igf1 | chr10:87859055-87937047 | -2.61 | 2.74E-59 | -2.8 | 0.000483 | t | Lu et al., 2015 (PMID: 26597826); Nakamura et al., 2010 (PMID: 20098680) |

| | | | | | | | | |
|---|---|---|---|---|---|---|---|---|
| 61 | Sparcl1 | chr5:104079108-104114088 | -3.39 | 3.70E-159 | -3.4 | 0.000483 | t | Le et al., 2025 (PMID: 39900499) |
| 62 | Vip | chr10:5639217-5647614 | -4.24 | 4.25E-88 | -4.22 | 0.000483 | r | Bensalma et al., 2019 (PMID: 30669581) |

**Table S1. List of differentially expressed genes in NIH3T3 cells expressing Foxc1 or pLXSH empty vector control.**

### CuffDiff results:

Selection criteria: q-value (FDR-adjusted p-value) ≤ 0.01, absolute log2 ratio ≥ 1, CuffLinks signal cut-off ≥ 1 FPKM in either condition.

| # | Gene | Locus | Status | log2 FPKM pLXSH | log2 FPKM Foxc1 | log2 ratio | q value | p value |
|---|---|---|---|---|---|---|---|---|
| 1 | Anxa10 | chr8:62057041-62123193 | OK | -10 | 2.79 | 12.79 | 0.000483 | 5.00E-05 |
| 2 | Arhgap36 | chrX:49470449-49500250 | OK | -10 | 2.78 | 12.78 | 0.000483 | 5.00E-05 |
| 3 | Kng1 | chr16:23058299-23082078 | OK | -10 | 1.47 | 11.47 | 0.000483 | 5.00E-05 |
| 4 | E330011O21I | chr16:78250862-78255454 | OK | -10 | 0.94 | 10.94 | 0.000483 | 5.00E-05 |
| 5 | Glra3 | chr8:55940824-56125352 | OK | -10 | 0.09 | 10.09 | 0.000483 | 5.00E-05 |
| 6 | Vnn1 | chr10:23894687-23905343 | OK | -2.22 | 4.2 | 6.42 | 0.000483 | 5.00E-05 |
| 7 | Akr1c18 | chr13:4132626-4150631 | OK | -1.24 | 4.65 | 5.89 | 0.000483 | 5.00E-05 |
| 8 | Wnt2 | chr6:17988939-18030445 | OK | -3.26 | 2.57 | 5.82 | 0.000483 | 5.00E-05 |
| 9 | Il33 | chr19:29925113-29960715 | OK | -4.06 | 1.05 | 5.11 | 0.00287 | 0.00035 |
| 10 | Itih2 | chr2:10094590-10130683 | OK | -2.2 | 2.68 | 4.88 | 0.000483 | 5.00E-05 |
| 11 | Agt | chr8:124556586-124569707 | OK | -2.03 | 2.84 | 4.87 | 0.000483 | 5.00E-05 |
| 12 | Foxc1 | chr13:31806645-31810635 | OK | 2.11 | 6.97 | 4.86 | 0.000483 | 5.00E-05 |
| 13 | Col10a1 | chr10:34303611-34418528 | OK | 0.34 | 4.93 | 4.59 | 0.000483 | 5.00E-05 |
| 14 | Il20ra | chr10:19712586-19760053 | OK | -1.57 | 2.54 | 4.1 | 0.000483 | 5.00E-05 |
| 15 | Ogn | chr13:49464058-49652731 | OK | 2.51 | 6.23 | 3.71 | 0.000483 | 5.00E-05 |
| 16 | Slc4a4 | chr5:88887259-89239656 | OK | -3.33 | 0.3 | 3.63 | 0.000483 | 5.00E-05 |
| 17 | Apba1 | chr19:23758875-23949597 | OK | 0.17 | 3.75 | 3.58 | 0.000483 | 5.00E-05 |
| 18 | Ifit3b | chr19:34607956-34613401 | OK | -3.43 | 0.1 | 3.53 | 0.00494 | 0.00065 |
| 19 | Itgbl1 | chr14:123660139-123974079 | OK | 0.67 | 4.16 | 3.49 | 0.000483 | 5.00E-05 |
| 20 | Itga1 | chr13:114958080-115101964 | OK | -1.16 | 2.19 | 3.35 | 0.000483 | 5.00E-05 |
| 21 | Ifi47 | chr11:49037659-49135387 | OK | -2.73 | 0.57 | 3.3 | 0.000483 | 5.00E-05 |
| 22 | Camk1g | chr1:193346345-193370282 | OK | -2.19 | 1.1 | 3.29 | 0.000483 | 5.00E-05 |
| 23 | Hid1 | chr11:115347708-115367719 | OK | -0.24 | 3 | 3.24 | 0.000483 | 5.00E-05 |
| 24 | Rspo2 | chr15:43020794-43170818 | OK | -1.33 | 1.85 | 3.19 | 0.000483 | 5.00E-05 |

| | | | | | | | | |
|---|---|---|---|---|---|---|---|---|
| 25 | Krt15 | chr11:100131758-100135949 | OK | -1.38 | 1.8 | 3.18 | 0.000483 | 5.00E-05 |
| 26 | Pdk4 | chr6:5483350-5496278 | OK | 2.66 | 5.83 | 3.17 | 0.000483 | 5.00E-05 |
| 27 | Fam20a | chr11:109672925-109722256 | OK | 0.17 | 3.22 | 3.05 | 0.000483 | 5.00E-05 |
| 28 | Nlrp2 | chr7:5298546-5360682 | OK | -2.48 | 0.48 | 2.96 | 0.000483 | 5.00E-05 |
| 29 | Myh14 | chr7:44605802-44670843 | OK | -2.5 | 0.36 | 2.86 | 0.000483 | 5.00E-05 |
| 30 | Sulf1 | chr1:12692429-12860372 | OK | 0.5 | 3.35 | 2.85 | 0.000483 | 5.00E-05 |
| 31 | C430002N11I | chr9:96765561-96774397 | OK | -2.6 | 0.24 | 2.84 | 0.000483 | 5.00E-05 |
| 32 | Rasgef1b | chr5:99217419-99252927 | OK | -1.72 | 1.09 | 2.81 | 0.000483 | 5.00E-05 |
| 33 | Omd | chr13:49464058-49652731 | OK | 0.1 | 2.9 | 2.8 | 0.000483 | 5.00E-05 |
| 34 | Pgbd5 | chr8:124369048-124433936 | OK | -0.09 | 2.7 | 2.78 | 0.000483 | 5.00E-05 |
| 35 | Scara5 | chr14:65666402-65764826 | OK | -1.79 | 0.97 | 2.76 | 0.000483 | 5.00E-05 |
| 36 | Dio2 | chr12:90724551-90738438 | OK | 1.18 | 3.9 | 2.73 | 0.000483 | 5.00E-05 |
| 37 | Ifit3 | chr19:34583528-34588982 | OK | -2.02 | 0.7 | 2.72 | 0.000483 | 5.00E-05 |
| 38 | Napsa | chr7:44572444-44586846 | OK | -2 | 0.53 | 2.53 | 0.000483 | 5.00E-05 |
| 39 | Pkia | chr3:7366603-7445365 | OK | 0.86 | 3.38 | 2.52 | 0.000483 | 5.00E-05 |
| 40 | Srpx2 | chrX:133908424-133932446 | OK | -0.33 | 2.14 | 2.48 | 0.000483 | 5.00E-05 |
| 41 | Plscr2 | chr9:92275601-92297752 | OK | -0.69 | 1.76 | 2.46 | 0.000483 | 5.00E-05 |
| 42 | Adra1b | chr11:43774604-43901237 | OK | -1.17 | 1.21 | 2.38 | 0.000483 | 5.00E-05 |
| 43 | Aldh3a1 | chr11:61208741-61218416 | OK | 1.34 | 3.63 | 2.3 | 0.000483 | 5.00E-05 |
| 44 | Gdf5 | chr2:155941024-155945364 | OK | 0.3 | 2.58 | 2.28 | 0.000483 | 5.00E-05 |
| 45 | Col11a1 | chr3:114030539-114220326 | OK | 1.03 | 3.27 | 2.23 | 0.000483 | 5.00E-05 |
| 46 | Ror1 | chr4:100095790-100442545 | OK | 0.69 | 2.85 | 2.16 | 0.000483 | 5.00E-05 |
| 47 | 8430408G22I | chr6:116633007-116673836 | OK | -1.76 | 0.39 | 2.15 | 0.00174 | 2.00E-04 |
| 48 | Nr3c2 | chr8:76902507-77243639 | OK | -1.94 | 0.21 | 2.15 | 0.000483 | 5.00E-05 |
| 49 | Tpd52l1 | chr10:31332379-31445921 | OK | -1.39 | 0.75 | 2.14 | 0.000483 | 5.00E-05 |
| 50 | C1qtnf1 | chr11:118428456-118454995 | OK | -0.08 | 2.05 | 2.12 | 0.000483 | 5.00E-05 |
| 51 | Postn | chr3:54361106-54391041 | OK | 4.91 | 7 | 2.09 | 0.000483 | 5.00E-05 |
| 52 | Enpp2 | chr15:54838678-54920146 | OK | -1.1 | 0.96 | 2.06 | 0.000483 | 5.00E-05 |
| 53 | Sorcs2 | chr5:36017180-36398139 | OK | -0.17 | 1.83 | 1.99 | 0.000483 | 5.00E-05 |
| 54 | Adamts6 | chr13:104287872-104494763 | OK | -0.95 | 0.98 | 1.93 | 0.000483 | 5.00E-05 |

| | | | | | | | | |
|---|---|---|---|---|---|---|---|---|
| 55 | Maf | chr8:115703252-115706894 | OK | -0.13 | 1.74 | 1.87 | 0.000483 | 5.00E-05 |
| 56 | Aspg | chr12:112106682-112127573 | OK | 0.13 | 1.99 | 1.86 | 0.000483 | 5.00E-05 |
| 57 | Fhl1 | chrX:56731760-56793346 | OK | -1.72 | 0.14 | 1.86 | 0.000483 | 5.00E-05 |
| 58 | Mtss1 | chr15:58941233-59082026 | OK | -1.83 | 0.03 | 1.86 | 0.000483 | 5.00E-05 |
| 59 | Wisp2 | chr2:163820833-163833147 | OK | 4.81 | 6.61 | 1.8 | 0.000483 | 5.00E-05 |
| 60 | Kcnh5 | chr12:74897216-75177332 | OK | -1.23 | 0.55 | 1.77 | 0.000483 | 5.00E-05 |
| 61 | Fam213a | chr14:40993739-41013775 | OK | 2.65 | 4.39 | 1.74 | 0.000483 | 5.00E-05 |
| 62 | Vamp5 | chr6:72368048-72380468 | OK | 1.35 | 3.08 | 1.73 | 0.000483 | 5.00E-05 |
| 63 | Msln | chr17:25748612-25754327 | OK | 2.33 | 4.05 | 1.72 | 0.000483 | 5.00E-05 |
| 64 | Tgfb2 | chr1:186623185-186705992 | OK | 0.83 | 2.54 | 1.71 | 0.000483 | 5.00E-05 |
| 65 | Aspn | chr13:49464058-49652731 | OK | 3.9 | 5.59 | 1.69 | 0.000483 | 5.00E-05 |
| 66 | Fjx1 | chr2:102449365-102451792 | OK | -0.63 | 1.06 | 1.69 | 0.000483 | 5.00E-05 |
| 67 | Sfrp1 | chr8:23411501-23449632 | OK | 2.49 | 4.18 | 1.69 | 0.000483 | 5.00E-05 |
| 68 | Ahr | chr12:35497978-35534989 | OK | 2.75 | 4.44 | 1.68 | 0.000483 | 5.00E-05 |
| 69 | Lhx8 | chr3:154306293-154330560 | OK | 0.38 | 2.03 | 1.65 | 0.000483 | 5.00E-05 |
| 70 | Loxl4 | chr19:42592278-42612806 | OK | 2.17 | 3.81 | 1.64 | 0.000483 | 5.00E-05 |
| 71 | 2610035D17I | chr11:113043905-113201838 | OK | -0.23 | 1.4 | 1.63 | 0.000483 | 5.00E-05 |
| 72 | Adgrg6 | chr10:14402584-14545036 | OK | -0.8 | 0.82 | 1.62 | 0.000483 | 5.00E-05 |
| 73 | Gm14005 | chr2:128298662-128429351 | OK | 1.57 | 3.19 | 1.62 | 0.000483 | 5.00E-05 |
| 74 | Rnf43 | chr11:87663086-87735539 | OK | 0.29 | 1.9 | 1.61 | 0.000483 | 5.00E-05 |
| 75 | Fst | chr13:114452261-114458951 | OK | 0.06 | 1.66 | 1.6 | 0.000483 | 5.00E-05 |
| 76 | Gadd45a | chr6:67035095-67080652 | OK | 3.01 | 4.62 | 1.6 | 0.000483 | 5.00E-05 |
| 77 | 2010111I01R | chr13:62964892-63431745 | OK | 2.79 | 4.37 | 1.58 | 0.000483 | 5.00E-05 |
| 78 | Alcam | chr16:52248995-52452997 | OK | 3.05 | 4.63 | 1.58 | 0.000483 | 5.00E-05 |
| 79 | Etl4 | chr2:20289912-20810535 | OK | 0.17 | 1.75 | 1.58 | 0.000483 | 5.00E-05 |
| 80 | Ly6a | chr15:74994876-74998031 | OK | 3.39 | 4.97 | 1.58 | 0.000483 | 5.00E-05 |
| 81 | S100a16 | chr3:90541222-90543151 | OK | -0.74 | 0.84 | 1.58 | 0.000483 | 5.00E-05 |
| 82 | Cped1 | chr6:21985909-22255606 | OK | 2.76 | 4.33 | 1.57 | 0.000483 | 5.00E-05 |
| 83 | Klf4 | chr4:55527136-55532475 | OK | 3.4 | 4.97 | 1.57 | 0.000483 | 5.00E-05 |
| 84 | Gjb4 | chr4:127351085-127354081 | OK | 0.76 | 2.33 | 1.56 | 0.000483 | 5.00E-05 |

| | | | | | | | | |
|---|---|---|---|---|---|---|---|---|
| 85 | Brinp3 | chr1:146495665-146902472 | OK | 0.92 | 2.46 | 1.55 | 0.000483 | 5.00E-05 |
| 86 | Cyp1b1 | chr17:79706952-79715041 | OK | 2.41 | 3.95 | 1.54 | 0.000483 | 5.00E-05 |
| 87 | Mfap3l | chr8:60632824-60676731 | OK | 0.52 | 2.07 | 1.54 | 0.000483 | 5.00E-05 |
| 88 | Ctgf | chr10:24595441-24598682 | OK | 6.6 | 8.13 | 1.53 | 0.000483 | 5.00E-05 |
| 89 | Efnb2 | chr8:8617438-8660773 | OK | 3.63 | 5.15 | 1.52 | 0.000483 | 5.00E-05 |
| 90 | Rbp1 | chr9:98422960-98446550 | OK | 4.78 | 6.29 | 1.51 | 0.000483 | 5.00E-05 |
| 91 | Nrep | chr18:33437018-33464029 | OK | 2.45 | 3.96 | 1.5 | 0.000483 | 5.00E-05 |
| 92 | Rerg | chr6:137054824-137170496 | OK | 0.2 | 1.69 | 1.49 | 0.000483 | 5.00E-05 |
| 93 | Tmem176b | chr6:48833810-48841374 | OK | 0.65 | 2.14 | 1.49 | 0.000483 | 5.00E-05 |
| 94 | Thrb | chr14:17660959-18038088 | OK | 0.42 | 1.9 | 1.48 | 0.000483 | 5.00E-05 |
| 95 | Actr3b | chr5:25760025-25850341 | OK | 0.7 | 2.17 | 1.47 | 0.000483 | 5.00E-05 |
| 96 | Dusp4 | chr8:34807609-34819894 | OK | 1.16 | 2.63 | 1.47 | 0.000483 | 5.00E-05 |
| 97 | Megf6 | chr4:154170712-154275721 | OK | -0.4 | 1.07 | 1.46 | 0.000483 | 5.00E-05 |
| 98 | Rbp2 | chr9:98490536-98509771 | OK | -0.14 | 1.32 | 1.46 | 0.0025 | 3.00E-04 |
| 99 | Tle4 | chr19:14448071-14598183 | OK | 1.55 | 2.98 | 1.44 | 0.000483 | 5.00E-05 |
| 100 | Cdsn | chr17:35552127-35557180 | OK | 3.26 | 4.69 | 1.43 | 0.000483 | 5.00E-05 |
| 101 | Dusp1 | chr17:26505590-26508472 | OK | 2.98 | 4.39 | 1.41 | 0.000483 | 5.00E-05 |
| 102 | Fzd1 | chr5:4753838-4758216 | OK | 4.95 | 6.36 | 1.41 | 0.000483 | 5.00E-05 |
| 103 | Irgm1 | chr11:48865248-48871346 | OK | 2.31 | 3.71 | 1.39 | 0.000483 | 5.00E-05 |
| 104 | Cpeb3 | chr19:37021290-37207471 | OK | 0.37 | 1.74 | 1.37 | 0.000483 | 5.00E-05 |
| 105 | Pam | chr1:97821093-98095632 | OK | 4.76 | 6.13 | 1.37 | 0.000483 | 5.00E-05 |
| 106 | Pde8b | chr13:95024104-95250304 | OK | 1.26 | 2.62 | 1.36 | 0.000483 | 5.00E-05 |
| 107 | Pld1 | chr3:27938679-28133362 | OK | -0.01 | 1.35 | 1.36 | 0.000483 | 5.00E-05 |
| 108 | Gchfr | chr2:119167787-119172389 | OK | 3.85 | 5.2 | 1.35 | 0.000483 | 5.00E-05 |
| 109 | Fam43a | chr16:30599722-30602797 | OK | 3.24 | 4.58 | 1.34 | 0.000483 | 5.00E-05 |
| 110 | Nrgn | chr9:37544492-37552745 | OK | -0.48 | 0.86 | 1.34 | 0.000483 | 5.00E-05 |
| 111 | Agpat9 | chr5:100846228-100899102 | OK | -0.75 | 0.58 | 1.33 | 0.000483 | 5.00E-05 |
| 112 | Il16 | chr7:83643062-83735490 | OK | -1.24 | 0.09 | 1.33 | 0.000483 | 5.00E-05 |
| 113 | Dpysl3 | chr18:43320978-43438286 | OK | 3.86 | 5.18 | 1.32 | 0.000483 | 5.00E-05 |
| 114 | Fam107b | chr2:3713457-3782134 | OK | 4.03 | 5.35 | 1.32 | 0.000483 | 5.00E-05 |

| | | | | | | | | |
|---|---|---|---|---|---|---|---|---|
| 115 | Smoc2 | chr17:14279505-14404790 | OK | 2.56 | 3.88 | 1.32 | 0.000483 | 5.00E-05 |
| 116 | Zdhhc2 | chr8:40423814-40484842 | OK | 1.97 | 3.29 | 1.32 | 0.000483 | 5.00E-05 |
| 117 | Fam110c | chr12:31073967-31079940 | OK | 1.4 | 2.71 | 1.31 | 0.000483 | 5.00E-05 |
| 118 | Sh3bp4 | chr1:89070461-89153793 | OK | 2.61 | 3.92 | 1.31 | 0.000483 | 5.00E-05 |
| 119 | Sdpr | chr1:51218385-51333779 | OK | 5.49 | 6.78 | 1.3 | 0.000483 | 5.00E-05 |
| 120 | Ltbp2 | chr12:84783211-84879755 | OK | 4.21 | 5.5 | 1.29 | 0.000483 | 5.00E-05 |
| 121 | Rgs20 | chr1:4909575-5070285 | OK | 2.63 | 3.91 | 1.29 | 0.000483 | 5.00E-05 |
| 122 | Grk5 | chr19:60889748-61140840 | OK | 1.48 | 2.76 | 1.28 | 0.000483 | 5.00E-05 |
| 123 | Sema5a | chr15:32244812-32696341 | OK | 1.5 | 2.79 | 1.28 | 0.000483 | 5.00E-05 |
| 124 | Fgd3 | chr13:49263109-49309208 | OK | 0.4 | 1.67 | 1.27 | 0.000483 | 5.00E-05 |
| 125 | Gpc1 | chr1:92831685-92860196 | OK | 5.61 | 6.88 | 1.27 | 0.000483 | 5.00E-05 |
| 126 | Cpe | chr8:64592550-64693040 | OK | 7.08 | 8.34 | 1.26 | 0.000483 | 5.00E-05 |
| 127 | Cryab | chr9:50751071-50756635 | OK | 5.09 | 6.35 | 1.26 | 0.000483 | 5.00E-05 |
| 128 | Pdlim2 | chr14:70164217-70177672 | OK | 3.44 | 4.7 | 1.26 | 0.000483 | 5.00E-05 |
| 129 | Dmxl2 | chr9:54365157-54501626 | OK | -0.42 | 0.82 | 1.25 | 0.000483 | 5.00E-05 |
| 130 | Lrrc8d | chr5:105699968-105815215 | OK | 0.91 | 2.16 | 1.25 | 0.000483 | 5.00E-05 |
| 131 | Phldb2 | chr16:45746230-45844378 | OK | 4.08 | 5.32 | 1.24 | 0.000483 | 5.00E-05 |
| 132 | Amigo2 | chr15:97244073-97385691 | OK | 3.27 | 4.5 | 1.23 | 0.000483 | 5.00E-05 |
| 133 | Arhgap31 | chr16:38598342-38713035 | OK | 1.42 | 2.65 | 1.23 | 0.000483 | 5.00E-05 |
| 134 | Gjb3 | chr4:127325234-127330836 | OK | 3.73 | 4.96 | 1.23 | 0.000483 | 5.00E-05 |
| 135 | Gli1 | chr10:127321963-127341579 | OK | -1.14 | 0.09 | 1.23 | 0.000483 | 5.00E-05 |
| 136 | Tec | chr5:72755717-72868448 | OK | -0.82 | 0.41 | 1.23 | 0.000483 | 5.00E-05 |
| 137 | Eva1c | chr16:90830858-90904885 | OK | 0.75 | 1.97 | 1.22 | 0.000483 | 5.00E-05 |
| 138 | Heg1 | chr16:33684465-33768195 | OK | 3.21 | 4.43 | 1.22 | 0.000483 | 5.00E-05 |
| 139 | Slc7a2 | chr8:40862366-40922070 | OK | 0.59 | 1.81 | 1.22 | 0.000483 | 5.00E-05 |
| 140 | Col12a1 | chr9:79598986-79718722 | OK | 4.42 | 5.64 | 1.21 | 0.000483 | 5.00E-05 |
| 141 | Tgm2 | chr2:158116404-158146392 | OK | 1.6 | 2.82 | 1.21 | 0.000483 | 5.00E-05 |
| 142 | Trim7 | chr11:48826137-48850195 | OK | -0.25 | 0.96 | 1.21 | 0.000483 | 5.00E-05 |
| 143 | Capn5 | chr7:98121558-98184799 | OK | 0.27 | 1.47 | 1.2 | 0.000483 | 5.00E-05 |
| 144 | Chrd | chr16:20724453-20742384 | OK | 0.58 | 1.78 | 1.2 | 0.000483 | 5.00E-05 |

| | | | | | | | | |
|---|---|---|---|---|---|---|---|---|
| 145 | Lxn | chr3:67430114-67475068 | OK | 6.6 | 7.8 | 1.2 | 0.000483 | 5.00E-05 |
| 146 | Ly6c1 | chr15:75044017-75048837 | OK | 2.25 | 3.46 | 1.2 | 0.000483 | 5.00E-05 |
| 147 | Rhou | chr8:123653928-123663880 | OK | 3.85 | 5.05 | 1.2 | 0.000483 | 5.00E-05 |
| 148 | Slit2 | chr5:47983154-48306282 | OK | 4.88 | 6.08 | 1.2 | 0.000483 | 5.00E-05 |
| 149 | AI606473 | chr3:154330820-154334687 | OK | 1.42 | 2.61 | 1.19 | 0.000483 | 5.00E-05 |
| 150 | Bicc1 | chr10:70925095-71159634 | OK | 4.85 | 6.03 | 1.18 | 0.000483 | 5.00E-05 |
| 151 | Col5a3 | chr9:20770049-20815034 | OK | 1 | 2.18 | 1.18 | 0.000483 | 5.00E-05 |
| 152 | Rab27b | chr18:69979130-70141605 | OK | 0.74 | 1.91 | 1.17 | 0.000483 | 5.00E-05 |
| 153 | Sepp1 | chr15:3270766-3280508 | OK | 2.84 | 4.01 | 1.17 | 0.000483 | 5.00E-05 |
| 154 | Adrb2 | chr18:62177712-62179981 | OK | -0.65 | 0.5 | 1.16 | 0.000483 | 5.00E-05 |
| 155 | Irs1 | chr1:82233104-82291439 | OK | 2.6 | 3.76 | 1.16 | 0.000483 | 5.00E-05 |
| 156 | Tmem176a | chr6:48841482-48845364 | OK | 0.23 | 1.39 | 1.16 | 0.000483 | 5.00E-05 |
| 157 | Asb4 | chr6:5383385-5433021 | OK | 2.68 | 3.84 | 1.15 | 0.000483 | 5.00E-05 |
| 158 | Gdpd5 | chr7:99381548-99460984 | OK | -0.65 | 0.5 | 1.15 | 0.000483 | 5.00E-05 |
| 159 | Kif13b | chr14:64652530-64806296 | OK | 1.08 | 2.23 | 1.15 | 0.000483 | 5.00E-05 |
| 160 | Ssfa2 | chr2:79635424-79672964 | OK | 2.2 | 3.35 | 1.15 | 0.000483 | 5.00E-05 |
| 161 | Arrb1 | chr7:99535485-99606771 | OK | 0.29 | 1.43 | 1.14 | 0.000483 | 5.00E-05 |
| 162 | Gstt1 | chr10:75783812-75798584 | OK | 1.37 | 2.51 | 1.14 | 0.000483 | 5.00E-05 |
| 163 | Kalrn | chr16:33969072-34514027 | OK | 1.11 | 2.25 | 1.14 | 0.000483 | 5.00E-05 |
| 164 | Mmp28 | chr11:83441875-83462961 | OK | -0.24 | 0.89 | 1.14 | 0.000483 | 5.00E-05 |
| 165 | Anpep | chr7:79821802-79842352 | OK | 0.49 | 1.62 | 1.13 | 0.000483 | 5.00E-05 |
| 166 | Atp10d | chr5:72203328-72298771 | OK | 0.77 | 1.89 | 1.11 | 0.000483 | 5.00E-05 |
| 167 | Phactr1 | chr13:42680798-43138512 | OK | 2.18 | 3.29 | 1.11 | 0.000483 | 5.00E-05 |
| 168 | Angpt1 | chr15:42424666-42676977 | OK | 0.53 | 1.63 | 1.09 | 0.000483 | 5.00E-05 |
| 169 | Col28a1 | chr6:7997807-8192617 | OK | 0.35 | 1.43 | 1.09 | 0.000483 | 5.00E-05 |
| 170 | Fgfbp3 | chr19:36917549-36919599 | OK | 2.14 | 3.23 | 1.09 | 0.000483 | 5.00E-05 |
| 171 | Pde3a | chr6:141249268-141499351 | OK | 0.73 | 1.82 | 1.09 | 0.000483 | 5.00E-05 |
| 172 | Aox1 | chr1:58029968-58106410 | OK | 0.48 | 1.57 | 1.08 | 0.000483 | 5.00E-05 |
| 173 | Bpgm | chr6:34476355-34505610 | OK | 3.01 | 4.09 | 1.08 | 0.000483 | 5.00E-05 |
| 174 | C1rl | chr6:124493112-124510643 | OK | -0.22 | 0.85 | 1.07 | 0.000483 | 5.00E-05 |

| | | | | | | | | |
|---|---|---|---|---|---|---|---|---|
| 175 | Ddr1 | chr17:35681566-35704139 | OK | 3.86 | 4.93 | 1.07 | 0.000483 | 5.00E-05 |
| 176 | Far1 | chr7:113513833-113570888 | OK | 5.51 | 6.58 | 1.07 | 0.000483 | 5.00E-05 |
| 177 | Ppp1r3c | chr19:36731730-36736604 | OK | 0.72 | 1.79 | 1.07 | 0.000483 | 5.00E-05 |
| 178 | Ptx3 | chr3:66053557-66296837 | OK | 7.25 | 8.32 | 1.07 | 0.000483 | 5.00E-05 |
| 179 | Dab2 | chr15:6299788-6440709 | OK | 3.84 | 4.91 | 1.06 | 0.000483 | 5.00E-05 |
| 180 | Paqr8 | chr1:20890621-20938756 | OK | -0.46 | 0.61 | 1.06 | 0.000483 | 5.00E-05 |
| 181 | 9930012K11F | chr14:70154404-70159502 | OK | 1.6 | 2.65 | 1.05 | 0.000483 | 5.00E-05 |
| 182 | Thbs1 | chr2:118111921-118127133 | OK | 6.48 | 7.53 | 1.05 | 0.000483 | 5.00E-05 |
| 183 | Rapgef3 | chr15:97744769-97767666 | OK | 1.05 | 2.09 | 1.04 | 0.000483 | 5.00E-05 |
| 184 | Stx11 | chr10:12939982-12964259 | OK | 1.56 | 2.6 | 1.04 | 0.000483 | 5.00E-05 |
| 185 | Dhrs3 | chr4:144892826-144927645 | OK | -0.34 | 0.69 | 1.03 | 0.000483 | 5.00E-05 |
| 186 | Dock4 | chr12:40446052-40846488 | OK | -0.48 | 0.54 | 1.03 | 0.000483 | 5.00E-05 |
| 187 | Glcci1 | chr6:8509599-8597549 | OK | 1.72 | 2.75 | 1.03 | 0.000483 | 5.00E-05 |
| 188 | Nupr1 | chr7:126623245-126625470 | OK | 4 | 5.02 | 1.02 | 0.000483 | 5.00E-05 |
| 189 | Per2 | chr1:91415981-91459328 | OK | 0.7 | 1.72 | 1.02 | 0.000483 | 5.00E-05 |
| 190 | Cemip | chr7:83932856-84086505 | OK | 4.83 | 5.84 | 1.01 | 0.000483 | 5.00E-05 |
| 191 | Mocos | chr18:24653690-24701556 | OK | 0.39 | 1.4 | 1.01 | 0.000483 | 5.00E-05 |
| 192 | Ngf | chr3:102469918-102521013 | OK | 2.88 | 3.89 | 1.01 | 0.000483 | 5.00E-05 |
| 193 | Cthrc1 | chr15:39076931-39087119 | OK | 6.44 | 7.44 | 1 | 0.000483 | 5.00E-05 |
| 194 | Ddhd1 | chr14:45593170-45658143 | OK | 2.3 | 3.3 | 1 | 0.000483 | 5.00E-05 |
| 195 | Gpr85 | chr6:13835073-13839848 | OK | 2.55 | 3.55 | 1 | 0.000483 | 5.00E-05 |
| 196 | Ugt1a7c | chr1:88055410-88220002 | OK | 3.32 | 2.31 | -1 | 0.000483 | 5.00E-05 |
| 197 | Unc13b | chr4:43058983-43264887 | OK | 0.04 | -0.96 | -1 | 0.000483 | 5.00E-05 |
| 198 | Inpp4b | chr8:81342561-82127917 | OK | 0.13 | -0.88 | -1.01 | 0.000483 | 5.00E-05 |
| 199 | Klk8 | chr7:43797576-43803822 | OK | 0.56 | -0.46 | -1.02 | 0.000483 | 5.00E-05 |
| 200 | Casp12 | chr9:5345475-5373034 | OK | 3.96 | 2.93 | -1.03 | 0.000483 | 5.00E-05 |
| 201 | Slc27a6 | chr18:58556239-58612869 | OK | 2.79 | 1.76 | -1.03 | 0.000483 | 5.00E-05 |
| 202 | Ablim1 | chr19:57033263-57216032 | OK | 0.94 | -0.1 | -1.04 | 0.000483 | 5.00E-05 |
| 203 | Slc16a9 | chr10:70245275-70285951 | OK | 0.26 | -0.78 | -1.04 | 0.000483 | 5.00E-05 |
| 204 | Car13 | chr3:14641726-14663002 | OK | 4.42 | 3.37 | -1.05 | 0.000483 | 5.00E-05 |

| | | | | | | | | |
|---|---|---|---|---|---|---|---|---|
| 205 | Cx3cl1 | chr8:94772179-94782426 | OK | 4.7 | 3.65 | -1.05 | 0.000483 | 5.00E-05 |
| 206 | Mcam | chr9:44134657-44142726 | OK | 1.23 | 0.18 | -1.05 | 0.000483 | 5.00E-05 |
| 207 | Notch1 | chr2:26457901-26503822 | OK | 1.35 | 0.3 | -1.05 | 0.000483 | 5.00E-05 |
| 208 | Casp4 | chr9:5308848-5336791 | OK | 2.91 | 1.85 | -1.06 | 0.000483 | 5.00E-05 |
| 209 | Il1rl1 | chr1:40429569-40465414 | OK | 9.88 | 8.82 | -1.06 | 0.000483 | 5.00E-05 |
| 210 | P2rx3 | chr2:84996551-85035834 | OK | 2.18 | 1.13 | -1.06 | 0.000483 | 5.00E-05 |
| 211 | Foxs1 | chr2:152931897-152933208 | OK | 1.62 | 0.55 | -1.07 | 0.000483 | 5.00E-05 |
| 212 | Myo1d | chr11:80482126-80780025 | OK | 4.76 | 3.69 | -1.07 | 0.000483 | 5.00E-05 |
| 213 | Il11 | chr7:4772377-4782857 | OK | 3.58 | 2.5 | -1.08 | 0.000483 | 5.00E-05 |
| 214 | Cacna1i | chr15:80287237-80398292 | OK | 0.3 | -0.8 | -1.1 | 0.000483 | 5.00E-05 |
| 215 | Ptn | chr6:36715662-36811361 | OK | 7.85 | 6.75 | -1.1 | 0.000483 | 5.00E-05 |
| 216 | Gm2115 | chr7:84528953-84578339 | OK | 5.97 | 4.86 | -1.11 | 0.000483 | 5.00E-05 |
| 217 | Siglec15 | chr18:78043613-78057268 | OK | 0.65 | -0.47 | -1.11 | 0.00463 | 6.00E-04 |
| 218 | Cish | chr9:107296688-107301961 | OK | 3.61 | 2.49 | -1.12 | 0.000483 | 5.00E-05 |
| 219 | Dnmt3l | chr10:78030021-78063622 | OK | 2.1 | 0.98 | -1.12 | 0.000483 | 5.00E-05 |
| 220 | Sox5 | chr6:143828424-144209568 | OK | 2.42 | 1.29 | -1.13 | 0.000483 | 5.00E-05 |
| 221 | Foxo6 | chr4:120267077-120287261 | OK | 0.26 | -0.9 | -1.16 | 0.000483 | 5.00E-05 |
| 222 | Abcg2 | chr6:58596671-58692451 | OK | 4.1 | 2.92 | -1.18 | 0.000483 | 5.00E-05 |
| 223 | Foxl1 | chr8:121127684-121130644 | OK | 1.04 | -0.15 | -1.19 | 0.000483 | 5.00E-05 |
| 224 | Trem2 | chr17:48346400-48352276 | OK | 2.47 | 1.28 | -1.19 | 0.000483 | 5.00E-05 |
| 225 | Mmp9 | chr2:164948218-164955849 | OK | 1.14 | -0.05 | -1.2 | 0.000483 | 5.00E-05 |
| 226 | Ank3 | chr10:69533707-70027436 | OK | 4.46 | 3.25 | -1.21 | 0.000483 | 5.00E-05 |
| 227 | Rcan2 | chr17:43801850-44039516 | OK | 3.92 | 2.69 | -1.22 | 0.000483 | 5.00E-05 |
| 228 | Hspb7 | chr4:141420778-141425310 | OK | 0.19 | -1.04 | -1.23 | 0.000483 | 5.00E-05 |
| 229 | Kcnab1 | chr3:65109367-65378225 | OK | 1.94 | 0.7 | -1.24 | 0.000483 | 5.00E-05 |
| 230 | Diras2 | chr13:52504374-52530836 | OK | 0.96 | -0.29 | -1.25 | 0.000483 | 5.00E-05 |
| 231 | Txnip | chr3:96555767-96566801 | OK | 4.95 | 3.71 | -1.25 | 0.000483 | 5.00E-05 |
| 232 | Eva1a | chr6:82041627-82093099 | OK | 0.01 | -1.26 | -1.27 | 0.00134 | 0.00015 |
| 233 | Mycn | chr12:12936092-12941836 | OK | 1.73 | 0.45 | -1.28 | 0.000483 | 5.00E-05 |
| 234 | Hivep3 | chr4:119814677-120135411 | OK | 1.58 | 0.28 | -1.31 | 0.000483 | 5.00E-05 |

| | | | | | | | | |
|---|---|---|---|---|---|---|---|---|
| 235 | Snap25 | chr2:136713449-136782428 | OK | 1.93 | 0.63 | -1.31 | 0.000483 | 5.00E-05 |
| 236 | Acot11 | chr4:106733914-106799831 | OK | 0.53 | -0.79 | -1.32 | 0.000483 | 5.00E-05 |
| 237 | Cd24a | chr10:43579168-43584265 | OK | 6.68 | 5.36 | -1.32 | 0.000483 | 5.00E-05 |
| 238 | Fxyd1 | chr7:31051677-31055656 | OK | 2.91 | 1.57 | -1.34 | 0.000483 | 5.00E-05 |
| 239 | Smarca1 | chrX:47809369-47892552 | OK | 1.33 | -0.01 | -1.34 | 0.000483 | 5.00E-05 |
| 240 | 9130024F11R | chr1:56971468-56975196 | OK | 1.38 | 0.01 | -1.38 | 0.000483 | 5.00E-05 |
| 241 | Dact1 | chr12:71309883-71320107 | OK | 2.71 | 1.32 | -1.39 | 0.000483 | 5.00E-05 |
| 242 | Sphk1 | chr11:116531910-116536675 | OK | 2.05 | 0.65 | -1.39 | 0.000483 | 5.00E-05 |
| 243 | Lpl | chr8:68880554-68906932 | OK | 2.64 | 1.23 | -1.41 | 0.000483 | 5.00E-05 |
| 244 | Lvrn | chr18:46850038-46905446 | OK | 0.4 | -1.01 | -1.42 | 0.000483 | 5.00E-05 |
| 245 | Cd34 | chr1:194938820-194976959 | OK | 6.19 | 4.76 | -1.43 | 0.000483 | 5.00E-05 |
| 246 | Eya1 | chr1:14168957-14310199 | OK | 1.37 | -0.07 | -1.44 | 0.000483 | 5.00E-05 |
| 247 | Scg5 | chr2:113776312-113829091 | OK | 1.55 | 0.11 | -1.44 | 0.000483 | 5.00E-05 |
| 248 | Fibin | chr2:110360924-110362993 | OK | 0.56 | -0.93 | -1.49 | 0.000483 | 5.00E-05 |
| 249 | Grpr | chrX:163513903-163549736 | OK | 1.23 | -0.28 | -1.51 | 0.000483 | 5.00E-05 |
| 250 | Spp1 | chr5:104435110-104441053 | OK | 10.87 | 9.32 | -1.54 | 0.000483 | 5.00E-05 |
| 251 | Sytl2 | chr7:90302354-90410719 | OK | 1.18 | -0.37 | -1.55 | 0.000483 | 5.00E-05 |
| 252 | AW551984 | chr9:39587395-39604124 | OK | 4.84 | 3.27 | -1.57 | 0.000483 | 5.00E-05 |
| 253 | Fam135b | chr15:71445677-71727838 | OK | 1.65 | 0.06 | -1.59 | 0.000483 | 5.00E-05 |
| 254 | Cytip | chr2:58129138-58160122 | OK | 0.41 | -1.2 | -1.61 | 0.000483 | 5.00E-05 |
| 255 | Catip | chr1:74362107-74369321 | OK | 1.4 | -0.25 | -1.65 | 0.000483 | 5.00E-05 |
| 256 | Dkk2 | chr3:132085291-132180304 | OK | 2.06 | 0.4 | -1.65 | 0.000483 | 5.00E-05 |
| 257 | Sulf2 | chr2:166073898-166155683 | OK | 2.17 | 0.53 | -1.65 | 0.000483 | 5.00E-05 |
| 258 | St3gal1 | chr15:67102874-67176882 | OK | 4.79 | 3.12 | -1.67 | 0.000483 | 5.00E-05 |
| 259 | Kcnk10 | chr12:98433993-98577940 | OK | 2.94 | 1.25 | -1.68 | 0.000483 | 5.00E-05 |
| 260 | Cxcl1 | chr5:90891244-90893115 | OK | 1.29 | -0.43 | -1.71 | 0.000483 | 5.00E-05 |
| 261 | Tusc1 | chr4:93334147-93335511 | OK | 1.82 | 0.11 | -1.72 | 0.000483 | 5.00E-05 |
| 262 | H19 | chr7:142575529-142578146 | OK | 4.1 | 2.36 | -1.74 | 0.000483 | 5.00E-05 |
| 263 | Rgcc | chr14:79288749-79301635 | OK | 2.89 | 1.15 | -1.74 | 0.000483 | 5.00E-05 |
| 264 | Tnc | chr4:63959784-64047015 | OK | 7.55 | 5.79 | -1.76 | 0.000483 | 5.00E-05 |

| | | | | | | | | |
|---|---|---|---|---|---|---|---|---|
| 265 | Mapkapk3 | chr9:107254926-107289877 | OK | 4.23 | 2.45 | -1.77 | 0.000483 | 5.00E-05 |
| 266 | Ptgfr | chr3:151798609-151837528 | OK | 5.42 | 3.64 | -1.78 | 0.000483 | 5.00E-05 |
| 267 | Il18rap | chr1:40515361-40551705 | OK | 1.18 | -0.61 | -1.79 | 0.000483 | 5.00E-05 |
| 268 | Has2 | chr15:56665626-56694546 | OK | 4.49 | 2.69 | -1.8 | 0.000483 | 5.00E-05 |
| 269 | 4833427G06I | chr9:51081312-51102078 | OK | 1.46 | -0.36 | -1.82 | 0.000483 | 5.00E-05 |
| 270 | Slc17a7 | chr7:45163920-45176139 | OK | 0.06 | -1.8 | -1.85 | 0.000483 | 5.00E-05 |
| 271 | Dcn | chr10:97479499-97518162 | OK | 6.89 | 5.02 | -1.87 | 0.000483 | 5.00E-05 |
| 272 | Ramp3 | chr11:6650147-6677475 | OK | 5.93 | 4.04 | -1.89 | 0.000483 | 5.00E-05 |
| 273 | Crabp1 | chr9:54764747-54773110 | OK | 7.83 | 5.91 | -1.92 | 0.000483 | 5.00E-05 |
| 274 | Serpine2 | chr1:79794320-79858665 | OK | 5.22 | 3.25 | -1.97 | 0.000483 | 5.00E-05 |
| 275 | Ptgir | chr7:16906489-16910905 | OK | 0.12 | -1.92 | -2.05 | 0.000483 | 5.00E-05 |
| 276 | Lum | chr10:97565500-97572703 | OK | 5.12 | 3.07 | -2.06 | 0.000483 | 5.00E-05 |
| 277 | Pkp2 | chr16:16213344-16272712 | OK | 0.15 | -1.93 | -2.07 | 0.000483 | 5.00E-05 |
| 278 | Tnn | chr1:160085031-160153575 | OK | 1.33 | -0.83 | -2.16 | 0.000483 | 5.00E-05 |
| 279 | Cxcl5 | chr5:90759297-90761625 | OK | 3.31 | 1.06 | -2.25 | 0.000483 | 5.00E-05 |
| 280 | Sfrp2 | chr3:83766320-83774314 | OK | 5.11 | 2.83 | -2.27 | 0.000483 | 5.00E-05 |
| 281 | Fcrlb | chr1:170907272-170912941 | OK | 0.25 | -2.05 | -2.3 | 0.00212 | 0.00025 |
| 282 | Osbp2 | chr11:3703730-3863903 | OK | 3.5 | 1.12 | -2.38 | 0.000483 | 5.00E-05 |
| 283 | Rgag4 | chrX:101849384-102092055 | OK | 0.14 | -2.28 | -2.42 | 0.000483 | 5.00E-05 |
| 284 | 4930550L24R | chrX:58911460-58920304 | OK | 2.46 | 0 | -2.46 | 0.000483 | 5.00E-05 |
| 285 | Gpr149 | chr3:62529962-62605140 | OK | 1.65 | -0.82 | -2.47 | 0.000483 | 5.00E-05 |
| 286 | Rtl1 | chr12:109590168-109595403 | OK | 0.46 | -2.08 | -2.53 | 0.000483 | 5.00E-05 |
| 287 | Igf1 | chr10:87859055-87937047 | OK | 1.62 | -1.18 | -2.8 | 0.000483 | 5.00E-05 |
| 288 | Agtr2 | chrX:21484623-21488833 | OK | 0.02 | -2.96 | -2.98 | 0.000483 | 5.00E-05 |
| 289 | Lmcd1 | chr6:112273757-112330423 | OK | 0.63 | -2.46 | -3.09 | 0.000483 | 5.00E-05 |
| 290 | Sparcl1 | chr5:104079108-104114088 | OK | 3.86 | 0.45 | -3.4 | 0.000483 | 5.00E-05 |
| 291 | Cck | chr9:121489823-121495694 | OK | 2.81 | -0.95 | -3.76 | 0.000483 | 5.00E-05 |
| 292 | Vip | chr10:5639217-5647614 | OK | 3.95 | -0.26 | -4.22 | 0.000483 | 5.00E-05 |

**Table S1. List of differentially expressed genes in NIH3T3 cells expressing Foxc1 or pLXSH empty vector control.**

### DESeq2 results:

Selection criteria: q-value (FDR-adjusted p-value) ≤ 0.01, absolute log2 ratio ≥ 1, CuffLinks signal cut-off ≥ 1 FPKM in either condition.

| # | Gene | Locus | baseMean | log2 ratio | log2 ratio s.e. | q value | p value |
|---|---|---|---|---|---|---|---|
| 1 | Il1rn | chr2:24336859-24351491 | 1660.92 | 12.34 | 1.02 | 1.43E-31 | 1.65E-33 |
| 2 | Arhgap36 | chrX:49470449-49500250 | 211.18 | 11.22 | 1.18 | 1.31E-19 | 2.71E-21 |
| 3 | Anxa10 | chr8:62057041-62123193 | 124.69 | 10.45 | 1.19 | 5.35E-17 | 1.27E-18 |
| 4 | Kng1 | chr16:23058299-23082078 | 55.60 | 9.29 | 1.20 | 2.81E-13 | 8.66E-15 |
| 5 | F5 | chr1:164115263-164220277 | 580.18 | 9.01 | 0.55 | 8.93E-58 | 4.83E-60 |
| 6 | Glra3 | chr8:55940824-56125352 | 17.17 | 7.60 | 1.23 | 1.47E-08 | 7.28E-10 |
| 7 | E330011O21I | chr16:78250862-78255454 | 16.42 | 7.53 | 1.24 | 2.17E-08 | 1.10E-09 |
| 8 | Serpina3h | chr12:104247895-104254405 | 58.63 | 6.50 | 0.75 | 1.63E-16 | 3.97E-18 |
| 9 | Vnn1 | chr10:23894687-23905343 | 436.13 | 6.39 | 0.27 | 2.52E-121 | 4.04E-124 |
| 10 | Fgf23 | chr6:127072901-127082296 | 92.47 | 6.34 | 0.57 | 4.77E-27 | 7.09E-29 |
| 11 | Akr1c18 | chr13:4132626-4150631 | 277.00 | 5.92 | 0.29 | 2.33E-88 | 8.36E-91 |
| 12 | Wnt2 | chr6:17988939-18030445 | 123.95 | 5.86 | 0.43 | 6.45E-41 | 5.56E-43 |
| 13 | Mc5r | chr18:68337602-68339711 | 14.34 | 5.45 | 1.10 | 1.06E-05 | 7.90E-07 |
| 14 | Il33 | chr19:29925113-29960715 | 52.88 | 5.15 | 0.52 | 5.15E-21 | 1.00E-22 |
| 15 | Agt | chr8:124556586-124569707 | 130.23 | 4.91 | 0.31 | 1.10E-52 | 6.81E-55 |
| 16 | Prss35 | chr9:86743632-86757506 | 49.85 | 4.91 | 0.51 | 1.85E-20 | 3.76E-22 |
| 17 | Itih2 | chr2:10094590-10130683 | 207.88 | 4.89 | 0.25 | 1.33E-82 | 5.18E-85 |
| 18 | Foxc1 | chr13:31806645-31810635 | 5178.65 | 4.86 | 0.08 | 0.00E+00 | 0.00E+00 |
| 19 | Otor | chr2:143078491-143081699 | 14.28 | 4.84 | 0.92 | 2.28E-06 | 1.54E-07 |
| 20 | Col10a1 | chr10:34303611-34418528 | 990.48 | 4.56 | 0.11 | 0.00E+00 | 1.00E-200 |
| 21 | Il20ra | chr10:19712586-19760053 | 126.46 | 4.12 | 0.26 | 1.98E-53 | 1.18E-55 |
| 22 | Ogn | chr13:49464058-49652731 | 2051.72 | 3.72 | 0.09 | 0.00E+00 | 0.00E+00 |
| 23 | Apba1 | chr19:23758875-23949597 | 983.90 | 3.60 | 0.10 | 0.00E+00 | 0.00E+00 |
| 24 | Slc4a4 | chr5:88887259-89239656 | 98.01 | 3.58 | 0.26 | 2.01E-40 | 1.77E-42 |
| 25 | Itgbl1 | chr14:123660139-123974079 | 397.21 | 3.50 | 0.14 | 2.37E-134 | 2.53E-137 |
| 26 | Ifit3b | chr19:34607956-34613401 | 22.65 | 3.42 | 0.53 | 2.29E-09 | 1.04E-10 |

| | | | | | | | |
|---|---|---|---|---|---|---|---|
| 27 | Itga1 | chr13:114958080-115101964 | 305.63 | 3.38 | 0.15 | 5.18E-109 | 1.11E-111 |
| 28 | Camk1g | chr1:193346345-193370282 | 58.66 | 3.33 | 0.33 | 9.12E-23 | 1.61E-24 |
| 29 | Ifi47 | chr11:49037659-49135387 | 26.42 | 3.31 | 0.47 | 7.74E-11 | 3.00E-12 |
| 30 | Hid1 | chr11:115347708-115367719 | 294.66 | 3.26 | 0.16 | 1.40E-95 | 4.48E-98 |
| 31 | Krt15 | chr11:100131758-100135949 | 62.01 | 3.20 | 0.30 | 5.07E-24 | 8.42E-26 |
| 32 | Pdk4 | chr6:5483350-5496278 | 2228.92 | 3.17 | 0.08 | 0.00E+00 | 1.00E-200 |
| 33 | Rspo2 | chr15:43020794-43170818 | 124.16 | 3.15 | 0.22 | 4.43E-46 | 3.17E-48 |
| 34 | Fam20a | chr11:109672925-109722256 | 266.02 | 3.08 | 0.15 | 4.27E-89 | 1.50E-91 |
| 35 | Nlrp2 | chr7:5298546-5360682 | 55.11 | 3.01 | 0.31 | 4.74E-20 | 9.68E-22 |
| 36 | C430002N11I | chr9:96765561-96774397 | 21.61 | 2.89 | 0.50 | 1.30E-07 | 7.21E-09 |
| 37 | Myh14 | chr7:44605802-44670843 | 99.00 | 2.87 | 0.23 | 2.37E-34 | 2.56E-36 |
| 38 | Omd | chr13:49464058-49652731 | 147.35 | 2.86 | 0.19 | 4.89E-48 | 3.35E-50 |
| 39 | Sulf1 | chr1:12692429-12860372 | 525.07 | 2.85 | 0.11 | 1.04E-136 | 1.03E-139 |
| 40 | Rasgef1b | chr5:99217419-99252927 | 72.43 | 2.82 | 0.27 | 1.21E-24 | 1.97E-26 |
| 41 | Pgbd5 | chr8:124369048-124433936 | 209.31 | 2.78 | 0.16 | 2.16E-65 | 1.01E-67 |
| 42 | Scara5 | chr14:65666402-65764826 | 85.31 | 2.76 | 0.25 | 9.57E-27 | 1.45E-28 |
| 43 | Dio2 | chr12:90724551-90738438 | 1017.36 | 2.72 | 0.09 | 9.25E-189 | 4.23E-192 |
| 44 | Ifit3 | chr19:34583528-34588982 | 35.43 | 2.72 | 0.37 | 7.40E-12 | 2.61E-13 |
| 45 | Wfdc12 | chr2:164189230-164190558 | 11.03 | 2.57 | 0.64 | 0.0005914 | 6.36E-05 |
| 46 | Srpx2 | chrX:133908424-133932446 | 149.81 | 2.50 | 0.18 | 1.16E-42 | 9.44E-45 |
| 47 | Pkia | chr3:7366603-7445365 | 434.28 | 2.50 | 0.13 | 9.69E-80 | 3.91E-82 |
| 48 | Plscr2 | chr9:92275601-92297752 | 71.95 | 2.46 | 0.25 | 8.60E-21 | 1.70E-22 |
| 49 | Adra1b | chr11:43774604-43901237 | 81.67 | 2.38 | 0.24 | 6.13E-22 | 1.15E-23 |
| 50 | Napsa | chr7:44572444-44586846 | 23.63 | 2.36 | 0.43 | 4.66E-07 | 2.84E-08 |
| 51 | Aldh3a1 | chr11:61208741-61218416 | 227.18 | 2.28 | 0.15 | 3.83E-49 | 2.57E-51 |
| 52 | Gdf5 | chr2:155941024-155945364 | 161.73 | 2.27 | 0.17 | 3.45E-40 | 3.05E-42 |
| 53 | Col11a1 | chr3:114030539-114220326 | 879.79 | 2.24 | 0.09 | 9.04E-133 | 1.03E-135 |
| 54 | 8430408G22I | chr6:116633007-116673836 | 19.11 | 2.14 | 0.47 | 5.32E-05 | 4.53E-06 |
| 55 | C1qtnf1 | chr11:118428456-118454995 | 223.49 | 2.11 | 0.15 | 4.34E-43 | 3.44E-45 |
| 56 | Ror1 | chr4:100095790-100442545 | 291.43 | 2.11 | 0.13 | 9.90E-56 | 5.51E-58 |
| 57 | Postn | chr3:54361106-54391041 | 4981.18 | 2.08 | 0.15 | 2.78E-41 | 2.35E-43 |
| 58 | Nr3c2 | chr8:76902507-77243639 | 54.55 | 2.08 | 0.29 | 1.74E-11 | 6.36E-13 |

| | | | | | | | |
|---|---|---|---|---|---|---|---|
| 59 | Enpp2 | chr15:54838678-54920146 | 87.21 | 2.06 | 0.23 | 1.45E-17 | 3.35E-19 |
| 60 | Sorcs2 | chr5:36017180-36398139 | 250.32 | 2.03 | 0.13 | 2.13E-49 | 1.40E-51 |
| 61 | Tpd52l1 | chr10:31332379-31445921 | 21.83 | 1.95 | 0.42 | 4.23E-05 | 3.50E-06 |
| 62 | Adamts6 | chr13:104287872-104494763 | 123.01 | 1.94 | 0.19 | 1.21E-23 | 2.08E-25 |
| 63 | Maf | chr8:115703252-115706894 | 154.48 | 1.88 | 0.17 | 5.68E-27 | 8.48E-29 |
| 64 | Fhl1 | chrX:56731760-56793346 | 31.58 | 1.88 | 0.35 | 8.50E-07 | 5.41E-08 |
| 65 | Aspg | chr12:112106682-112127573 | 140.16 | 1.84 | 0.17 | 2.83E-25 | 4.49E-27 |
| 66 | Mtss1 | chr15:58941233-59082026 | 63.40 | 1.83 | 0.25 | 1.72E-11 | 6.28E-13 |
| 67 | Wisp2 | chr2:163820833-163833147 | 2112.67 | 1.80 | 0.06 | 2.19E-172 | 1.17E-175 |
| 68 | Kcnh5 | chr12:74897216-75177332 | 73.76 | 1.77 | 0.23 | 4.04E-13 | 1.27E-14 |
| 69 | Fam213a | chr14:40993739-41013775 | 414.31 | 1.75 | 0.10 | 1.56E-60 | 7.95E-63 |
| 70 | Msln | chr17:25748612-25754327 | 475.55 | 1.74 | 0.10 | 1.77E-63 | 8.48E-66 |
| 71 | Vamp5 | chr6:72368048-72380468 | 171.91 | 1.73 | 0.16 | 4.22E-27 | 6.23E-29 |
| 72 | Tgfb2 | chr1:186623185-186705992 | 353.65 | 1.71 | 0.11 | 3.13E-50 | 1.95E-52 |
| 73 | Fjx1 | chr2:102449365-102451792 | 65.19 | 1.70 | 0.25 | 3.26E-10 | 1.34E-11 |
| 74 | Aspn | chr13:49464058-49652731 | 1437.46 | 1.70 | 0.17 | 2.13E-21 | 4.05E-23 |
| 75 | Sfrp1 | chr8:23411501-23449632 | 1042.09 | 1.69 | 0.09 | 1.53E-82 | 6.05E-85 |
| 76 | Ahr | chr12:35497978-35534989 | 1606.88 | 1.68 | 0.07 | 6.54E-114 | 1.35E-116 |
| 77 | Rnf43 | chr11:87663086-87735539 | 209.15 | 1.68 | 0.14 | 2.23E-29 | 2.85E-31 |
| 78 | Lhx8 | chr3:154306293-154330560 | 105.26 | 1.65 | 0.19 | 4.48E-16 | 1.12E-17 |
| 79 | Loxl4 | chr19:42592278-42612806 | 1028.28 | 1.63 | 0.08 | 5.94E-102 | 1.50E-104 |
| 80 | Mest | chr6:30733505-30896794 | 93.08 | 1.60 | 0.21 | 1.98E-12 | 6.66E-14 |
| 81 | Fst | chr13:114452261-114458951 | 108.24 | 1.59 | 0.19 | 5.16E-16 | 1.30E-17 |
| 82 | Brinp3 | chr1:146495665-146902472 | 180.97 | 1.59 | 0.16 | 2.76E-22 | 5.06E-24 |
| 83 | S100a16 | chr3:90541222-90543151 | 21.99 | 1.58 | 0.42 | 0.0013337 | 0.000157 |
| 84 | Adgrg6 | chr10:14402584-14545036 | 151.41 | 1.58 | 0.16 | 1.59E-20 | 3.16E-22 |
| 85 | 2010111I01R | chr13:62964892-63431745 | 588.56 | 1.58 | 0.09 | 2.32E-60 | 1.20E-62 |
| 86 | Actr3b | chr5:25760025-25850341 | 107.65 | 1.58 | 0.19 | 6.00E-15 | 1.63E-16 |
| 87 | Gadd45a | chr6:67035095-67080652 | 363.23 | 1.58 | 0.11 | 7.67E-44 | 5.96E-46 |
| 88 | Alcam | chr16:52248995-52452997 | 1623.55 | 1.58 | 0.07 | 9.17E-118 | 1.68E-120 |
| 89 | Klf4 | chr4:55527136-55532475 | 1263.39 | 1.57 | 0.07 | 1.96E-104 | 4.49E-107 |
| 90 | Ly6a | chr15:74994876-74998031 | 344.82 | 1.57 | 0.12 | 2.57E-38 | 2.47E-40 |

| | | | | | | | |
|---|---|---|---|---|---|---|---|
| 91 | Cped1 | chr6:21985909-22255606 | 1299.61 | 1.57 | 0.09 | 3.19E-64 | 1.51E-66 |
| 92 | Mfap3l | chr8:60632824-60676731 | 363.27 | 1.55 | 0.11 | 5.50E-41 | 4.69E-43 |
| 93 | Etl4 | chr2:20289912-20810535 | 310.73 | 1.55 | 0.12 | 2.40E-37 | 2.39E-39 |
| 94 | Rerg | chr6:137054824-137170496 | 90.63 | 1.55 | 0.20 | 5.63E-13 | 1.80E-14 |
| 95 | Ctgf | chr10:24595441-24598682 | 8793.21 | 1.53 | 0.06 | 5.47E-160 | 4.17E-163 |
| 96 | Cyp1b1 | chr17:79706952-79715041 | 1089.53 | 1.53 | 0.08 | 1.45E-72 | 6.18E-75 |
| 97 | Gjb4 | chr4:127351085-127354081 | 90.82 | 1.53 | 0.20 | 1.61E-12 | 5.32E-14 |
| 98 | Rbp1 | chr9:98422960-98446550 | 2776.66 | 1.52 | 0.07 | 6.85E-104 | 1.62E-106 |
| 99 | Efnb2 | chr8:8617438-8660773 | 2090.50 | 1.52 | 0.07 | 5.52E-106 | 1.22E-108 |
| 100 | Rbp2 | chr9:98490536-98509771 | 17.91 | 1.51 | 0.44 | 0.0049683 | 0.00069 |
| 101 | Nrep | chr18:33437018-33464029 | 432.06 | 1.50 | 0.11 | 1.01E-38 | 9.55E-41 |
| 102 | 2610035D17I | chr11:113043905-113201838 | 56.79 | 1.50 | 0.26 | 8.87E-08 | 4.81E-09 |
| 103 | Thrb | chr14:17660959-18038088 | 301.36 | 1.49 | 0.13 | 7.78E-29 | 1.04E-30 |
| 104 | Tmem176b | chr6:48833810-48841374 | 66.19 | 1.49 | 0.23 | 4.07E-09 | 1.91E-10 |
| 105 | Gm14005 | chr2:128298662-128429351 | 69.85 | 1.49 | 0.24 | 8.07E-09 | 3.91E-10 |
| 106 | Megf6 | chr4:154170712-154275721 | 203.20 | 1.49 | 0.15 | 1.46E-22 | 2.60E-24 |
| 107 | Dusp4 | chr8:34807609-34819894 | 212.10 | 1.46 | 0.14 | 1.50E-23 | 2.59E-25 |
| 108 | Cdsn | chr17:35552127-35557180 | 923.83 | 1.43 | 0.08 | 1.26E-73 | 5.27E-76 |
| 109 | Tle4 | chr19:14448071-14598183 | 482.94 | 1.42 | 0.10 | 4.09E-47 | 2.84E-49 |
| 110 | Fzd1 | chr5:4753838-4758216 | 4950.70 | 1.41 | 0.06 | 2.22E-128 | 3.04E-131 |
| 111 | Dusp1 | chr17:26505590-26508472 | 534.38 | 1.40 | 0.10 | 1.49E-45 | 1.10E-47 |
| 112 | Irgm1 | chr11:48865248-48871346 | 416.18 | 1.40 | 0.10 | 5.75E-40 | 5.17E-42 |
| 113 | Pam | chr1:97821093-98095632 | 4112.66 | 1.39 | 0.06 | 1.78E-129 | 2.31E-132 |
| 114 | Cpeb3 | chr19:37021290-37207471 | 271.27 | 1.38 | 0.14 | 1.17E-22 | 2.06E-24 |
| 115 | Agpat9 | chr5:100846228-100899102 | 59.46 | 1.36 | 0.25 | 6.53E-07 | 4.06E-08 |
| 116 | Pde8b | chr13:95024104-95250304 | 395.88 | 1.35 | 0.11 | 2.88E-30 | 3.48E-32 |
| 117 | Pld1 | chr3:27938679-28133362 | 174.98 | 1.35 | 0.16 | 7.09E-15 | 1.94E-16 |
| 118 | Gchfr | chr2:119167787-119172389 | 248.27 | 1.34 | 0.13 | 5.77E-24 | 9.64E-26 |
| 119 | Fam43a | chr16:30599722-30602797 | 1006.43 | 1.34 | 0.08 | 1.74E-67 | 7.94E-70 |
| 120 | Il16 | chr7:83643062-83735490 | 75.98 | 1.34 | 0.21 | 9.64E-09 | 4.68E-10 |
| 121 | Nrgn | chr9:37544492-37552745 | 29.80 | 1.33 | 0.35 | 0.0010762 | 0.000122 |
| 122 | D030025P21I | chr12:84783211-84879755 | 116.60 | 1.33 | 0.19 | 2.29E-11 | 8.44E-13 |

| | | | | | | | |
|---|---|---|---|---|---|---|---|
| 123 | Zdhhc2 | chr8:40423814-40484842 | 339.02 | 1.33 | 0.11 | 6.35E-30 | 7.84E-32 |
| 124 | Fam107b | chr2:3713457-3782134 | 1728.84 | 1.32 | 0.07 | 6.13E-75 | 2.52E-77 |
| 125 | Smoc2 | chr17:14279505-14404790 | 575.82 | 1.32 | 0.09 | 1.04E-45 | 7.57E-48 |
| 126 | Dpysl3 | chr18:43320978-43438286 | 2769.92 | 1.32 | 0.06 | 5.06E-94 | 1.70E-96 |
| 127 | Fam110c | chr12:31073967-31079940 | 240.77 | 1.31 | 0.14 | 7.69E-19 | 1.65E-20 |
| 128 | Sh3bp4 | chr1:89070461-89153793 | 776.61 | 1.31 | 0.09 | 7.77E-43 | 6.22E-45 |
| 129 | Pdlim2 | chr14:70164217-70177672 | 543.43 | 1.31 | 0.10 | 1.13E-40 | 9.84E-43 |
| 130 | Sdpr | chr1:51218385-51333779 | 4665.96 | 1.30 | 0.06 | 1.72E-96 | 5.39E-99 |
| 131 | Tec | chr5:72755717-72868448 | 49.55 | 1.29 | 0.27 | 2.45E-05 | 1.94E-06 |
| 132 | Sema5a | chr15:32244812-32696341 | 1039.86 | 1.29 | 0.09 | 2.48E-45 | 1.87E-47 |
| 133 | Ltbp2 | chr12:84783211-84879755 | 4423.59 | 1.29 | 0.08 | 6.38E-50 | 4.04E-52 |
| 134 | Fgd3 | chr13:49263109-49309208 | 143.88 | 1.28 | 0.16 | 1.83E-13 | 5.59E-15 |
| 135 | Lrrc8d | chr5:105699968-105815215 | 237.32 | 1.27 | 0.13 | 3.97E-21 | 7.69E-23 |
| 136 | Rgs20 | chr1:4909575-5070285 | 346.20 | 1.27 | 0.11 | 1.47E-28 | 2.00E-30 |
| 137 | Gpc1 | chr1:92831685-92860196 | 5975.98 | 1.27 | 0.06 | 0.00E+00 | 2.94E-103 |
| 138 | Grk5 | chr19:60889748-61140840 | 283.41 | 1.27 | 0.12 | 1.94E-23 | 3.39E-25 |
| 139 | Cpe | chr8:64592550-64693040 | 8929.50 | 1.25 | 0.05 | 7.27E-121 | 1.22E-123 |
| 140 | Phldb2 | chr16:45746230-45844378 | 3220.34 | 1.24 | 0.06 | 1.41E-95 | 4.64E-98 |
| 141 | Arhgap31 | chr16:38598342-38713035 | 695.31 | 1.24 | 0.09 | 3.59E-44 | 2.77E-46 |
| 142 | Gjb3 | chr4:127325234-127330836 | 722.77 | 1.24 | 0.10 | 5.62E-34 | 6.12E-36 |
| 143 | Amigo2 | chr15:97244073-97385691 | 931.59 | 1.24 | 0.09 | 1.68E-45 | 1.25E-47 |
| 144 | Dmxl2 | chr9:54365157-54501626 | 277.43 | 1.24 | 0.13 | 6.89E-21 | 1.35E-22 |
| 145 | Tmem176a | chr6:48841482-48845364 | 35.38 | 1.23 | 0.31 | 0.0006533 | 7.10E-05 |
| 146 | Slc7a2 | chr8:40862366-40922070 | 397.85 | 1.22 | 0.11 | 1.04E-26 | 1.58E-28 |
| 147 | Heg1 | chr16:33684465-33768195 | 1515.09 | 1.22 | 0.09 | 2.09E-39 | 1.93E-41 |
| 148 | Capn5 | chr7:98121558-98184799 | 179.81 | 1.22 | 0.15 | 3.58E-14 | 1.06E-15 |
| 149 | Tgm2 | chr2:158116404-158146392 | 362.26 | 1.22 | 0.12 | 1.81E-23 | 3.15E-25 |
| 150 | Col12a1 | chr9:79598986-79718722 | 8819.25 | 1.21 | 0.22 | 3.69E-07 | 2.20E-08 |
| 151 | Chrd | chr16:20724453-20742384 | 165.18 | 1.21 | 0.15 | 1.76E-13 | 5.34E-15 |
| 152 | Slit2 | chr5:47983154-48306282 | 8327.68 | 1.20 | 0.08 | 1.05E-53 | 6.15E-56 |
| 153 | AI606473 | chr3:154330820-154334687 | 186.22 | 1.20 | 0.15 | 7.02E-15 | 1.92E-16 |
| 154 | Gli1 | chr10:127321963-127341579 | 55.50 | 1.20 | 0.25 | 2.11E-05 | 1.65E-06 |

| | | | | | | | |
|---|---|---|---|---|---|---|---|
| 155 | Lxn | chr3:67430114-67475068 | 3111.75 | 1.20 | 0.06 | 3.52E-84 | 1.34E-86 |
| 156 | Rhou | chr8:123653928-123663880 | 1590.14 | 1.20 | 0.07 | 2.60E-70 | 1.17E-72 |
| 157 | Col5a3 | chr9:20770049-20815034 | 426.09 | 1.20 | 0.10 | 2.24E-30 | 2.70E-32 |
| 158 | Rab27b | chr18:69979130-70141605 | 379.10 | 1.20 | 0.11 | 8.35E-24 | 1.42E-25 |
| 159 | Cryab | chr9:50751071-50756635 | 900.17 | 1.20 | 0.08 | 6.19E-45 | 4.72E-47 |
| 160 | Eva1c | chr16:90830858-90904885 | 115.62 | 1.19 | 0.18 | 6.49E-10 | 2.76E-11 |
| 161 | Ly6c1 | chr15:75044017-75048837 | 111.86 | 1.19 | 0.19 | 3.60E-09 | 1.68E-10 |
| 162 | Adrb2 | chr18:62177712-62179981 | 44.82 | 1.19 | 0.28 | 0.0003171 | 3.15E-05 |
| 163 | Bicc1 | chr10:70925095-71159634 | 2805.54 | 1.18 | 0.07 | 1.52E-70 | 6.71E-73 |
| 164 | Gstt1 | chr10:75783812-75798584 | 76.36 | 1.18 | 0.23 | 2.78E-06 | 1.92E-07 |
| 165 | Mmp28 | chr11:83441875-83462961 | 64.38 | 1.16 | 0.24 | 9.98E-06 | 7.43E-07 |
| 166 | Sepp1 | chr15:3270766-3280508 | 452.61 | 1.16 | 0.11 | 5.91E-24 | 9.92E-26 |
| 167 | Irs1 | chr1:82233104-82291439 | 1841.92 | 1.16 | 0.08 | 2.35E-49 | 1.56E-51 |
| 168 | Gdpd5 | chr7:99381548-99460984 | 74.75 | 1.16 | 0.22 | 1.62E-06 | 1.07E-07 |
| 169 | Asb4 | chr6:5383385-5433021 | 678.38 | 1.15 | 0.09 | 1.59E-39 | 1.44E-41 |
| 170 | Ssfa2 | chr2:79635424-79672964 | 766.39 | 1.14 | 0.09 | 5.25E-36 | 5.48E-38 |
| 171 | Kalrn | chr16:33969072-34514027 | 694.15 | 1.14 | 0.11 | 6.60E-24 | 1.11E-25 |
| 172 | Kif13b | chr14:64652530-64806296 | 397.11 | 1.14 | 0.11 | 1.77E-23 | 3.05E-25 |
| 173 | Anpep | chr7:79821802-79842352 | 156.16 | 1.13 | 0.17 | 1.02E-09 | 4.46E-11 |
| 174 | Phactr1 | chr13:42680798-43138512 | 728.45 | 1.13 | 0.09 | 1.31E-34 | 1.40E-36 |
| 175 | Arrb1 | chr7:99535485-99606771 | 281.14 | 1.13 | 0.12 | 2.94E-18 | 6.59E-20 |
| 176 | Atp10d | chr5:72203328-72298771 | 332.87 | 1.12 | 0.11 | 1.17E-21 | 2.21E-23 |
| 177 | Fgfbp3 | chr19:36917549-36919599 | 237.35 | 1.12 | 0.13 | 4.70E-17 | 1.11E-18 |
| 178 | Trim7 | chr11:48826137-48850195 | 41.35 | 1.10 | 0.29 | 0.0015151 | 0.00018 |
| 179 | Angpt1 | chr15:42424666-42676977 | 192.93 | 1.09 | 0.14 | 2.06E-13 | 6.30E-15 |
| 180 | Col28a1 | chr6:7997807-8192617 | 172.83 | 1.08 | 0.15 | 1.19E-11 | 4.25E-13 |
| 181 | Pde3a | chr6:141249268-141499351 | 210.82 | 1.08 | 0.13 | 1.73E-14 | 4.93E-16 |
| 182 | Aox1 | chr1:58029968-58106410 | 194.65 | 1.08 | 0.15 | 3.95E-12 | 1.37E-13 |
| 183 | Gfra1 | chr19:58235580-58455398 | 125.73 | 1.08 | 0.18 | 2.04E-08 | 1.03E-09 |
| 184 | C1rl | chr6:124493112-124510643 | 78.84 | 1.07 | 0.22 | 1.57E-05 | 1.20E-06 |
| 185 | Far1 | chr7:113513833-113570888 | 6182.92 | 1.07 | 0.07 | 6.85E-53 | 4.18E-55 |
| 186 | Ptx3 | chr3:66053557-66296837 | 8650.91 | 1.07 | 0.05 | 5.62E-93 | 1.93E-95 |

| | | | | | | | |
|---|---|---|---|---|---|---|---|
| 187 | Ddr1 | chr17:35681566-35704139 | 1698.82 | 1.06 | 0.08 | 2.62E-42 | 2.18E-44 |
| 188 | Dab2 | chr15:6299788-6440709 | 2027.57 | 1.06 | 0.06 | 1.24E-61 | 6.12E-64 |
| 189 | Paqr8 | chr1:20890621-20938756 | 102.12 | 1.06 | 0.19 | 2.60E-07 | 1.51E-08 |
| 190 | Ppp1r3c | chr19:36731730-36736604 | 135.69 | 1.06 | 0.17 | 5.33E-09 | 2.53E-10 |
| 191 | Rapgef3 | chr15:97744769-97767666 | 252.50 | 1.06 | 0.14 | 3.89E-13 | 1.22E-14 |
| 192 | Stx11 | chr10:12939982-12964259 | 198.47 | 1.05 | 0.15 | 1.18E-11 | 4.21E-13 |
| 193 | Bpgm | chr6:34476355-34505610 | 517.34 | 1.05 | 0.09 | 2.21E-27 | 3.19E-29 |
| 194 | Thbs1 | chr2:118111921-118127133 | 16400.84 | 1.05 | 0.07 | 6.62E-55 | 3.79E-57 |
| 195 | Gdpd1 | chr11:86583864-87555823 | 122.25 | 1.05 | 0.18 | 1.58E-07 | 8.93E-09 |
| 196 | 9930012K11F | chr14:70154404-70159502 | 179.21 | 1.05 | 0.14 | 1.33E-11 | 4.79E-13 |
| 197 | Ngf | chr3:102469918-102521013 | 203.31 | 1.03 | 0.14 | 5.01E-12 | 1.75E-13 |
| 198 | Glcci1 | chr6:8509599-8597549 | 587.02 | 1.03 | 0.09 | 4.21E-29 | 5.55E-31 |
| 199 | Dock4 | chr12:40446052-40846488 | 179.22 | 1.02 | 0.15 | 8.46E-11 | 3.29E-12 |
| 200 | Gpr85 | chr6:13835073-13839848 | 517.80 | 1.01 | 0.09 | 9.31E-26 | 1.45E-27 |
| 201 | Mocos | chr18:24653690-24701556 | 112.93 | 1.01 | 0.18 | 4.44E-07 | 2.69E-08 |
| 202 | Cemip | chr7:83932856-84086505 | 6219.68 | 1.01 | 0.08 | 5.80E-37 | 5.92E-39 |
| 203 | Cthrc1 | chr15:39076931-39087119 | 2779.31 | 1.00 | 0.07 | 7.40E-50 | 4.74E-52 |
| 204 | Per2 | chr1:91415981-91459328 | 293.79 | 1.00 | 0.12 | 1.94E-15 | 5.14E-17 |
| 205 | Ugt1a7c | chr1:88055410-88220002 | 182.14 | -1.01 | 0.14 | 6.94E-11 | 2.68E-12 |
| 206 | Unc13b | chr4:43058983-43264887 | 100.46 | -1.01 | 0.19 | 2.66E-06 | 1.83E-07 |
| 207 | Ablim1 | chr19:57033263-57216032 | 175.09 | -1.02 | 0.15 | 1.06E-10 | 4.18E-12 |
| 208 | Olfm1 | chr2:28193092-28230736 | 291.56 | -1.02 | 0.12 | 2.39E-15 | 6.34E-17 |
| 209 | Slc27a6 | chr18:58556239-58612869 | 267.28 | -1.03 | 0.12 | 6.79E-16 | 1.75E-17 |
| 210 | Slc16a9 | chr10:70245275-70285951 | 64.55 | -1.03 | 0.23 | 8.66E-05 | 7.63E-06 |
| 211 | P2rx3 | chr2:84996551-85035834 | 290.84 | -1.03 | 0.12 | 1.12E-16 | 2.72E-18 |
| 212 | Casp12 | chr9:5345475-5373034 | 599.40 | -1.03 | 0.09 | 1.96E-27 | 2.81E-29 |
| 213 | Car13 | chr3:14641726-14663002 | 709.28 | -1.04 | 0.09 | 4.17E-31 | 4.83E-33 |
| 214 | Mcam | chr9:44134657-44142726 | 100.25 | -1.04 | 0.19 | 9.28E-07 | 5.92E-08 |
| 215 | Il1rl1 | chr1:40429569-40465414 | 45249.83 | -1.04 | 0.05 | 3.07E-97 | 9.13E-100 |
| 216 | Cx3cl1 | chr8:94772179-94782426 | 1202.51 | -1.05 | 0.08 | 6.29E-38 | 6.09E-40 |
| 217 | Notch1 | chr2:26457901-26503822 | 370.77 | -1.06 | 0.12 | 6.67E-18 | 1.52E-19 |
| 218 | Dnmt3l | chr10:78030021-78063622 | 72.82 | -1.06 | 0.22 | 2.39E-05 | 1.88E-06 |

| | | | | | | | |
|---|---|---|---|---|---|---|---|
| 219 | Myo1d | chr11:80482126-80780025 | 2077.25 | -1.06 | 0.08 | 3.75E-40 | 3.34E-42 |
| 220 | Gm16062 | chr11:59810079-59839767 | 81.47 | -1.07 | 0.23 | 4.04E-05 | 3.33E-06 |
| 221 | Foxs1 | chr2:152931897-152933208 | 53.95 | -1.08 | 0.26 | 0.0002694 | 2.63E-05 |
| 222 | Casp4 | chr9:5308848-5336791 | 150.65 | -1.08 | 0.16 | 2.68E-10 | 1.10E-11 |
| 223 | Il11 | chr7:4772377-4782857 | 329.99 | -1.09 | 0.12 | 2.30E-19 | 4.83E-21 |
| 224 | Sox5 | chr6:143828424-144209568 | 522.31 | -1.11 | 0.11 | 2.60E-22 | 4.73E-24 |
| 225 | Gm2115 | chr7:84528953-84578339 | 1624.45 | -1.11 | 0.08 | 1.84E-46 | 1.31E-48 |
| 226 | Cacna1i | chr15:80287237-80398292 | 182.55 | -1.11 | 0.15 | 2.01E-11 | 7.39E-13 |
| 227 | Ptn | chr6:36715662-36811361 | 5861.30 | -1.12 | 0.06 | 1.49E-71 | 6.47E-74 |
| 228 | Cish | chr9:107296688-107301961 | 375.52 | -1.12 | 0.11 | 3.58E-24 | 5.90E-26 |
| 229 | Abcg2 | chr6:58596671-58692451 | 597.70 | -1.16 | 0.09 | 6.59E-33 | 7.38E-35 |
| 230 | Trem2 | chr17:48346400-48352276 | 76.95 | -1.18 | 0.22 | 1.15E-06 | 7.39E-08 |
| 231 | Foxo6 | chr4:120267077-120287261 | 43.66 | -1.18 | 0.30 | 0.0005821 | 6.24E-05 |
| 232 | Mmp9 | chr2:164948218-164955849 | 100.28 | -1.19 | 0.20 | 4.11E-08 | 2.15E-09 |
| 233 | Foxl1 | chr8:121127684-121130644 | 88.97 | -1.21 | 0.21 | 1.18E-07 | 6.52E-09 |
| 234 | Hspb7 | chr4:141420778-141425310 | 44.09 | -1.21 | 0.28 | 0.0001674 | 1.57E-05 |
| 235 | Rcan2 | chr17:43801850-44039516 | 633.15 | -1.21 | 0.10 | 1.76E-30 | 2.09E-32 |
| 236 | Ank3 | chr10:69533707-70027436 | 3106.32 | -1.22 | 0.08 | 1.64E-49 | 1.06E-51 |
| 237 | Txnip | chr3:96555767-96566801 | 1216.77 | -1.25 | 0.07 | 2.84E-62 | 1.39E-64 |
| 238 | Diras2 | chr13:52504374-52530836 | 120.05 | -1.25 | 0.18 | 4.16E-11 | 1.57E-12 |
| 239 | Kcnab1 | chr3:65109367-65378225 | 162.30 | -1.25 | 0.16 | 1.77E-13 | 5.39E-15 |
| 240 | Glipr1 | chr10:111972694-111997264 | 24.28 | -1.28 | 0.38 | 0.0057093 | 0.000808 |
| 241 | Mycn | chr12:12936092-12941836 | 119.46 | -1.29 | 0.18 | 6.85E-11 | 2.64E-12 |
| 242 | Hivep3 | chr4:119814677-120135411 | 388.63 | -1.30 | 0.13 | 1.09E-21 | 2.07E-23 |
| 243 | Smarca1 | chrX:47809369-47892552 | 146.29 | -1.31 | 0.17 | 1.47E-12 | 4.85E-14 |
| 244 | Acot11 | chr4:106733914-106799831 | 117.14 | -1.31 | 0.19 | 4.00E-11 | 1.51E-12 |
| 245 | Fxyd1 | chr7:31051677-31055656 | 45.81 | -1.31 | 0.29 | 5.03E-05 | 4.25E-06 |
| 246 | Cd24a | chr10:43579168-43584265 | 2456.65 | -1.32 | 0.06 | 1.26E-97 | 3.65E-100 |
| 247 | Snap25 | chr2:136713449-136782428 | 111.58 | -1.33 | 0.19 | 1.23E-10 | 4.89E-12 |
| 248 | Dact1 | chr12:71309883-71320107 | 332.60 | -1.39 | 0.12 | 2.94E-29 | 3.81E-31 |
| 249 | Sphk1 | chr11:116531910-116536675 | 101.33 | -1.41 | 0.19 | 5.84E-12 | 2.04E-13 |
| 250 | Lpl | chr8:68880554-68906932 | 348.32 | -1.41 | 0.12 | 1.53E-32 | 1.74E-34 |

| | | | | | | | |
|---|---|---|---|---|---|---|---|
| 251 | Cd34 | chr1:194938820-194976959 | 2459.85 | -1.43 | 0.06 | 3.74E-118 | 6.56E-121 |
| 252 | Eya1 | chr1:14168957-14310199 | 154.61 | -1.44 | 0.18 | 1.07E-14 | 3.00E-16 |
| 253 | Lvrn | chr18:46850038-46905446 | 63.93 | -1.46 | 0.25 | 1.52E-07 | 8.50E-09 |
| 254 | Scg5 | chr2:113776312-113829091 | 46.97 | -1.48 | 0.28 | 2.05E-06 | 1.38E-07 |
| 255 | Fibin | chr2:110360924-110362993 | 40.17 | -1.49 | 0.31 | 1.41E-05 | 1.07E-06 |
| 256 | Grpr | chrX:163513903-163549736 | 80.93 | -1.52 | 0.23 | 7.19E-10 | 3.10E-11 |
| 257 | Sytl2 | chr7:90302354-90410719 | 141.35 | -1.52 | 0.19 | 8.49E-15 | 2.34E-16 |
| 258 | Spp1 | chr5:104435110-104441053 | 34834.73 | -1.55 | 0.06 | 2.62E-166 | 1.60E-169 |
| 259 | AW551984 | chr9:39587395-39604124 | 1687.78 | -1.57 | 0.07 | 7.17E-97 | 2.19E-99 |
| 260 | Fam135b | chr15:71445677-71727838 | 193.24 | -1.59 | 0.15 | 2.96E-25 | 4.72E-27 |
| 261 | Cytip | chr2:58129138-58160122 | 102.57 | -1.62 | 0.21 | 1.26E-13 | 3.81E-15 |
| 262 | 9130024F11F | chr1:56971468-56975196 | 83.34 | -1.63 | 0.22 | 1.99E-12 | 6.71E-14 |
| 263 | Sulf2 | chr2:166073898-166155683 | 233.31 | -1.65 | 0.14 | 8.78E-31 | 1.03E-32 |
| 264 | St3gal1 | chr15:67102874-67176882 | 2141.72 | -1.67 | 0.06 | 1.11E-148 | 1.01E-151 |
| 265 | Catip | chr1:74362107-74369321 | 76.42 | -1.67 | 0.23 | 3.65E-12 | 1.26E-13 |
| 266 | Dkk2 | chr3:132085291-132180304 | 194.47 | -1.67 | 0.15 | 1.40E-25 | 2.20E-27 |
| 267 | Kcnk10 | chr12:98433993-98577940 | 244.88 | -1.68 | 0.14 | 3.72E-32 | 4.26E-34 |
| 268 | Tusc1 | chr4:93334147-93335511 | 59.08 | -1.70 | 0.27 | 5.61E-09 | 2.67E-10 |
| 269 | Rgcc | chr14:79288749-79301635 | 74.98 | -1.73 | 0.23 | 3.58E-12 | 1.23E-13 |
| 270 | H19 | chr7:142575529-142578146 | 493.30 | -1.73 | 0.11 | 1.09E-57 | 5.96E-60 |
| 271 | Cxcl1 | chr5:90891244-90893115 | 26.87 | -1.74 | 0.39 | 7.80E-05 | 6.79E-06 |
| 272 | Tnc | chr4:63959784-64047015 | 17780.03 | -1.76 | 0.08 | 1.50E-115 | 2.85E-118 |
| 273 | Mapkapk3 | chr9:107254926-107289877 | 683.27 | -1.78 | 0.09 | 4.38E-85 | 1.63E-87 |
| 274 | Ptgfr | chr3:151798609-151837528 | 2483.86 | -1.79 | 0.08 | 2.76E-100 | 7.57E-103 |
| 275 | Il18rap | chr1:40515361-40551705 | 144.55 | -1.79 | 0.18 | 4.23E-22 | 7.81E-24 |
| 276 | Has2 | chr15:56665626-56694546 | 1236.24 | -1.80 | 0.08 | 8.75E-102 | 2.27E-104 |
| 277 | 4833427G06I | chr9:51081312-51102078 | 24.21 | -1.82 | 0.40 | 7.88E-05 | 6.88E-06 |
| 278 | Dcn | chr10:97479499-97518162 | 2705.20 | -1.87 | 0.07 | 1.22E-164 | 8.37E-168 |
| 279 | Slc17a7 | chr7:45163920-45176139 | 38.84 | -1.88 | 0.32 | 5.21E-08 | 2.77E-09 |
| 280 | Ramp3 | chr11:6650147-6677475 | 862.80 | -1.89 | 0.09 | 1.90E-102 | 4.63E-105 |
| 281 | Crabp1 | chr9:54764747-54773110 | 1898.86 | -1.92 | 0.08 | 9.81E-124 | 1.42E-126 |
| 282 | Serpine2 | chr1:79794320-79858665 | 930.26 | -1.97 | 0.08 | 3.54E-122 | 5.39E-125 |

| | | | | | | | |
|---|---|---|---|---|---|---|---|
| 283 | Pkp2 | chr16:16213344-16272712 | 39.20 | -2.06 | 0.32 | 4.10E-09 | 1.93E-10 |
| 284 | Lum | chr10:97565500-97572703 | 857.83 | -2.06 | 0.09 | 4.82E-115 | 9.55E-118 |
| 285 | Ptgir | chr7:16906489-16910905 | 44.79 | -2.06 | 0.31 | 6.74E-10 | 2.88E-11 |
| 286 | Tnn | chr1:160085031-160153575 | 184.31 | -2.16 | 0.16 | 6.47E-39 | 6.06E-41 |
| 287 | Cxcl5 | chr5:90759297-90761625 | 188.77 | -2.22 | 0.17 | 2.05E-37 | 2.01E-39 |
| 288 | Fcrlb | chr1:170907272-170912941 | 17.89 | -2.25 | 0.49 | 4.44E-05 | 3.70E-06 |
| 289 | Sfrp2 | chr3:83766320-83774314 | 802.45 | -2.27 | 0.09 | 7.73E-131 | 9.42E-134 |
| 290 | Osbp2 | chr11:3703730-3863903 | 450.55 | -2.39 | 0.11 | 3.06E-98 | 8.63E-101 |
| 291 | Rgag4 | chrX:101849384-102092055 | 57.75 | -2.41 | 0.28 | 6.48E-16 | 1.67E-17 |
| 292 | Gpr149 | chr3:62529962-62605140 | 118.88 | -2.45 | 0.21 | 4.17E-29 | 5.46E-31 |
| 293 | 4930550L24R | chrX:58911460-58920304 | 94.61 | -2.47 | 0.22 | 2.44E-26 | 3.73E-28 |
| 294 | Rtl1 | chr12:109590168-109595403 | 85.78 | -2.56 | 0.25 | 6.54E-23 | 1.15E-24 |
| 295 | Igf1 | chr10:87859055-87937047 | 215.17 | -2.61 | 0.16 | 2.74E-59 | 1.46E-61 |
| 296 | Agtr2 | chrX:21484623-21488833 | 32.11 | -2.98 | 0.41 | 1.28E-11 | 4.57E-13 |
| 297 | Lmcd1 | chr6:112273757-112330423 | 28.66 | -3.13 | 0.45 | 6.75E-11 | 2.59E-12 |
| 298 | Sparcl1 | chr5:104079108-104114088 | 488.70 | -3.39 | 0.13 | 3.70E-159 | 3.10E-162 |
| 299 | Cck | chr9:121489823-121495694 | 41.31 | -3.71 | 0.43 | 1.76E-16 | 4.28E-18 |
| 300 | Vip | chr10:5639217-5647614 | 235.80 | -4.24 | 0.21 | 4.25E-88 | 1.55E-90 |

**Table S1. List of differentially expressed genes in NIH3T3 cells expressing Foxc1 or pLXSH empty vector control.**

**DESeq2 CuffDiff intersected:**

| | | | DESeq2: | | CuffDiff: | |
|---|---|---|---|---|---|---|
| # | Gene | Locus | log2 ratio | q value | log2 ratio | q value |
| 1 | Arhgap36 | chrX:49470449-49500250 | 11.22 | 1.31E-19 | 12.78 | 0.000483 |
| 2 | Anxa10 | chr8:62057041-62123193 | 10.45 | 5.35E-17 | 12.79 | 0.000483 |
| 3 | Kng1 | chr16:23058299-23082078 | 9.29 | 2.81E-13 | 11.47 | 0.000483 |
| 4 | Glra3 | chr8:55940824-56125352 | 7.60 | 1.47E-08 | 10.09 | 0.000483 |
| 5 | E330011O | chr16:78250862-78255454 | 7.53 | 2.17E-08 | 10.94 | 0.000483 |
| 6 | Vnn1 | chr10:23894687-23905343 | 6.39 | 2.52E-121 | 6.42 | 0.000483 |
| 7 | Akr1c18 | chr13:4132626-4150631 | 5.92 | 2.33E-88 | 5.89 | 0.000483 |
| 8 | Wnt2 | chr6:17988939-18030445 | 5.86 | 6.45E-41 | 5.82 | 0.000483 |
| 9 | Il33 | chr19:29925113-29960715 | 5.15 | 5.15E-21 | 5.11 | 0.00287 |
| 10 | Agt | chr8:124556586-124569707 | 4.91 | 1.10E-52 | 4.87 | 0.000483 |
| 11 | Itih2 | chr2:10094590-10130683 | 4.89 | 1.33E-82 | 4.88 | 0.000483 |
| 12 | Foxc1 | chr13:31806645-31810635 | 4.86 | 0.00E+00 | 4.86 | 0.000483 |
| 13 | Col10a1 | chr10:34303611-34418528 | 4.56 | 0.00E+00 | 4.59 | 0.000483 |
| 14 | Il20ra | chr10:19712586-19760053 | 4.12 | 1.98E-53 | 4.1 | 0.000483 |
| 15 | Ogn | chr13:49464058-49652731 | 3.72 | 0.00E+00 | 3.71 | 0.000483 |
| 16 | Apba1 | chr19:23758875-23949597 | 3.60 | 0.00E+00 | 3.58 | 0.000483 |
| 17 | Slc4a4 | chr5:88887259-89239656 | 3.58 | 2.01E-40 | 3.63 | 0.000483 |
| 18 | Itgbl1 | chr14:123660139-123974079 | 3.50 | 2.37E-134 | 3.49 | 0.000483 |
| 19 | Ifit3b | chr19:34607956-34613401 | 3.42 | 2.29E-09 | 3.53 | 0.00494 |
| 20 | Itga1 | chr13:114958080-115101964 | 3.38 | 5.18E-109 | 3.35 | 0.000483 |
| 21 | Camk1g | chr1:193346345-193370282 | 3.33 | 9.12E-23 | 3.29 | 0.000483 |
| 22 | Ifi47 | chr11:49037659-49135387 | 3.31 | 7.74E-11 | 3.3 | 0.000483 |
| 23 | Hid1 | chr11:115347708-115367719 | 3.26 | 1.40E-95 | 3.24 | 0.000483 |
| 24 | Krt15 | chr11:100131758-100135949 | 3.20 | 5.07E-24 | 3.18 | 0.000483 |
| 25 | Pdk4 | chr6:5483350-5496278 | 3.17 | 0.00E+00 | 3.17 | 0.000483 |

| | | | | | | |
|---|---|---|---|---|---|---|
| 26 | Rspo2 | chr15:43020794-43170818 | 3.15 | 4.43E-46 | 3.19 | 0.000483 |
| 27 | Fam20a | chr11:109672925-109722256 | 3.08 | 4.27E-89 | 3.05 | 0.000483 |
| 28 | Nlrp2 | chr7:5298546-5360682 | 3.01 | 4.74E-20 | 2.96 | 0.000483 |
| 29 | C430002N | chr9:96765561-96774397 | 2.89 | 1.30E-07 | 2.84 | 0.000483 |
| 30 | Myh14 | chr7:44605802-44670843 | 2.87 | 2.37E-34 | 2.86 | 0.000483 |
| 31 | Omd | chr13:49464058-49652731 | 2.86 | 4.89E-48 | 2.8 | 0.000483 |
| 32 | Sulf1 | chr1:12692429-12860372 | 2.85 | 1.04E-136 | 2.85 | 0.000483 |
| 33 | Rasgef1b | chr5:99217419-99252927 | 2.82 | 1.21E-24 | 2.81 | 0.000483 |
| 34 | Pgbd5 | chr8:124369048-124433936 | 2.78 | 2.16E-65 | 2.78 | 0.000483 |
| 35 | Scara5 | chr14:65666402-65764826 | 2.76 | 9.57E-27 | 2.76 | 0.000483 |
| 36 | Dio2 | chr12:90724551-90738438 | 2.72 | 9.25E-189 | 2.73 | 0.000483 |
| 37 | Ifit3 | chr19:34583528-34588982 | 2.72 | 7.40E-12 | 2.72 | 0.000483 |
| 38 | Srpx2 | chrX:133908424-133932446 | 2.50 | 1.16E-42 | 2.48 | 0.000483 |
| 39 | Pkia | chr3:7366603-7445365 | 2.50 | 9.69E-80 | 2.52 | 0.000483 |
| 40 | Plscr2 | chr9:92275601-92297752 | 2.46 | 8.60E-21 | 2.46 | 0.000483 |
| 41 | Adra1b | chr11:43774604-43901237 | 2.38 | 6.13E-22 | 2.38 | 0.000483 |
| 42 | Napsa | chr7:44572444-44586846 | 2.36 | 4.66E-07 | 2.53 | 0.000483 |
| 43 | Aldh3a1 | chr11:61208741-61218416 | 2.28 | 3.83E-49 | 2.3 | 0.000483 |
| 44 | Gdf5 | chr2:155941024-155945364 | 2.27 | 3.45E-40 | 2.28 | 0.000483 |
| 45 | Col11a1 | chr3:114030539-114220326 | 2.24 | 9.04E-133 | 2.23 | 0.000483 |
| 46 | 8430408G | chr6:116633007-116673836 | 2.14 | 5.32E-05 | 2.15 | 0.00174 |
| 47 | C1qtnf1 | chr11:118428456-118454995 | 2.11 | 4.34E-43 | 2.12 | 0.000483 |
| 48 | Ror1 | chr4:100095790-100442545 | 2.11 | 9.90E-56 | 2.16 | 0.000483 |
| 49 | Postn | chr3:54361106-54391041 | 2.08 | 2.78E-41 | 2.09 | 0.000483 |
| 50 | Nr3c2 | chr8:76902507-77243639 | 2.08 | 1.74E-11 | 2.15 | 0.000483 |
| 51 | Enpp2 | chr15:54838678-54920146 | 2.06 | 1.45E-17 | 2.06 | 0.000483 |
| 52 | Sorcs2 | chr5:36017180-36398139 | 2.03 | 2.13E-49 | 1.99 | 0.000483 |
| 53 | Tpd52l1 | chr10:31332379-31445921 | 1.95 | 4.23E-05 | 2.14 | 0.000483 |
| 54 | Adamts6 | chr13:104287872-104494763 | 1.94 | 1.21E-23 | 1.93 | 0.000483 |
| 55 | Maf | chr8:115703252-115706894 | 1.88 | 5.68E-27 | 1.87 | 0.000483 |
| 56 | Fhl1 | chrX:56731760-56793346 | 1.88 | 8.50E-07 | 1.86 | 0.000483 |
| 57 | Aspg | chr12:112106682-112127573 | 1.84 | 2.83E-25 | 1.86 | 0.000483 |

| | | | | | | |
|---|---|---|---|---|---|---|
| 58 | Mtss1 | chr15:58941233-59082026 | 1.83 | 1.72E-11 | 1.86 | 0.000483 |
| 59 | Wisp2 | chr2:163820833-163833147 | 1.80 | 2.19E-172 | 1.8 | 0.000483 |
| 60 | Kcnh5 | chr12:74897216-75177332 | 1.77 | 4.04E-13 | 1.77 | 0.000483 |
| 61 | Fam213a | chr14:40993739-41013775 | 1.75 | 1.56E-60 | 1.74 | 0.000483 |
| 62 | Msln | chr17:25748612-25754327 | 1.74 | 1.77E-63 | 1.72 | 0.000483 |
| 63 | Vamp5 | chr6:72368048-72380468 | 1.73 | 4.22E-27 | 1.73 | 0.000483 |
| 64 | Tgfb2 | chr1:186623185-186705992 | 1.71 | 3.13E-50 | 1.71 | 0.000483 |
| 65 | Fjx1 | chr2:102449365-102451792 | 1.70 | 3.26E-10 | 1.69 | 0.000483 |
| 66 | Aspn | chr13:49464058-49652731 | 1.70 | 2.13E-21 | 1.69 | 0.000483 |
| 67 | Sfrp1 | chr8:23411501-23449632 | 1.69 | 1.53E-82 | 1.69 | 0.000483 |
| 68 | Ahr | chr12:35497978-35534989 | 1.68 | 6.54E-114 | 1.68 | 0.000483 |
| 69 | Rnf43 | chr11:87663086-87735539 | 1.68 | 2.23E-29 | 1.61 | 0.000483 |
| 70 | Lhx8 | chr3:154306293-154330560 | 1.65 | 4.48E-16 | 1.65 | 0.000483 |
| 71 | Loxl4 | chr19:42592278-42612806 | 1.63 | 5.94E-102 | 1.64 | 0.000483 |
| 72 | Fst | chr13:114452261-114458951 | 1.59 | 5.16E-16 | 1.6 | 0.000483 |
| 73 | Brinp3 | chr1:146495665-146902472 | 1.59 | 2.76E-22 | 1.55 | 0.000483 |
| 74 | S100a16 | chr3:90541222-90543151 | 1.58 | 0.001333708 | 1.58 | 0.000483 |
| 75 | Adgrg6 | chr10:14402584-14545036 | 1.58 | 1.59E-20 | 1.62 | 0.000483 |
| 76 | 2010111I0 | chr13:62964892-63431745 | 1.58 | 2.32E-60 | 1.58 | 0.000483 |
| 77 | Actr3b | chr5:25760025-25850341 | 1.58 | 6.00E-15 | 1.47 | 0.000483 |
| 78 | Gadd45a | chr6:67035095-67080652 | 1.58 | 7.67E-44 | 1.6 | 0.000483 |
| 79 | Alcam | chr16:52248995-52452997 | 1.58 | 9.17E-118 | 1.58 | 0.000483 |
| 80 | Klf4 | chr4:55527136-55532475 | 1.57 | 1.96E-104 | 1.57 | 0.000483 |
| 81 | Ly6a | chr15:74994876-74998031 | 1.57 | 2.57E-38 | 1.58 | 0.000483 |
| 82 | Cped1 | chr6:21985909-22255606 | 1.57 | 3.19E-64 | 1.57 | 0.000483 |
| 83 | Mfap3l | chr8:60632824-60676731 | 1.55 | 5.50E-41 | 1.54 | 0.000483 |
| 84 | Etl4 | chr2:20289912-20810535 | 1.55 | 2.40E-37 | 1.58 | 0.000483 |
| 85 | Rerg | chr6:137054824-137170496 | 1.55 | 5.63E-13 | 1.49 | 0.000483 |
| 86 | Ctgf | chr10:24595441-24598682 | 1.53 | 5.47E-160 | 1.53 | 0.000483 |
| 87 | Cyp1b1 | chr17:79706952-79715041 | 1.53 | 1.45E-72 | 1.54 | 0.000483 |
| 88 | Gjb4 | chr4:127351085-127354081 | 1.53 | 1.61E-12 | 1.56 | 0.000483 |
| 89 | Rbp1 | chr9:98422960-98446550 | 1.52 | 6.85E-104 | 1.51 | 0.000483 |

| | | | | | | |
|---|---|---|---|---|---|---|
| 90 | Efnb2 | chr8:8617438-8660773 | 1.52 | 5.52E-106 | 1.52 | 0.000483 |
| 91 | Rbp2 | chr9:98490536-98509771 | 1.51 | 0.004968317 | 1.46 | 0.0025 |
| 92 | Nrep | chr18:33437018-33464029 | 1.50 | 1.01E-38 | 1.5 | 0.000483 |
| 93 | 2610035D | chr11:113043905-113201838 | 1.50 | 8.87E-08 | 1.63 | 0.000483 |
| 94 | Thrb | chr14:17660959-18038088 | 1.49 | 7.78E-29 | 1.48 | 0.000483 |
| 95 | Tmem176l | chr6:48833810-48841374 | 1.49 | 4.07E-09 | 1.49 | 0.000483 |
| 96 | Gm14005 | chr2:128298662-128429351 | 1.49 | 8.07E-09 | 1.62 | 0.000483 |
| 97 | Megf6 | chr4:154170712-154275721 | 1.49 | 1.46E-22 | 1.46 | 0.000483 |
| 98 | Dusp4 | chr8:34807609-34819894 | 1.46 | 1.50E-23 | 1.47 | 0.000483 |
| 99 | Cdsn | chr17:35552127-35557180 | 1.43 | 1.26E-73 | 1.43 | 0.000483 |
| 100 | Tle4 | chr19:14448071-14598183 | 1.42 | 4.09E-47 | 1.44 | 0.000483 |
| 101 | Fzd1 | chr5:4753838-4758216 | 1.41 | 2.22E-128 | 1.41 | 0.000483 |
| 102 | Dusp1 | chr17:26505590-26508472 | 1.40 | 1.49E-45 | 1.41 | 0.000483 |
| 103 | Irgm1 | chr11:48865248-48871346 | 1.40 | 5.75E-40 | 1.39 | 0.000483 |
| 104 | Pam | chr1:97821093-98095632 | 1.39 | 1.78E-129 | 1.37 | 0.000483 |
| 105 | Cpeb3 | chr19:37021290-37207471 | 1.38 | 1.17E-22 | 1.37 | 0.000483 |
| 106 | Agpat9 | chr5:100846228-100899102 | 1.36 | 6.53E-07 | 1.33 | 0.000483 |
| 107 | Pde8b | chr13:95024104-95250304 | 1.35 | 2.88E-30 | 1.36 | 0.000483 |
| 108 | Pld1 | chr3:27938679-28133362 | 1.35 | 7.09E-15 | 1.36 | 0.000483 |
| 109 | Gchfr | chr2:119167787-119172389 | 1.34 | 5.77E-24 | 1.35 | 0.000483 |
| 110 | Fam43a | chr16:30599722-30602797 | 1.34 | 1.74E-67 | 1.34 | 0.000483 |
| 111 | Il16 | chr7:83643062-83735490 | 1.34 | 9.64E-09 | 1.33 | 0.000483 |
| 112 | Nrgn | chr9:37544492-37552745 | 1.33 | 0.00107622 | 1.34 | 0.000483 |
| 113 | Zdhhc2 | chr8:40423814-40484842 | 1.33 | 6.35E-30 | 1.32 | 0.000483 |
| 114 | Fam107b | chr2:3713457-3782134 | 1.32 | 6.13E-75 | 1.32 | 0.000483 |
| 115 | Smoc2 | chr17:14279505-14404790 | 1.32 | 1.04E-45 | 1.32 | 0.000483 |
| 116 | Dpysl3 | chr18:43320978-43438286 | 1.32 | 5.06E-94 | 1.32 | 0.000483 |
| 117 | Fam110c | chr12:31073967-31079940 | 1.31 | 7.69E-19 | 1.31 | 0.000483 |
| 118 | Sh3bp4 | chr1:89070461-89153793 | 1.31 | 7.77E-43 | 1.31 | 0.000483 |
| 119 | Pdlim2 | chr14:70164217-70177672 | 1.31 | 1.13E-40 | 1.26 | 0.000483 |
| 120 | Sdpr | chr1:51218385-51333779 | 1.30 | 1.72E-96 | 1.3 | 0.000483 |
| 121 | Tec | chr5:72755717-72868448 | 1.29 | 2.45E-05 | 1.23 | 0.000483 |

| | | | | | | |
|---|---|---|---|---|---|---|
| 122 | Sema5a | chr15:32244812-32696341 | 1.29 | 2.48E-45 | 1.28 | 0.000483 |
| 123 | Ltbp2 | chr12:84783211-84879755 | 1.29 | 6.38E-50 | 1.29 | 0.000483 |
| 124 | Fgd3 | chr13:49263109-49309208 | 1.28 | 1.83E-13 | 1.27 | 0.000483 |
| 125 | Lrrc8d | chr5:105699968-105815215 | 1.27 | 3.97E-21 | 1.25 | 0.000483 |
| 126 | Rgs20 | chr1:4909575-5070285 | 1.27 | 1.47E-28 | 1.29 | 0.000483 |
| 127 | Gpc1 | chr1:92831685-92860196 | 1.27 | 1.10E-100 | 1.27 | 0.000483 |
| 128 | Grk5 | chr19:60889748-61140840 | 1.27 | 1.94E-23 | 1.28 | 0.000483 |
| 129 | Cpe | chr8:64592550-64693040 | 1.25 | 7.27E-121 | 1.26 | 0.000483 |
| 130 | Phldb2 | chr16:45746230-45844378 | 1.24 | 1.41E-95 | 1.24 | 0.000483 |
| 131 | Arhgap31 | chr16:38598342-38713035 | 1.24 | 3.59E-44 | 1.23 | 0.000483 |
| 132 | Gjb3 | chr4:127325234-127330836 | 1.24 | 5.62E-34 | 1.23 | 0.000483 |
| 133 | Amigo2 | chr15:97244073-97385691 | 1.24 | 1.68E-45 | 1.23 | 0.000483 |
| 134 | Dmxl2 | chr9:54365157-54501626 | 1.24 | 6.89E-21 | 1.25 | 0.000483 |
| 135 | Tmem176a | chr6:48841482-48845364 | 1.23 | 0.000653279 | 1.16 | 0.000483 |
| 136 | Slc7a2 | chr8:40862366-40922070 | 1.22 | 1.04E-26 | 1.22 | 0.000483 |
| 137 | Heg1 | chr16:33684465-33768195 | 1.22 | 2.09E-39 | 1.22 | 0.000483 |
| 138 | Capn5 | chr7:98121558-98184799 | 1.22 | 3.58E-14 | 1.2 | 0.000483 |
| 139 | Tgm2 | chr2:158116404-158146392 | 1.22 | 1.81E-23 | 1.21 | 0.000483 |
| 140 | Col12a1 | chr9:79598986-79718722 | 1.21 | 3.69E-07 | 1.21 | 0.000483 |
| 141 | Chrd | chr16:20724453-20742384 | 1.21 | 1.76E-13 | 1.2 | 0.000483 |
| 142 | Slit2 | chr5:47983154-48306282 | 1.20 | 1.05E-53 | 1.2 | 0.000483 |
| 143 | AI606473 | chr3:154330820-154334687 | 1.20 | 7.02E-15 | 1.19 | 0.000483 |
| 144 | Gli1 | chr10:127321963-127341579 | 1.20 | 2.11E-05 | 1.23 | 0.000483 |
| 145 | Lxn | chr3:67430114-67475068 | 1.20 | 3.52E-84 | 1.2 | 0.000483 |
| 146 | Rhou | chr8:123653928-123663880 | 1.20 | 2.60E-70 | 1.2 | 0.000483 |
| 147 | Col5a3 | chr9:20770049-20815034 | 1.20 | 2.24E-30 | 1.18 | 0.000483 |
| 148 | Rab27b | chr18:69979130-70141605 | 1.20 | 8.35E-24 | 1.17 | 0.000483 |
| 149 | Cryab | chr9:50751071-50756635 | 1.20 | 6.19E-45 | 1.26 | 0.000483 |
| 150 | Eva1c | chr16:90830858-90904885 | 1.19 | 6.49E-10 | 1.22 | 0.000483 |
| 151 | Ly6c1 | chr15:75044017-75048837 | 1.19 | 3.60E-09 | 1.2 | 0.000483 |
| 152 | Adrb2 | chr18:62177712-62179981 | 1.19 | 0.000317111 | 1.16 | 0.000483 |
| 153 | Bicc1 | chr10:70925095-71159634 | 1.18 | 1.52E-70 | 1.18 | 0.000483 |

| | | | | | | |
|---|---|---|---|---|---|---|
| 154 | Gstt1 | chr10:75783812-75798584 | 1.18 | 2.78E-06 | 1.14 | 0.000483 |
| 155 | Mmp28 | chr11:83441875-83462961 | 1.16 | 9.98E-06 | 1.14 | 0.000483 |
| 156 | Sepp1 | chr15:3270766-3280508 | 1.16 | 5.91E-24 | 1.17 | 0.000483 |
| 157 | Irs1 | chr1:82233104-82291439 | 1.16 | 2.35E-49 | 1.16 | 0.000483 |
| 158 | Gdpd5 | chr7:99381548-99460984 | 1.16 | 1.62E-06 | 1.15 | 0.000483 |
| 159 | Asb4 | chr6:5383385-5433021 | 1.15 | 1.59E-39 | 1.15 | 0.000483 |
| 160 | Ssfa2 | chr2:79635424-79672964 | 1.14 | 5.25E-36 | 1.15 | 0.000483 |
| 161 | Kalrn | chr16:33969072-34514027 | 1.14 | 6.60E-24 | 1.14 | 0.000483 |
| 162 | Kif13b | chr14:64652530-64806296 | 1.14 | 1.77E-23 | 1.15 | 0.000483 |
| 163 | Anpep | chr7:79821802-79842352 | 1.13 | 1.02E-09 | 1.13 | 0.000483 |
| 164 | Phactr1 | chr13:42680798-43138512 | 1.13 | 1.31E-34 | 1.11 | 0.000483 |
| 165 | Arrb1 | chr7:99535485-99606771 | 1.13 | 2.94E-18 | 1.14 | 0.000483 |
| 166 | Atp10d | chr5:72203328-72298771 | 1.12 | 1.17E-21 | 1.11 | 0.000483 |
| 167 | Fgfbp3 | chr19:36917549-36919599 | 1.12 | 4.70E-17 | 1.09 | 0.000483 |
| 168 | Trim7 | chr11:48826137-48850195 | 1.10 | 0.001515125 | 1.21 | 0.000483 |
| 169 | Angpt1 | chr15:42424666-42676977 | 1.09 | 2.06E-13 | 1.09 | 0.000483 |
| 170 | Col28a1 | chr6:7997807-8192617 | 1.08 | 1.19E-11 | 1.09 | 0.000483 |
| 171 | Pde3a | chr6:141249268-141499351 | 1.08 | 1.73E-14 | 1.09 | 0.000483 |
| 172 | Aox1 | chr1:58029968-58106410 | 1.08 | 3.95E-12 | 1.08 | 0.000483 |
| 173 | C1rl | chr6:124493112-124510643 | 1.07 | 1.57E-05 | 1.07 | 0.000483 |
| 174 | Far1 | chr7:113513833-113570888 | 1.07 | 6.85E-53 | 1.07 | 0.000483 |
| 175 | Ptx3 | chr3:66053557-66296837 | 1.07 | 5.62E-93 | 1.07 | 0.000483 |
| 176 | Ddr1 | chr17:35681566-35704139 | 1.06 | 2.62E-42 | 1.07 | 0.000483 |
| 177 | Dab2 | chr15:6299788-6440709 | 1.06 | 1.24E-61 | 1.06 | 0.000483 |
| 178 | Paqr8 | chr1:20890621-20938756 | 1.06 | 2.60E-07 | 1.06 | 0.000483 |
| 179 | Ppp1r3c | chr19:36731730-36736604 | 1.06 | 5.33E-09 | 1.07 | 0.000483 |
| 180 | Rapgef3 | chr15:97744769-97767666 | 1.06 | 3.89E-13 | 1.04 | 0.000483 |
| 181 | Stx11 | chr10:12939982-12964259 | 1.05 | 1.18E-11 | 1.04 | 0.000483 |
| 182 | Bpgm | chr6:34476355-34505610 | 1.05 | 2.21E-27 | 1.08 | 0.000483 |
| 183 | Thbs1 | chr2:118111921-118127133 | 1.05 | 6.62E-55 | 1.05 | 0.000483 |
| 184 | 9930012K | chr14:70154404-70159502 | 1.05 | 1.33E-11 | 1.05 | 0.000483 |
| 185 | Ngf | chr3:102469918-102521013 | 1.03 | 5.01E-12 | 1.01 | 0.000483 |

| | | | | | | |
|---|---|---|---|---|---|---|
| 186 | Glcci1 | chr6:8509599-8597549 | 1.03 | 4.21E-29 | 1.03 | 0.000483 |
| 187 | Dock4 | chr12:40446052-40846488 | 1.02 | 8.46E-11 | 1.03 | 0.000483 |
| 188 | Gpr85 | chr6:13835073-13839848 | 1.01 | 9.31E-26 | 1 | 0.000483 |
| 189 | Mocos | chr18:24653690-24701556 | 1.01 | 4.44E-07 | 1.01 | 0.000483 |
| 190 | Cemip | chr7:83932856-84086505 | 1.01 | 5.80E-37 | 1.01 | 0.000483 |
| 191 | Cthrc1 | chr15:39076931-39087119 | 1.00 | 7.40E-50 | 1 | 0.000483 |
| 192 | Per2 | chr1:91415981-91459328 | 1.00 | 1.94E-15 | 1.02 | 0.000483 |
| 193 | Ugt1a7c | chr1:88055410-88220002 | -1.01 | 6.94E-11 | -1 | 0.000483 |
| 194 | Unc13b | chr4:43058983-43264887 | -1.01 | 2.66E-06 | -1 | 0.000483 |
| 195 | Ablim1 | chr19:57033263-57216032 | -1.02 | 1.06E-10 | -1.04 | 0.000483 |
| 196 | Slc27a6 | chr18:58556239-58612869 | -1.03 | 6.79E-16 | -1.03 | 0.000483 |
| 197 | Slc16a9 | chr10:70245275-70285951 | -1.03 | 8.66E-05 | -1.04 | 0.000483 |
| 198 | P2rx3 | chr2:84996551-85035834 | -1.03 | 1.12E-16 | -1.06 | 0.000483 |
| 199 | Casp12 | chr9:5345475-5373034 | -1.03 | 1.96E-27 | -1.03 | 0.000483 |
| 200 | Car13 | chr3:14641726-14663002 | -1.04 | 4.17E-31 | -1.05 | 0.000483 |
| 201 | Mcam | chr9:44134657-44142726 | -1.04 | 9.28E-07 | -1.05 | 0.000483 |
| 202 | Il1rl1 | chr1:40429569-40465414 | -1.04 | 3.07E-97 | -1.06 | 0.000483 |
| 203 | Cx3cl1 | chr8:94772179-94782426 | -1.05 | 6.29E-38 | -1.05 | 0.000483 |
| 204 | Notch1 | chr2:26457901-26503822 | -1.06 | 6.67E-18 | -1.05 | 0.000483 |
| 205 | Dnmt3l | chr10:78030021-78063622 | -1.06 | 2.39E-05 | -1.12 | 0.000483 |
| 206 | Myo1d | chr11:80482126-80780025 | -1.06 | 3.75E-40 | -1.07 | 0.000483 |
| 207 | Foxs1 | chr2:152931897-152933208 | -1.08 | 0.000269378 | -1.07 | 0.000483 |
| 208 | Casp4 | chr9:5308848-5336791 | -1.08 | 2.68E-10 | -1.06 | 0.000483 |
| 209 | Il11 | chr7:4772377-4782857 | -1.09 | 2.30E-19 | -1.08 | 0.000483 |
| 210 | Sox5 | chr6:143828424-144209568 | -1.11 | 2.60E-22 | -1.13 | 0.000483 |
| 211 | Gm2115 | chr7:84528953-84578339 | -1.11 | 1.84E-46 | -1.11 | 0.000483 |
| 212 | Cacna1i | chr15:80287237-80398292 | -1.11 | 2.01E-11 | -1.1 | 0.000483 |
| 213 | Ptn | chr6:36715662-36811361 | -1.12 | 1.49E-71 | -1.1 | 0.000483 |
| 214 | Cish | chr9:107296688-107301961 | -1.12 | 3.58E-24 | -1.12 | 0.000483 |
| 215 | Abcg2 | chr6:58596671-58692451 | -1.16 | 6.59E-33 | -1.18 | 0.000483 |
| 216 | Trem2 | chr17:48346400-48352276 | -1.18 | 1.15E-06 | -1.19 | 0.000483 |
| 217 | Foxo6 | chr4:120267077-120287261 | -1.18 | 0.000582099 | -1.16 | 0.000483 |

| | | | | | | |
|---|---|---|---|---|---|---|
| 218 | Mmp9 | chr2:164948218-164955849 | -1.19 | 4.11E-08 | -1.2 | 0.000483 |
| 219 | Foxl1 | chr8:121127684-121130644 | -1.21 | 1.18E-07 | -1.19 | 0.000483 |
| 220 | Hspb7 | chr4:141420778-141425310 | -1.21 | 0.000167378 | -1.23 | 0.000483 |
| 221 | Rcan2 | chr17:43801850-44039516 | -1.21 | 1.76E-30 | -1.22 | 0.000483 |
| 222 | Ank3 | chr10:69533707-70027436 | -1.22 | 1.64E-49 | -1.21 | 0.000483 |
| 223 | Txnip | chr3:96555767-96566801 | -1.25 | 2.84E-62 | -1.25 | 0.000483 |
| 224 | Diras2 | chr13:52504374-52530836 | -1.25 | 4.16E-11 | -1.25 | 0.000483 |
| 225 | Kcnab1 | chr3:65109367-65378225 | -1.25 | 1.77E-13 | -1.24 | 0.000483 |
| 226 | Mycn | chr12:12936092-12941836 | -1.29 | 6.85E-11 | -1.28 | 0.000483 |
| 227 | Hivep3 | chr4:119814677-120135411 | -1.30 | 1.09E-21 | -1.31 | 0.000483 |
| 228 | Smarca1 | chrX:47809369-47892552 | -1.31 | 1.47E-12 | -1.34 | 0.000483 |
| 229 | Acot11 | chr4:106733914-106799831 | -1.31 | 4.00E-11 | -1.32 | 0.000483 |
| 230 | Fxyd1 | chr7:31051677-31055656 | -1.31 | 5.03E-05 | -1.34 | 0.000483 |
| 231 | Cd24a | chr10:43579168-43584265 | -1.32 | 1.26E-97 | -1.32 | 0.000483 |
| 232 | Snap25 | chr2:136713449-136782428 | -1.33 | 1.23E-10 | -1.31 | 0.000483 |
| 233 | Dact1 | chr12:71309883-71320107 | -1.39 | 2.94E-29 | -1.39 | 0.000483 |
| 234 | Sphk1 | chr11:116531910-116536675 | -1.41 | 5.84E-12 | -1.39 | 0.000483 |
| 235 | Lpl | chr8:68880554-68906932 | -1.41 | 1.53E-32 | -1.41 | 0.000483 |
| 236 | Cd34 | chr1:194938820-194976959 | -1.43 | 3.74E-118 | -1.43 | 0.000483 |
| 237 | Eya1 | chr1:14168957-14310199 | -1.44 | 1.07E-14 | -1.44 | 0.000483 |
| 238 | Lvrn | chr18:46850038-46905446 | -1.46 | 1.52E-07 | -1.42 | 0.000483 |
| 239 | Scg5 | chr2:113776312-113829091 | -1.48 | 2.05E-06 | -1.44 | 0.000483 |
| 240 | Fibin | chr2:110360924-110362993 | -1.49 | 1.41E-05 | -1.49 | 0.000483 |
| 241 | Grpr | chrX:163513903-163549736 | -1.52 | 7.19E-10 | -1.51 | 0.000483 |
| 242 | Sytl2 | chr7:90302354-90410719 | -1.52 | 8.49E-15 | -1.55 | 0.000483 |
| 243 | Spp1 | chr5:104435110-104441053 | -1.55 | 2.62E-166 | -1.54 | 0.000483 |
| 244 | AW55198 | chr9:39587395-39604124 | -1.57 | 7.17E-97 | -1.57 | 0.000483 |
| 245 | Fam135b | chr15:71445677-71727838 | -1.59 | 2.96E-25 | -1.59 | 0.000483 |
| 246 | Cytip | chr2:58129138-58160122 | -1.62 | 1.26E-13 | -1.61 | 0.000483 |
| 247 | 9130024F | chr1:56971468-56975196 | -1.63 | 1.99E-12 | -1.38 | 0.000483 |
| 248 | Sulf2 | chr2:166073898-166155683 | -1.65 | 8.78E-31 | -1.65 | 0.000483 |
| 249 | St3gal1 | chr15:67102874-67176882 | -1.67 | 1.11E-148 | -1.67 | 0.000483 |

| | | | | | | |
|---|---|---|---|---|---|---|
| 250 | Catip | chr1:74362107-74369321 | -1.67 | 3.65E-12 | -1.65 | 0.000483 |
| 251 | Dkk2 | chr3:132085291-132180304 | -1.67 | 1.40E-25 | -1.65 | 0.000483 |
| 252 | Kcnk10 | chr12:98433993-98577940 | -1.68 | 3.72E-32 | -1.68 | 0.000483 |
| 253 | Tusc1 | chr4:93334147-93335511 | -1.70 | 5.61E-09 | -1.72 | 0.000483 |
| 254 | Rgcc | chr14:79288749-79301635 | -1.73 | 3.58E-12 | -1.74 | 0.000483 |
| 255 | H19 | chr7:142575529-142578146 | -1.73 | 1.09E-57 | -1.74 | 0.000483 |
| 256 | Cxcl1 | chr5:90891244-90893115 | -1.74 | 7.80E-05 | -1.71 | 0.000483 |
| 257 | Tnc | chr4:63959784-64047015 | -1.76 | 1.50E-115 | -1.76 | 0.000483 |
| 258 | Mapkapk3 | chr9:107254926-107289877 | -1.78 | 4.38E-85 | -1.77 | 0.000483 |
| 259 | Ptgfr | chr3:151798609-151837528 | -1.79 | 2.76E-100 | -1.78 | 0.000483 |
| 260 | Il18rap | chr1:40515361-40551705 | -1.79 | 4.23E-22 | -1.79 | 0.000483 |
| 261 | Has2 | chr15:56665626-56694546 | -1.80 | 8.75E-102 | -1.8 | 0.000483 |
| 262 | 4833427G | chr9:51081312-51102078 | -1.82 | 7.88E-05 | -1.82 | 0.000483 |
| 263 | Dcn | chr10:97479499-97518162 | -1.87 | 1.22E-164 | -1.87 | 0.000483 |
| 264 | Slc17a7 | chr7:45163920-45176139 | -1.88 | 5.21E-08 | -1.85 | 0.000483 |
| 265 | Ramp3 | chr11:6650147-6677475 | -1.89 | 1.90E-102 | -1.89 | 0.000483 |
| 266 | Crabp1 | chr9:54764747-54773110 | -1.92 | 9.81E-124 | -1.92 | 0.000483 |
| 267 | Serpine2 | chr1:79794320-79858665 | -1.97 | 3.54E-122 | -1.97 | 0.000483 |
| 268 | Pkp2 | chr16:16213344-16272712 | -2.06 | 4.10E-09 | -2.07 | 0.000483 |
| 269 | Lum | chr10:97565500-97572703 | -2.06 | 4.82E-115 | -2.06 | 0.000483 |
| 270 | Ptgir | chr7:16906489-16910905 | -2.06 | 6.74E-10 | -2.05 | 0.000483 |
| 271 | Tnn | chr1:160085031-160153575 | -2.16 | 6.47E-39 | -2.16 | 0.000483 |
| 272 | Cxcl5 | chr5:90759297-90761625 | -2.22 | 2.05E-37 | -2.25 | 0.000483 |
| 273 | Fcrlb | chr1:170907272-170912941 | -2.25 | 4.44E-05 | -2.3 | 0.00212 |
| 274 | Sfrp2 | chr3:83766320-83774314 | -2.27 | 7.73E-131 | -2.27 | 0.000483 |
| 275 | Osbp2 | chr11:3703730-3863903 | -2.39 | 3.06E-98 | -2.38 | 0.000483 |
| 276 | Rgag4 | chrX:101849384-102092055 | -2.41 | 6.48E-16 | -2.42 | 0.000483 |
| 277 | Gpr149 | chr3:62529962-62605140 | -2.45 | 4.17E-29 | -2.47 | 0.000483 |
| 278 | 4930550L2 | chrX:58911460-58920304 | -2.47 | 2.44E-26 | -2.46 | 0.000483 |
| 279 | Rtl1 | chr12:109590168-109595403 | -2.56 | 6.54E-23 | -2.53 | 0.000483 |
| 280 | Igf1 | chr10:87859055-87937047 | -2.61 | 2.74E-59 | -2.8 | 0.000483 |
| 281 | Agtr2 | chrX:21484623-21488833 | -2.98 | 1.28E-11 | -2.98 | 0.000483 |

| | | | | | | |
|---|---|---|---|---|---|---|
| 282 | Lmcd1 | chr6:112273757-112330423 | -3.13 | 6.75E-11 | -3.09 | 0.000483 |
| 283 | Sparcl1 | chr5:104079108-104114088 | -3.39 | 3.70E-159 | -3.4 | 0.000483 |
| 284 | Cck | chr9:121489823-121495694 | -3.71 | 1.76E-16 | -3.76 | 0.000483 |
| 285 | Vip | chr10:5639217-5647614 | -4.24 | 4.25E-88 | -4.22 | 0.000483 |
