## Supplemental Table 2 for "Mechanistic insights into transcriptional regulation of ARHGAP36 expression identify a factor predictive of neuroblastoma survival"

### Table S2. Functional profiling of the list of differentially expressed genes from File S1.

Gene set enrichment analysis performed using Enrichr platform [https://maayanlab.cloud/Enrichr/]
Results of annotation using different gene set libraries and databases are shown in separate tabs.

Significant gene set enrichment (FDR-adjusted P-values < 0.05) shown in blue
Gene set enrichment (non-adjusted P-value < 0.05) shown in gray

| **Tab** | **Gene set library / database** |
|---|---|
| *Minimum 3 genes fom the list associated with each term cut-off:* | |
| Elsevier | Elsevier Pathway Collection |
| MSigDB | MSigDB (Molecular Signature Database) hallmark gene sets |
| JL Disease | Jensen Lab Diseases (disease-gene associations mined from literature) |
| JL Tissue | Jensen Lab Tissues (tissue expression database) |
| MGA | Mouse Gene Atlas (expression in mouse cells and tissues) |
| MGI | MGI (Mouse Genome Informatics) mammalian phenotypes level 4 |
| Wiki Mouse | WikiPathways Mouse |
| Wiki Human | WikiPathways Human |
| GO | Gene Ontology Resource (biological process) |
| GWAS | GWAS Catalog (The NHGRI-EBI catalog of human genome-wide association studies) |
| *Minimum 5 genes from the list associated with each term cut-off:* | |
| Localisation | Jensen Lab Compartments (subcellular localisation database) |

| # | Term | Overlap | P-value | Adjusted P-value | Odds Ratio | Combined Score | Genes |
|---|---|---|---|---|---|---|---|
| 1 | Proteins Involved in Arterial Hypertension | 17/255 | 1.63E-07 | 1.24E-04 | 5.19109808 | 81.14031308 | RAMP3;ANGPT1;LPL;IGF1;AHR;ADRB2;NGF;SLC4A4;MMP9;AGT;KNG1;NR3C2;AN |
| 2 | Proteins Involved in Osteoarthritis | 10/86 | 4.02E-07 | 1.53E-04 | 9.39665072 | 138.3773334 | TGFB2;SPP1;COL10A1;PTN;IGF1;MMP9;THBS1;GDF5;ASPN;TGM2 |
| 3 | Proteins Involved in Myocardial Ischemia | 16/252 | 7.27E-07 | 1.70E-04 | 4.90933149 | 69.38928479 | POSTN;ANGPT1;SPHK1;LPL;ARRB1;IGF1;PTN;MMP9;AGT;KNG1;CXCL5;NR3C2;SPP |
| 4 | Proteins Involved in Endometriosis | 16/256 | 8.96E-07 | 1.70E-04 | 4.82651797 | 67.21095763 | IL11;POSTN;TNC;IL16;GSTT1;IGF1;AHR;SULF1;NGF;MMP9;THBS1;DCN;SFRP1;SPP |
| 5 | Proteins Involved in Glaucoma | 9/84 | 3.09E-06 | 4.69E-04 | 8.53913043 | 108.3428497 | EFNB2;SFRP1;FOXC1;CASP12;CYP1B1;NGF;SLC4A4;MMP9;TGM2 |
| 6 | Proteins Involved in Glioma | 16/290 | 4.53E-06 | 5.15E-04 | 4.2202263 | 51.93258584 | DUSP4;TGFB2;DOCK4;NOTCH1;DUSP1;TNC;IGF1;AHR;GRPR;GLI1;NGF;MMP9;KN |
| 7 | Proteins Involved in Dilated Cardiomyopathy | 12/167 | 5.13E-06 | 5.15E-04 | 5.54696916 | 67.5640883 | NOTCH1;TNC;SPP1;AGTR2;ADRB2;IGF1;ADRA1B;MMP9;CRYAB;DCN;AGT;NR3C2 |
| 8 | Proteins Involved in Medulloblastoma | 8/68 | 5.43E-06 | 5.15E-04 | 9.46089049 | 114.6949941 | TGFB2;MYCN;NOTCH1;IRS1;IGF1;GLI1;NGF;MMP9 |
| 9 | Proteins Involved in Osteoporosis | 9/100 | 1.31E-05 | 0.001103583 | 7.03201147 | 79.06772876 | IL11;IL1RL1;IL33;TGFB2;IRS1;DUSP1;SPP1;IGF1;AGT |
| 10 | Proteins with Altered Expression in Asthma | 5/24 | 1.93E-05 | 0.001465623 | 18.5112782 | 200.9379212 | IL33;POSTN;IGF1;NGF;MMP9 |
| 11 | Proteins Involved in Pulmonary Hypertension | 11/165 | 2.57E-05 | 0.001770248 | 5.09932221 | 53.90362147 | ANGPT1;DUSP1;SPP1;CCK;VIP;MMP9;THBS1;KNG1;CX3CL1;DCN;AGT |
| 12 | Proteins Involved in Atherosclerosis | 12/200 | 3.15E-05 | 0.001995429 | 4.56558335 | 47.31766626 | THRB;DUSP1;LPL;CXCL1;PTN;MMP9;THBS1;CX3CL1;AGT;CXCL5;DCN;TGM2 |
| 13 | SNS Related Adrenal Cortex Response | 5/27 | 3.54E-05 | 0.002067694 | 15.9845779 | 163.8159287 | ADRB2;ADRA1B;NGF;AGT;RAPGEF3 |
| 14 | Proteins Involved in Melanoma | 13/238 | 3.96E-05 | 0.002149479 | 4.14003268 | 41.9611943 | POSTN;TGFB2;NOTCH1;MCAM;TNC;IGF1;PTN;GLI1;MMP9;EFNB2;ALCAM;SPP1;T |
| 15 | Epithelial to Mesenchymal Transition in Cancer: Overview | 8/90 | 4.35E-05 | 0.002200089 | 6.91485427 | 69.44732788 | DDR1;FOXC1;NOTCH1;IRS1;TNC;IGF1;GLI1;MMP9 |
| 16 | Proteins Involved in Age-Related Macular Degeneration | 10/148 | 5.33E-05 | 0.002530114 | 5.15862978 | 50.75526311 | IL33;ANGPT1;CAPN5;SPP1;COL10A1;IGF1;NGF;MMP9;CRYAB;ABCG2 |
| 17 | Proteins Involved in Cataract | 8/94 | 5.95E-05 | 0.002654294 | 6.59188985 | 64.14149397 | ALDH3A1;TGFB2;IRS1;IGF1;SLC4A4;MMP9;CRYAB;TGM2 |
| 18 | Genes with Mutations Associated with Glaucoma | 4/16 | 6.42E-05 | 0.002661885 | 23.3724792 | 225.6292821 | FOXC1;CYP1B1;LTBP2;SLC4A4 |
| 19 | Proteins Involved in Neuroblastoma | 12/216 | 6.66E-05 | 0.002661885 | 4.20405085 | 40.42734577 | EFNB2;MYCN;SPP1;IGF1;PTN;NGF;VIP;MMP9;THBS1;CX3CL1;AGT;KNG1 |
| 20 | Proteins Involved in Chronic Obstructive Pulmonary Disease | 10/170 | 1.69E-04 | 0.006412839 | 4.44431818 | 38.60211523 | NOTCH1;SERPINE2;SPP1;GSTT1;CXCL1;PTX3;ADRB2;SLIT2;NGF;MMP9 |
| 21 | IL17F Signaling in Bronchial Epithelial Cell in Asthma | 4/21 | 2.00E-04 | 0.007212779 | 16.4940339 | 140.5189478 | IL11;IL33;IL1RL1;IGF1 |
| 22 | Proteins Involved in Diabetic Nephropathy | 10/175 | 2.14E-04 | 0.007378747 | 4.30853994 | 36.40763987 | NOTCH1;SPP1;TXNIP;AGTR2;IGF1;MMP9;THBS1;DCN;AGT;NR3C2 |
| 23 | Proteins Involved in Glioblastoma | 10/178 | 2.45E-04 | 0.008095993 | 4.23095238 | 35.17145675 | MYCN;NOTCH1;ANGPT1;FST;TNC;IGF1;PTN;GLI1;MMP9;CRYAB |
| 24 | Proteins Involved in Hypothyroidism | 6/63 | 2.73E-04 | 0.00863699 | 7.41671383 | 60.85895084 | THRB;DIO2;IGF1;VIP;PAM;NRGN |
| 25 | Neuroinflammation in Amyotrophic Lateral Sclerosis | 3/10 | 3.19E-04 | 0.009685791 | 29.9513678 | 241.1151384 | IGF1;NGF;CX3CL1 |
| 26 | Glioma Invasion Signaling | 6/75 | 7.02E-04 | 0.020493036 | 6.12311049 | 44.46342853 | DDR1;EFNB2;ANGPT1;COL11A1;IGF1;MMP9 |
| 27 | Proteins with Altered Expression in Cancer Metastases | 7/106 | 8.12E-04 | 0.022837139 | 4.9891723 | 35.50061551 | DDR1;TGFB2;NOTCH1;TNC;SPP1;GLI1;MMP9 |
| 28 | P2RXs -> Synaptic Transmission | 3/14 | 9.28E-04 | 0.023457907 | 19.0560928 | 133.0655786 | SNAP25;P2RX3;SLC17A7 |
| 29 | Proteins Involved in Prolactinoma | 4/31 | 9.38E-04 | 0.023457907 | 10.3798603 | 72.36548233 | IGF1;NGF;VIP;THBS1 |
| 30 | Proteins Involved in Ulcerative Colitis | 8/141 | 9.52E-04 | 0.023457907 | 4.252219 | 29.58159721 | IL11;IL33;SPP1;TXNIP;GSTT1;CD34;MMP9;KNG1 |
| 31 | Ca2+ Flux Regulation | 7/109 | 9.58E-04 | 0.023457907 | 4.84169135 | 33.65247706 | PTGFR;PTGIR;P2RX3;IRS1;ADRB2;ADRA1B;NGF |
| 32 | Proteins Involved in Amyotrophic Lateral Sclerosis | 9/181 | 0.00118899 | 0.028201452 | 3.70506825 | 24.95232927 | PTGFR;CASP4;SCG5;SPP1;CCK;IGF1;MMP9;CRYAB;AGT |
| 33 | Proteins Involved in Vascular Dementia | 3/16 | 0.00139739 | 0.032139972 | 16.1227496 | 105.9772353 | IGF1;AGTR2;NGF |
| 34 | Steroids Induced Cataract | 5/59 | 0.00151523 | 0.033825241 | 6.50165344 | 42.20996438 | MAF;IRS1;IGF1;MMP9;CRYAB |
| 35 | Vitamin C Related Norepinephrine Synthesis | 3/17 | 0.00167905 | 0.035399876 | 14.9703647 | 95.65359163 | ADRB2;ADRA1B;AGT |
| 36 | NTRK -> FOXO/MYCN Signaling | 3/17 | 0.00167905 | 0.035399876 | 14.9703647 | 95.65359163 | MYCN;IRS1;NGF |
| 37 | ARRB1/ARRB2 and GPCRs Internalization | 2/5 | 0.00196689 | 0.040347896 | 46.4358068 | 289.3554382 | GRK5;ARRB1 |
| 38 | Receptors and Adaptor Proteins Activated in Cancer | 7/125 | 0.00211606 | 0.04226561 | 4.18177661 | 25.75220481 | EFNB2;DDR1;IL11;IRS1;COL11A1;IGF1;PTN |
| 39 | Aryl Hydrocarbon Receptor Genomic and non-Genomic Signa | 4/39 | 0.00224284 | 0.043649154 | 8.00406711 | 48.82490153 | ALDH3A1;MAF;CYP1B1;AHR |
| 40 | Renin-Angiotensin-Aldosterone System in Myocardial ischemi | 3/19 | 0.0023428 | 0.044454583 | 13.0977394 | 79.32527373 | AGTR2;AGT;NR3C2 |
| 41 | Proteins Involved in Alzheimer's Disease | 7/130 | 0.00264038 | 0.048651168 | 4.01076212 | 23.81121702 | NOTCH1;LPL;APBA1;IGF1;NGF;MMP9;TGM2 |
| 42 | Thyrotropin Releasing Hormone (TRH) Hypothalamic Release | 3/20 | 0.0027274 | 0.048651168 | 12.3266583 | 72.78158675 | THRB;DIO2;ADRA1B |
| 43 | Neuroblastoma | 7/131 | 0.00275626 | 0.048651168 | 3.9782142 | 23.4471216 | MYCN;NOTCH1;IRS1;IGF1;PTN;GLI1;NGF |
| 44 | IGF1R -> ARRB1/ERK1/3 Signaling | 2/6 | 0.00292261 | 0.049635078 | 34.8250883 | 203.2140946 | ARRB1;IGF1 |
| 45 | Proteins Involved in Polycystic Ovary Syndrome | 6/99 | 0.00294279 | 0.049635078 | 4.53740317 | 26.44578591 | IRS1;FST;ADRB2;IGF1;NGF;AGT |
| 46 | Graves Ophthalmopathy | 4/43 | 0.00321882 | 0.053110462 | 7.18167716 | 41.21379044 | IRS1;HAS2;IL16;IGF1 |
| 47 | Proteins with Altered Expression in Cancer-Associated Sustair | 8/175 | 0.00369362 | 0.059648015 | 3.38061783 | 18.93534274 | EFNB2;DDR1;IL11;NOTCH1;IRS1;IGF1;PTN;GLI1 |
| 48 | Proteins Involved in Obesity | 6/106 | 0.00412629 | 0.065246955 | 4.21827957 | 23.15994372 | NOTCH1;IRS1;SPP1;CCK;IGF1;AGT |
| 49 | Integrins in Cancer Cell Motility, Invasion and Survival | 5/75 | 0.00434552 | 0.067311158 | 5.01147959 | 27.25548673 | IRS1;TNC;SPP1;MMP9;THBS1 |
| 50 | Growth Factor Signaling in Neuroblastoma | 4/47 | 0.00445022 | 0.067554356 | 6.51228999 | 35.26275761 | MYCN;IRS1;IGF1;NGF |

| | | | | | | | |
|---|---|---|---|---|---|---|---|
| 51 | G-Proteins Signaling in Insulin Secretion in beta-Cell | 3/24 | 0.00464324 | 0.069102404 | 9.97669706 | 53.59822762 | CCK;ADRB2;VIP |
| 52 | Hematopoietic Cell Lineage: B-cell (mouse) | 4/49 | 0.00517048 | 0.073880111 | 6.22222222 | 32.7586878 | ALCAM;NOTCH1;ANGPT1;CD34 |
| 53 | Proteins Involved in Chronic Bronchitis | 3/25 | 0.00522142 | 0.073880111 | 9.52272727 | 50.04179619 | CYP1B1;ADRB2;KNG1 |
| 54 | Proteins Involved in non-Alcoholic Fatty Liver Disease | 8/186 | 0.00531813 | 0.073880111 | 3.16991847 | 16.599702 | IRS1;TNC;SPP1;LPL;CXCL1;AHR;IGF1;AGT |
| 55 | Proteins with Altered Expression in Glomerulonephritis | 2/8 | 0.00535363 | 0.073880111 | 23.2143698 | 121.4106951 | AGT;KNG1 |
| 56 | Atopic Dermatitis | 5/81 | 0.00602234 | 0.08162425 | 4.61442669 | 23.5902365 | IL1RL1;IL33;MAF;CDSN;PKP2 |
| 57 | Glutamate Overdose and Aura Effect Caused by Mutations in | 3/27 | 0.00650274 | 0.083693439 | 8.72828014 | 43.95153526 | SNAP25;SLC17A7;SLC4A4 |
| 58 | Glutamate Overdose and Aura Effect Overview | 3/27 | 0.00650274 | 0.083693439 | 8.72828014 | 43.95153526 | SNAP25;SLC17A7;SLC4A4 |
| 59 | Adrenoceptors Physiological Effects | 2/9 | 0.00681876 | 0.083693439 | 19.8970217 | 99.24788618 | ADRB2;ADRA1B |
| 60 | GPCRs Desentization | 2/9 | 0.00681876 | 0.083693439 | 19.8970217 | 99.24788618 | GRK5;ARRB1 |
| 61 | IGF1R -> CEBPA/FOXO1A Signaling | 2/9 | 0.00681876 | 0.083693439 | 19.8970217 | 99.24788618 | IRS1;IGF1 |
| 62 | Muscular Dystrophies | 4/53 | 0.00683662 | 0.083693439 | 5.71312368 | 28.48256147 | IRS1;IGF1;MMP9;AGT |
| 63 | Proteins Involved in Multiple Sclerosis | 4/54 | 0.00730243 | 0.087976835 | 5.59857651 | 27.5424703 | SPP1;VIP;MMP9;CRYAB |
| 64 | Proteins Involved in Glomerulonephritis | 6/121 | 0.00779806 | 0.092480118 | 3.66526414 | 17.79075339 | ITGA1;SPP1;SLIT2;MMP9;CX3CL1;AGT |
| 65 | Vasodilation Activation | 2/10 | 0.0084437 | 0.097752245 | 17.4090106 | 83.11644343 | PTGIR;ADRB2 |
| 66 | Smooth Muscle Cell Dysfunction in Pulmonary Hypertension | 5/88 | 0.0085002 | 0.097752245 | 4.22375215 | 20.13744017 | PTGIR;IRS1;TNC;IGF1;MMP9 |
| 67 | Thyroid Hormones in Adipose Tissue Metabolism | 3/30 | 0.00874753 | 0.099095129 | 7.7572892 | 36.76167061 | THRB;IRS1;LPL |
| 68 | Proteins Involved in Myocarditis | 8/207 | 0.0098713 | 0.106297722 | 2.83235673 | 13.08017307 | IL33;SPP1;TNC;ADRB2;MMP9;THBS1;AGT;NR3C2 |
| 69 | Bone Remodeling in Hyperthyroidism | 4/59 | 0.0099452 | 0.106297722 | 5.08832093 | 23.4605421 | THRB;COL10A1;IGF1;RAPGEF3 |
| 70 | Proteins Involved in Ependymoma | 2/11 | 0.01022363 | 0.106297722 | 15.4738909 | 70.91767359 | TGFB2;TNC |
| 71 | Proteins with Altered Expression in Atopic Dermatitis | 2/11 | 0.01022363 | 0.106297722 | 15.4738909 | 70.91767359 | IL33;CDSN |
| 72 | Aryl Hydrocarbon Receptor/Tryptophan Metabolites Signaling | 2/11 | 0.01022363 | 0.106297722 | 15.4738909 | 70.91767359 | CYP1B1;AHR |
| 73 | Dioxin Role in Endometriosis | 2/11 | 0.01022363 | 0.106297722 | 15.4738909 | 70.91767359 | CYP1B1;AHR |
| 74 | Ovulation Block | 3/32 | 0.01046607 | 0.106653287 | 7.22157007 | 32.92759154 | IRS1;FST;IGF1 |
| 75 | Proteins Involved in Multiple Myeloma | 4/60 | 0.01053886 | 0.106653287 | 4.99720386 | 22.75069847 | PTN;IGF1;CD34;MMP9 |
| 76 | DDR1 -> NF-kB Signaling | 2/12 | 0.01215383 | 0.11826612 | 13.9257951 | 61.4143004 | DDR1;COL11A1 |
| 77 | InsulinR -> CTNNB/FOXA/FOXO Signaling | 2/12 | 0.01215383 | 0.11826612 | 13.9257951 | 61.4143004 | IRS1;FOXO6 |
| 78 | FOXO3 and PTEN Inactivation in Premature Ovarian Failure (M | 2/12 | 0.01215383 | 0.11826612 | 13.9257951 | 61.4143004 | IRS1;IGF1 |
| 79 | Proteins Involved in Polycystic Kidney Disease | 5/97 | 0.01262029 | 0.121250644 | 3.80881211 | 16.65383808 | POSTN;IGF1;AGT;BICC1;SYTL2 |
| 80 | Thyroid Stimulating Hormone (TSH) Secretion in Overt Hypoth | 3/35 | 0.013386 | 0.125770155 | 6.54355053 | 28.22590686 | THRB;CCK;ADRA1B |
| 81 | Androgens in Sebocyte Maturation | 4/65 | 0.01384951 | 0.125770155 | 4.5864302 | 19.6276523 | IRS1;KRT15;IGF1;GLI1 |
| 82 | Cardiomyocyte Hypertrophy | 4/65 | 0.01384951 | 0.125770155 | 4.5864302 | 19.6276523 | IRS1;IGF1;MMP9;AGT |
| 83 | Dopamine Mediated Glutamate Release/Uptake Circle in Neu | 2/13 | 0.01422969 | 0.125770155 | 12.6591712 | 53.83216844 | SNAP25;SLC17A7 |
| 84 | Retinoic Acid Role in Endometriosis | 2/13 | 0.01422969 | 0.125770155 | 12.6591712 | 53.83216844 | RBP1;CYP1B1 |
| 85 | Chemokines Signaling in Atherosclerosis | 2/13 | 0.01422969 | 0.125770155 | 12.6591712 | 53.83216844 | CXCL1;CX3CL1 |
| 86 | Norepinephrine Release Regulation | 3/36 | 0.01445193 | 0.125770155 | 6.34493875 | 26.8830457 | UNC13B;SNAP25;ADRB2 |
| 87 | Tumor Infiltrating Macrophages in Cancer Progression and Im | 3/36 | 0.01445193 | 0.125770155 | 6.34493875 | 26.8830457 | SPP1;MMP9;CX3CL1 |
| 88 | Toll-like Receptor Independent Sterile Inflammation | 4/66 | 0.01458205 | 0.125770155 | 4.51222592 | 19.07752952 | IL1RL1;IL33;TXNIP;CXCL1 |
| 89 | TERT Activation in Cancer | 3/37 | 0.01556462 | 0.132226268 | 6.15801001 | 25.63428796 | MYCN;GLI1;KLF4 |
| 90 | Mast-Cell Activation without Degranulation through IL33/IL1F | 2/14 | 0.0164467 | 0.132226268 | 11.6036514 | 47.66350821 | IL33;IL1RL1 |
| 91 | Genes with Mutations Associated with Atopic Dermatitis | 2/14 | 0.0164467 | 0.132226268 | 11.6036514 | 47.66350821 | IL1RL1;ADRB2 |
| 92 | Glutamate, D-Serine, and ATP Release from Astrocytes | 2/14 | 0.0164467 | 0.132226268 | 11.6036514 | 47.66350821 | SNAP25;SLC17A7 |
| 93 | Parathyroid Glands Development (Mouse Model) | 2/14 | 0.0164467 | 0.132226268 | 11.6036514 | 47.66350821 | EYA1;GLI1 |
| 94 | Medulloblastoma | 5/104 | 0.0166323 | 0.132226268 | 3.53823954 | 14.49407551 | MYCN;NOTCH1;IGF1;GLI1;NGF |
| 95 | VEGFA/NOTCH1/WNT Cross-talk in Blood Vessel Sprouting an | 3/38 | 0.01672427 | 0.132226268 | 5.98176292 | 24.47075955 | FOXC1;NOTCH1;WNT2 |
| 96 | Hematopoietic Cell Lineage: B-cell | 3/38 | 0.01672427 | 0.132226268 | 5.98176292 | 24.47075955 | NOTCH1;AHR;CD34 |
| 97 | Proteins with Altered Expression in Osteoarthritis | 2/15 | 0.01880045 | 0.144136768 | 10.7105192 | 42.56225984 | SPP1;COL10A1 |
| 98 | Longevity Related Drugs | 2/15 | 0.01880045 | 0.144136768 | 10.7105192 | 42.56225984 | AHR;KNG1 |
| 99 | EphrinR -> STAT Signaling | 2/15 | 0.01880045 | 0.144136768 | 10.7105192 | 42.56225984 | EFNB2;NOTCH1 |
| 100 | Proteins with Altered Expression in Endometriosis | 3/41 | 0.02048651 | 0.14822508 | 5.50867861 | 21.4176793 | ANGPT1;RBP1;MMP9 |
| 101 | Steroidogenesis Impairement in Polycystic Ovary Syndrome | 3/41 | 0.02048651 | 0.14822508 | 5.50867861 | 21.4176793 | IRS1;FST;IGF1 |

| # | Term | Overlap | P-value | Adjusted P-value | Odds Ratio | Combined Score | Genes |
|---|---|---|---|---|---|---|---|
| 1 | Epithelial Mesenchymal Transition | 21/200 | 1.15E-12 | 4.81E-11 | 8.68156425 | 238.703215 | POSTN;SERPINE2;GADD45A;LUM;COL11A1;COL12A1;TNC;CXCL1;THBS1;DCN;SFRP1;DAB2;ANPE[illegible] |
| 2 | KRAS Signaling Up | 15/200 | 1.94E-07 | 4.08E-06 | 5.86486486 | 90.63365809 | IL33;SNAP25;TMEM176B;TMEM176A;ANXA10;BPGM;NGF;KLF4;MMP9;MYCN;SPP1;SCG5;SPAR[illegible] |
| 3 | Myogenesis | 11/200 | 1.46E-04 | 0.002043766 | 4.14756884 | 36.6314044 | ABLIM1;NOTCH1;PPP1R3C;PKIA;SPHK1;FST;FXYD1;FHL1;IGF1;TPD52L1;CRYAB |
| 4 | Notch Signaling | 4/32 | 0.00106014 | 0.009165676 | 10.0086426 | 68.55275919 | FZD1;NOTCH1;ARRB1;WNT2 |
| 5 | UV Response Dn | 8/144 | 0.00109115 | 0.009165676 | 4.15778297 | 28.35824756 | PTGFR;DAB2;GRK5;IRS1;DUSP1;COL11A1;HAS2;KALRN |
| 6 | Angiogenesis | 4/36 | 0.0016608 | 0.011625617 | 8.75578292 | 56.04098917 | POSTN;LUM;SPP1;LPL |
| 7 | TNF-alpha Signaling via NF-kB | 9/200 | 0.00235875 | 0.012383451 | 3.33325746 | 20.16494888 | DUSP4;GADD45A;DUSP1;SPHK1;TNC;CXCL1;PTX3;KLF4;FJX1 |
| 8 | Inflammatory Response | 9/200 | 0.00235875 | 0.012383451 | 3.33325746 | 20.16494888 | PTGIR;IL18RAP;SPHK1;HAS2;AHR;VIP;SLC4A4;CX3CL1;SLC7A2 |
| 9 | Estrogen Response Early | 8/200 | 0.00811539 | 0.030986041 | 2.93667268 | 14.13712118 | FOXC1;ABLIM1;CISH;KRT15;TPD52L1;KLF4;SLC7A2;TGM2 |
| 10 | Apical Junction | 8/200 | 0.00811539 | 0.030986041 | 2.93667268 | 14.13712118 | CDSN;IRS1;AMIGO2;SLIT2;ADRA1B;CD34;MMP9;CX3CL1 |
| 11 | p53 Pathway | 8/200 | 0.00811539 | 0.030986041 | 2.93667268 | 14.13712118 | NOTCH1;MAPKAPK3;GADD45A;SPHK1;KIF13B;TXNIP;KLF4;TPD52L1 |
| 12 | Hypoxia | 7/200 | 0.02487275 | 0.074618242 | 2.54694897 | 9.408385063 | LXN;PPP1R3C;DUSP1;GPC1;PAM;DCN;TGM2 |
| 13 | Estrogen Response Late | 7/200 | 0.02487275 | 0.074618242 | 2.54694897 | 9.408385063 | UNC13B;FOXC1;CISH;GJB3;CPE;TPD52L1;KLF4 |
| 14 | Xenobiotic Metabolism | 7/200 | 0.02487275 | 0.074618242 | 2.54694897 | 9.408385063 | ALDH3A1;TGFB2;VNN1;TMEM176B;PDK4;AOX1;IGF1 |
| 15 | Apoptosis | 6/161 | 0.02806368 | 0.078578293 | 2.71383975 | 9.697307159 | TGFB2;GADD45A;LUM;CASP4;TXNIP;DCN |

| |
|---|
| P;GPC1;COL5A3;DPYSL3;SPP1;PTX3;SLIT2;TGM2;CTHRC1 |
| L1;ITGBL1;CPE |

| # | Term | Overlap | P-value | Adjusted P-value | Odds Ratio | Combined Score | Genes |
|---|---|---|---|---|---|---|---|
| 1 | Hypertension | 17/289 | 9.46E-07 | 4.05E-04 | 4.53428172 | 62.8941203 | PTGIR;RAMP3;NOTCH1;ANGPT1;IRS1;LPL;IGF1;ADRB2;MMP9;AGT;KNG1;NR3C2;CTGF;SPP1;AGTR2;VIP;CD34 |
| 2 | Coronary artery disease | 12/205 | 4.02E-05 | 0.008597904 | 4.44616523 | 45.00503089 | PTGIR;ANGPT1;SPP1;LPL;PHACTR1;IGF1;ADRB2;CD34;MMP9;CX3CL1;AGT;KNG1 |
| 3 | Skin cancer | 18/454 | 9.82E-05 | 0.014010598 | 2.98098134 | 27.50984357 | KCNH5;NOTCH1;ANGPT1;COL11A1;TNC;ANK3;KALRN;CPED1;FAM135B;COL5A3;DMXL2;NLRP2;ITGBL1;AOX1;SLC16A9;SLIT2;PHLDB2;ADAMTS6 |
| 4 | Skin disease | 5/41 | 2.79E-04 | 0.029849182 | 9.76140873 | 79.87843493 | GJB4;GJB3;IGF1;CD34;MMP9 |
| 5 | Arthritis | 10/186 | 3.49E-04 | 0.029849182 | 4.03698347 | 32.13956841 | IL11;IL33;DUSP1;SPP1;CXCL1;VIP;MMP9;KNG1;CXCL5;CTGF |
| 6 | Fibroma | 3/13 | 7.37E-04 | 0.052545485 | 20.962766 | 151.2136509 | COL12A1;TNC;CD34 |
| 7 | Kidney cancer | 55/2584 | 0.00140234 | 0.085742874 | 1.62502794 | 10.67580847 | SEMA5A;DOCK4;SERPINE2;IRS1;COL12A1;TNC;GLI1;SLC4A4;NR3C2;IL18RAP;TNN;ANPEP;CAPN5;RSPO2;KIF13B;ENPP2;AOX1;SOX5;TGM2;UNC13B;TLE4;RNF43;POSTN;KCNH5;KCNAB1;ANK3;MMP9;DKK2;CPED1;PKP2;ROR1;PHLDB2;PTGFR;NOTCH1;ITIH2;COL11 [illegible] |
| 8 | Spiradenoma | 2/5 | 0.00196689 | 0.096791483 | 46.4358068 | 289.3554382 | KRT15;DKK2 |
| 9 | Diabetic retinopathy | 4/38 | 0.00203533 | 0.096791483 | 8.23989952 | 51.06343924 | TGFB2;ANGPT1;IGF1;CTGF |
| 10 | Alstrom syndrome | 2/6 | 0.00292261 | 0.108424061 | 34.8250883 | 203.2140946 | RTL1;SLC4A4 |
| 11 | Carcinoma | 184/11318 | 0.00351334 | 0.108424061 | 1.40405181 | 7.934562304 | SEMA5A;RTL1;SERPINE2;IRS1;COL12A1;TNC;LOXL4;AHR;GRPR;SLC4A4;ZDHHC2;IFIT3;HID1;NR3C2;CTGF;FAM107B;ALCAM;IL18RAP;TNN;ANPEP;DPYSL3;CAPN5;C1RL;PDK4;ENPP2;CYP1B1;PDE8B;SOX5;TGM2;PAQR8;UNC13B;TLE4;RNF43;POSTN;KCNH5;KCNK10; [illegible] |
| 12 | Rhinitis | 3/22 | 0.0036078 | 0.108424061 | 11.0279955 | 62.02870512 | NGF;VIP;KNG1 |
| 13 | Alpha thalassemia | 2/7 | 0.00405324 | 0.108424061 | 27.8586572 | 153.4521625 | DNMT3L;SMARCA1 |
| 14 | Xanthinuria | 2/7 | 0.00405324 | 0.108424061 | 27.8586572 | 153.4521625 | MOCOS;AOX1 |
| 15 | cartilage-hair hypoplasia | 2/7 | 0.00405324 | 0.108424061 | 27.8586572 | 153.4521625 | RTL1;COL10A1 |
| 16 | MEDNIK syndrome | 2/7 | 0.00405324 | 0.108424061 | 27.8586572 | 153.4521625 | GJB4;GJB3 |
| 17 | Cerebrovascular disease | 9/225 | 0.00512113 | 0.128932011 | 2.94368961 | 15.52613736 | NOTCH1;ANGPT1;AGTR2;IGF1;NGF;CD34;MMP9;AGT;KNG1 |
| 18 | Irritable bowel syndrome | 4/50 | 0.00555822 | 0.13216209 | 6.08664707 | 31.60477851 | P2RX3;CCK;VIP;TGM2 |
| 19 | Melanoma | 15/505 | 0.00614902 | 0.132655896 | 2.17970522 | 11.09788844 | KCNH5;ANGPT1;COL11A1;TNC;ANK3;KALRN;CPED1;COL5A3;NLRP2;ITGBL1;AOX1;SLC16A9;SLIT2;PHLDB2;ADAMTS6 |
| 20 | Heart conduction disease | 5/82 | 0.00634043 | 0.132655896 | 4.55426716 | 23.04827201 | SFRP2;PDE3A;SLC4A4;ADAMTS6;SULF2 |
| 21 | Clouston syndrome | 2/9 | 0.00681876 | 0.132655896 | 19.8970217 | 99.24788618 | GJB4;GJB3 |
| 22 | Interstitial cystitis | 2/9 | 0.00681876 | 0.132655896 | 19.8970217 | 99.24788618 | P2RX3;ZDHHC2 |
| 23 | Bipolar disorder | 6/124 | 0.00874828 | 0.162794119 | 3.57153271 | 16.92512898 | TLE4;DDR1;ABLIM1;FAM110C;FOXO6;ANK3 |
| 24 | Schizophrenia | 6/126 | 0.00942612 | 0.168099177 | 3.51164875 | 16.37927966 | CACNA1I;ARHGAP31;GPC1;ANK3;ADAMTS6;NRGN |
| 25 | Transient neonatal diabetes mellitus | 2/11 | 0.01022363 | 0.175028513 | 15.4738909 | 70.91767359 | DNMT3L;NLRP2 |
| 26 | Degenerative disc disease | 3/33 | 0.01139338 | 0.187552595 | 6.98049645 | 31.23578541 | GDF5;DCN;ASPN |
| 27 | Spondylolisthesis | 2/12 | 0.01215383 | 0.192660721 | 13.9257951 | 61.4143004 | CDSN;AMIGO2 |
| 28 | Oculodentodigital dysplasia | 2/13 | 0.01422969 | 0.207034986 | 12.6591712 | 53.83216844 | GJB4;GJB3 |
| 29 | Weill-Marchesani syndrome | 2/13 | 0.01422969 | 0.207034986 | 12.6591712 | 53.83216844 | LTBP2;ADAMTS6 |
| 30 | Pain agnosia | 5/101 | 0.01482184 | 0.207034986 | 3.64936756 | 15.36987209 | P2RX3;RGS20;CCK;NGF;KNG1 |
| 31 | Obesity | 5/103 | 0.01601332 | 0.207034986 | 3.57452624 | 14.77828729 | SFRP2;DIRAS2;ENPP2;ST3GAL1;CAMK1G |
| 32 | Sensorineural hearing loss | 5/103 | 0.01601332 | 0.207034986 | 3.57452624 | 14.77828729 | NOTCH1;GJB4;GJB3;COL11A1;MYH14 |
| 33 | Proliferative vitreoretinopathy | 2/14 | 0.0164467 | 0.207034986 | 11.6036514 | 47.66350821 | TGFB2;CTGF |
| 34 | Cryoglobulinemia | 2/14 | 0.0164467 | 0.207034986 | 11.6036514 | 47.66350821 | CDSN;DUSP1 |
| 35 | Adams-Oliver syndrome | 2/15 | 0.01880045 | 0.229059666 | 10.7105192 | 42.56225984 | ARHGAP31;NOTCH1 |
| 36 | Hyperglycemia | 5/108 | 0.0192667 | 0.229059666 | 3.4001387 | 13.42842962 | IRS1;TXNIP;LPL;CCK;IGF1 |
| 37 | Oppositional defiant disorder | 2/16 | 0.02128661 | 0.238400227 | 9.94497728 | 38.28495278 | LXN;OSBP2 |
| 38 | Hyperprolactinemia | 2/16 | 0.02128661 | 0.238400227 | 9.94497728 | 38.28495278 | IGF1;VIP |
| 39 | Common cold | 3/42 | 0.02183532 | 0.238400227 | 5.36715767 | 20.52522532 | IL33;POSTN;MMP9 |
| 40 | Myopia | 4/75 | 0.02228381 | 0.238400227 | 3.9384492 | 14.98144697 | TGFB2;LUM;COL11A1;DCN |
| 41 | Acquired metabolic disease | 10/338 | 0.02364808 | 0.238400227 | 2.14933481 | 8.048126505 | UNC13B;DOCK4;KCNH5;LPL;CCK;IGF1;SULF1;GDF5;NR3C2;LHX8 |
| 42 | Coffin-Siris syndrome | 2/17 | 0.02390095 | 0.238400227 | 9.28150766 | 34.65563604 | CPED1;ADAMTS6 |
| 43 | Brain disease | 4/77 | 0.0242748 | 0.238400227 | 3.83015649 | 14.24173443 | SNAP25;NOTCH1;IGF1;NGF |
| 44 | Ulcerative colitis | 5/115 | 0.02450843 | 0.238400227 | 3.18262987 | 11.80354019 | DDR1;RSPO2;SH3BP4;SORCS2;CAMK1G |
| 45 | Vascular disease | 4/78 | 0.02530935 | 0.240720054 | 3.77820525 | 13.89087888 | TNC;LPL;KALRN;KNG1 |
| 46 | Oligohydramnios | 2/19 | 0.02949779 | 0.274457657 | 8.18873415 | 28.85251413 | SRPX2;AGT |
| 47 | Gestational trophoblastic neoplasm | 2/20 | 0.03247228 | 0.290317637 | 7.73341186 | 26.50525231 | NLRP2;PAM |
| 48 | Brain edema | 2/21 | 0.03555898 | 0.290317637 | 7.32601823 | 24.44371887 | MMP9;KNG1 |
| 49 | Ureteral disease | 2/22 | 0.0387541 | 0.290317637 | 6.95936396 | 22.62154281 | SPP1;CTGF |
| 50 | Hodgkin's lymphoma, nodular sclerosis | 2/22 | 0.0387541 | 0.290317637 | 6.95936396 | 22.62154281 | GJB3;MYH14 |
| 51 | Hyperinsulinism | 3/54 | 0.0416959 | 0.290317637 | 4.10179391 | 13.03284478 | IRS1;LPL;IGF1 |
| 52 | Ankylosis | 2/23 | 0.04205396 | 0.290317637 | 6.62762914 | 21.00164306 | SPP1;GDF5 |
| 53 | Aniridia | 2/24 | 0.04545494 | 0.290317637 | 6.32605204 | 19.55404091 | FOXC1;CYP1B1 |
| 54 | Corneal disease | 2/24 | 0.04545494 | 0.290317637 | 6.32605204 | 19.55404091 | ADAMTS6;NR3C2 |
| 55 | Gastroesophageal reflux disease | 2/24 | 0.04545494 | 0.290317637 | 6.32605204 | 19.55404091 | CCK;PAM |
| 56 | Goiter | 2/24 | 0.04545494 | 0.290317637 | 6.32605204 | 19.55404091 | THRB;IGF1 |
| 57 | Heart disease | 4/94 | 0.04547996 | 0.290317637 | 3.10399367 | 9.592841388 | PKIA;SLIT2;CD34;DCN |
| 58 | Pancreatic cancer | 5/137 | 0.04658653 | 0.290317637 | 2.64921537 | 8.123670353 | MYO1D;RNF43;DAB2;SLIT2;SOX5 |
| 59 | Liver cancer | 14/597 | 0.04777391 | 0.290317637 | 1.69531562 | 5.155922044 | DOCK4;KCNH5;NOTCH1;COL11A1;COL12A1;ITGA1;ANK3;SULF1;KALRN;SYTL2;CACNA1I;DMXL2;HIVEP3;PAM |
| 60 | Hyperaldosteronism | 2/25 | 0.04895351 | 0.290317637 | 6.05069903 | 18.25425842 | AGT;NR3C2 |

| # | Term | Overlap | P-value | Adjusted P-value | Odds Ratio | Combined Score | Genes |
|---|---|---|---|---|---|---|---|
| 1 | Mesenchyme | 22/264 | 3.36E-11 | 2.13E-08 | 6.73107501 | 162.3341407 | FOXC1;POSTN;TGFB2;EYA1;MCAM;CRABP1;TNC;IGF1;FOXL1;GLI1;GDF5;MMP9;DCN;CTGF;ALCAM;ANPEP;SPP1;COL10A1;HAS2;CD34;SOX5;LHX8 |
| 2 | Mesenchymal stem cell | 17/159 | 1.28E-10 | 4.07E-08 | 8.74345701 | 199.1594891 | NOTCH1;ANGPT1;MCAM;ITGA1;TNC;LPL;IGF1;NGF;KLF4;GDF5;MMP9;DCN;ALCAM;ANPEP;SPP1;COL10A1;CD34 |
| 3 | Adult | 38/918 | 3.58E-09 | 7.58E-07 | 3.29283217 | 64.03794374 | SNAP25;NOTCH1;IRS1;FHL1;TNC;LPL;PTN;ADRB2;GLI1;KNG1;NR3C2;CTGF;NRGN;EFNB2;ALCAM;ANPEP;SPP1;SLC17A7;CD34;TGM2;POSTN;TGFB2;ANGPT1;MCAM;FST;CCK;IGF1;NGF;KLF4;GDF5;MMP9;AGT;DCN;MYCN;RBP1;AGTR2;VIP;ABCG2 |
| 4 | Stromal cell | 16/182 | 8.01E-09 | 1.27E-06 | 7.00461325 | 130.579873 | IL11;POSTN;TGFB2;NOTCH1;ANGPT1;MCAM;TNC;CXCL1;IGF1;NGF;MMP9;DCN;ALCAM;ANPEP;SPP1;CD34 |
| 5 | Vasculature | 12/121 | 1.64E-07 | 2.08E-05 | 7.90644218 | 123.5401217 | EFNB2;PTGIR;NOTCH1;ANPEP;MCAM;AGTR2;IGF1;VIP;CD34;MMP9;THBS1;KNG1 |
| 6 | Periodontal ligament | 8/49 | 4.19E-07 | 4.43E-05 | 13.8585894 | 203.5244045 | POSTN;ALCAM;NOTCH1;MCAM;SPP1;GDF5;DCN;ASPN |
| 7 | Plasma cell | 17/286 | 8.19E-07 | 6.87E-05 | 4.58555734 | 64.26704114 | POSTN;ITIH2;LUM;HEG1;ITGA1;TNC;PTN;SEPP1;THBS1;AGT;KNG1;ALCAM;VNN1;TNN;ANPEP;C1RL;SPARCL1 |
| 8 | Cranium | 12/141 | 8.66E-07 | 6.87E-05 | 6.6738223 | 93.16617267 | TGFB2;NOTCH1;OMD;OGN;TNC;SPP1;IGF1;NGF;GDF5;CD34;MMP9;KNG1 |
| 9 | Keloid | 6/29 | 2.86E-06 | 2.02E-04 | 18.4123422 | 235.0019687 | POSTN;TGFB2;TNC;CD34;DCN;CTGF |
| 10 | Neural crest | 11/138 | 4.71E-06 | 2.99E-04 | 6.19196506 | 75.95103238 | FOXC1;MYCN;NOTCH1;EYA1;RBP1;CRABP1;TNC;CHRD;GLI1;NGF;SOX5 |
| 11 | Immune system | 34/1046 | 6.47E-06 | 3.74E-04 | 2.50343291 | 29.91120546 | NOTCH1;TNC;LPL;GSTT1;CXCL1;ADRB2;MSLN;THBS1;STX11;KNG1;CXCL5;CTGF;IL1RL1;ALCAM;ANPEP;SPP1;CYP1B1;CD34;TGM2;IL11;IL33;TGFB2;CDSN;CISH;GADD45A;DUSP1;SPHK1;IL16;IGF1;NGF;MMP9;MAF;MYCN;VIP |
| 12 | Myofibroblast | 8/74 | 1.03E-05 | 5.44E-04 | 8.59818401 | 98.74531648 | POSTN;TGFB2;TNC;SPP1;CD34;MMP9;DCN;CTGF |
| 13 | Ear | 12/186 | 1.53E-05 | 7.50E-04 | 4.93646583 | 54.71821399 | POSTN;DAB2;FOXC1;COL11A1;OGN;SPP1;CYP1B1;CPE;NREP;IGF1;PTN;CTGF |
| 14 | Fiber | 14/263 | 2.67E-05 | 0.001210908 | 4.03864906 | 42.53082049 | NOTCH1;LUM;TNC;CCK;IGF1;NGF;MMP9;KNG1;DCN;P2RX3;SPP1;SLC17A7;VIP;CD34 |
| 15 | Tendon | 9/111 | 3.03E-05 | 0.001284099 | 6.27014066 | 65.22998418 | NOTCH1;LUM;TNC;IGF1;GDF5;CD34;MMP9;DCN;CTGF |
| 16 | Sympathetic nervous system | 4/14 | 3.61E-05 | 0.00143308 | 28.0498221 | 286.9206693 | MYCN;ADRB2;ADRA1B;NGF |
| 17 | Perichondrium | 5/28 | 4.26E-05 | 0.001591769 | 15.2888199 | 153.856302 | TNC;COL10A1;IGF1;GDF5;SOX5 |
| 18 | Alveolar bone | 7/70 | 6.17E-05 | 0.002177103 | 7.85451639 | 76.13393204 | POSTN;NOTCH1;TNC;SPP1;FAM20A;MMP9;LHX8 |
| 19 | Blood vessel endothelium | 9/124 | 7.25E-05 | 0.002423835 | 5.55765595 | 52.97330016 | PTGIR;NOTCH1;SPP1;AGTR2;IGF1;MMP9;KNG1;DCN;ABCG2 |
| 20 | Ectoderm | 9/136 | 1.48E-04 | 0.004389244 | 5.02944197 | 44.36807535 | FZD1;NOTCH1;EYA1;FST;CRABP1;TNC;CHRD;GLI1;NGF |
| 21 | Pulp | 5/36 | 1.49E-04 | 0.004389244 | 11.3387097 | 99.91723689 | P2RX3;MCAM;OMD;SPP1;KNG1 |
| 22 | Leiomyosarcoma cell line | 3/8 | 1.52E-04 | 0.004389244 | 41.9361702 | 368.6684572 | ADRB2;ADRA1B;KNG1 |
| 23 | Neointima | 6/58 | 1.73E-04 | 0.004759555 | 8.13192721 | 70.45695925 | TNC;SPP1;AGTR2;KLF4;CD34;MMP9 |
| 24 | Knee | 8/110 | 1.80E-04 | 0.004759555 | 5.55333758 | 47.88738708 | NOTCH1;COL10A1;IGF1;GDF5;MMP9;KNG1;DCN;ASPN |
| 25 | Mesoderm | 7/86 | 2.28E-04 | 0.005789794 | 6.25862854 | 52.4874105 | EFNB2;FZD1;SFRP2;NOTCH1;FST;CHRD;WNT2 |
| 26 | Femur | 6/63 | 2.73E-04 | 0.006670099 | 7.41671383 | 60.85895084 | NOTCH1;AMIGO2;SPP1;IGF1;ZDHHC2;CD34 |
| 27 | Dental pulp | 5/42 | 3.13E-04 | 0.007371379 | 9.49710425 | 76.62205363 | ALCAM;MCAM;OMD;TNC;SPP1 |
| 28 | Artery | 14/336 | 3.54E-04 | 0.008024302 | 3.11134285 | 24.72491158 | PTGIR;NOTCH1;IRS1;IGF1;ADRB2;MMP9;AGT;KNG1;DCN;NR3C2;CASP12;SPP1;AGTR2;VIP |
| 29 | Bone matrix | 5/44 | 3.91E-04 | 0.008560754 | 9.00915751 | 70.69392362 | OGN;SPP1;IGF1;MMP9;DCN |
| 30 | Tibia | 6/71 | 5.24E-04 | 0.011081527 | 6.50124069 | 49.11624546 | NOTCH1;SPP1;PTN;IGF1;ZDHHC2;GDF5 |
| 31 | Dentin | 6/72 | 5.64E-04 | 0.011561137 | 6.40241121 | 47.88839612 | SDPR;MCAM;OMD;SPP1;MMP9;DCN |
| 32 | Cementum | 4/28 | 6.31E-04 | 0.012525751 | 11.6791222 | 86.05011144 | POSTN;NOTCH1;SPP1;GDF5 |
| 33 | SK-N-SH cell | 3/13 | 7.37E-04 | 0.014174334 | 20.962766 | 151.2136509 | MYCN;NGF;GRPR |
| 34 | Aorta endothelium | 4/30 | 8.26E-04 | 0.015430126 | 10.7796332 | 76.52135984 | NOTCH1;MCAM;THBS1;PHLDB2 |
| 35 | Cancer stem cell | 6/78 | 8.64E-04 | 0.015679488 | 5.86708483 | 41.38453204 | ALCAM;NOTCH1;GLI1;KLF4;CD34;ABCG2 |
| 36 | Molaris | 5/53 | 9.31E-04 | 0.016419225 | 7.31659226 | 51.06548564 | SDPR;MCAM;SPP1;FAM20A;PTN |
| 37 | HUVEC cell | 7/113 | 0.00118354 | 0.019935208 | 4.65803584 | 31.39166255 | ANGPT1;MCAM;CD34;MMP9;THBS1;KNG1;CX3CL1 |
| 38 | Epiphyseal growth plate | 4/33 | 0.00119297 | 0.019935208 | 9.66302614 | 65.04479274 | COL10A1;IGF1;GDF5;SOX5 |
| 39 | Tooth enamel | 5/62 | 0.00189192 | 0.029442595 | 6.1585213 | 38.61494256 | NOTCH1;SDPR;SPP1;FAM20A;SLC4A4 |
| 40 | i10 | 2/5 | 0.00196689 | 0.029442595 | 46.4358068 | 289.3554382 | TNN;ASPN |
| 41 | pa317 | 2/5 | 0.00196689 | 0.029442595 | 46.4358068 | 289.3554382 | CD34;ABCG2 |
| 42 | MSTO-211H cell | 2/5 | 0.00196689 | 0.029442595 | 46.4358068 | 289.3554382 | DIO2;ROR1 |
| 43 | GH4-C1 cell | 3/18 | 0.00199375 | 0.029442595 | 13.9716312 | 86.87193826 | SCG5;CPE;VIP |
| 44 | Satellite cell | 4/39 | 0.00224284 | 0.032368286 | 8.00406711 | 48.82490153 | NOTCH1;GPC1;FST;IGF1 |
| 45 | NG-108-15 cell | 3/19 | 0.0023428 | 0.033059476 | 13.0977394 | 79.32527373 | CACNA1I;AGTR2;KNG1 |
| 46 | Granulation tissue | 3/20 | 0.0027274 | 0.037650024 | 12.3266583 | 72.78158675 | TNC;MMP9;CTGF |
| 47 | Cornea | 2/6 | 0.00292261 | 0.037874624 | 34.8250883 | 203.2140946 | LUM;COL12A1 |
| 48 | Placental membrane | 2/6 | 0.00292261 | 0.037874624 | 34.8250883 | 203.2140946 | IGF1;ABCG2 |
| 49 | CH-1 cell | 2/6 | 0.00292261 | 0.037874624 | 34.8250883 | 203.2140946 | WISP2;SOX5 |
| 50 | Osteocyte | 4/43 | 0.00321882 | 0.040077413 | 7.18167716 | 41.21379044 | ALCAM;SPP1;IGF1;CD34 |
| 51 | Vascular cell | 4/43 | 0.00321882 | 0.040077413 | 7.18167716 | 41.21379044 | MCAM;THBS1;CD34;MMP9 |
| 52 | Cerebrospinal fluid | 4/44 | 0.0035016 | 0.042759949 | 7.00177936 | 39.59180418 | OGN;SPARCL1;CX3CL1;KNG1 |
| 53 | Adductor | 2/7 | 0.00405324 | 0.047401723 | 27.8586572 | 153.4521625 | AMIGO2;PDE8B |
| 54 | Pyloric region | 2/7 | 0.00405324 | 0.047401723 | 27.8586572 | 153.4521625 | SRPX2;CCK |
| 55 | Microvessel | 3/23 | 0.00410566 | 0.047401723 | 10.4760638 | 57.57004088 | MMP9;KNG1;ABCG2 |
| 56 | Hip | 6/107 | 0.00432039 | 0.04899019 | 4.1763015 | 22.73749144 | NOTCH1;AMIGO2;IGF1;ZDHHC2;GDF5;ASPN |
| 57 | Cartilage | 3/24 | 0.00464324 | 0.051630164 | 9.97669706 | 53.59822762 | LUM;TNC;ASPN |
| 58 | Vascular system | 23/894 | 0.00471583 | 0.051630164 | 1.89924716 | 10.17394641 | POSTN;SERPINE2;MCAM;AMIGO2;FHL1;BPGM;THBS1;ASPN;DCN;CTGF;DAB2;ALCAM;SFRP2;SRPX2;PPP1R3C;OGN;SPARCL1;ITGBL1;CPE;FAM43A;PAM;CD34;TGM2 |
| 59 | Prepuce | 4/49 | 0.00517048 | 0.053961203 | 6.22222222 | 32.7586878 | CDSN;DUSP1;PTX3;IFIT3 |
| 60 | Peripheral nervous system | 8/186 | 0.00531813 | 0.053961203 | 3.16991847 | 16.599702 | KCNH5;RAMP3;AGPAT9;MCAM;ARHGAP36;SEPP1;FAM107B;ABCG2 |
| 61 | Clonei | 2/8 | 0.00535363 | 0.053961203 | 23.2143698 | 121.4106951 | LXN;NLRP2 |
| 62 | imr5 | 2/8 | 0.00535363 | 0.053961203 | 23.2143698 | 121.4106951 | EFNB2;MYCN |
| 63 | Mantle muscle | 2/8 | 0.00535363 | 0.053961203 | 23.2143698 | 121.4106951 | GJB4;GJB3 |
| 64 | Submucosa | 4/51 | 0.00596489 | 0.059182927 | 5.95684107 | 30.51013036 | SEMA5A;VIP;CD34;KNG1 |
| 65 | Mast cell | 7/154 | 0.00666184 | 0.064193048 | 3.35183282 | 16.7972413 | IL1RL1;IL33;NGF;VIP;CD34;MMP9;KNG1 |
| 66 | Microglia | 2/9 | 0.00681876 | 0.064193048 | 19.8970217 | 99.24788618 | TREM2;CX3CL1 |
| 67 | Intervertebral disc | 4/53 | 0.00683662 | 0.064193048 | 5.71312368 | 28.48256147 | LUM;NGF;GDF5;DCN |
| 68 | Ganglion | 11/323 | 0.00687422 | 0.064193048 | 2.4966428 | 12.43322537 | PER2;SNAP25;NOTCH1;P2RX3;CPE;SLC17A7;CCK;IGF1;NGF;VIP;KNG1 |
| 69 | Swiss-3T3 cell | 3/28 | 0.00720732 | 0.066328246 | 8.3787234 | 41.32937649 | IGF1;GRPR;KNG1 |
| 70 | Endothelial cell | 4/55 | 0.00778868 | 0.070654491 | 5.48852139 | 26.64722889 | IL33;PTX3;THBS1;TGM2 |
| 71 | AtT-20 cell | 3/29 | 0.00795539 | 0.070714446 | 8.05605565 | 38.94221156 | SCG5;CPE;PAM |
| 72 | Trachea | 22/882 | 0.00801802 | 0.070714446 | 1.83398178 | 8.850913973 | DDR1;SCARA5;MCAM;FHL1;LOXL4;LPL;ASPN;DCN;HID1;CTGF;FAM107B;ALDH3A1;ALCAM;SFRP2;PPP1R3C;OGN;PDK4;CYP1B1;TXNIP;SPARCL1;PAM;TGM2 |
| 73 | Trophoblast | 7/161 | 0.00841835 | 0.072456073 | 3.19833224 | 15.27952357 | IL11;TGFB2;GJB3;IGF1;PAM;MMP9;ABCG2 |
| 74 | Tail fin | 2/10 | 0.0084437 | 0.072456073 | 17.4090106 | 83.11644343 | THRB;CHRD |
| 75 | MG-63 cell | 3/30 | 0.00874753 | 0.073726594 | 7.7572892 | 36.76167061 | SPP1;IGF1;MMP9 |
| 76 | Long bone | 4/57 | 0.00882397 | 0.073726594 | 5.28087021 | 24.98001247 | SPP1;COL10A1;IGF1;GDF5 |
| 77 | BTO:0000868 | 3/31 | 0.00958426 | 0.078025717 | 7.47986322 | 34.76365891 | TNC;SPP1;LPL |
| 78 | Trigeminal nerve | 3/31 | 0.00958426 | 0.078025717 | 7.47986322 | 34.76365891 | P2RX3;SLC17A7;NGF |
| 79 | ATDC-5 cell | 2/11 | 0.01022363 | 0.081150048 | 15.4738909 | 70.91767359 | COL10A1;SOX5 |
| 80 | Sclera | 2/11 | 0.01022363 | 0.081150048 | 15.4738909 | 70.91767359 | LUM;DCN |
| 81 | adipose-derived stem cell | 3/33 | 0.01139338 | 0.089318488 | 6.98049645 | 31.23578541 | MCAM;SPP1;CD34 |
| 82 | Monocytic leukemia cell | 2/12 | 0.01215383 | 0.092984125 | 13.9257951 | 61.4143004 | CASP4;MMP9 |
| 83 | Stellate cell | 2/12 | 0.01215383 | 0.092984125 | 13.9257951 | 61.4143004 | CD34;CTGF |
| 84 | Decidua | 3/34 | 0.01236658 | 0.093485451 | 6.75497598 | 29.67297247 | IL11;TGFB2;FST |
| 85 | Marrow | 5/98 | 0.01314842 | 0.098226412 | 3.76766513 | 16.31946774 | IL11;ANPEP;SPP1;IGF1;CD34 |
| 86 | Secretory cell | 3/35 | 0.013386 | 0.098838468 | 6.54355053 | 28.22590686 | UNC13B;NOTCH1;RAB27B |
| 87 | Outer ear | 2/13 | 0.01422969 | 0.103860408 | 12.6591712 | 53.83216844 | NOTCH1;EYA1 |
| 88 | Pancreatic beta cell | 4/68 | 0.01611955 | 0.116040637 | 4.37077402 | 18.04134302 | SNAP25;MAF;TXNIP;RAPGEF3 |
| 89 | Breast cancer cell line | 8/227 | 0.01644123 | 0.116040637 | 2.57105649 | 10.56180464 | NOTCH1;GADD45A;IRS1;CYP1B1;IGF1;MMP9;ABCG2;WISP2 |
| 90 | Epithelial stem cell | 2/14 | 0.0164467 | 0.116040637 | 11.6036514 | 47.66350821 | KRT15;ABCG2 |
| 91 | Notochord | 3/38 | 0.01672427 | 0.116702331 | 5.98176292 | 24.47075955 | FST;CHRD;GLI1 |
| 92 | Muscle stem cell | 2/15 | 0.01880045 | 0.12836865 | 10.7105192 | 42.56225984 | NOTCH1;IGF1 |
| 93 | Myelocyte | 2/15 | 0.01880045 | 0.12836865 | 10.7105192 | 42.56225984 | ANPEP;CD34 |

| # | Term | Overlap | P-value | Adjusted P-value | Odds Ratio | Combined Score | Genes |
|---|---|---|---|---|---|---|---|
| 1 | osteoblast day21 | 26/264 | 9.89E-15 | 9.10E-13 | 8.21521041 | 264.9159594 | MEGF6;COL11A1;COL12A1;AI606473;LTBP2;THBS1;WISP2;CASP12;SRPX2;CYP1B1;COL10A1;CTHRC1;FZD1;IL[illegible] |
| 2 | osteoblast day14 | 26/301 | 2.20E-13 | 1.01E-11 | 7.0963847 | 206.8165619 | COL11A1;COL12A1;GPR85;LTBP2;AI606473;THBS1;WISP2;CASP12;SRPX2;CYP1B1;COL10A1;HAS2;DACT1;CTH[illegible] |
| 3 | osteoblast day5 | 15/175 | 3.33E-08 | 1.02E-06 | 6.78993056 | 116.9070284 | IL11;MEGF6;COL11A1;TNC;LOXL4;SULF1;THBS1;CTGF;WISP2;EFNB2;TNN;GM2115;HAS2;PTX3;CTHRC1 |
| 4 | umbilical cord | 15/208 | 3.23E-07 | 7.43E-06 | 5.61945884 | 83.98785273 | POSTN;TGFB2;LUM;COL12A1;ITGA1;TNC;MSLN;DKK2;DCN;SULF2;H19;ROR1;ASB4;AGTR2;WNT2 |
| 5 | MEF | 14/300 | 1.10E-04 | 0.002029909 | 3.50948314 | 31.97881468 | TGFB2;MCAM;COL12A1;TNC;LTBP2;SORCS2;THBS1;CTGF;EFNB2;SDPR;DPYSL3;FIBIN;SLIT2;PHLDB2 |
| 6 | 3T3-L1 | 10/181 | 2.81E-04 | 0.004302202 | 4.15608719 | 33.99121343 | CDSN;P2RX3;GM2115;TMEM176A;LOXL4;CYP1B1;PTN;SLIT2;GRPR;CXCL5 |
| 7 | uterus | 8/141 | 9.52E-04 | 0.012514527 | 4.252219 | 29.58159721 | SEMA5A;SMOC2;MAF;RAMP3;ASPG;DIO2;GLI1;TGM2 |
| 8 | C3H 10T1 2 | 8/155 | 0.00174501 | 0.020067586 | 3.8444952 | 24.41637531 | CDSN;PDLIM2;SCARA5;OMD;RSPO2;LOXL4;PTX3;PTN |
| 9 | cornea | 12/427 | 0.01999254 | 0.204368144 | 2.04422084 | 7.997802014 | DDR1;ALDH3A1;ABLIM1;S100A16;KIF13B;TREM2;PHACTR1;SULF1;KLF4;SOX5;TGM2;MFAP3L |
| 10 | mammary gland non-lactating | 7/201 | 0.02546891 | 0.234313931 | 2.53369057 | 9.299396809 | PPP1R3C;IRS1;PKIA;COL5A3;FHL1;PDK4;LMCD1 |
| 11 | mast cells | 13/515 | 0.0339872 | 0.278608233 | 1.82921989 | 6.186003408 | CISH;TUSC1;RAB27B;AHR;SMARCA1;MTSS1;IL1RL1;GLRA3;GRK5;FAM110C;HIVEP3;LY6A;CD34 |
| 12 | mast cells IgE+antigen 1hr | 8/265 | 0.03671558 | 0.278608233 | 2.18662996 | 7.225836824 | FGD3;MYO1D;CISH;FAM110C;C430002N11RIK;2010111I01RIK;MTSS1;ADAMTS6 |
| 13 | retinal pigment epithelium | 12/473 | 0.03936855 | 0.278608233 | 1.83585612 | 5.938605133 | DUSP4;2610035D17RIK;ENPP2;CHRD;SLC16A9;GDPD5;SULF1;VAMP5;RAPGEF3;SOX5;OSBP2;MFAP3L |
| 14 | nucleus accumbens | 10/383 | 0.04863162 | 0.319579247 | 1.88564465 | 5.701211239 | ALCAM;RGS20;PHACTR1;KCNAB1;ANK3;PDE8B;KALRN;CX3CL1;CAMK1G;LHX8 |

| |
|---|
| 33;POSTN;LUM;OMD;IGF1;ASPN;BICC1;H19;SMOC2;SFRP1;SFRP2;OGN;ITGBL1 |
| IRC1;FZD1;POSTN;EYA1;LUM;ASPN;BICC1;H19;SMOC2;SFRP1;SFRP2;OGN;ITGBL1 |

| # | Term | Overlap | P-value | Adjusted P-value | Odds Ratio | Combined Score | Genes |
|---|---|---|---|---|---|---|---|
| 1 | abnormal trabecular bone morphology MP:0000130 | 11/122 | 1.40E-06 | 0.001723419 | 7.09028737 | 95.54791806 | POSTN;SFRP1;THRB;IRS1;COL11A1;PDK4;COL10A1;LTBP2;HIVEP3;IGF1;MMP9 |
| 2 | abnormal glucose homeostasis MP:0002078 | 13/178 | 1.73E-06 | 0.001723419 | 5.66287879 | 75.13902563 | IRS1;TREM2;IGF1;ADRA1B;PPP1R3C;COL5A3;PDK4;SCG5;TXNIP;CPE;AGTR2;RAPGEF3;ABCG2 |
| 3 | dilated renal tubules MP:0002705 | 8/77 | 1.38E-05 | 0.006558252 | 8.22309423 | 91.99656577 | TGFB2;MAF;SPP1;FAM20A;SLC4A4;DCN;MTSS1;AGT |
| 4 | increased bone volume MP:0010875 | 5/23 | 1.55E-05 | 0.006558252 | 19.5406746 | 216.4488258 | PER2;IL20RA;HIVEP3;ADRB2;AGTR2 |
| 5 | abnormal long bone epiphyseal plate proliferative zone MP:0( | 7/58 | 1.79E-05 | 0.006558252 | 9.70856256 | 106.119731 | TLE4;THRB;IRS1;COL11A1;COL10A1;LTBP2;IGF1 |
| 6 | abnormal kidney morphology MP:0002135 | 18/401 | 1.97E-05 | 0.006558252 | 3.40282219 | 36.86505595 | TLE4;FOXC1;TGFB2;SPHK1;CCK;ADRB2;STX11;AGT;BICC1;C1QTNF1;MAF;GRK5;PHACTR1;AGTR2;FAM20A;SLIT2;ADAMTS6;DACT1 |
| 7 | abnormal osteoblast physiology MP:0005006 | 6/45 | 4.06E-05 | 0.010399966 | 10.8497381 | 109.7199677 | IRS1;COL12A1;SPP1;HIVEP3;ADRB2;CTHRC1 |
| 8 | no abnormal phenotype detected MP:0002169 | 44/1654 | 4.17E-05 | 0.010399966 | 2.05309142 | 20.70524331 | SEMA5A;SNAP25;FOXC1;NOTCH1;CRABP1;TNC;CHRD;GRPR;GLI1;PLD1;MSLN;KALRN;EFNB2;IL1RL1;DNMT3L;ABLIM1;GRK5;ENPP2;HAS2;MYH14;SLC17A7;ST3 |
| 9 | hydronephrosis MP:0000519 | 8/93 | 5.51E-05 | 0.012205364 | 6.66978127 | 65.41090866 | FOXC1;TGFB2;HIVEP3;AGTR2;SLIT2;DCN;AGT;DACT1 |
| 10 | decreased body weight MP:0001262 | 37/1329 | 7.26E-05 | 0.014482863 | 2.12739376 | 20.27534634 | SNAP25;RAMP3;THRB;IRS1;COL12A1;FHL1;GPR85;LTBP2;AHR;ADRB2;SLC4A4;KALRN;NR3C2;FAM107B;TNN;COL10A1;SLC17A7;WNT2;TLE4;POSTN;TGFB2;DU |
| 11 | parturition failure MP:0012010 | 3/7 | 9.61E-05 | 0.017420977 | 52.4228723 | 484.9422001 | PTGFR;GADD45A;KALRN |
| 12 | abnormal chemokine level MP:0008721 | 5/34 | 1.13E-04 | 0.018382453 | 12.1219212 | 110.2181904 | DUSP1;ENPP2;SPP1;AHR;MMP9 |
| 13 | decreased leukocyte cell number MP:0000221 | 11/196 | 1.22E-04 | 0.018382453 | 4.23811403 | 38.18568039 | FGD3;TLE4;SFRP1;GADD45A;OGN;KIF13B;SPP1;PHACTR1;CYTIP;ZDHHC2;STX11 |
| 14 | lung inflammation MP:0001861 | 9/134 | 1.32E-04 | 0.018382453 | 5.11043478 | 45.65887135 | IL33;CISH;SCARA5;DUSP1;PTX3;AHR;VIP;MMP9;THBS1 |
| 15 | insulin resistance MP:0005331 | 9/135 | 1.39E-04 | 0.018382453 | 5.06961698 | 45.00700941 | SNAP25;C1QTNF1;PPP1R3C;IRS1;COL5A3;CPE;ADRA1B;RAPGEF3;CTHRC1 |
| 16 | small vertebral body MP:0004670 | 3/8 | 1.52E-04 | 0.018382453 | 41.9361702 | 368.6684572 | FOXC1;MYCN;CHRD |
| 17 | abnormal cochlea morphology MP:0000031 | 6/57 | 1.57E-04 | 0.018382453 | 8.29179844 | 72.64896275 | TGFB2;THRB;EYA1;COL11A1;ROR1;IGF1 |
| 18 | abnormal femur morphology MP:0000559 | 6/59 | 1.90E-04 | 0.019824244 | 7.97808886 | 68.36331594 | FOXC1;TGFB2;IRS1;COL12A1;COL10A1;GDF5 |
| 19 | short radius MP:0004355 | 5/38 | 1.94E-04 | 0.019824244 | 10.6504329 | 91.0508812 | FOXC1;POSTN;TGFB2;SFRP2;GDF5 |
| 20 | abnormal enamel morphology MP:0002576 | 4/21 | 2.00E-04 | 0.019824244 | 16.4940339 | 140.5189478 | POSTN;TNN;LTBP2;FAM20A |
| 21 | increased systemic arterial diastolic blood pressure MP:00061 | 5/39 | 2.20E-04 | 0.019824244 | 10.3366597 | 87.07030286 | IRS1;IGF1;AGTR2;SLC4A4;THBS1 |
| 22 | abnormal laryngeal cartilage morphology MP:0002256 | 3/9 | 2.26E-04 | 0.019824244 | 34.9450355 | 293.4093291 | FOXC1;CHRD;RSPO2 |
| 23 | abnormal long bone epiphyseal plate morphology MP:000305 | 6/61 | 2.29E-04 | 0.019824244 | 7.68719453 | 64.44755359 | TLE4;THRB;COL11A1;COL10A1;MMP9;GDF5 |
| 24 | short metatarsal bones MP:0004635 | 3/10 | 3.19E-04 | 0.024117373 | 29.9513678 | 241.1151384 | SFRP2;RSPO2;GDF5 |
| 25 | abnormal periodontal ligament morphology MP:0003668 | 3/10 | 3.19E-04 | 0.024117373 | 29.9513678 | 241.1151384 | POSTN;TNN;DCN |
| 26 | abnormal cardiovascular development MP:0002925 | 4/24 | 3.43E-04 | 0.024117373 | 14.0177936 | 111.847618 | EFNB2;TGFB2;EYA1;CHRD |
| 27 | short ulna MP:0004359 | 5/43 | 3.51E-04 | 0.024117373 | 9.24671053 | 73.56614753 | FOXC1;POSTN;TGFB2;SFRP2;GDF5 |
| 28 | abnormal lung vasculature morphology MP:0004007 | 5/43 | 3.51E-04 | 0.024117373 | 9.24671053 | 73.56614753 | RSPO2;LPL;VIP;THBS1;WNT2 |
| 29 | abnormal response to cardiac infarction MP:0000343 | 5/43 | 3.51E-04 | 0.024117373 | 9.24671053 | 73.56614753 | POSTN;SPP1;AGTR2;MMP9;TGM2 |
| 30 | decreased cornea thickness MP:0005543 | 5/44 | 3.91E-04 | 0.02514336 | 9.00915751 | 70.69392362 | TGFB2;LUM;NLRP2;GDPD5;CX3CL1 |
| 31 | cleft secondary palate MP:0009890 | 7/94 | 3.94E-04 | 0.02514336 | 5.68080708 | 44.52884099 | TGFB2;EYA1;FST;RSPO2;CHRD;SOX5;LHX8 |
| 32 | decreased circulating sodium level MP:0005634 | 4/25 | 4.03E-04 | 0.02514336 | 13.3496018 | 104.3381494 | DUSP4;TXNIP;SLC4A4;NR3C2 |
| 33 | increased ventricle muscle contractility MP:0005600 | 3/11 | 4.34E-04 | 0.025429306 | 26.206117 | 202.896864 | ARRB1;IGF1;MMP9 |
| 34 | increased interleukin-4 secretion MP:0008699 | 5/45 | 4.35E-04 | 0.025429306 | 8.78348214 | 67.99036303 | DUSP4;CISH;SPP1;COL10A1;MMP9 |
| 35 | abnormal spleen morphology MP:0000689 | 15/385 | 4.46E-04 | 0.025429306 | 2.90465465 | 22.40913253 | TLE4;SPHK1;AHR;ADRB2;GDPD5;CACNA1I;C1QTNF1;MAF;GJB4;MAPKAPK3;GPC1;COL10A1;PHACTR1;ASB4;SLIT2 |
| 36 | short femur MP:0003109 | 7/97 | 4.77E-04 | 0.026151213 | 5.49060751 | 41.98931157 | POSTN;THRB;IRS1;COL12A1;COL10A1;GDF5;MMP9 |
| 37 | decreased susceptibility to injury MP:0005166 | 6/70 | 4.85E-04 | 0.026151213 | 6.6031586 | 50.3909558 | SMOC2;POSTN;PLD1;MMP9;SORCS2;TGM2 |
| 38 | decreased bone resorption MP:0004993 | 4/27 | 5.47E-04 | 0.028724438 | 12.187529 | 91.53833895 | IL20RA;SPP1;HIVEP3;ADRB2 |
| 39 | neonatal lethality, complete penetrance MP:0011087 | 17/481 | 5.76E-04 | 0.029285622 | 2.63177914 | 19.6301419 | RTL1;SNAP25;FOXC1;TGFB2;EYA1;COL11A1;FST;LPL;IGF1;KLF4;SULF2;EFNB2;MAF;MYCN;RSPO2;ROR1;SOX5 |
| 40 | abnormal long bone hypertrophic chondrocyte zone MP:0000 | 5/48 | 5.89E-04 | 0.029285622 | 8.16943522 | 60.76358706 | TLE4;THRB;IRS1;IGF1;MMP9 |
| 41 | abnormal osteoclast differentiation MP:0008396 | 6/73 | 6.08E-04 | 0.029285622 | 6.30653186 | 46.70507692 | IRS1;PDK4;IL20RA;HIVEP3;ADRB2;DKK2 |
| 42 | abnormal blood vessel morphology MP:0001614 | 9/165 | 6.17E-04 | 0.029285622 | 4.08841973 | 30.21750984 | SEMA5A;EFNB2;PER2;SNAP25;MAF;ENPP2;SPP1;HAS2;AHR |
| 43 | abnormal renal tubule epithelium morphology MP:0009640 | 4/28 | 6.31E-04 | 0.029285622 | 11.6791222 | 86.05011144 | DDR1;TGFB2;SLC4A4;DCN |
| 44 | increased physiological sensitivity to xenobiotic MP:0008873 | 7/102 | 6.46E-04 | 0.029288475 | 5.20030292 | 38.19503349 | FST;COL5A3;SPP1;AHR;AGTR2;TGM2;ABCG2 |
| 45 | decreased physiological sensitivity to xenobiotic MP:0008874 | 8/138 | 8.28E-04 | 0.035988183 | 4.35101361 | 30.87809842 | SPP1;CYP1B1;TXNIP;AHR;PDE8B;KALRN;CD34;MMP9 |
| 46 | absent kidney MP:0000520 | 5/52 | 8.53E-04 | 0.035988183 | 7.47264438 | 52.80870386 | TGFB2;EYA1;RSPO2;AGTR2;DACT1 |
| 47 | thick aortic valve cusps MP:0010593 | 3/14 | 9.28E-04 | 0.035988183 | 19.0560928 | 133.0655786 | FOXC1;TGFB2;NOTCH1 |
| 48 | abnormal renal/urinary system morphology MP:0000516 | 3/14 | 9.28E-04 | 0.035988183 | 19.0560928 | 133.0655786 | FOXC1;AGTR2;DACT1 |
| 49 | decreased tympanic ring size MP:0006020 | 3/14 | 9.28E-04 | 0.035988183 | 19.0560928 | 133.0655786 | EYA1;CHRD;RSPO2 |
| 50 | decreased long bone epiphyseal plate size MP:0006396 | 4/31 | 9.38E-04 | 0.035988183 | 10.3798603 | 72.36548233 | TLE4;THRB;IRS1;IGF1 |
| 51 | impaired passive avoidance behavior MP:0004000 | 4/31 | 9.38E-04 | 0.035988183 | 10.3798603 | 72.36548233 | IL33;TNC;ADRA1B;KALRN |
| 52 | pancreatic islet hyperplasia MP:0005491 | 4/31 | 9.38E-04 | 0.035988183 | 10.3798603 | 72.36548233 | SCG5;CPE;THBS1;RAPGEF3 |
| 53 | increased insulin sensitivity MP:0002891 | 8/143 | 0.00104314 | 0.039166205 | 4.18879529 | 28.75824694 | PPP1R3C;RCAN2;SPP1;TXNIP;FOXO6;AGTR2;RAPGEF3;TGM2 |
| 54 | decreased sensitivity to xenobiotic induced morbidity/mortali | 4/32 | 0.00106014 | 0.039166205 | 10.0086426 | 68.55275919 | IL33;CASP4;AHR;MMP9 |
| 55 | decreased body temperature MP:0005534 | 5/56 | 0.00119684 | 0.04341255 | 6.88515406 | 46.32381809 | GRK5;DUSP1;SCG5;AGTR2;STX11 |
| 56 | short mandible MP:0000088 | 5/57 | 0.00129686 | 0.045155176 | 6.75240385 | 44.88871212 | FOXC1;TGFB2;COL11A1;RSPO2;CHRD |
| 57 | abnormal bone mineralization MP:0002896 | 7/115 | 0.00131083 | 0.045155176 | 4.57130962 | 30.34021838 | DDR1;PER2;THRB;IRS1;SPP1;DKK2;SOX5 |
| 58 | abnormal cardiac epithelial to mesenchymal transition MP:00 | 3/16 | 0.00139739 | 0.045155176 | 16.1227496 | 105.9772353 | NOTCH1;MYCN;HAS2 |
| 59 | abnormal kidney blood vessel morphology MP:0000530 | 3/16 | 0.00139739 | 0.045155176 | 16.1227496 | 105.9772353 | FOXC1;AHR;AGT |
| 60 | abnormal hindlimb morphology MP:0000556 | 5/58 | 0.00140291 | 0.045155176 | 6.62466307 | 43.51876205 | FOXC1;COL11A1;RSPO2;THBS1;ADAMTS6 |
| 61 | decreased susceptibility to bacterial infection MP:0002411 | 5/58 | 0.00140291 | 0.045155176 | 6.62466307 | 43.51876205 | MMP28;SPP1;SULF1;MMP9;DCN |
| 62 | decreased length of long bones MP:0004686 | 6/86 | 0.0014392 | 0.045155176 | 5.27822581 | 34.53897211 | SFRP2;COL11A1;LTBP2;SLC4A4;MMP9;GDF5 |
| 63 | hyperglycemia MP:0001559 | 6/86 | 0.0014392 | 0.045155176 | 5.27822581 | 34.53897211 | SNAP25;COL5A3;CPE;ADRA1B;RAPGEF3;TGM2 |
| 64 | abnormal cornea morphology MP:0001312 | 7/117 | 0.00144859 | 0.045155176 | 4.48773708 | 29.33708501 | FOXC1;LUM;ANPEP;OGN;CRYAB;THBS1;DKK2 |
| 65 | respiratory failure MP:0001953 | 8/152 | 0.00154209 | 0.047330401 | 3.92519053 | 25.41409562 | SNAP25;FOXC1;MYCN;FST;COL11A1;RSPO2;IGF1;SOX5 |
| 66 | increased width of hypertrophic chondrocyte zone MP:00034( | 4/36 | 0.0016608 | 0.049546271 | 8.75578292 | 56.04098917 | FOXC1;COL10A1;LTBP2;MMP9 |
| 67 | increased blood urea nitrogen level MP:0005565 | 9/190 | 0.00166396 | 0.049546271 | 3.51921691 | 22.5179029 | EYA1;DUSP1;C1RL;SPP1;TXNIP;RHOU;SLC7A2;STX11;BICC1 |
| 68 | abnormal heart morphology MP:0000266 | 16/485 | 0.00169518 | 0.049733634 | 2.44081769 | 15.5723325 | SPHK1;AHR;ADRB2;THBS1;STX11;BICC1;EFNB2;MAF;RGCC;MAPKAPK3;GRK5;GJB3;PKP2;HAS2;PHACTR1;SLIT2 |
| 69 | increased osteoclast cell number MP:0004984 | 5/61 | 0.00175951 | 0.050129545 | 6.26881378 | 39.76133755 | POSTN;THRB;IRS1;SPP1;DKK2 |

| | | | | | | | |
|---|---|---|---|---|---|---|---|
| 72 | increased trabecular bone thickness MP:0009347 | 4/37 | 0.00184148 | 0.050129545 | 8.4900248 | 53.46325648 | IL20RA;SPP1;HIVEP3;AGTR2 |
| 73 | decreased susceptibility to endotoxin shock MP:0008734 | 5/62 | 0.00189192 | 0.050129545 | 6.1585213 | 38.61494256 | IL33;SPHK1;CASP4;ENPP2;TGM2 |
| 74 | small kidney MP:0002989 | 9/194 | 0.00191885 | 0.050129545 | 3.44242068 | 21.53588691 | TLE4;C1QTNF1;MAF;MYCN;SPHK1;STX11;ADAMTS6;BICC1;SULF2 |
| 75 | hydrometra MP:0009709 | 6/91 | 0.00192353 | 0.050129545 | 4.96647691 | 31.05832249 | GJB4;SPARCL1;FAM20A;SLIT2;FJX1;DACT1 |
| 76 | sternebra fusion MP:0010082 | 3/18 | 0.00199375 | 0.050129545 | 13.9716312 | 86.87193826 | SULF1;ADAMTS6;SULF2 |
| 77 | hydroureter MP:0000536 | 3/18 | 0.00199375 | 0.050129545 | 13.9716312 | 86.87193826 | FOXC1;AGTR2;SLIT2 |
| 78 | increased circulating renin level MP:0003352 | 3/18 | 0.00199375 | 0.050129545 | 13.9716312 | 86.87193826 | SLC4A4;AGT;NR3C2 |
| 79 | abnormal osteoclast physiology MP:0001541 | 5/63 | 0.00203147 | 0.050129545 | 6.05203202 | 37.51651928 | THRB;IRS1;SPP1;HIVEP3;ADRB2 |
| 80 | decreased systemic arterial systolic blood pressure MP:00062 | 4/38 | 0.00203533 | 0.050129545 | 8.23989952 | 51.06343924 | DUSP1;SPP1;ADRA1B;AGT |
| 81 | increased fibroblast proliferation MP:0011703 | 4/38 | 0.00203533 | 0.050129545 | 8.23989952 | 51.06343924 | GADD45A;LUM;SULF1;SULF2 |
| 82 | increased T cell proliferation MP:0005348 | 6/94 | 0.00226799 | 0.053005495 | 4.79643206 | 29.2048053 | DUSP4;RGCC;CISH;GADD45A;TXNIP;MMP9 |
| 83 | increased circulating VLDL cholesterol level MP:0005145 | 3/19 | 0.0023428 | 0.053005495 | 13.0977394 | 79.32527373 | LPL;CPE;TXNIP |
| 84 | increased circulating insulin level MP:0002079 | 8/163 | 0.00239044 | 0.053005495 | 3.64457901 | 21.99969557 | SNAP25;C1QTNF1;PPP1R3C;IRS1;PDK4;CPE;IGF1;ADRA1B |
| 85 | decreased litter size MP:0001935 | 11/283 | 0.00256975 | 0.053005495 | 2.86969998 | 17.11473555 | FZD1;PER2;FST;ASB4;PTX3;AHR;ADRA1B;GDF5;MMP9;THBS1;SULF2 |
| 86 | decreased food intake MP:0011940 | 6/97 | 0.002657 | 0.053005495 | 4.63759896 | 27.50355589 | SNAP25;TNN;RCAN2;FOXO6;CPE;RAPGEF3 |
| 87 | abnormal blood circulation MP:0002128 | 4/41 | 0.00270071 | 0.053005495 | 7.57064538 | 44.77462196 | ENPP2;PKP2;SLC4A4;DACT1 |
| 88 | palatal shelves fail to meet at midline MP:0009888 | 4/41 | 0.00270071 | 0.053005495 | 7.57064538 | 44.77462196 | CHRD;RSPO2;SOX5;LHX8 |
| 89 | abnormal metatarsal bone morphology MP:0003072 | 3/20 | 0.0027274 | 0.053005495 | 12.3266583 | 72.78158675 | FOXC1;GDF5;MMP9 |
| 90 | abnormal thoracic cage shape MP:0010099 | 3/20 | 0.0027274 | 0.053005495 | 12.3266583 | 72.78158675 | TGFB2;LTBP2;SOX5 |
| 91 | hypoalgesia MP:0003043 | 3/20 | 0.0027274 | 0.053005495 | 12.3266583 | 72.78158675 | AGTR2;NGF;MMP9 |
| 92 | increased interleukin-13 secretion MP:0008672 | 3/20 | 0.0027274 | 0.053005495 | 12.3266583 | 72.78158675 | DUSP4;CISH;MMP9 |
| 93 | abnormal uterus morphology MP:0001120 | 7/132 | 0.002876 | 0.053005495 | 3.94618705 | 23.09054581 | FZD1;GJB4;SPARCL1;PHACTR1;AHR;FAM20A;FJX1 |
| 94 | small mandibular condyloid process MP:0030258 | 2/6 | 0.00292261 | 0.053005495 | 34.8250883 | 203.2140946 | TGFB2;RSPO2 |
| 95 | small Meckel's cartilage MP:0030026 | 2/6 | 0.00292261 | 0.053005495 | 34.8250883 | 203.2140946 | FOXC1;COL11A1 |
| 96 | tubular nephritis MP:0002704 | 2/6 | 0.00292261 | 0.053005495 | 34.8250883 | 203.2140946 | MAF;DCN |
| 97 | abnormal lacrimal gland physiology MP:0001348 | 2/6 | 0.00292261 | 0.053005495 | 34.8250883 | 203.2140946 | FOXC1;THBS1 |
| 98 | abnormal renal/urinary system physiology MP:0005502 | 2/6 | 0.00292261 | 0.053005495 | 34.8250883 | 203.2140946 | AGTR2;AGT |
| 99 | absent eye anterior chamber MP:0010709 | 2/6 | 0.00292261 | 0.053005495 | 34.8250883 | 203.2140946 | FOXC1;TGFB2 |
| 100 | absent Schlemm's canal MP:0004226 | 2/6 | 0.00292261 | 0.053005495 | 34.8250883 | 203.2140946 | FOXC1;CYP1B1 |
| 101 | blind ureter MP:0011797 | 2/6 | 0.00292261 | 0.053005495 | 34.8250883 | 203.2140946 | FOXC1;DACT1 |
| 102 | bruising MP:0009275 | 2/6 | 0.00292261 | 0.053005495 | 34.8250883 | 203.2140946 | LUM;SCG5 |
| 103 | decreased osteoblast proliferation MP:0030439 | 2/6 | 0.00292261 | 0.053005495 | 34.8250883 | 203.2140946 | IRS1;CTHRC1 |
| 104 | decreased posterior semicircular canal size MP:0003164 | 2/6 | 0.00292261 | 0.053005495 | 34.8250883 | 203.2140946 | EFNB2;EYA1 |
| 105 | delayed inner ear development MP:0004664 | 2/6 | 0.00292261 | 0.053005495 | 34.8250883 | 203.2140946 | THRB;IGF1 |
| 106 | growth retardation of incisors MP:0005359 | 2/6 | 0.00292261 | 0.053005495 | 34.8250883 | 203.2140946 | FST;RSPO2 |
| 107 | hypoplastic trabecular meshwork MP:0004223 | 2/6 | 0.00292261 | 0.053005495 | 34.8250883 | 203.2140946 | FOXC1;CYP1B1 |
| 108 | increased cardiac cell glucose uptake MP:0030018 | 2/6 | 0.00292261 | 0.053005495 | 34.8250883 | 203.2140946 | THRB;TXNIP |
| 109 | increased urine uric acid level MP:0009810 | 2/6 | 0.00292261 | 0.053005495 | 34.8250883 | 203.2140946 | AHR;ABCG2 |
| 110 | micromelia MP:0008736 | 2/6 | 0.00292261 | 0.053005495 | 34.8250883 | 203.2140946 | COL11A1;GDF5 |
| 111 | abnormal placenta vasculature MP:0003231 | 5/69 | 0.00303143 | 0.054483806 | 5.48297991 | 31.79427016 | MAF;NOTCH1;ASB4;SLIT2;WNT2 |
| 112 | abnormal physiological neovascularization MP:0003710 | 3/21 | 0.00314871 | 0.056086429 | 11.641253 | 67.0624861 | SPP1;MMP9;THBS1 |
| 113 | trabecula carnea hypoplasia MP:0000295 | 4/44 | 0.0035016 | 0.060996215 | 7.00177936 | 39.59180418 | TGFB2;NOTCH1;MYCN;PKP2 |
| 114 | increased systemic arterial blood pressure MP:0002842 | 4/44 | 0.0035016 | 0.060996215 | 7.00177936 | 39.59180418 | ADRB2;IGF1;AGTR2;AGT |
| 115 | abnormal circulating insulin level MP:0001560 | 3/22 | 0.0036078 | 0.060996215 | 11.0279955 | 62.02870512 | SCG5;TXNIP;AGTR2 |
| 116 | abnormal endochondral bone ossification MP:0008272 | 3/22 | 0.0036078 | 0.060996215 | 11.0279955 | 62.02870512 | COL11A1;COL10A1;GDF5 |
| 117 | abnormal tracheal cartilage morphology MP:0003120 | 3/22 | 0.0036078 | 0.060996215 | 11.0279955 | 62.02870512 | FOXC1;COL11A1;RSPO2 |
| 118 | early reproductive senescence MP:0008995 | 3/22 | 0.0036078 | 0.060996215 | 11.0279955 | 62.02870512 | PER2;IL33;FST |
| 119 | enlarged kidney MP:0003068 | 7/138 | 0.00368027 | 0.061259134 | 3.76429238 | 21.09799067 | FOXC1;MAF;C1QTNF1;PGBD5;ADRB2;SLIT2;BICC1 |
| 120 | short tibia MP:0002764 | 9/214 | 0.00369886 | 0.061259134 | 3.10339343 | 17.37816793 | POSTN;THRB;IRS1;RCAN2;RSPO2;FAM20A;GDF5;MMP9;SLC7A2 |
| 121 | increased respiratory quotient MP:0010378 | 4/45 | 0.00380081 | 0.061259134 | 6.83065706 | 38.06412274 | DUSP1;SPP1;CPE;RAPGEF3 |
| 122 | abnormal canal of Schlemm morphology MP:0005204 | 2/7 | 0.00405324 | 0.061259134 | 27.8586572 | 153.4521625 | FOXC1;CYP1B1 |
| 123 | short basicranium MP:0030420 | 2/7 | 0.00405324 | 0.061259134 | 27.8586572 | 153.4521625 | FOXC1;LTBP2 |
| 124 | abnormal common crus morphology MP:0004922 | 2/7 | 0.00405324 | 0.061259134 | 27.8586572 | 153.4521625 | EFNB2;EYA1 |
| 125 | thick pulmonary valve cusps MP:0010605 | 2/7 | 0.00405324 | 0.061259134 | 27.8586572 | 153.4521625 | FOXC1;TGFB2 |
| 126 | abnormal foramen magnum morphology MP:0010941 | 2/7 | 0.00405324 | 0.061259134 | 27.8586572 | 153.4521625 | FOXC1;LTBP2 |
| 127 | abnormal juxtaglomerular apparatus morphology MP:000282 | 2/7 | 0.00405324 | 0.061259134 | 27.8586572 | 153.4521625 | AGT;NR3C2 |
| 128 | abnormal kidney arterial blood vessel morphology MP:00113 | 2/7 | 0.00405324 | 0.061259134 | 27.8586572 | 153.4521625 | FOXC1;FAM20A |
| 129 | abnormal ovary physiology MP:0003507 | 2/7 | 0.00405324 | 0.061259134 | 27.8586572 | 153.4521625 | IL33;AHR |
| 130 | abnormal pupillary reflex MP:0002638 | 2/7 | 0.00405324 | 0.061259134 | 27.8586572 | 153.4521625 | MAF;EVA1C |
| 131 | calcified aortic valve MP:0006116 | 2/7 | 0.00405324 | 0.061259134 | 27.8586572 | 153.4521625 | NOTCH1;SPP1 |
| 132 | decreased gastrocnemius weight MP:0009422 | 2/7 | 0.00405324 | 0.061259134 | 27.8586572 | 153.4521625 | FST;SLC4A4 |
| 133 | abnormal interleukin level MP:0008751 | 4/46 | 0.00411687 | 0.061292195 | 6.66768344 | 36.62333352 | DUSP1;TREM2;AHR;ZDHHC2 |
| 134 | delayed endochondral bone ossification MP:0003419 | 4/46 | 0.00411687 | 0.061292195 | 6.66768344 | 36.62333352 | FOXC1;SFRP2;IGF1;GDF5 |
| 135 | abnormal liver morphology MP:0000598 | 11/306 | 0.00463406 | 0.068112301 | 2.64283063 | 14.20342466 | TLE4;C1QTNF1;MAF;MYCN;GJB4;GPC1;SPP1;LPL;AHR;THBS1;BICC1 |
| 136 | abnormal epiphyseal plate morphology MP:0006395 | 3/24 | 0.00464324 | 0.068112301 | 9.97669706 | 53.59822762 | POSTN;THRB;COL11A1 |
| 137 | decreased susceptibility to diet-induced obesity MP:0005659 | 6/109 | 0.00472872 | 0.068859778 | 4.09479069 | 21.92392474 | FOXS1;C1QTNF1;DUSP1;COL5A3;RCAN2;AGTR2 |
| 138 | decreased epididymal fat pad weight MP:0009289 | 4/48 | 0.00480129 | 0.069139601 | 6.36395988 | 33.97636268 | DUSP1;RCAN2;AGT;RAPGEF3 |
| 139 | enlarged spleen MP:0000691 | 14/444 | 0.00481725 | 0.069139601 | 2.3169141 | 12.36201799 | TLE4;SPHK1;AHR;ADRB2;BPGM;SLC4A4;MTSS1;C1QTNF1;MAF;GJB4;MAPKAPK3;GPC1;PHACTR1;SLIT2 |
| 140 | abnormal neuron differentiation MP:0009937 | 6/111 | 0.00516475 | 0.072656423 | 4.01638505 | 21.14987745 | FOXC1;SFRP2;FST;DPYSL3;SMARCA1;GDPD5 |
| 141 | increased liver glycogen level MP:0010400 | 3/25 | 0.00522142 | 0.072656423 | 9.52272727 | 50.04179619 | SCG5;ADRA1B;CTHRC1 |

| # | Term | Overlap | P-value | Adjusted P-value | Odds Ratio | Combined Score | Genes |
|---|---|---|---|---|---|---|---|
| 1 | BMP Signaling Pathway in Eyelid Development WP3663 | 4/20 | 1.63E-04 | 0.008084917 | 17.5258007 | 152.8156133 | FOXC1;SFRP1;NOTCH1;DKK2 |
| 2 | ACE Inhibitor Pathway WP396 | 3/9 | 2.26E-04 | 0.008084917 | 34.9450355 | 293.4093291 | AGTR2;AGT;KNG1 |
| 3 | Endochondral Ossification WP1270 | 6/62 | 2.50E-04 | 0.008084917 | 7.54953917 | 62.61477381 | TGFB2;SPP1;COL10A1;IGF1;MMP9;SOX5 |
| 4 | Calcium Regulation in the Cardiac Cell WP553 | 8/147 | 0.00124586 | 0.030212029 | 4.06742332 | 27.2026495 | GRK5;GJB4;GJB3;PKIA;RGS20;ARRB1;ADRB2;ADRA1B |
| 5 | Retinol metabolism WP1259 | 4/39 | 0.00224284 | 0.043511133 | 8.00406711 | 48.82490153 | RBP2;CRABP1;RBP1;LPL |
| 6 | Adipogenesis genes WP447 | 7/134 | 0.0031274 | 0.050559567 | 3.88364584 | 22.39913942 | FZD1;IRS1;GADD45A;LPL;AHR;IGF1;AGT |
| 7 | Wnt Signaling Pathway NetPath WP539 | 6/108 | 0.00452116 | 0.062650346 | 4.13514653 | 22.32560196 | FZD1;SFRP1;DAB2;SFRP2;ARRB1;WNT2 |
| 8 | Factors and pathways affecting insulin-like growth factor (IGF | 3/31 | 0.00958426 | 0.116209169 | 7.47986322 | 34.76365891 | IRS1;IGF1;PLD1 |
| 9 | Lung fibrosis WP3632 | 4/61 | 0.01115496 | 0.120225668 | 4.90928389 | 22.07150774 | SPP1;PTX3;IGF1;MMP9 |
| 10 | Spinal Cord Injury WP2432 | 5/99 | 0.01369127 | 0.132805328 | 3.72739362 | 15.99423411 | EFNB2;GADD45A;CXCL1;SLIT2;MMP9 |
| 11 | Focal Adhesion WP85 | 7/185 | 0.01704963 | 0.147607533 | 2.7637014 | 11.25276007 | TNN;COL5A3;COL11A1;TNC;SPP1;IGF1;THBS1 |
| 12 | Focal Adhesion-PI3K-Akt-mTOR-signaling pathway WP2841 | 10/324 | 0.01826073 | 0.147607533 | 2.24678633 | 8.993891679 | ANGPT1;IRS1;TNN;COL11A1;COL5A3;SPP1;TNC;IGF1;N |
| 13 | PluriNetWork WP1763 | 9/292 | 0.02464833 | 0.172267862 | 2.23905362 | 8.291318549 | FZD1;TLE4;DNMT3L;MYCN;NOTCH1;GADD45A;IRS1;SF |
| 14 | MAPK signaling pathway WP493 | 6/159 | 0.02661219 | 0.172267862 | 2.7495959 | 9.971095835 | DUSP4;TGFB2;GADD45A;DUSP1;ARRB1;NGF |
| 15 | Small Ligand GPCRs WP353 | 2/18 | 0.02663936 | 0.172267862 | 8.70097173 | 31.54420231 | PTGFR;PTGIR |
| 16 | MicroRNAs in Cardiomyocyte Hypertrophy WP1560 | 4/82 | 0.02971071 | 0.180121151 | 3.58372114 | 12.60125177 | FZD1;CISH;IGF1;AGT |
| 17 | Glutathione metabolism WP164 | 2/20 | 0.03247228 | 0.185283002 | 7.73341186 | 26.50525231 | ANPEP;GSTT1 |
| 18 | Id Signaling Pathway WP512 | 3/51 | 0.03610114 | 0.19359228 | 4.35882092 | 14.47752219 | IRS1;IGF1;NGF |
| 19 | TGF Beta Signaling Pathway WP113 | 3/52 | 0.03792014 | 0.19359228 | 4.26964828 | 13.97145472 | FST;SPP1;THBS1 |

NGF;THBS1
PP1;KLF4

| # | Term | Overlap | P-value | Adjusted P-value | Odds Ratio | Combined Score | Genes |
|---|---|---|---|---|---|---|---|
| 1 | Amplification and Expansion of Oncogenic Pathways as Metas | 5/17 | 3.05E-06 | 7.20E-04 | 29.3199405 | 372.366288 | POSTN;NOTCH1;TNC;CYTIP;WNT2 |
| 2 | Hair Follicle Development: Cytodifferentiation (Part 3 of 3) W | 8/87 | 3.40E-05 | 0.004012614 | 7.17854042 | 73.85998064 | FZD1;SFRP1;NOTCH1;FST;KRT15;IGF1;CD34;CTGF |
| 3 | Differentiation Pathway WP2848 | 6/48 | 5.90E-05 | 0.004638477 | 10.0732207 | 98.09895349 | IL11;NOTCH1;FST;IGF1;WNT2;GDF5 |
| 4 | ACE Inhibitor Pathway WP554 | 4/17 | 8.30E-05 | 0.004897292 | 21.5735012 | 202.7177862 | AGTR2;AGT;KNG1;NR3C2 |
| 5 | BMP Signaling Pathway in Eyelid Development WP3927 | 4/20 | 1.63E-04 | 0.007711277 | 17.5258007 | 152.8156133 | FOXC1;SFRP1;NOTCH1;DKK2 |
| 6 | ncRNAs involved in Wnt signaling in hepatocellular carcinoma | 7/86 | 2.28E-04 | 0.008965823 | 6.25862854 | 52.4874105 | FZD1;SFRP1;SFRP2;ROR1;KLF4;WNT2;DKK2 |
| 7 | Endochondral Ossification WP474 | 6/64 | 2.98E-04 | 0.010040151 | 7.28846867 | 59.17569515 | TGFB2;SPP1;COL10A1;IGF1;MMP9;SOX5 |
| 8 | Vitamin D Receptor Pathway WP2877 | 9/182 | 0.00123559 | 0.03561906 | 3.68346318 | 24.66521682 | IL1RL1;SFRP1;TGFB2;GADD45A;SPP1;ADRB2;ADRA1B; |
| 9 | Calcium Regulation in the Cardiac Cell WP536 | 8/149 | 0.00135835 | 0.03561906 | 4.00931971 | 26.46745041 | GRK5;GJB4;GJB3;PKIA;RGS20;ARRB1;ADRB2;ADRA1B |
| 10 | Lung fibrosis WP3624 | 5/63 | 0.00203147 | 0.046075018 | 6.05203202 | 37.51651928 | SPP1;PTX3;IGF1;MMP9;CTGF |
| 11 | LncRNA involvement in canonical Wnt signaling and colorecta | 6/94 | 0.00226799 | 0.046075018 | 4.79643206 | 29.2048053 | FZD1;SFRP1;SFRP2;ROR1;WNT2;DKK2 |
| 12 | Extracellular vesicles in the crosstalk of cardiac cells WP4300 | 3/19 | 0.0023428 | 0.046075018 | 13.0977394 | 79.32527373 | SPP1;IGF1;MMP9 |
| 13 | Adipogenesis WP236 | 7/130 | 0.00264038 | 0.04793312 | 4.01076212 | 23.81121702 | FZD1;IRS1;GADD45A;LPL;AHR;IGF1;AGT |
| 14 | Vitamin A and Carotenoid Metabolism WP716 | 4/43 | 0.00321882 | 0.054260039 | 7.18167716 | 41.21379044 | RBP2;CRABP1;RBP1;LPL |
| 15 | miRNA targets in ECM and membrane receptors WP2911 | 3/22 | 0.0036078 | 0.05676266 | 11.0279955 | 62.02870512 | COL5A3;ITGA1;THBS1 |
| 16 | Gastric acid production WP2596 | 2/7 | 0.00405324 | 0.05978523 | 27.8586572 | 153.4521625 | CCK;VIP |
| 17 | Hypothesized Pathways in Pathogenesis of Cardiovascular Dis | 3/25 | 0.00522142 | 0.072485604 | 9.52272727 | 50.04179619 | POSTN;LTBP2;CTGF |
| 18 | Wnt Signaling WP428 | 6/115 | 0.00612424 | 0.080295539 | 3.86820558 | 19.71044644 | FZD1;SFRP1;SFRP2;ROR1;WNT2;DKK2 |
| 19 | Oligodendrocyte Specification and differentiation(including re | 3/30 | 0.00874753 | 0.108653504 | 7.7572892 | 36.76167061 | CXCL1;IGF1;SOX5 |
| 20 | Factors and pathways affecting insulin-like growth factor (IGF | 3/31 | 0.00958426 | 0.113094284 | 7.47986322 | 34.76365891 | IRS1;IGF1;PLD1 |
| 21 | Focal Adhesion-PI3K-Akt-mTOR-signaling pathway WP3932 | 10/303 | 0.01195407 | 0.134340969 | 2.41042507 | 10.67018897 | ANGPT1;IRS1;TNN;COL11A1;COL5A3;SPP1;TNC;IGF1;N |
| 22 | BMP2-WNT4-FOXO1 Pathway in Human Primary Endometrial | 2/13 | 0.01422969 | 0.139925322 | 12.6591712 | 53.83216844 | SFRP1;DCN |
| 23 | Estrogen Receptor Pathway WP2881 | 2/13 | 0.01422969 | 0.139925322 | 12.6591712 | 53.83216844 | PDK4;CYP1B1 |
| 24 | Osteopontin Signaling WP1434 | 2/13 | 0.01422969 | 0.139925322 | 12.6591712 | 53.83216844 | SPP1;MMP9 |
| 25 | Role of Osx and miRNAs in tooth development WP3971 | 2/15 | 0.01880045 | 0.177476229 | 10.7105192 | 42.56225984 | NOTCH1;KLF4 |
| 26 | Deregulation of Rab and Rab Effector Genes in Bladder Cance | 2/16 | 0.02128661 | 0.192302433 | 9.94497728 | 38.28495278 | RAB27B;SYTL2 |
| 27 | Focal Adhesion WP306 | 7/198 | 0.02370931 | 0.192302433 | 2.57388226 | 9.631177252 | TNN;COL5A3;ITGA1;SPP1;TNC;IGF1;THBS1 |
| 28 | NOTCH1 regulation of human endothelial cell calcification WF | 2/17 | 0.02390095 | 0.192302433 | 9.28150766 | 34.65563604 | NOTCH1;ITGA1 |
| 29 | MAPK Signaling Pathway WP382 | 8/246 | 0.02516974 | 0.192302433 | 2.36349847 | 8.702668179 | DUSP4;CACNA1I;TGFB2;MAPKAPK3;GADD45A;DUSP1; |
| 30 | Epithelial to mesenchymal transition in colorectal cancer WP4 | 6/159 | 0.02661219 | 0.192302433 | 2.7495959 | 9.971095835 | FZD1;TGFB2;NOTCH1;PKP2;WNT2;MMP9 |
| 31 | Simplified Interaction Map Between LOXL4 and Oxidative Stre | 2/18 | 0.02663936 | 0.192302433 | 8.70097173 | 31.54420231 | DDR1;LOXL4 |
| 32 | Spinal Cord Injury WP2431 | 5/118 | 0.02700967 | 0.192302433 | 3.09766119 | 11.18739069 | EFNB2;GADD45A;CXCL1;SLIT2;MMP9 |
| 33 | Aryl Hydrocarbon Receptor Pathway WP2873 | 3/46 | 0.02770459 | 0.192302433 | 4.86689758 | 17.45346005 | ALDH3A1;CYP1B1;AHR |
| 34 | Aryl Hydrocarbon Receptor WP2586 | 3/46 | 0.02770459 | 0.192302433 | 4.86689758 | 17.45346005 | LPL;CYP1B1;AHR |
| 35 | Small Ligand GPCRs WP247 | 2/19 | 0.02949779 | 0.198899354 | 8.18873415 | 28.85251413 | PTGFR;PTGIR |
| 36 | MicroRNAs in cardiomyocyte hypertrophy WP1544 | 4/84 | 0.0320706 | 0.207120481 | 3.49377224 | 12.01793183 | FZD1;CISH;IGF1;AGT |
| 37 | Imatinib and Chronic Myeloid Leukemia WP3640 | 2/20 | 0.03247228 | 0.207120481 | 7.73341186 | 26.50525231 | GADD45A;ABCG2 |
| 38 | Synaptic Vesicle Pathway WP2267 | 3/51 | 0.03610114 | 0.224207081 | 4.35882092 | 14.47752219 | UNC13B;SNAP25;SLC17A7 |
| 39 | Cardiac Progenitor Differentiation WP2406 | 3/53 | 0.03978515 | 0.235529317 | 4.18404255 | 13.49044741 | NOTCH1;ANPEP;IGF1 |
| 40 | Nuclear Receptors Meta-Pathway WP2882 | 9/319 | 0.03992022 | 0.235529317 | 2.04119916 | 6.574441699 | IL11;ALDH3A1;TGFB2;SRPX2;AMIGO2;PDK4;CYP1B1;A |
| 41 | TGF-beta Receptor Signaling WP560 | 3/54 | 0.0416959 | 0.240005683 | 4.10179391 | 13.03284478 | FST;SPP1;THBS1 |
| 42 | Hematopoietic Stem Cell Differentiation WP2849 | 3/55 | 0.04365209 | 0.245283178 | 4.02270867 | 12.59712869 | THRB;NOTCH1;CD34 |
| 43 | Angiogenesis WP1539 | 2/24 | 0.04545494 | 0.249473608 | 6.32605204 | 19.55404091 | ANGPT1;MMP9 |
| 44 | Preimplantation Embryo WP3527 | 3/58 | 0.04978998 | 0.262364645 | 3.80270793 | 11.40790151 | DNMT3L;KLF4;NR3C2 |

KLF4;KNG1

NGF;THBS1

ARRB1;NGF

HR;SMARCA1

| # | Term | Overlap | P-value | Adjusted P-value | Odds Ratio | Combined Score | Genes |
|---|---|---|---|---|---|---|---|
| 1 | extracellular matrix organization (GO:0030198) | 21/300 | 2.32E-09 | 4.94E-06 | 5.54138156 | 110.1825965 | DDR1;COL28A1;POSTN;TGFB2;LUM;COL11A1;COL12A1;ITGA1;TNC;LOXL4;MMP9;THBS1;DCN;SMOC2;COL5A3; |
| 2 | extracellular structure organization (GO:0043062) | 16/216 | 8.98E-08 | 6.81E-05 | 5.80371747 | 94.16661406 | DDR1;COL28A1;POSTN;LUM;COL11A1;ITGA1;TNC;MMP9;THBS1;DCN;SMOC2;COL5A3;MMP28;SPP1;COL10A1 |
| 3 | external encapsulating structure organization (GO:0045229) | 16/217 | 9.58E-08 | 6.81E-05 | 5.77454734 | 93.32458625 | DDR1;COL28A1;POSTN;LUM;COL11A1;ITGA1;TNC;MMP9;THBS1;DCN;SMOC2;COL5A3;MMP28;SPP1;COL10A1 |
| 4 | negative regulation of cell population proliferation (GO:000828 | 21/379 | 1.33E-07 | 7.11E-05 | 4.3010094 | 68.08649209 | TGFB2;PTGIR;NOTCH1;ITGA1;TREM2;CXCL1;NGF;KLF4;GDF5;IFIT3;MTSS1;RERG;PER2;SFRP1;SFRP2;RGCC;TNN |
| 5 | positive regulation of cell differentiation (GO:0045597) | 17/258 | 1.92E-07 | 7.93E-05 | 5.12568898 | 79.26155785 | FOXC1;TGFB2;LPL;IGF1;PTN;NGF;GDF5;AGT;SFRP1;DAB2;SFRP2;RGCC;GPC1;ASB4;CD34;BRINP3;SOX5 |
| 6 | regulation of angiogenesis (GO:0045765) | 15/203 | 2.36E-07 | 7.93E-05 | 5.77039007 | 88.05811753 | SEMA5A;FOXC1;TGFB2;SPHK1;SULF1;KLF4;THBS1;DCN;SMOC2;SFRP1;SFRP2;RGCC;CYP1B1;CD34;RAPGEF3 |
| 7 | regulation of fibroblast growth factor receptor signaling pathw | 6/20 | 2.60E-07 | 7.93E-05 | 30.2626728 | 458.8247439 | SMOC2;GPC1;SULF1;FGFBP3;THBS1;SULF2 |
| 8 | positive regulation of protein phosphorylation (GO:0001934) | 20/371 | 4.13E-07 | 1.07E-04 | 4.16362952 | 61.20058588 | IL11;CEMIP;TGFB2;RAMP3;ANGPT1;SPHK1;ARRB1;TREM2;IGF1;NGF;GDF5;MMP9;CX3CL1;AGT;DAB2;SFRP2;T |
| 9 | regulation of cell migration (GO:0030334) | 21/408 | 4.51E-07 | 1.07E-04 | 3.97275076 | 58.04696809 | SEMA5A;CEMIP;NOTCH1;SERPINE2;SPHK1;CHRD;IGF1;SULF1;MMP9;THBS1;CX3CL1;SFRP1;DAB2;SFRP2;FAM1 |
| 10 | negative regulation of endothelial cell migration (GO:0010596) | 7/38 | 9.58E-07 | 1.91E-04 | 15.9883964 | 221.5681889 | NOTCH1;RGCC;GADD45A;SLIT2;AGTR2;THBS1;DCN |
| 11 | cellular response to oxygen-containing compound (GO:190170 | 18/323 | 9.87E-07 | 1.91E-04 | 4.29029287 | 59.3270105 | RAMP3;IRS1;LPL;CXCL1;AHR;MMP9;CXCL5;SFRP1;P2RX3;PDK4;PDE3A;SH3BP4;SLIT2;CPEB3;WNT2;BRINP3;RA |
| 12 | negative regulation of cellular response to growth factor stimul | 9/80 | 2.04E-06 | 3.55E-04 | 9.02204532 | 118.1913297 | NOTCH1;GPC1;CHRD;AGTR2;SLIT2;SULF1;THBS1;AGT;SULF2 |
| 13 | negative regulation of cellular process (GO:0048523) | 24/566 | 2.16E-06 | 3.55E-04 | 3.25283115 | 42.43020714 | SEMA5A;TGFB2;NOTCH1;ANGPT1;DUSP1;FHL1;ITGA1;TREM2;CXCL1;NGF;KLF4;IFIT3;KNG1;RERG;SFRP1;SFRP2 |
| 14 | positive regulation of macromolecule biosynthetic process (GO | 11/129 | 2.44E-06 | 3.71E-04 | 6.66729556 | 86.16966724 | IL33;SMOC2;TGFB2;RGCC;IRS1;IGF1;PLD1;CPEB3;GLI1;KLF4;THBS1 |
| 15 | glomerular filtration (GO:0003094) | 4/9 | 4.81E-06 | 6.00E-04 | 56.113879 | 687.1202686 | MCAM;SULF1;CD34;SULF2 |
| 16 | positive regulation of cell population proliferation (GO:000828 | 21/474 | 4.88E-06 | 6.00E-04 | 3.38234999 | 41.36950623 | PTGFR;IL11;TGFB2;NOTCH1;IRS1;SPHK1;CHRD;IGF1;PTN;GLI1;THBS1;CX3CL1;AGT;CXCL5;EFNB2;SFRP1;SFRP2; |
| 17 | collagen fibril organization (GO:0030199) | 9/89 | 5.01E-06 | 6.00E-04 | 8.00339674 | 97.67117005 | COL28A1;TGFB2;LUM;COL11A1;COL5A3;COL12A1;LOXL4;COL10A1;CYP1B1 |
| 18 | regulation of cell population proliferation (GO:0042127) | 28/764 | 5.07E-06 | 6.00E-04 | 2.80944426 | 34.25522233 | PTGFR;NOTCH1;IRS1;CXCL1;PTN;GLI1;THBS1;IFIT3;CX3CL1;RERG;CXCL5;EFNB2;GRK5;CYP1B1;HAS2;SH3BP4;W |
| 19 | regulation of neuroinflammatory response (GO:0150077) | 5/19 | 5.60E-06 | 6.29E-04 | 25.1288265 | 303.8748757 | IL33;SPHK1;TREM2;IGF1;MMP9 |
| 20 | regulation of peptidyl-tyrosine phosphorylation (GO:0050730) | 9/92 | 6.60E-06 | 7.04E-04 | 7.71293871 | 92.00370396 | IL11;SFRP1;SFRP2;TEC;ANGPT1;ENPP2;TREM2;IGF1;AGT |
| 21 | negative regulation of signal transduction (GO:0009968) | 15/267 | 7.26E-06 | 7.35E-04 | 4.29078483 | 50.77568153 | RNF43;PTGIR;IRS1;ARRB1;SULF1;THBS1;CX3CL1;AGT;DCN;SULF2;SFRP1;SFRP2;GPC1;AGTR2;DACT1 |
| 22 | negative regulation of blood vessel endothelial cell migration ( | 6/34 | 7.64E-06 | 7.35E-04 | 15.1205837 | 178.1566107 | NOTCH1;RGCC;GADD45A;AGTR2;KLF4;THBS1 |
| 23 | renal filtration (GO:0097205) | 4/10 | 7.92E-06 | 7.35E-04 | 46.7591934 | 549.2126167 | MCAM;SULF1;CD34;SULF2 |
| 24 | negative regulation of epithelial cell proliferation (GO:0050680 | 8/72 | 8.37E-06 | 7.44E-04 | 8.86777978 | 103.6673184 | SFRP1;TGFB2;SFRP2;RGCC;SULF1;THBS1;GDF5;MTSS1 |
| 25 | glycosaminoglycan biosynthetic process (GO:0006024) | 9/97 | 1.02E-05 | 8.71E-04 | 7.27285079 | 83.5849178 | CEMIP;ANGPT1;LUM;GPC1;OMD;OGN;HAS2;ST3GAL1;DCN |
| 26 | positive regulation of cellular process (GO:0048522) | 24/625 | 1.15E-05 | 9.46E-04 | 2.92447453 | 33.25186403 | PTGFR;IL11;TGFB2;RAMP3;NOTCH1;IRS1;SPHK1;CHRD;TREM2;IGF1;PTN;GLI1;THBS1;CX3CL1;CXCL5;EFNB2;SF |
| 27 | positive regulation of signal transduction (GO:0009967) | 14/252 | 1.66E-05 | 0.001312244 | 4.22769698 | 46.52776757 | TGFB2;NOTCH1;IRS1;TREM2;IGF1;GLI1;SULF1;TPD52L1;SULF2;SMOC2;SFRP1;RSPO2;FGFBP3;DACT1 |
| 28 | response to amyloid-beta (GO:1904645) | 6/44 | 3.56E-05 | 0.002655886 | 11.1358234 | 114.073751 | RAMP3;CASP4;TREM2;ADRB2;IGF1;MMP9 |
| 29 | negative regulation of fibroblast growth factor receptor signali | 4/14 | 3.61E-05 | 0.002655886 | 28.0498221 | 286.9206693 | GPC1;SULF1;THBS1;SULF2 |
| 30 | positive regulation of MAPK cascade (GO:0043410) | 14/274 | 4.18E-05 | 0.002867406 | 3.8655975 | 38.97360964 | IL11;TGFB2;RAMP3;NOTCH1;ANGPT1;DIRAS2;ARRB1;TREM2;IGF1;ADRB2;ADRA1B;TPD52L1;CX3CL1;ROR1 |
| 31 | negative regulation of cell migration (GO:0030336) | 10/144 | 4.23E-05 | 0.002867406 | 5.31370421 | 53.51911458 | SFRP1;SFRP2;TNN;DPYSL3;CHRD;CYP1B1;SLIT2;SULF1;PHLDB2;CX3CL1 |
| 32 | positive regulation of angiogenesis (GO:0045766) | 9/116 | 4.30E-05 | 0.002867406 | 5.97561967 | 60.07826041 | SEMA5A;SMOC2;SFRP2;SPHK1;CYP1B1;KLF4;CD34;THBS1;RAPGEF3 |
| 33 | regulation of endothelial cell chemotaxis (GO:2001026) | 4/15 | 4.87E-05 | 0.003147059 | 25.4985442 | 253.2021796 | SEMA5A;SMOC2;NOTCH1;THBS1 |
| 34 | regulation of smooth muscle cell proliferation (GO:0048660) | 6/49 | 6.64E-05 | 0.004165315 | 9.83845961 | 94.64487604 | PTGIR;OGN;IGF1;VIP;THBS1;CX3CL1 |
| 35 | negative regulation of cell adhesion (GO:0007162) | 7/73 | 8.09E-05 | 0.004785123 | 7.49634838 | 70.63405543 | SEMA5A;NOTCH1;ANGPT1;DUSP1;CYP1B1;CX3CL1;KNG1 |
| 36 | negative regulation of growth (GO:0045926) | 9/126 | 8.21E-05 | 0.004785123 | 5.46209588 | 51.3845526 | SFRP1;TGFB2;SFRP2;NOTCH1;FHL1;SH3BP4;AGTR2;SLIT2;RERG |
| 37 | positive regulation of stem cell differentiation (GO:2000738) | 4/17 | 8.30E-05 | 0.004785123 | 21.5735012 | 202.7177862 | FOXC1;TGFB2;PTN;SOX5 |
| 38 | regulation of extracellular matrix assembly (GO:1901201) | 3/7 | 9.61E-05 | 0.005391747 | 52.4228723 | 484.9422001 | NOTCH1;RGCC;AGT |
| 39 | regulation of epithelial to mesenchymal transition (GO:001071 | 7/76 | 1.05E-04 | 0.005662413 | 7.16932541 | 65.70536592 | SFRP1;FOXC1;TGFB2;DAB2;SFRP2;RGCC;PHLDB2 |
| 40 | positive regulation of vasculature development (GO:1904018) | 8/102 | 1.06E-04 | 0.005662413 | 6.02842 | 55.16191397 | SEMA5A;SMOC2;SFRP2;SPHK1;CYP1B1;CD34;THBS1;RAPGEF3 |
| 41 | cellular response to organonitrogen compound (GO:0071417) | 8/103 | 1.14E-04 | 0.005916635 | 5.96465894 | 54.16924319 | SFRP1;P2RX3;PDE3A;SH3BP4;AHR;SLIT2;CPEB3;RAPGEF3 |
| 42 | positive regulation of intracellular signal transduction (GO:190 | 20/546 | 1.19E-04 | 0.006022638 | 2.75328216 | 24.87395588 | SEMA5A;IL11;TGFB2;RAMP3;ANGPT1;IRS1;SPHK1;TREM2;IGF1;ADRB2;ADRA1B;THBS1;CX3CL1;AGT;DCN;NR3 |
| 43 | negative regulation of blood vessel morphogenesis (GO:20001 | 7/78 | 1.23E-04 | 0.006022638 | 6.96666329 | 62.69750272 | FOXC1;TGFB2;RGCC;KLF4;SULF1;THBS1;DCN |
| 44 | negative regulation of programmed cell death (GO:0043069) | 16/381 | 1.24E-04 | 0.006022638 | 3.15323115 | 28.35803222 | CDSN;ANGPT1;SPHK1;AMIGO2;TREM2;IGF1;NGF;MMP9;THBS1;IFIT3;CX3CL1;SFRP1;DAB2;GRK5;VIP;CRYAB |
| 45 | positive regulation of multicellular organismal process (GO:005 | 15/345 | 1.38E-04 | 0.006551693 | 3.26346801 | 29.0013111 | IL33;FOXC1;TGFB2;FST;DIO2;TREM2;PTN;ADRB2;ADRA1B;AGT;PER2;DAB2;RGCC;VIP;SOX5 |
| 46 | glycosaminoglycan catabolic process (GO:0006027) | 6/56 | 1.42E-04 | 0.006576644 | 8.45806452 | 74.94584454 | CEMIP;LUM;GPC1;OMD;OGN;DCN |
| 47 | endoderm formation (GO:0001706) | 5/36 | 1.49E-04 | 0.006758804 | 11.3387097 | 99.91723689 | DUSP4;DUSP1;COL11A1;COL12A1;MMP9 |
| 48 | renal system development (GO:0072001) | 6/57 | 1.57E-04 | 0.006960799 | 8.29179844 | 72.64896275 | FOXC1;TGFB2;HAS2;SULF1;AGT;SULF2 |
| 49 | regulation of Wnt signaling pathway (GO:0030111) | 8/111 | 1.92E-04 | 0.008336317 | 5.49914128 | 47.07593878 | RNF43;SFRP1;SFRP2;RSPO2;FOXL1;SULF1;DACT1;SULF2 |
| 50 | negative regulation of cell motility (GO:2000146) | 8/114 | 2.30E-04 | 0.009816295 | 5.34268783 | 44.75555283 | SFRP1;SFRP2;DPYSL3;CHRD;CYP1B1;SLIT2;SULF1;CX3CL1 |
| 51 | transmembrane receptor protein tyrosine kinase signaling path | 16/404 | 2.41E-04 | 0.010046202 | 2.96278695 | 24.67665084 | DDR1;FOXC1;ANGPT1;IRS1;IGF1;ADRB2;SULF1;NGF;KALRN;MMP9;MTSS1;SULF2;EFNB2;MAPKAPK3;PDK4;RO |
| 52 | negative regulation of angiogenesis (GO:0016525) | 7/87 | 2.45E-04 | 0.010046202 | 6.18008094 | 51.38491086 | FOXC1;TGFB2;RGCC;KLF4;SULF1;THBS1;DCN |
| 53 | positive regulation of cellular biosynthetic process (GO:003132 | 10/180 | 2.68E-04 | 0.01080134 | 4.18074866 | 34.37862203 | SMOC2;RGCC;IRS1;AGTR2;IGF1;PLD1;CPEB3;GLI1;KLF4;THBS1 |
| 54 | regulation of endothelial cell migration (GO:0010594) | 7/89 | 2.82E-04 | 0.011135245 | 6.02873311 | 49.27850643 | SMOC2;ANGPT1;PTN;SLIT2;THBS1;DCN;AGT |
| 55 | regulation of cell growth (GO:0001558) | 11/217 | 2.96E-04 | 0.011484987 | 3.80198072 | 30.88982388 | SFRP1;TGFB2;SFRP2;CISH;SPHK1;FHL1;SH3BP4;AGTR2;SLIT2;AGT;RERG |
| 56 | cellular response to organic cyclic compound (GO:0071407) | 9/150 | 3.08E-04 | 0.011715496 | 4.52682701 | 36.60743562 | SFRP1;RAMP3;P2RX3;PDE3A;SPP1;CYP1B1;AHR;RAPGEF3;TGM2 |

| | | | | | | | |
|---|---|---|---|---|---|---|---|
| 57 | positive regulation of epithelial to mesenchymal transition (GO | 5/42 | 3.13E-04 | 0.011728833 | 9.49710425 | 76.62205363 | FOXC1;TGFB2;DAB2;NOTCH1;RGCC |
| 58 | cellular response to lipid (GO:0071396) | 11/219 | 3.20E-04 | 0.011778571 | 3.7650372 | 30.2946757 | SFRP1;RAMP3;IRS1;PDK4;SPP1;LPL;CXCL1;AHR;BRINP3;WNT2;CXCL5 |
| 59 | regulation of cardiac muscle hypertrophy (GO:0010611) | 4/24 | 3.43E-04 | 0.01238552 | 14.0177936 | 111.847618 | NOTCH1;IGF1;LMCD1;AGT |
| 60 | regulation of epithelial cell proliferation (GO:0050678) | 7/93 | 3.69E-04 | 0.013129962 | 5.74715576 | 45.42436202 | SFRP1;TGFB2;SFRP2;ANGPT1;IGF1;GDF5;MTSS1 |
| 61 | cardiac ventricle morphogenesis (GO:0003208) | 5/44 | 3.91E-04 | 0.013348905 | 9.00915751 | 70.69392362 | TGFB2;NOTCH1;SFRP2;PKP2;CPE |
| 62 | regulation of endothelial cell apoptotic process (GO:2000351) | 5/44 | 3.91E-04 | 0.013348905 | 9.00915751 | 70.69392362 | SEMA5A;IL11;RGCC;ANGPT1;THBS1 |
| 63 | positive regulation of epithelial cell migration (GO:0010634) | 7/94 | 3.94E-04 | 0.013348905 | 5.68080708 | 44.52884099 | SMOC2;TGFB2;ANGPT1;ENPP2;MMP9;THBS1;AGT |
| 64 | cAMP-mediated signaling (GO:0019933) | 4/25 | 4.03E-04 | 0.0134413 | 13.3496018 | 104.3381494 | ADGRG6;PDE3A;AHR;RAPGEF3 |
| 65 | regulation of apoptotic process (GO:0042981) | 23/742 | 4.25E-04 | 0.013953396 | 2.31931542 | 18.00468272 | UNC13B;ANGPT1;GADD45A;SPHK1;IGF1;NGF;KALRN;MMP9;THBS1;IFIT3;CX3CL1;AGT;KNG1;FGD3;SFRP1;DAB |
| 66 | positive regulation of chemotaxis (GO:0050921) | 5/45 | 4.35E-04 | 0.014050783 | 8.78348214 | 67.99036303 | SEMA5A;SMOC2;TREM2;PTN;THBS1 |
| 67 | cellular response to growth factor stimulus (GO:0071363) | 9/158 | 4.51E-04 | 0.014226914 | 4.28202509 | 32.99320324 | SFRP1;FOXC1;NOTCH1;SPHK1;PDE3A;HAS2;KLF4;WNT2;SOX5 |
| 68 | negative regulation of cell growth (GO:0030308) | 8/126 | 4.54E-04 | 0.014226914 | 4.7964266 | 36.92479502 | SFRP1;TGFB2;SFRP2;FHL1;SH3BP4;AGTR2;SLIT2;RERG |
| 69 | proteoglycan metabolic process (GO:0006029) | 4/26 | 4.71E-04 | 0.014569993 | 12.7421546 | 97.60451443 | FOXL1;IGF1;SULF1;SULF2 |
| 70 | regulation of fibroblast proliferation (GO:0048145) | 5/46 | 4.82E-04 | 0.014570792 | 8.56881533 | 65.44199607 | SFRP1;SPHK1;IGF1;WNT2;AGT |
| 71 | kidney development (GO:0001822) | 6/70 | 4.85E-04 | 0.014570792 | 6.6031586 | 50.3909558 | FOXC1;TGFB2;HAS2;SULF1;AGT;SULF2 |
| 72 | positive regulation of cell migration (GO:0030335) | 12/269 | 5.08E-04 | 0.015037119 | 3.32800274 | 25.24573659 | SEMA5A;CEMIP;TGFB2;DAB2;NOTCH1;FAM110C;SPHK1;ENPP2;HAS2;IGF1;MMP9;THBS1 |
| 73 | positive regulation of apoptotic process (GO:0043065) | 13/310 | 5.38E-04 | 0.015268243 | 3.12480194 | 23.52311726 | UNC13B;TGFB2;GADD45A;NGF;TPD52L1;KALRN;KNG1;FGD3;SFRP1;SFRP2;CYP1B1;SLIT2;TGM2 |
| 74 | regulation of blood coagulation (GO:0030193) | 4/27 | 5.47E-04 | 0.015268243 | 12.187529 | 91.53833895 | SERPINE2;THBS1;CD34;KNG1 |
| 75 | positive regulation of secretion by cell (GO:1903532) | 6/72 | 5.64E-04 | 0.015268243 | 6.40241121 | 47.88839612 | UNC13B;TGFB2;RGCC;TREM2;RAB27B;IGF1 |
| 76 | cellular response to fluid shear stress (GO:0071498) | 3/12 | 5.73E-04 | 0.015268243 | 23.2931442 | 173.8888723 | HAS2;KLF4;MTSS1 |
| 77 | keratan sulfate catabolic process (GO:0042340) | 3/12 | 5.73E-04 | 0.015268243 | 23.2931442 | 173.8888723 | LUM;OGN;OMD |
| 78 | negative regulation of hemostasis (GO:1900047) | 3/12 | 5.73E-04 | 0.015268243 | 23.2931442 | 173.8888723 | SERPINE2;CD34;KNG1 |
| 79 | negative regulation of vascular permeability (GO:0043116) | 3/12 | 5.73E-04 | 0.015268243 | 23.2931442 | 173.8888723 | ANGPT1;PDE3A;SLIT2 |
| 80 | positive regulation of vascular permeability (GO:0043117) | 3/12 | 5.73E-04 | 0.015268243 | 23.2931442 | 173.8888723 | ANGPT1;PDE3A;FGFBP3 |
| 81 | positive regulation of gene expression (GO:0010628) | 17/482 | 5.90E-04 | 0.015532299 | 2.62598299 | 19.52594078 | IL33;RAMP3;NOTCH1;ANGPT1;TREM2;ANK3;IGF1;PLD1;NGF;KLF4;THBS1;AGT;C1QTNF1;MYCN;RGCC;CPEB3;C |
| 82 | negative regulation of canonical Wnt signaling pathway (GO:00 | 9/165 | 6.17E-04 | 0.015861919 | 4.08841973 | 30.21750984 | TLE4;SFRP1;DAB2;SFRP2;TNN;GLI1;DKK2;BICC1;DACT1 |
| 83 | keratan sulfate biosynthetic process (GO:0018146) | 4/28 | 6.31E-04 | 0.015861919 | 11.6791222 | 86.05011144 | LUM;OMD;OGN;ST3GAL1 |
| 84 | positive regulation of fibroblast proliferation (GO:0048146) | 4/28 | 6.31E-04 | 0.015861919 | 11.6791222 | 86.05011144 | SPHK1;IGF1;WNT2;AGT |
| 85 | negative regulation of apoptotic process (GO:0043066) | 17/485 | 6.32E-04 | 0.015861919 | 2.60874314 | 19.21722172 | NOTCH1;ANGPT1;SPHK1;TREM2;IGF1;NGF;KLF4;MMP9;THBS1;IFIT3;CX3CL1;SFRP1;DAB2;GRK5;PDK4;VIP;CRY |
| 86 | regulation of cell adhesion (GO:0030155) | 8/133 | 6.50E-04 | 0.016113672 | 4.52620939 | 33.21796208 | SEMA5A;ANGPT1;DUSP1;CHRD;CYTIP;ZDHHC2;KNG1;TGM2 |
| 87 | negative regulation of wound healing (GO:0061045) | 4/29 | 7.24E-04 | 0.017550497 | 11.2113879 | 81.06529558 | SERPINE2;CD34;PHLDB2;KNG1 |
| 88 | positive regulation of extrinsic apoptotic signaling pathway (GC | 4/29 | 7.24E-04 | 0.017550497 | 11.2113879 | 81.06529558 | SFRP1;AGTR2;THBS1;AGT |
| 89 | positive regulation of cellular metabolic process (GO:0031325) | 7/105 | 7.68E-04 | 0.018350808 | 5.04033916 | 36.14841275 | RAMP3;ANGPT1;GADD45A;ARRB1;KLF4;THBS1;AGT |
| 90 | regulation of inflammatory response (GO:0050727) | 10/206 | 7.74E-04 | 0.018350808 | 3.62133581 | 25.94164449 | IL33;CASP12;SPHK1;CASP4;LPL;IL16;KLF4;MMP9;CX3CL1;AGT |
| 91 | positive regulation of phosphatidylinositol 3-kinase signaling (( | 6/77 | 8.07E-04 | 0.018921379 | 5.95002272 | 42.37539553 | TGFB2;TREM2;ROR1;IGF1;DCN;AGT |
| 92 | positive regulation of small GTPase mediated signal transductic | 4/30 | 8.26E-04 | 0.019154803 | 10.7796332 | 76.52135984 | NOTCH1;IGF1;NGF;TGM2 |
| 93 | sprouting angiogenesis (GO:0002040) | 5/52 | 8.53E-04 | 0.019560338 | 7.47264438 | 52.80870386 | SEMA5A;EFNB2;ANGPT1;SLIT2;THBS1 |
| 94 | positive regulation of response to external stimulus (GO:00321 | 8/139 | 8.68E-04 | 0.019689578 | 4.3175793 | 30.43744779 | IL33;LPL;TREM2;IL16;PTN;THBS1;CX3CL1;AGT |
| 95 | cardiac left ventricle morphogenesis (GO:0003214) | 3/14 | 9.28E-04 | 0.020190983 | 19.0560928 | 133.0655786 | NOTCH1;SFRP2;CPE |
| 96 | negative regulation of cell migration involved in sprouting angi | 3/14 | 9.28E-04 | 0.020190983 | 19.0560928 | 133.0655786 | NOTCH1;KLF4;THBS1 |
| 97 | negative regulation of coagulation (GO:0050819) | 3/14 | 9.28E-04 | 0.020190983 | 19.0560928 | 133.0655786 | SERPINE2;CD34;KNG1 |
| 98 | negative regulation of neuroinflammatory response (GO:0150C | 3/14 | 9.28E-04 | 0.020190983 | 19.0560928 | 133.0655786 | TREM2;IGF1;CX3CL1 |
| 99 | regulation of potassium ion transport (GO:0043266) | 4/31 | 9.38E-04 | 0.020210452 | 10.3798603 | 72.36548233 | FHL1;KCNAB1;ANK3;VIP |
| 100 | positive regulation of developmental process (GO:0051094) | 9/177 | 0.00101657 | 0.02168352 | 3.79406056 | 26.14607511 | FST;ENPP2;FOXO6;PTN;CPEB3;PHLDB2;DCN;RAPGEF3;SOX5 |
| 101 | regulation of cell migration involved in sprouting angiogenesis | 4/32 | 0.00106014 | 0.021991912 | 10.0086426 | 68.55275919 | NOTCH1;SRPX2;KLF4;THBS1 |
| 102 | negative regulation of chemotaxis (GO:0050922) | 4/32 | 0.00106014 | 0.021991912 | 10.0086426 | 68.55275919 | SEMA5A;NOTCH1;SLIT2;THBS1 |
| 103 | positive regulation of phosphorylation (GO:0042327) | 11/253 | 0.00106196 | 0.021991912 | 3.23042468 | 22.12077324 | DDR1;DAB2;ANGPT1;ROR1;ARRB1;TREM2;IGF1;FAM20A;SLC4A4;MMP9;THBS1 |
| 104 | regulation of blood vessel endothelial cell migration (GO:0043 | 5/55 | 0.00110264 | 0.02261472 | 7.02321429 | 47.82842699 | RGCC;ANGPT1;GADD45A;AGTR2;THBS1 |
| 105 | regulation of extracellular matrix organization (GO:1903053) | 3/15 | 0.00114741 | 0.023089008 | 17.4671986 | 118.2572005 | DDR1;NOTCH1;AGT |
| 106 | regulation of microglial cell activation (GO:1903978) | 3/15 | 0.00114741 | 0.023089008 | 17.4671986 | 118.2572005 | SPHK1;TREM2;CX3CL1 |
| 107 | sulfur compound biosynthetic process (GO:0044272) | 7/113 | 0.00118354 | 0.023561218 | 4.65803584 | 31.39166255 | ANGPT1;LUM;OMD;OGN;GSTT1;ST3GAL1;DCN |
| 108 | keratan sulfate metabolic process (GO:0042339) | 4/33 | 0.00119297 | 0.023561218 | 9.66302614 | 65.04479274 | LUM;OMD;OGN;ST3GAL1 |
| 109 | cellular response to transforming growth factor beta stimulus ( | 7/114 | 0.00124591 | 0.024380974 | 4.61426746 | 30.85970889 | SFRP1;TGFB2;PDE3A;LTBP2;WNT2;GDF5;SOX5 |
| 110 | positive regulation of macromolecule metabolic process (GO:0( | 14/384 | 0.00129744 | 0.02515851 | 2.70100728 | 17.95457705 | IL33;RAMP3;NOTCH1;ANGPT1;ARRB1;TREM2;ANK3;IGF1;NGF;KLF4;C1QTNF1;MYCN;RGCC;CD34 |
| 111 | positive regulation of cell motility (GO:2000147) | 10/221 | 0.00131894 | 0.025238024 | 3.36130978 | 22.28860645 | SEMA5A;CEMIP;DAB2;NOTCH1;FAM110C;SPHK1;HAS2;IGF1;MMP9;THBS1 |
| 112 | positive regulation of calcium ion transport into cytosol (GO:00 | 4/34 | 0.00133704 | 0.025238024 | 9.34045077 | 61.80856906 | CEMIP;RAMP3;P2RX3;CX3CL1 |
| 113 | positive regulation of DNA binding (GO:0043388) | 4/34 | 0.00133704 | 0.025238024 | 9.34045077 | 61.80856906 | FOXC1;IGF1;NGF;MMP9 |

| # | Term | Overlap | P-value | Adjusted P-value | Odds Ratio | Combined Score | Genes |
|---|---|---|---|---|---|---|---|
| 1 | Systolic blood pressure (alcohol consumption interaction) | 4/10 | 7.92E-06 | 0.004604124 | 46.7591934 | 549.2126167 | PTGFR;LUM;PDE3A;SLC16A9 |
| 2 | Intraocular pressure | 18/401 | 1.97E-05 | 0.005429054 | 3.40282219 | 36.86505595 | FOXC1;TGFB2;ANGPT1;LUM;FST;LTBP2;IGF1;FAM213A;KALRN;DCN;ALDH3A1;ALCAM;ANPEP;PKIA;HIVEP3;GLCCI1;ST3GAL1;ADAMTS6 |
| 3 | Thyroid function | 3/5 | 2.80E-05 | 0.005429054 | 104.856383 | 1099.11811 | MAF;PDE8B;NR3C2 |
| 4 | Coronary artery calcified atherosclerotic plaque (130 HU thre: | 5/38 | 1.94E-04 | 0.028139679 | 10.6504329 | 91.0508812 | NOTCH1;P2RX3;PDE3A;ANK3;MTSS1 |
| 5 | Nicotine dependence and major depression (severity of como | 4/25 | 4.03E-04 | 0.041648821 | 13.3496018 | 104.3381494 | CPED1;SMOC2;SLC16A9;SULF2 |
| 6 | Spherical equivalent or myopia (age of diagnosis) | 10/191 | 4.30E-04 | 0.041648821 | 3.92446007 | 30.42034822 | MYO1D;MYCN;MAF;THRB;RASGEF1B;PDE3A;COL10A1;HIVEP3;CD34;BICC1 |
| 7 | Paclitaxel disposition in epithelial ovarian cancer | 5/47 | 5.33E-04 | 0.044267892 | 8.36437075 | 63.03671047 | ABLIM1;GLRA3;MAF;DIO2;PTN |
| 8 | Blood protein levels | 47/2091 | 0.0010906 | 0.073807604 | 1.70726578 | 11.64530351 | SEMA5A;ITIH2;SERPINE2;AMIGO2;TNC;TREM2;CXCL1;PTN;CX3CL1;KNG1;CXCL5;CTGF;WISP2;NRGN;IL1RL1;ALCAM;IL18RAP;EVA1C;GPC1;MMP28;C1RL;RSPO2 |
| 9 | Systolic blood pressure | 20/657 | 0.00124678 | 0.073807604 | 2.26035959 | 15.11546366 | FZD1;FOXC1;HSPB7;FOXO6;LPL;LTBP2;IGF1;ADRB2;SMARCA1;ADRA1B;NGF;AGT;H19;TEC;PDE3A;GPR149;HIVEP3;PHACTR1;PAM;SOX5 |
| 10 | Serum metabolite concentrations in chronic kidney disease | 4/34 | 0.00133704 | 0.073807604 | 9.34045077 | 61.80856906 | TMEM176A;ASPG;SLIT2;SLC16A9 |
| 11 | Cotinine glucuronidation | 3/16 | 0.00139739 | 0.073807604 | 16.1227496 | 105.9772353 | KCNH5;KLF4;FAM107B |
| 12 | Response to platinum-based agents | 2/5 | 0.00196689 | 0.095230426 | 46.4358068 | 289.3554382 | FOXC1;ITGA1 |
| 13 | Central corneal thickness | 5/66 | 0.00249524 | 0.111518137 | 5.75351288 | 34.48292827 | TGFB2;COL12A1;ADAMTS6;DCN;NR3C2 |
| 14 | Fasting blood insulin (BMI interaction) | 3/21 | 0.00314871 | 0.130671539 | 11.641253 | 67.0624861 | IRS1;IGF1;SULF1 |
| 15 | Osteoarthritis (hip) | 3/23 | 0.00410566 | 0.159025937 | 10.4760638 | 57.57004088 | IL11;COL11A1;TNC |
| 16 | Pelvic organ prolapse (moderate/severe) | 4/47 | 0.00445022 | 0.161598654 | 6.51228999 | 35.26275761 | RTL1;PGBD5;CPE;SOX5 |
| 17 | Serum protein levels (sST2) | 2/8 | 0.00535363 | 0.182968219 | 23.2143698 | 121.4106951 | IL1RL1;IL18RAP |
| 18 | Serum thyroid-stimulating hormone levels | 3/26 | 0.005841 | 0.18853453 | 9.10823312 | 46.84230496 | MAF;ITGA1;PDE8B |
| 19 | Anxiety disorder | 2/9 | 0.00681876 | 0.192850407 | 19.8970217 | 99.24788618 | CXCL1;MFAP3L |
| 20 | Paneth cell defects in Crohn's disease | 2/9 | 0.00681876 | 0.192850407 | 19.8970217 | 99.24788618 | EYA1;ACTR3B |
| 21 | Thyroid hormone levels | 3/28 | 0.00720732 | 0.192850407 | 8.3787234 | 41.32937649 | MAF;PDE8B;NR3C2 |
| 22 | Bone mineral density (hip) | 4/54 | 0.00730243 | 0.192850407 | 5.59857651 | 27.5424703 | TGFB2;GADD45A;FOXL1;FJX1 |
| 23 | Local histogram emphysema pattern | 2/10 | 0.0084437 | 0.197997598 | 17.4090106 | 83.11644343 | MYO1D;TGFB2 |
| 24 | Adiponectin levels | 3/30 | 0.00874753 | 0.197997598 | 7.7572892 | 36.76167061 | IRS1;PDE3A;KNG1 |
| 25 | Uric acid levels | 3/31 | 0.00958426 | 0.197997598 | 7.47986322 | 34.76365891 | SEMA5A;SLC16A9;ABCG2 |
| 26 | Age-related nuclear cataracts | 2/11 | 0.01022363 | 0.197997598 | 15.4738909 | 70.91767359 | KCNAB1;SLC4A4 |
| 27 | Response to antidepressants in depression | 2/11 | 0.01022363 | 0.197997598 | 15.4738909 | 70.91767359 | RASGEF1B;RHOU |
| 28 | Fasting blood insulin | 2/11 | 0.01022363 | 0.197997598 | 15.4738909 | 70.91767359 | IRS1;IGF1 |
| 29 | LDL peak particle diameter (total fat intake interaction) | 2/11 | 0.01022363 | 0.197997598 | 15.4738909 | 70.91767359 | ABCG2;SOX5 |
| 30 | Low high density lipoprotein cholesterol levels | 2/11 | 0.01022363 | 0.197997598 | 15.4738909 | 70.91767359 | P2RX3;LMCD1 |
| 31 | Age-related macular degeneration | 4/67 | 0.01533865 | 0.281454191 | 4.44037734 | 18.54914214 | DDR1;POSTN;COL10A1;CD34 |
| 32 | Self-reported allergy | 3/37 | 0.01556462 | 0.281454191 | 6.15801001 | 25.63428796 | IL1RL1;IL33;DAB2 |
| 33 | Obstetric antiphospholipid syndrome | 2/14 | 0.0164467 | 0.281454191 | 11.6036514 | 47.66350821 | SH3BP4;NGF |
| 34 | Myopia | 4/70 | 0.01775513 | 0.281454191 | 4.23789496 | 17.0832961 | MYO1D;THRB;COL10A1;BICC1 |
| 35 | Heel bone mineral density | 21/898 | 0.01833668 | 0.281454191 | 1.70863999 | 6.832597918 | FOXC1;TGFB2;THRB;EYA1;IRS1;COL11A1;ITGA1;ANK3;FAM213A;KLF4;SLC4A4;MMP9;DCN;BICC1;CPED1;DAB2;RGCC;RSPO2;HIVEP3;ADAMTS6;SOX5 |
| 36 | Midgestational circulating levels of PCBs | 2/15 | 0.01880045 | 0.281454191 | 10.7105192 | 42.56225984 | FST;BICC1 |
| 37 | Cholangiocarcinoma in primary sclerosing cholangitis (time to | 2/16 | 0.02128661 | 0.281454191 | 9.94497728 | 38.28495278 | CISH;MAPKAPK3 |
| 38 | Systolic blood pressure response to hydrochlorothiazide in hy | 2/16 | 0.02128661 | 0.281454191 | 9.94497728 | 38.28495278 | PKIA;ZDHHC2 |
| 39 | Mitral valve prolapse | 2/16 | 0.02128661 | 0.281454191 | 9.94497728 | 38.28495278 | LMCD1;CTGF |
| 40 | Prostate cancer (SNP x SNP interaction) | 4/74 | 0.0213271 | 0.281454191 | 3.99491612 | 15.37154557 | ABLIM1;PKIA;AMIGO2;TNC |
| 41 | Urate levels | 4/74 | 0.0213271 | 0.281454191 | 3.99491612 | 15.37154557 | DAB2;MAF;SLC16A9;ABCG2 |
| 42 | Migraine | 4/74 | 0.0213271 | 0.281454191 | 3.99491612 | 15.37154557 | DOCK4;GPR149;PHACTR1;NGF |
| 43 | Glomerular filtration rate | 5/111 | 0.0214129 | 0.281454191 | 3.30340296 | 12.69749408 | H19;CEMIP;KNG1;SOX5;ABCG2 |
| 44 | Systemic juvenile idiopathic arthritis | 3/42 | 0.02183532 | 0.281454191 | 5.36715767 | 20.52522532 | COL11A1;COL12A1;FOXL1 |
| 45 | Primary biliary cholangitis | 5/112 | 0.02216147 | 0.281454191 | 3.27236315 | 12.46574038 | IL1RL1;IL18RAP;GLRA3;IL16;PAM |
| 46 | Dental caries | 4/75 | 0.02228381 | 0.281454191 | 3.9384492 | 14.98144697 | FZD1;SPP1;RHOU;ABCG2 |
| 47 | Glaucoma (primary open-angle) | 4/76 | 0.02326633 | 0.287611412 | 3.88355081 | 14.60505645 | FOXC1;ANGPT1;COL11A1;BICC1 |
| 48 | Fractional exhaled nitric oxide (childhood) | 2/17 | 0.02390095 | 0.289301124 | 9.28150766 | 34.65563604 | ANGPT1;SLC4A4 |
| 49 | Extremely high intelligence | 4/78 | 0.02530935 | 0.300096596 | 3.77820525 | 13.89087888 | THRB;COL11A1;FOXO6;APBA1 |
| 50 | Coffee consumption (cups per day) | 2/18 | 0.02663936 | 0.30347977 | 8.70097173 | 31.54420231 | AHR;ABCG2 |
| 51 | Migraine without aura | 2/18 | 0.02663936 | 0.30347977 | 8.70097173 | 31.54420231 | PHACTR1;NGF |
| 52 | Metabolic traits | 3/46 | 0.02770459 | 0.309545491 | 4.86689758 | 17.45346005 | LPL;AHR;SLC16A9 |
| 53 | HIV-1 control | 2/19 | 0.02949779 | 0.323362518 | 8.18873415 | 28.85251413 | DDR1;CDSN |
| 54 | Cardiovascular disease risk factors | 2/20 | 0.03247228 | 0.331754545 | 7.73341186 | 26.50525231 | LPL;ABCG2 |
| 55 | Takayasu arteritis | 4/88 | 0.0371116 | 0.331754545 | 3.32672428 | 10.95764977 | DDR1;CDSN;KCNK10;KLF4 |
| 56 | Asthma (childhood onset) | 3/52 | 0.03792014 | 0.331754545 | 4.26964828 | 13.97145472 | TLE4;IL1RL1;IL33 |
| 57 | Glaucoma | 3/52 | 0.03792014 | 0.331754545 | 4.26964828 | 13.97145472 | FOXC1;ANGPT1;BICC1 |
| 58 | Asthma and hay fever | 2/22 | 0.0387541 | 0.331754545 | 6.95936396 | 22.62154281 | IL33;IL1RL1 |
| 59 | Cutaneous malignant melanoma | 2/22 | 0.0387541 | 0.331754545 | 6.95936396 | 22.62154281 | CYP1B1;KLF4 |
| 60 | Pulmonary function (smoking interaction) | 3/54 | 0.0416959 | 0.331754545 | 4.10179391 | 13.03284478 | MYCN;LUM;ITGA1 |
| 61 | Multiple system atrophy | 2/23 | 0.04205396 | 0.331754545 | 6.62762914 | 21.00164306 | LOXL4;FOXL1 |
| 62 | Cognitive function | 4/92 | 0.04258282 | 0.331754545 | 3.1748625 | 10.02083237 | RNF43;LRRC8D;GLCCI1;MTSS1 |
| 63 | Serum uric acid levels | 3/55 | 0.04365209 | 0.331754545 | 4.02270867 | 12.59712869 | RGS20;BICC1;ABCG2 |
| 64 | HDL cholesterol levels | 4/93 | 0.04401794 | 0.331754545 | 3.13902995 | 9.80368671 | IRS1;TMEM176A;PDE3A;LPL |
| 65 | Diastolic blood pressure | 15/646 | 0.04448148 | 0.331754545 | 1.68022539 | 5.230008116 | TGFB2;DUSP1;ANK3;ADRA1B;NGF;SLC7A2;AGT;RERG;H19;TNN;PDE3A;GPR149;HIVEP3;PHACTR1;PAM |
| 66 | Oppositional defiant disorder dimensions in attention-deficit I | 2/24 | 0.04545494 | 0.331754545 | 6.32605204 | 19.55404091 | FOXS1;SOX5 |
| 67 | Dupuytren's disease | 3/56 | 0.0456534 | 0.331754545 | 3.94660779 | 12.18190461 | RSPO2;SULF1;WNT2 |
| 68 | Gut microbiota (functional units) | 3/56 | 0.0456534 | 0.331754545 | 3.94660779 | 12.18190461 | SEMA5A;HIVEP3;SORCS2 |
| 69 | Amyotrophic lateral sclerosis (sporadic) | 6/182 | 0.04650441 | 0.331754545 | 2.38746334 | 7.325234383 | ALDH3A1;DOCK4;ALCAM;THRB;ANK3;NR3C2 |

CDSN;SCARA5;ANGPT1;LUM;IL16;BPGM;FAM213A;GDF5;MMP9;DKK2;AGT;ASPN;DCN;ALDH3A1;SFRP1;MAPKAPK3;PKP2;SPARCL1;ROR1;FAM20A;PAM

| # | Term | Overlap | P-value | Adjusted P-value | Odds Ratio | Combined Score | Genes |
|---|---|---|---|---|---|---|---|
| 1 | Extracellular space | 60/1348 | 1.61E-15 | 1.71E-12 | 3.81511387 | 129.9473289 | DDR1;SERPINE2;COL12A1;TNC;LOXL4;CXCL1;MSLN;CX3CL1;CXCL5;CTGF;WISP2;C1QTNF1;ANPEP;DPYSL3;C1RL;ENPP2;LMCD1;IL11;POSTN;OMD;IL16;SEPP1;MMP9;DKK2;DCN;ALDH3A1;SFRP1;SFRP2;SPARCL1;FJX1;ITIH2;CHRD;LPL;LTBP2;PTN |
| 2 | Extracellular matrix | 35/489 | 3.84E-15 | 2.03E-12 | 5.93951542 | 197.1489545 | SERPINE2;COL11A1;COL12A1;TNC;LPL;LTBP2;PTN;THBS1;CTGF;WISP2;IL1RL1;TNN;GPC1;MMP28;COL10A1;SLIT2;FGFBP3;WNT2;ADAMTS6;TGM2;CTHRC1;COL28A1;POSTN;TGFB2;LUM;OMD;MMP9;ASPN;DCN;SMOC2;SFRP1;SFRP2;COL5A3;C |
| 3 | Thrombospondin complex | 28/317 | 1.43E-14 | 5.04E-12 | 7.32336111 | 233.466153 | SEMA5A;NOTCH1;SERPINE2;TNC;CXCL1;PTN;THBS1;KNG1;CTGF;WISP2;EFNB2;MMP28;RSPO2;SPP1;COL10A1;CD34;ADAMTS6;IL33;POSTN;TGFB2;LUM;IGF1;MMP9;ASPN;DCN;CPED1;OGN;SPARCL1 |
| 4 | Proteinaceous extracellular matrix | 28/347 | 1.37E-13 | 3.63E-11 | 6.62439774 | 196.1962006 | COL11A1;COL12A1;TNC;LTBP2;PTN;CTGF;WISP2;IL1RL1;TNN;GPC1;MMP28;COL10A1;SLIT2;WNT2;ADAMTS6;CTHRC1;COL28A1;POSTN;LUM;OMD;MMP9;ASPN;DCN;SMOC2;SFRP1;COL5A3;OGN;SPARCL1 |
| 5 | Extracellular region | 117/4434 | 4.87E-13 | 1.03E-10 | 2.4840415 | 70.42372033 | SEMA5A;SERPINE2;COL12A1;TNC;LOXL4;SLC4A4;HID1;CTGF;ALCAM;TNN;ANPEP;DPYSL3;CAPN5;C1RL;ENPP2;TGM2;POSTN;FST;OMD;ACOT11;SEPP1;DKK2;ALDH3A1;SFRP1;SFRP2;SCG5;SPARCL1;AGTR2;PTGFR;LXN;NOTCH1;CHRD;LPL;LTBP2 |
| 6 | Extracellular region part | 103/3766 | 3.42E-12 | 6.03E-10 | 2.48003648 | 65.47458726 | SEMA5A;SERPINE2;COL12A1;TNC;LOXL4;SLC4A4;HID1;CTGF;ALCAM;TNN;ANPEP;DPYSL3;CAPN5;C1RL;ENPP2;TGM2;POSTN;OMD;ACOT11;SEPP1;DKK2;ALDH3A1;SFRP1;SFRP2;SPARCL1;LXN;CHRD;LPL;LTBP2;GSTT1;ACTR3B;KALRN;KNG1;S10 |
| 7 | NF-kappaB complex | 38/864 | 6.70E-10 | 9.72E-08 | 3.51815981 | 74.31728759 | NOTCH1;IRS1;TREM2;CXCL1;GLI1;THBS1;IFIT3;CX3CL1;KNG1;CXCL5;NR3C2;CTGF;IL1RL1;IL18RAP;PDLIM2;SPP1;NLRP2;CYP1B1;HAS2;HIVEP3;CD34;TGM2;IL11;IL33;CISH;ANGPT1;GADD45A;DUSP1;CCK;IGF1;FOXL1;NGF;MMP9;AGT;MAF;MYC |
| 8 | VEGF-A complex | 32/639 | 7.35E-10 | 9.72E-08 | 3.98158506 | 83.73651666 | PTGFR;FOXC1;MEGF6;NOTCH1;TNC;CXCL1;PTN;THBS1;CX3CL1;CXCL5;CTGF;EFNB2;ALCAM;SPP1;COL10A1;PHACTR1;PDE8B;CD34;IL11;POSTN;TGFB2;ANGPT1;MCAM;IGF1;SULF1;NGF;MMP9;AGT;DCN;DAB2;MYCN;FIBIN |
| 9 | Type III intermediate filament | 39/946 | 2.44E-09 | 2.87E-07 | 3.28749294 | 65.19650012 | SNAP25;NOTCH1;TNC;CXCL1;PTN;GLI1;MSLN;THBS1;KNG1;CXCL5;ALCAM;ANPEP;DPYSL3;SPP1;HAS2;SLC17A7;CD34;NAPSA;POSTN;TGFB2;LUM;MCAM;ITGA1;CCK;IGF1;NGF;KLF4;MMP9;AGT;DCN;MYCN;P2RX3;KRT15;RBP1;OGN;PKP2;VIP;C |
| 10 | platelet-derived growth factor complex | 18/268 | 6.25E-08 | 6.61E-06 | 5.24898876 | 87.07011821 | TGFB2;NOTCH1;ANGPT1;IRS1;TNC;CXCL1;IGF1;NGF;GDF5;MMP9;THBS1;AGT;KNG1;DCN;CTGF;SPP1;HAS2;CD34 |
| 11 | Cell surface | 30/715 | 1.36E-07 | 1.31E-05 | 3.26835552 | 51.67392322 | RAMP3;NOTCH1;SERPINE2;LPL;PTN;MSLN;THBS1;CX3CL1;IL1RL1;ALCAM;SRPX2;TNN;ANPEP;CAPN5;RSPO2;SLIT2;CD34;FZD1;KCNH5;SCARA5;MCAM;HEG1;ITGA1;ANK3;SULF1;SULF2;SFRP1;PAM;VAMP5;CRYAB |
| 12 | Integrin alphav-beta3 complex | 12/142 | 9.34E-07 | 8.23E-05 | 6.62214708 | 91.94369056 | THRB;ANGPT1;IRS1;MCAM;OMD;TNC;SPP1;IGF1;CD34;MMP9;THBS1;CTGF |
| 13 | Elastic fiber | 13/171 | 1.10E-06 | 8.95E-05 | 5.91588328 | 81.16888913 | TGFB2;LUM;TNC;LOXL4;LTBP2;IGF1;MMP9;DCN;CTGF;H19;SMOC2;SPP1;CD34 |
| 14 | Spanning component of plasma membrane | 69/2798 | 2.46E-06 | 1.86E-04 | 1.98830463 | 25.68170813 | DDR1;SEMA5A;SNAP25;SERPINE2;IRS1;TNC;ARRB1;TREM2;CXCL1;GRPR;GLI1;ADRA1B;SLC4A4;CX3CL1;ALCAM;CASP12;EVA1C;ANPEP;ENPP2;AOX1;CD34;IL11;RNF43;PTGIR;KCNH5;KCNK10;SCARA5;CISH;HEG1;FST;NGF;MMP9;ASPN;SLC27A6 |
| 15 | Collagen trimer | 9/84 | 3.09E-06 | 2.18E-04 | 8.53913043 | 108.3428497 | COL28A1;C1QTNF1;LUM;COL11A1;COL5A3;COL12A1;COL10A1;DCN;CTHRC1 |
| 16 | Activin complex | 17/329 | 5.46E-06 | 3.61E-04 | 3.94483113 | 47.80071632 | IL11;TGFB2;NOTCH1;EYA1;FST;TNC;CHRD;CXCL1;IGF1;NGF;KLF4;GDF5;MMP9;CTGF;MYCN;HAS2;CD34 |
| 17 | Secretory vesicle | 20/451 | 8.12E-06 | 5.06E-04 | 3.37678939 | 39.57817699 | SNAP25;NAPSA;TGFB2;NOTCH1;SERPINE2;ITGA1;RAB27B;IGF1;SEPP1;THBS1;STX11;KNG1;SYTL2;DPYSL3;DMXL2;SCG5;CPE;SLC17A7;APBA1;PAM |
| 18 | Bcl-2 family protein complex | 37/1233 | 1.48E-05 | 8.67E-04 | 2.31012987 | 25.69849819 | NOTCH1;IRS1;IL20RA;CXCL1;GLI1;MSLN;THBS1;NR3C2;CTGF;CASP12;ANPEP;CASP4;SPP1;CYP1B1;CD34;TGM2;IL11;IL33;TGFB2;CISH;ANGPT1;GADD45A;DUSP1;MCAM;SPHK1;IGF1;NGF;KLF4;MMP9;AGT;SFRP1;MAF;MYCN;RGCC;TXNIP;AGTR2 |
| 19 | Activin AB complex | 10/133 | 2.13E-05 | 0.001186393 | 5.79216556 | 62.30362866 | PER2;IL11;TGFB2;FST;OMD;FHL1;DIO2;HAS2;SLIT2;CTGF |
| 20 | membrane-bounded vesicle | 78/3516 | 2.40E-05 | 0.001221606 | 1.78399137 | 18.9762615 | DDR1;SEMA5A;SNAP25;THRB;SERPINE2;COL12A1;LOXL4;ARRB1;SLC4A4;HID1;WISP2;NRGN;ALCAM;ANPEP;DPYSL3;CAPN5;C1RL;AOX1;TGM2;OMD;ACOT11;ITGA1;BPGM;NGF;SEPP1;MMP9;MTSS1;SYTL2;GCHFR;SFRP1;SCG5;SPARCL1;FAM20 |
| 21 | BMP receptor complex | 11/164 | 2.42E-05 | 0.001221606 | 5.13291351 | 54.54850852 | SMOC2;POSTN;TGFB2;ANGPT1;FST;CHRD;SPP1;COL10A1;HAS2;PHLDB2;GDF5 |
| 22 | BCL-2 complex | 33/1079 | 3.06E-05 | 0.00147195 | 2.33723709 | 24.29385096 | NOTCH1;IRS1;CXCL1;GLI1;MSLN;THBS1;NR3C2;CTGF;CASP12;ANPEP;CASP4;SPP1;CYP1B1;CD34;TGM2;IL11;TGFB2;CISH;ANGPT1;GADD45A;DUSP1;MCAM;SPHK1;IGF1;NGF;KLF4;MMP9;SFRP1;MAF;MYCN;TXNIP;AGTR2;ABCG2 |
| 23 | DRM complex | 10/140 | 3.32E-05 | 0.001526957 | 5.47832168 | 56.49858729 | SFRP1;DNMT3L;SFRP2;FST;AMIGO2;SCG5;CHRD;APBA1;GDF5;CRYAB |
| 24 | BAX complex | 24/689 | 5.51E-05 | 0.002430376 | 2.63417164 | 25.83013643 | TGFB2;NOTCH1;ANGPT1;GADD45A;DUSP1;SPHK1;IGF1;GLI1;NGF;KLF4;MMP9;AGT;CTGF;SFRP1;CASP12;MYCN;CASP4;SPP1;CYP1B1;TXNIP;AGTR2;CD34;TGM2;ABCG2 |
| 25 | Apical complex | 37/1315 | 5.83E-05 | 0.002467669 | 2.1523329 | 20.98468875 | NOTCH1;FHL1;TNC;CXCL1;PTN;ADRB2;GLI1;SLC4A4;CX3CL1;KNG1;CXCL5;NR3C2;ANPEP;GPC1;RSPO2;SPP1;ENPP2;SLC16A9;LMCD1;TGM2;FZD1;TGFB2;MCAM;RAB27B;CCK;IGF1;NGF;MMP9;AGT;SYTL2;DAB2;MAF;RBP1;AGTR2;VIP;RAPGEF3; |
| 26 | Extracellular matrix component | 9/122 | 6.39E-05 | 0.002600216 | 5.65659869 | 54.63256009 | COL28A1;SMOC2;LUM;COL5A3;COL11A1;COL12A1;TNC;PTN;DCN |
| 27 | Integrin alpha5-beta1 complex | 10/154 | 7.46E-05 | 0.00292187 | 4.94217172 | 46.96956022 | IL11;ANPEP;ITGA1;TNC;SPP1;IGF1;CD34;MMP9;RGAG4;CTGF |
| 28 | Activin A complex | 14/294 | 8.91E-05 | 0.003367593 | 3.58579336 | 33.43927337 | IL11;TGFB2;EYA1;FST;CHRD;CXCL1;IGF1;NGF;KLF4;GDF5;MMP9;CTGF;MYCN;CD34 |
| 29 | Protein complex involved in cell-cell adhesion | 15/334 | 9.67E-05 | 0.003411066 | 3.3779171 | 31.22434663 | POSTN;LUM;MCAM;TNC;PTN;MSLN;MMP9;THBS1;ASPN;CTGF;ALCAM;ANPEP;CAPN5;SPP1;ST3GAL1 |
| 30 | Galectin complex | 15/334 | 9.67E-05 | 0.003411066 | 3.3779171 | 31.22434663 | POSTN;LUM;MCAM;TNC;PTN;MSLN;MMP9;THBS1;ASPN;CTGF;ALCAM;ANPEP;CAPN5;SPP1;ST3GAL1 |
| 31 | Complex of collagen trimers | 4/20 | 1.63E-04 | 0.005575814 | 17.5258007 | 152.8156133 | LUM;COL11A1;COL5A3;DCN |
| 32 | Rad51C-XRCC3 complex | 7/83 | 1.83E-04 | 0.006038371 | 6.50667361 | 56.00958171 | IL1RL1;IL33;HAS2;IL16;CX3CL1;CXCL5;ADAMTS6 |
| 33 | Fibrillar collagen trimer | 3/9 | 2.26E-04 | 0.007236039 | 34.9450355 | 293.4093291 | LUM;COL5A3;COL11A1 |
| 34 | Plasma membrane part | 57/2517 | 2.42E-04 | 0.007352118 | 1.75355591 | 14.5986665 | DDR1;SNAP25;SERPINE2;IRS1;ARRB1;GRPR;ADRA1B;SLC4A4;ZDHHC2;ALCAM;C1QTNF1;ANPEP;ENPP2;CD34;TGM2;RNF43;PTGIR;KCNH5;KCNK10;SCARA5;HEG1;ITGA1;KCNAB1;ANK3;SYTL2;SDPR;ROR1;AGTR2;VAMP5;CRYAB;PTGFR;RAMP3; |
| 35 | Mucus layer | 17/446 | 2.43E-04 | 0.007352118 | 2.85166823 | 23.73031222 | UNC13B;IL11;IL33;POSTN;NOTCH1;CXCL1;CCK;IGF1;ADRB2;KLF4;SLC4A4;MMP9;KNG1;CPED1;ANPEP;VIP;TGM2 |
| 36 | RNA polymerase II transcription repressor complex | 9/151 | 3.23E-04 | 0.00949464 | 4.49471831 | 36.12694821 | DNMT3L;SFRP2;FST;AMIGO2;SCG5;CHRD;APBA1;GDF5;CRYAB |
| 37 | Activin receptor complex | 7/92 | 3.46E-04 | 0.009884654 | 5.81506559 | 46.34601215 | PER2;TGFB2;FST;OMD;SLIT2;MSLN;GDF5 |
| 38 | Synapse | 23/737 | 3.87E-04 | 0.010294213 | 2.33617187 | 18.35521038 | UNC13B;SNAP25;POSTN;SERPINE2;ARRB1;CCK;ANK3;PTN;KALRN;STX11;NRGN;GLRA3;SRPX2;P2RX3;DMXL2;CPE;RGS20;SLC17A7;APBA1;PHACTR1;CPEB3;CRYAB;DACT1 |
| 39 | PCNA complex | 24/785 | 3.89E-04 | 0.010294213 | 2.29027142 | 17.98190465 | IL11;TGFB2;NOTCH1;ANGPT1;GADD45A;DUSP1;IRS1;MCAM;TNC;IGF1;GLI1;NGF;MMP9;THBS1;CTGF;EFNB2;MYCN;RGCC;PPP1R3C;KRT15;SPP1;CYP1B1;CD34;ABCG2 |
| 40 | DNA polymerase processivity factor complex | 24/785 | 3.89E-04 | 0.010294213 | 2.29027142 | 17.98190465 | IL11;TGFB2;NOTCH1;ANGPT1;GADD45A;DUSP1;IRS1;MCAM;TNC;IGF1;GLI1;NGF;MMP9;THBS1;CTGF;EFNB2;MYCN;RGCC;PPP1R3C;KRT15;SPP1;CYP1B1;CD34;ABCG2 |
| 41 | Axon collateral | 8/124 | 4.07E-04 | 0.010443868 | 4.87962156 | 38.08903307 | FZD1;PPP1R3C;SLC17A7;CCK;PTN;SLIT2;NGF;VIP |
| 42 | Granular vesicle | 6/68 | 4.15E-04 | 0.010443868 | 6.81685744 | 53.09109171 | UNC13B;SCG5;CCK;NGF;VIP;PAM |
| 43 | Peptidase inhibitor complex | 7/96 | 4.48E-04 | 0.011027158 | 5.55258265 | 42.81224269 | ALDH3A1;IL1RL1;SFRP1;ANPEP;MTSS1;KNG1;AGT |
| 44 | cytoplasmic, membrane-bounded vesicle | 31/1147 | 4.80E-04 | 0.011544381 | 2.03401137 | 15.54290286 | SNAP25;NOTCH1;SERPINE2;ARRB1;PLD1;THBS1;STX11;KNG1;NRGN;DPYSL3;SH3BP4;RGS20;SLC17A7;IL33;NAPSA;TGFB2;ITGA1;RAB27B;IGF1;NGF;SEPP1;MTSS1;SYTL2;GCHFR;DAB2;DMXL2;SCG5;CPE;APBA1;PAM;VAMP5 |
| 45 | Glycocalyx | 10/196 | 5.27E-04 | 0.012383968 | 3.81798481 | 28.82131317 | LXN;ITIH2;ANGPT1;LUM;GPC1;TREM2;ST3GAL1;MMP9;KNG1;ADAMTS6 |
| 46 | Apical dendrite | 3/12 | 5.73E-04 | 0.013170918 | 23.2931442 | 173.8888723 | MYO1D;CPEB3;OSBP2 |
| 47 | Growth cone filopodium | 5/49 | 6.48E-04 | 0.01458331 | 7.98336039 | 58.61275858 | UNC13B;ABLIM1;DPYSL3;IGF1;NGF |
| 48 | Exocytic vesicle | 8/135 | 7.17E-04 | 0.015321346 | 4.45447568 | 32.25371051 | SNAP25;DPYSL3;DMXL2;SLC17A7;APBA1;IGF1;STX11;SYTL2 |
| 49 | Lamina reticularis | 4/29 | 7.24E-04 | 0.015321346 | 11.2113879 | 81.06529558 | POSTN;TGFB2;TNC;IGF1 |
| 50 | Integrin alpha9-beta1 complex | 4/29 | 7.24E-04 | 0.015321346 | 11.2113879 | 81.06529558 | TNC;SPP1;THBS1;TGM2 |
| 51 | Extracellular vesicle | 59/2754 | 7.74E-04 | 0.01588294 | 1.64871033 | 11.8120552 | DDR1;SEMA5A;THRB;SERPINE2;COL12A1;LOXL4;SLC4A4;HID1;WISP2;ALCAM;ANPEP;CAPN5;C1RL;AOX1;TGM2;OMD;ACOT11;ITGA1;BPGM;SEPP1;MMP9;SFRP1;SPARCL1;FAM20A;VAMP5;CRYAB;RAPGEF3;LXN;ITIH2;LPL;LTBP2;GSTT1;ACTR3 |
| 52 | Extracellular organelle | 59/2755 | 7.81E-04 | 0.01588294 | 1.64800196 | 11.79211588 | DDR1;SEMA5A;THRB;SERPINE2;COL12A1;LOXL4;SLC4A4;HID1;WISP2;ALCAM;ANPEP;CAPN5;C1RL;AOX1;TGM2;OMD;ACOT11;ITGA1;BPGM;SEPP1;MMP9;SFRP1;SPARCL1;FAM20A;VAMP5;CRYAB;RAPGEF3;LXN;ITIH2;LPL;LTBP2;GSTT1;ACTR3 |
| 53 | Cytoplasmic vesicle | 32/1245 | 9.07E-04 | 0.017658833 | 1.92924478 | 13.51529356 | SNAP25;NOTCH1;SERPINE2;ARRB1;PLD1;THBS1;STX11;KNG1;NRGN;DPYSL3;SH3BP4;RGS20;SLC17A7;IL33;NAPSA;TGFB2;ITGA1;RAB27B;IGF1;NGF;SEPP1;MTSS1;SYTL2;GCHFR;MYO1D;DAB2;DMXL2;SCG5;CPE;APBA1;PAM;VAMP5 |
| 54 | G-protein coupled receptor complex | 20/641 | 9.27E-04 | 0.017658833 | 2.32054204 | 16.20658354 | FZD1;PTGIR;RAMP3;KCNK10;SPHK1;GPR85;ARRB1;CCK;ADRB2;GRPR;GDPD5;PLD1;ADRA1B;KNG1;GRK5;ENPP2;RGS20;AGTR2;VIP;RAPGEF3 |
| 55 | Intracellular vesicle | 32/1247 | 9.31E-04 | 0.017658833 | 1.92586086 | 13.44066482 | SNAP25;NOTCH1;SERPINE2;ARRB1;PLD1;THBS1;STX11;KNG1;NRGN;DPYSL3;SH3BP4;RGS20;SLC17A7;IL33;NAPSA;TGFB2;ITGA1;RAB27B;IGF1;NGF;SEPP1;MTSS1;SYTL2;GCHFR;MYO1D;DAB2;DMXL2;SCG5;CPE;APBA1;PAM;VAMP5 |
| 56 | Neuron projection | 26/936 | 9.35E-04 | 0.017658833 | 2.07446222 | 14.47000266 | SNAP25;ARRB1;GDPD5;NRGN;ALCAM;GLRA3;DPYSL3;KIF13B;MYH14;SLC17A7;UNC13B;TGFB2;ITGA1;KCNAB1;CCK;ANK3;GCHFR;MYO1D;P2RX3;GPR149;APBA1;CPEB3;PAM;CRYAB;CAMK1G;OSBP2 |
| 57 | Neuromuscular junction | 5/54 | 0.00101405 | 0.018822184 | 7.16690962 | 49.40726631 | UNC13B;POSTN;SERPINE2;ANK3;PTN |
| 58 | CGRP receptor complex | 5/55 | 0.00110264 | 0.019772763 | 7.02321429 | 47.82842699 | RAMP3;P2RX3;VIP;PAM;KNG1 |
| 59 | Main axon | 5/55 | 0.00110264 | 0.019772763 | 7.02321429 | 47.82842699 | MYO1D;KIF13B;KCNAB1;CCK;ANK3 |
| 60 | Extracellular exosome | 58/2740 | 0.00115674 | 0.020397164 | 1.62268607 | 10.97284739 | DDR1;SEMA5A;THRB;COL12A1;LOXL4;SLC4A4;HID1;WISP2;ALCAM;ANPEP;CAPN5;C1RL;AOX1;TGM2;OMD;ACOT11;ITGA1;BPGM;SEPP1;MMP9;SFRP1;SPARCL1;FAM20A;VAMP5;CRYAB;RAPGEF3;LXN;ITIH2;LPL;LTBP2;GSTT1;ACTR3B;KALRN;T |
| 61 | Survivin complex | 15/429 | 0.00133501 | 0.023154779 | 2.59004294 | 17.14301728 | IL11;NOTCH1;ANGPT1;GADD45A;IGF1;GLI1;MSLN;MMP9;CTGF;SFRP1;MYCN;SPP1;CD34;WNT2;ABCG2 |
| 62 | Secretory granule | 13/344 | 0.00139302 | 0.023771296 | 2.79891594 | 18.40644854 | NAPSA;TGFB2;NOTCH1;SERPINE2;ITGA1;RAB27B;IGF1;SEPP1;THBS1;KNG1;SCG5;CPE;PAM |
| 63 | Calcitonin family receptor complex | 5/59 | 0.00151523 | 0.025446213 | 6.50165344 | 42.20996438 | RAMP3;P2RX3;VIP;PAM;KNG1 |
| 64 | Neuron part | 32/1287 | 0.00155282 | 0.025670049 | 1.86044754 | 12.03278473 | SNAP25;DOCK4;ARRB1;GDPD5;KALRN;STX11;NRGN;ALCAM;GLRA3;DPYSL3;KIF13B;MYH14;RGS20;SLC17A7;UNC13B;TGFB2;ITGA1;KCNAB1;CCK;ANK3;GCHFR;MYO1D;P2RX3;DMXL2;GPR149;CPE;APBA1;CPEB3;PAM;CRYAB;CAMK1G;OSBP2 |
| 65 | Transforming growth factor beta receptor homodimeric comp | 8/157 | 0.00189161 | 0.030328004 | 3.79250357 | 23.7802325 | IL11;TGFB2;TNC;GDF5;MMP9;THBS1;ASPN;CTGF |
| 66 | Growth hormone receptor complex | 5/62 | 0.00189192 | 0.030328004 | 6.1585213 | 38.61494256 | PER2;IL11;OMD;SLIT2;IGF1 |
| 67 | Axon | 14/403 | 0.00203198 | 0.032087109 | 2.56655821 | 15.9094371 | UNC13B;TGFB2;KCNAB1;CCK;ANK3;GDPD5;NRGN;MYO1D;ALCAM;P2RX3;KIF13B;MYH14;SLC17A7;CRYAB |
| 68 | nitric-oxide synthase complex | 10/236 | 0.00214178 | 0.033323638 | 3.13580048 | 19.27299414 | SPHK1;HSPB7;SPP1;ANXA10;AGTR2;NGF;VIP;MMP9;SLC7A2;KNG1 |
| 69 | Golgi lumen | 6/94 | 0.00226799 | 0.033980375 | 4.79643206 | 29.2048053 | LUM;GPC1;OMD;OGN;NGF;DCN |
| 70 | Synapse part | 18/594 | 0.00226938 | 0.033980375 | 2.2400515 | 13.63799471 | UNC13B;SNAP25;ARRB1;CCK;ANK3;KALRN;STX11;NRGN;GLRA3;SRPX2;P2RX3;DMXL2;CPE;RGS20;SLC17A7;APBA1;CPEB3;CRYAB |
| 71 | Protein complex involved in cell-matrix adhesion | 9/199 | 0.00228035 | 0.033980375 | 3.35097254 | 20.38539971 | MEGF6;POSTN;SRPX2;MMP28;TNC;TREM2;THBS1;MTSS1;CTHRC1 |
| 72 | 6-phosphofructo-2-kinase/fructose-2,6-biphosphatase compl | 4/40 | 0.00246448 | 0.036139127 | 7.7813365 | 46.73296259 | ANGPT1;LUM;FST;AOX1 |
| 73 | Glutamatergic synapse | 10/241 | 0.00249353 | 0.036139127 | 3.06713892 | 18.38460034 | UNC13B;FZD1;SEMA5A;EFNB2;SNAP25;SPARCL1;SLC17A7;CCK;CPEB3;RAPGEF3 |
| 74 | Region of cytosol | 3/20 | 0.0027274 | 0.038994496 | 12.3266583 | 72.78158675 | SLC17A7;PAM;TGM2 |
| 75 | Fatty acid synthase complex | 18/607 | 0.00285768 | 0.040312314 | 2.18912268 | 12.8233242 | IL11;TGFB2;GADD45A;DUSP1;IRS1;LPL;IL16;CXCL1;IGF1;NGF;MMP9;TRIM7;CASP12;CASP4;PDK4;SPP1;SLC27A6;CD34 |
| 76 | ATP-sensitive potassium channel complex | 6/99 | 0.00294279 | 0.040966751 | 4.53740317 | 26.44578591 | PTGIR;CCK;GLI1;KNG1;RAPGEF3;DACT1 |
| 77 | Transport vesicle | 12/333 | 0.00310235 | 0.042627027 | 2.65571189 | 15.33832154 | IL33;SNAP25;DPYSL3;DMXL2;CPE;RAB27B;SLC17A7;APBA1;IGF1;STX11;SYTL2;NRGN |
| 78 | EFF-1 complex | 3/21 | 0.00314871 | 0.04270945 | 11.641253 | 67.0624861 | NOTCH1;FXYD1;VIP |
| 79 | Integrin alphav-beta6 complex | 4/43 | 0.00321882 | 0.043107686 | 7.18167716 | 41.21379044 | LUM;TNC;SPP1;MMP9 |
| 80 | Ryanodine receptor complex | 8/172 | 0.00332575 | 0.043983033 | 3.4429867 | 19.64588979 | CASP12;CISH;FXYD1;CCK;KNG1;RERG;RAPGEF3;SSFA2 |
| 81 | External side of plasma membrane | 9/212 | 0.0034775 | 0.045422137 | 3.13429 | 17.74460149 | IL1RL1;ALCAM;SERPINE2;ANPEP;MCAM;HEG1;ITGA1;CD34;THBS1 |
| 82 | EMILIN complex | 7/137 | 0.00353558 | 0.045476937 | 3.79344217 | 21.41352063 | POSTN;MEGF6;THRB;SRPX2;TREM2;MTSS1;CTHRC1 |
| 83 | Proximal dendrite | 8/174 | 0.00356766 | 0.045476937 | 3.40115697 | 19.1683934 | CACNA1I;CDSN;SPARCL1;SLC17A7;CCK;ANK3;CPEB3;SLC4A4 |
| 84 | Platelet alpha granule | 5/72 | 0.00364615 | 0.04592415 | 5.23667377 | 29.39912106 | TGFB2;SERPINE2;IGF1;THBS1;KNG1 |
| 85 | Perikaryon | 6/104 | 0.00375752 | 0.046770091 | 4.30480579 | 24.03801707 | GLRA3;ITGA1;KCNAB1;CCK;PAM;CRYAB |
| 86 | Transforming growth factor beta type II receptor homodimer | 6/105 | 0.00393871 | 0.048106649 | 4.26110568 | 23.59332482 | TGFB2;TNC;GDF5;THBS1;ASPN;CTGF |
| 87 | C-fiber | 8/177 | 0.00395584 | 0.048106649 | 3.34026873 | 18.48024506 | PTGIR;KCNK10;P2RX3;CCK;GRPR;NGF;VIP;KNG1 |
| 88 | Receptor complex | 11/302 | 0.00420278 | 0.050528837 | 2.67971004 | 14.6633998 | DDR1;RAMP3;NOTCH1;P2RX3;IRS1;ITGA1;LOXL4;ROR1;AHR;ADRB2;NR3C2 |
| 89 | Somatodendritic compartment | 18/642 | 0.00510278 | 0.060660029 | 2.06255402 | 10.8860973 | TGFB2;ITGA1;ARRB1;KCNAB1;CCK;ANK3;GDPD5;NRGN;GCHFR;MYO1D;ALCAM;GLRA3;P2RX3;CPE;CPEB3;PAM;CRYAB;OSBP2 |

| | | | | | | | |
|---|---|---|---|---|---|---|---|
| 125 | Glucose transporter complex | 12/395 | 0.01150846 | 0.097407632 | 2.21868993 | 9.905724091 | C1QTNF1;IRS1;TXNIP;SLC27A6;LPL;IGF1;CD34;VAMP5;SLC7A2;AGT;KNG1;ABCG2 |
| 126 | Ooplasm | 6/132 | 0.01168378 | 0.098106646 | 3.34340331 | 14.87665315 | CRABP1;LPL;IL16;IGF1;KLF4;WNT2 |
| 127 | Cell projection | 37/1774 | 0.01224309 | 0.101993587 | 1.54415752 | 6.798607253 | SNAP25;DOCK4;ARRB1;GLI1;GDPD5;ACTR3B;NRGN;FGD3;ABLIM1;ALCAM;GLRA3;DPYSL3;KIF13B;SPP1;MYH14;SLC17A7;UNC13B;TGFB2;ANGPT1;ITGA1;KCNAB1;CCK;ANK3;MTSS1;GCHFR;MYO1D;ARHGAP31;P2RX3;GPR149;RHOU;APBA1;CPI |
| 128 | Activin B complex | 4/64 | 0.0131408 | 0.107837408 | 4.66310795 | 20.20074063 | TGFB2;FST;CHRD;HAS2 |
| 129 | Integrin alpha2-beta1 complex | 5/98 | 0.01314842 | 0.107837408 | 3.76766513 | 16.31946774 | DDR1;IL11;PTGIR;ITGA1;CD34 |
| 130 | Inhibin complex | 7/176 | 0.01329636 | 0.10810981 | 2.9122217 | 12.58156845 | TGFB2;FST;MCAM;IGF1;MSLN;TPD52L1;CD34 |
| 131 | Lobed nucleus | 3/35 | 0.013386 | 0.10810981 | 6.54355053 | 28.22590686 | RGCC;ANPEP;KLF4 |
| 132 | Weibel-Palade body | 5/99 | 0.01369127 | 0.109737611 | 3.72739362 | 15.99423411 | IL11;ANGPT1;MCAM;CD34;RAPGEF3 |
| 133 | Dendrite | 13/457 | 0.01445428 | 0.114279913 | 2.07441541 | 8.788810588 | ARRB1;KCNAB1;CCK;ANK3;NRGN;GCHFR;MYO1D;ALCAM;GLRA3;P2RX3;CPEB3;CRYAB;OSBP2 |
| 134 | Phospholamban complex | 7/179 | 0.01447402 | 0.114279913 | 2.86098795 | 12.11742917 | FXYD1;PKP2;PDE3A;ADRB2;IGF1;RAPGEF3;AGT |
| 135 | Dendritic branch | 6/141 | 0.01574758 | 0.123414378 | 3.11907607 | 12.94749839 | SNAP25;DOCK4;SLC17A7;PTN;NGF;NRGN |
| 136 | Integral component of plasma membrane | 33/1570 | 0.01595168 | 0.124094688 | 1.54876537 | 6.409087117 | DDR1;PTGFR;RAMP3;IRS1;ADRB2;GRPR;ADRA1B;SLC4A4;ZDHHC2;SLC7A2;EFNB2;ALCAM;C1QTNF1;GLRA3;ANPEP;GPC1;ENPP2;HAS2;LRRC8D;CD34;RNF43;PTGIR;KCNH5;KCNK10;SCARA5;ITGA1;KCNAB1;P2RX3;FXYD1;GPR149;ROR1;AGTR2;\ |
| 137 | Intrinsic component of plasma membrane | 34/1632 | 0.01636984 | 0.126418181 | 1.53572942 | 6.315402618 | DDR1;PTGFR;RAMP3;IRS1;ADRB2;GRPR;ADRA1B;SLC4A4;ZDHHC2;SLC7A2;EFNB2;ALCAM;C1QTNF1;GLRA3;ANPEP;GPC1;ENPP2;HAS2;LRRC8D;CD34;TGM2;RNF43;PTGIR;KCNH5;KCNK10;SCARA5;ITGA1;KCNAB1;P2RX3;FXYD1;GPR149;ROR1;A |
| 138 | Cytoplasmic membrane-bounded vesicle lumen | 5/104 | 0.0166323 | 0.127073412 | 3.53823954 | 14.49407551 | TGFB2;IGF1;SEPP1;THBS1;KNG1 |
| 139 | Subapical complex | 8/228 | 0.01683639 | 0.127073412 | 2.5592386 | 10.45247432 | MYO1D;ARHGAP31;ANPEP;RAB27B;RHOU;SLC4A4;BICC1;SYTL2 |
| 140 | Aryl hydrocarbon receptor complex | 8/228 | 0.01683639 | 0.127073412 | 2.5592386 | 10.45247432 | PER2;ALDH3A1;CDSN;RSPO2;SPP1;CYP1B1;GSTT1;ABCG2 |
| 141 | Integrin alpha4-beta1 complex | 7/185 | 0.01704963 | 0.127073412 | 2.7637014 | 11.25276007 | ALCAM;ANPEP;ITGA1;TNC;SPP1;CD34;MMP9 |
| 142 | Golgi apparatus part | 21/892 | 0.01715926 | 0.127073412 | 1.72095815 | 6.996068953 | SNAP25;POSTN;NOTCH1;LUM;OMD;RAB27B;ARRB1;SULF1;PLD1;NGF;DCN;SULF2;HID1;NRGN;GPC1;OGN;RGS20;RHOU;ST3GAL1;PAM;CAMK1G |
| 143 | Vesicle lumen | 5/105 | 0.01726698 | 0.127073412 | 3.50267857 | 14.21722985 | TGFB2;IGF1;SEPP1;THBS1;KNG1 |
| 144 | laminin-1 complex | 6/144 | 0.01729544 | 0.127073412 | 3.05080256 | 12.37805973 | ITGA1;TNC;NGF;MMP9;THBS1;DCN |
| 145 | Dinoflagellate epicone | 9/276 | 0.01787292 | 0.130410678 | 2.37518319 | 9.558850231 | DNMT3L;MYCN;MAPKAPK3;CPE;FAR1;TREM2;IGF1;KLF4;CD34 |
| 146 | Lytic vacuole | 14/522 | 0.01808084 | 0.131024192 | 1.95323532 | 7.838142489 | NAPSA;RAMP3;LUM;OMD;ARRB1;ANK3;ADRB2;PLD1;DCN;DAB2;ANPEP;GPC1;OGN;CD34 |
| 147 | Lysosome | 14/523 | 0.01834832 | 0.132057963 | 1.94929643 | 7.793710825 | NAPSA;RAMP3;LUM;OMD;ARRB1;ANK3;ADRB2;PLD1;DCN;DAB2;ANPEP;GPC1;OGN;CD34 |
| 148 | Female germ cell nucleus | 10/326 | 0.01896888 | 0.133229866 | 2.23233602 | 8.851113572 | PAQR8;FGD3;DNMT3L;DUSP1;FST;PDE3A;GPR149;HAS2;IGF1;RAPGEF3 |
| 149 | Germinal vesicle | 10/326 | 0.01896888 | 0.133229866 | 2.23233602 | 8.851113572 | PAQR8;FGD3;DNMT3L;DUSP1;FST;PDE3A;GPR149;HAS2;IGF1;RAPGEF3 |
| 150 | Large latent transforming growth factor-beta complex | 3/40 | 0.01918511 | 0.133229866 | 5.65784934 | 22.36899106 | TGFB2;CHRD;TGM2 |
| 151 | Cytoneme | 3/40 | 0.01918511 | 0.133229866 | 5.65784934 | 22.36899106 | NOTCH1;GPC1;GLI1 |
| 152 | Transforming growth factor beta type I receptor homodimeri | 5/108 | 0.0192667 | 0.133229866 | 3.4001387 | 13.42842962 | TGFB2;MMP9;ASPN;CTGF;DACT1 |
| 153 | Microspike | 5/108 | 0.0192667 | 0.133229866 | 3.4001387 | 13.42842962 | FGD3;NGF;THBS1;KNG1;MTSS1 |
| 154 | Membrane region | 10/329 | 0.02006868 | 0.137874421 | 2.21100028 | 8.641904686 | MYO1D;DAB2;ANGPT1;P2RX3;SDPR;IRS1;ITGA1;SH3BP4;ARRB1;SULF1 |
| 155 | L-type voltage-gated calcium channel complex | 5/110 | 0.020681 | 0.141164497 | 3.33503401 | 12.93506255 | DUSP4;CDSN;ANK3;NGF;SOX5 |
| 156 | Basal part of cell | 3/44 | 0.02467522 | 0.167348629 | 5.10482615 | 18.89784002 | ITGA1;CD34;PHLDB2 |
| 157 | GO:0098791 | 9/293 | 0.02512496 | 0.169313436 | 2.23105481 | 8.218968161 | SNAP25;POSTN;RGS20;RAB27B;SULF1;ST3GAL1;PAM;SULF2;HID1 |
| 158 | Inward rectifier potassium channel complex | 5/117 | 0.0261586 | 0.172910694 | 3.12547832 | 11.38792159 | CACNA1I;ABLIM1;PKP2;KCNAB1;APBA1 |
| 159 | Stress fiber | 3/45 | 0.02616626 | 0.172910694 | 4.98302938 | 18.15459445 | ABLIM1;PDLIM2;MYH14 |
| 160 | Contractile actin filament bundle | 3/45 | 0.02616626 | 0.172910694 | 4.98302938 | 18.15459445 | ABLIM1;PDLIM2;MYH14 |
| 161 | piccolo-bassoon transport vesicle | 2/18 | 0.02663936 | 0.172910694 | 8.70097173 | 31.54420231 | UNC13B;SLC17A7 |
| 162 | Integrin alphav-beta1 complex | 2/18 | 0.02663936 | 0.172910694 | 8.70097173 | 31.54420231 | TNC;SPP1 |
| 163 | Hyaluranon cable | 2/18 | 0.02663936 | 0.172910694 | 8.70097173 | 31.54420231 | TGFB2;HAS2 |
| 164 | Golgi stack | 5/118 | 0.02700967 | 0.174245286 | 3.09766119 | 11.18739069 | RAB27B;SULF1;ST3GAL1;HID1;SULF2 |
| 165 | Side of membrane | 11/397 | 0.02766435 | 0.176575023 | 2.01031542 | 7.21222884 | IL1RL1;ALCAM;TEC;SERPINE2;ANPEP;MCAM;HEG1;ITGA1;KCNAB1;CD34;THBS1 |
| 166 | SNARE complex | 3/46 | 0.02770459 | 0.176575023 | 4.86689758 | 17.45346005 | SNAP25;VAMP5;STX11 |
| 167 | Kir2 inward rectifier potassium channel complex | 5/120 | 0.02876426 | 0.182231058 | 3.04347826 | 10.80015284 | CACNA1I;ABLIM1;PKP2;KCNAB1;APBA1 |
| 168 | Intercalated disc | 3/47 | 0.02929012 | 0.18445799 | 4.75604449 | 16.79123966 | PKP2;ANK3;VAMP5 |
| 169 | Axon part | 7/208 | 0.02991755 | 0.186949851 | 2.44457568 | 8.578773805 | UNC13B;MYO1D;P2RX3;KIF13B;KCNAB1;CCK;ANK3 |
| 170 | voltage-gated sodium channel complex | 5/122 | 0.03058943 | 0.186949851 | 2.99114774 | 10.43043401 | P2RX3;TNC;PKP2;ANK3;NGF |
| 171 | Synaptic vesicle | 5/122 | 0.03058943 | 0.186949851 | 2.99114774 | 10.43043401 | SNAP25;DMXL2;SLC17A7;APBA1;STX11 |
| 172 | Annuli extracellular matrix | 3/48 | 0.03092271 | 0.186949851 | 4.6501182 | 16.16504088 | IL1RL1;LUM;ADAMTS6 |
| 173 | Collagen and cuticulin-based cuticle extracellular matrix | 3/48 | 0.03092271 | 0.186949851 | 4.6501182 | 16.16504088 | IL1RL1;LUM;ADAMTS6 |
| 174 | Collagen and cuticulin-based cuticle extracellular matrix part | 3/48 | 0.03092271 | 0.186949851 | 4.6501182 | 16.16504088 | IL1RL1;LUM;ADAMTS6 |
| 175 | Cortical layer of collagen and cuticulin-based cuticle extracell | 3/48 | 0.03092271 | 0.186949851 | 4.6501182 | 16.16504088 | IL1RL1;LUM;ADAMTS6 |
| 176 | Membrane microdomain | 8/257 | 0.03148881 | 0.188221267 | 2.25781103 | 7.807788097 | MYO1D;ANGPT1;P2RX3;SDPR;IRS1;GPC1;ITGA1;SULF1 |
| 177 | Membrane raft | 8/257 | 0.03148881 | 0.188221267 | 2.25781103 | 7.807788097 | MYO1D;ANGPT1;P2RX3;SDPR;IRS1;GPC1;ITGA1;SULF1 |
| 178 | Connexon complex | 2/20 | 0.03247228 | 0.193009387 | 7.73341186 | 26.50525231 | GJB4;GJB3 |
| 179 | Basal cortex | 2/21 | 0.03555898 | 0.208725258 | 7.32601823 | 24.44371887 | NOTCH1;PHLDB2 |
| 180 | Organelle subcompartment | 9/313 | 0.03607943 | 0.208725258 | 2.08212958 | 6.916901712 | SNAP25;POSTN;RGS20;RAB27B;SULF1;ST3GAL1;PAM;SULF2;HID1 |
| 181 | G-protein coupled receptor dimeric complex | 3/51 | 0.03610114 | 0.208725258 | 4.35882092 | 14.47752219 | RAMP3;RGS20;ADRB2 |
| 182 | Hippocampal mossy fiber | 3/51 | 0.03610114 | 0.208725258 | 4.35882092 | 14.47752219 | UNC13B;KCNAB1;SLC17A7 |
| 183 | Dendritic tree | 6/171 | 0.03610276 | 0.208725258 | 2.54806126 | 8.463094751 | CACNA1I;CCK;PTN;NGF;VIP;KALRN |
| 184 | Caspase complex | 10/365 | 0.03712939 | 0.213493997 | 1.98309859 | 6.531030642 | CASP12;NOTCH1;GADD45A;PLSCR2;CASP4;IGF1;NGF;CD34;MMP9;ABCG2 |
| 185 | Actin filament bundle | 3/52 | 0.03792014 | 0.216862191 | 4.26964828 | 13.97145472 | ABLIM1;MYH14;CRYAB |
| 186 | Postsynapse | 10/367 | 0.03830368 | 0.217780905 | 1.97178508 | 6.432375556 | GLRA3;P2RX3;RGS20;SLC17A7;ARRB1;ANK3;CPEB3;KALRN;CRYAB;NRGN |
| 187 | oncostatin-M receptor complex | 2/22 | 0.0387541 | 0.217780905 | 6.95936396 | 22.62154281 | CISH;TGM2 |
| 188 | Pyroptosome complex | 2/22 | 0.0387541 | 0.217780905 | 6.95936396 | 22.62154281 | CASP4;ARRB1 |
| 189 | Cell periphery | 24/1148 | 0.03910999 | 0.217780905 | 1.52092281 | 4.929884658 | FZD1;MCAM;SPHK1;AMIGO2;FHL1;ATP10D;RAB27B;IL16;PLD1;ARHGAP36;CX3CL1;SYTL2;SSFA2;ALDH3A1;DAB2;GRK5;S100A16;CASP4;KIF13B;TXNIP;LMCD1;CRYAB;TGM2;ABCG2 |
| 190 | Plasma membrane | 24/1148 | 0.03910999 | 0.217780905 | 1.52092281 | 4.929884658 | FZD1;MCAM;SPHK1;AMIGO2;FHL1;ATP10D;RAB27B;IL16;PLD1;ARHGAP36;CX3CL1;SYTL2;SSFA2;ALDH3A1;DAB2;GRK5;S100A16;CASP4;KIF13B;TXNIP;LMCD1;CRYAB;TGM2;ABCG2 |
| 191 | PSII associated light-harvesting complex II | 3/53 | 0.03978515 | 0.220380576 | 4.18404255 | 13.49044741 | CDSN;SDPR;TGM2 |
| 192 | Synaptic membrane | 8/270 | 0.04026333 | 0.221867731 | 2.14434922 | 6.88832327 | SNAP25;SRPX2;GLRA3;CPE;ARRB1;ANK3;CPEB3;CRYAB |
| 193 | Cell cortex | 7/223 | 0.04114624 | 0.222950049 | 2.27306488 | 7.252492362 | PDLIM2;FAM110C;COL10A1;RGS20;CYTIP;PHLDB2;CTGF |
| 194 | Basement membrane | 4/91 | 0.04117463 | 0.222950049 | 3.2115188 | 10.24453003 | COL28A1;SMOC2;TNC;PTN |
| 195 | Postsynaptic density | 6/177 | 0.04156984 | 0.222950049 | 2.45790102 | 7.817060227 | RGS20;ARRB1;CPEB3;KALRN;CRYAB;NRGN |
| 196 | Postsynaptic specialization | 6/177 | 0.04156984 | 0.222950049 | 2.45790102 | 7.817060227 | RGS20;ARRB1;CPEB3;KALRN;CRYAB;NRGN |
| 197 | Nuclear aryl hydrocarbon receptor complex | 2/23 | 0.04205396 | 0.222950049 | 6.62762914 | 21.00164306 | MYCN;CYP1B1 |
| 198 | Columella | 2/23 | 0.04205396 | 0.222950049 | 6.62762914 | 21.00164306 | CD34;DCN |
| 199 | Desmosome | 2/23 | 0.04205396 | 0.222950049 | 6.62762914 | 21.00164306 | CDSN;PKP2 |
| 200 | insulin-like growth factor binding protein complex | 1/3 | 0.04214557 | 0.222950049 | 34.7059859 | 109.9008695 | IGF1 |
| 201 | PET complex | 4/92 | 0.04258282 | 0.223032797 | 3.1748625 | 10.02083237 | IL33;CPE;PTN;KNG1 |
| 202 | Perineuronal net | 4/92 | 0.04258282 | 0.223032797 | 3.1748625 | 10.02083237 | FGD3;PKP2;HAS2;SLC17A7 |
| 203 | Ski complex | 7/225 | 0.04282491 | 0.22319583 | 2.25198007 | 7.095168044 | NOTCH1;EYA1;SPHK1;PHACTR1;GLI1;BICC1;LHX8 |
| 204 | Presynapse | 8/274 | 0.04326102 | 0.224363503 | 2.11166906 | 6.63170383 | UNC13B;SNAP25;P2RX3;DMXL2;SLC17A7;APBA1;CCK;STX11 |
| 205 | Actomyosin | 3/55 | 0.04365209 | 0.225287377 | 4.02270867 | 12.59712869 | ABLIM1;PDLIM2;MYH14 |
| 206 | Glycoprotein Ib-IX-V complex | 4/94 | 0.04547996 | 0.233581528 | 3.10399367 | 9.592841388 | IL11;ANPEP;CD34;KNG1 |
| 207 | Cytoplasmic vesicle part | 14/594 | 0.04616529 | 0.235955941 | 1.7043517 | 5.24177967 | TGFB2;RAB27B;ARRB1;IGF1;SEPP1;THBS1;KNG1;NRGN;DAB2;DMXL2;CPE;SLC17A7;PAM;VAMP5 |
| 208 | Dentate gyrus mossy fiber | 4/95 | 0.04696886 | 0.238908899 | 3.06972743 | 9.388056858 | UNC13B;KCNAB1;SLC17A7;CCK |
| 209 | Thylakoid light-harvesting complex | 3/57 | 0.04769948 | 0.240314512 | 3.87332545 | 11.78588959 | CDSN;SDPR;TGM2 |
| 210 | Grb2-Sos complex | 3/57 | 0.04769948 | 0.240314512 | 3.87332545 | 11.78588959 | DAB2;IRS1;IGF1 |
| 211 | Tubulin complex | 17/766 | 0.04817282 | 0.241549043 | 1.60623119 | 4.871635329 | SNAP25;KCNH5;NOTCH1;SERPINE2;IGF1;GLI1;NGF;MMP9;ASPN;CTGF;ALCAM;FAM110C;DPYSL3;SPP1;CD34;TGM2;ABCG2 |
| 212 | Transforming growth factor beta1-type II receptor complex | 2/25 | 0.04895351 | 0.24202249 | 6.05069903 | 18.25425842 | POSTN;ITGA1 |
| 213 | Smooth muscle contractile fiber | 2/25 | 0.04895351 | 0.24202249 | 6.05069903 | 18.25425842 | VIP;CD34 |
| 214 | Glutamate synthase complex | 2/25 | 0.04895351 | 0.24202249 | 6.05069903 | 18.25425842 | SNAP25;HEG1 |
