## Supplemental Table 3 for "Mechanistic insights into transcriptional regulation of ARHGAP36 expression identify a factor predictive of neuroblastoma survival"

**Table S3. List of predicted Fox and Fos-Jun transcription factor binding sites in Prox 3 and Dist 2 regions of mouse Arhgap36**

(JASPAR CORE 2024 predicted TF binding motifs; cutoff score 350)

### *Prox 3:*

| Region | Conservation | TF motifs | Peak | Foxc1 | Foxc2 | Fos-Jun |
|---|---|---|---|---|---|---|
| tf1 | partial | Fox | | 1 | 1 | |
| tf2 | conserved | Fos-Jun, Fox | | | | 1 |
| tf3 | conserved | Fos-Jun | | | | 1 |
| tf4 | conserved | Fox | | | | |
| tf5 | conserved | Fox | | | 1 | |
| tf6 | none | Fox | | 1 | 1 | |
| tf7 | partial | Fos-Jun, Fox | | 1 | | 1 |
| tf8 | conserved | Fox | | 1 | 2 | |
| tf9 | conserved | Fos-Jun, Fox | y | | | 1 |
| tf10 | conserved | Fox | y | 3 | 3 | |
| tf11 | none | Fox | | 1 | | |
| tf12 | none | Fox | | | | |
| tf13 | none | Fos-Jun, Fox | | 1 | 1 | 1 |
| tf14 | partial | Fox | | 1 | 1 | |
| tf15 | conserved | Fox | | 2 | 2 | |
| tf16 | partial | Fos-Jun, Fox | | 1 | 1 | 1 |
| tf17 | none | Fos-Jun, Fox | | | | 1 |
| tf18 | conserved | Fox | | 1 | | |
| tf19 | none | Fox | | | | |
| | | | | **14** | **13** | **7** |

**JASPAR Foxc1 motifs:**

| | |
|---|---|
| conserved | 4 |
| partial | 4 |
| non-conserved | 6 |
| total | 14 |

### *Dist 2:*

| Region | Conservation | TF motifs | Peak | Foxc1 | Foxc2 | Fos-Jun |
|---|---|---|---|---|---|---|
| tf20 | conserved | Fox | | 1 | | |
| tf21 | none | Fox | | | 1 | |

**JASPAR Foxc1 motifs:**

| | |
|---|---|
| conserved | 6 |
| partial | 2 |

| | | | | | | |
|---|---|---|---|---|---|---|
| tf22 | conserved | Fox | y | 1 | 1 | |
| tf23 | conserved | Fox | y | 1 | 1 | |
| tf24 | conserved | Fos-Jun, Fox | y | 1 | | 1 |
| tf25 | conserved | Fox | y | 2 | 2 | |
| tf26 | partial | Fos-Jun, Fox | y | 1 | 1 | 1 |
| tf27 | none | Fox | | | | |
| tf28 | none | Fox | | 1 | 1 | |
| tf29 | none | Fox | | 1 | | |
| tf30 | none | Fox | | 1 | 1 | |
| tf31 | partial | Fos-Jun, Fox | | 1 | 1 | 2 |
| tf32 | partial | Fox | | | | |
| | | | | **11** | **9** | **4** |

| | |
|---|---|
| non-conserved | 3 |
| total | 11 |
