## Supplemental Table 4 for "Mechanistic insights into transcriptional regulation of ARHGAP36 expression identify a factor predictive of neuroblastoma survival"

### CCLE dataset (DepMap portal):

- Neuroblastoma
- Glioma
- some breast
- some lung
- sporadic other

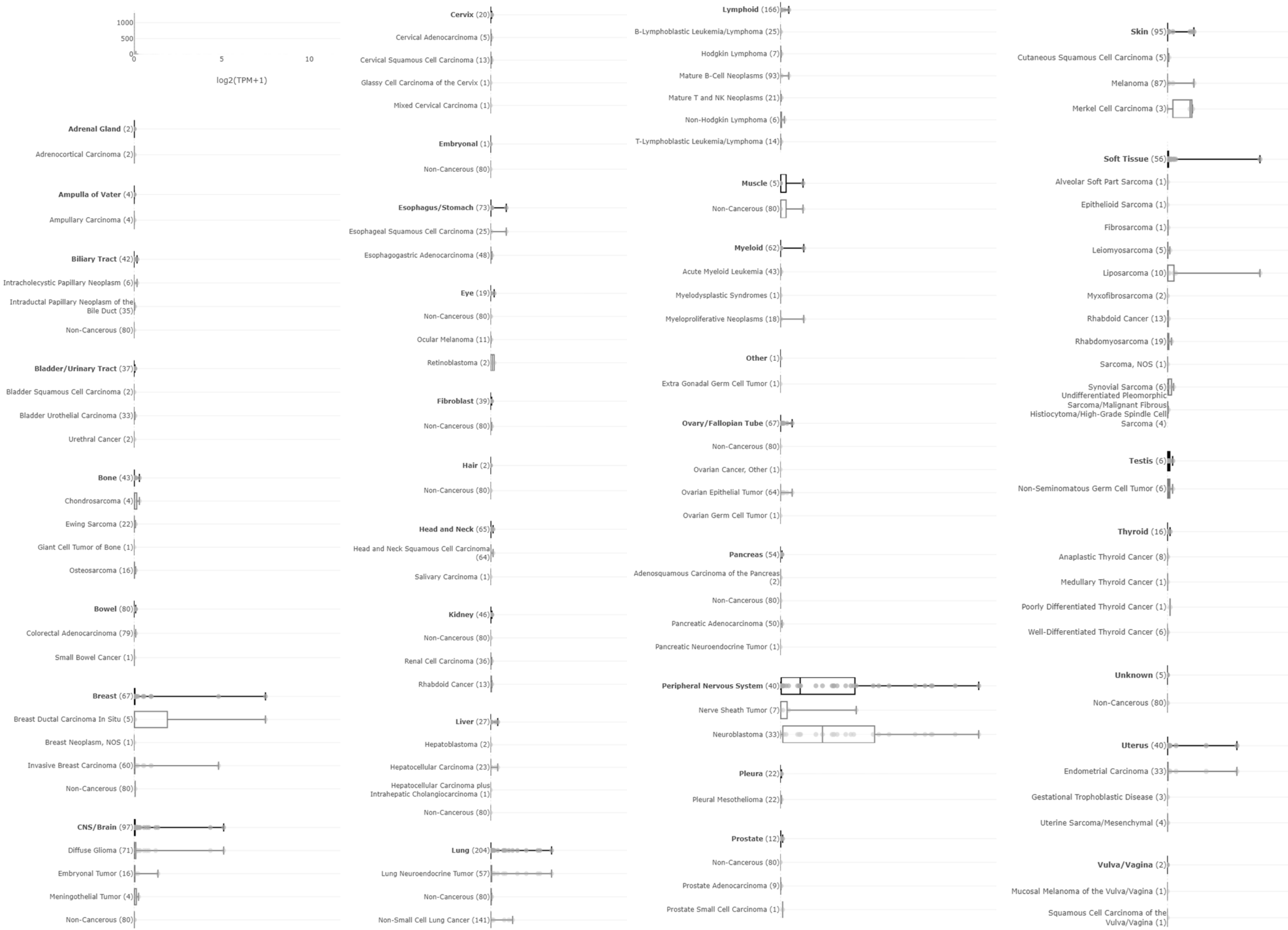

### TCGA Pan-Cancer cohort (cBioPortal):

- Pheochromocytoma, paraganglioma
- Thyroid
- GBM
- some breast cancer cases

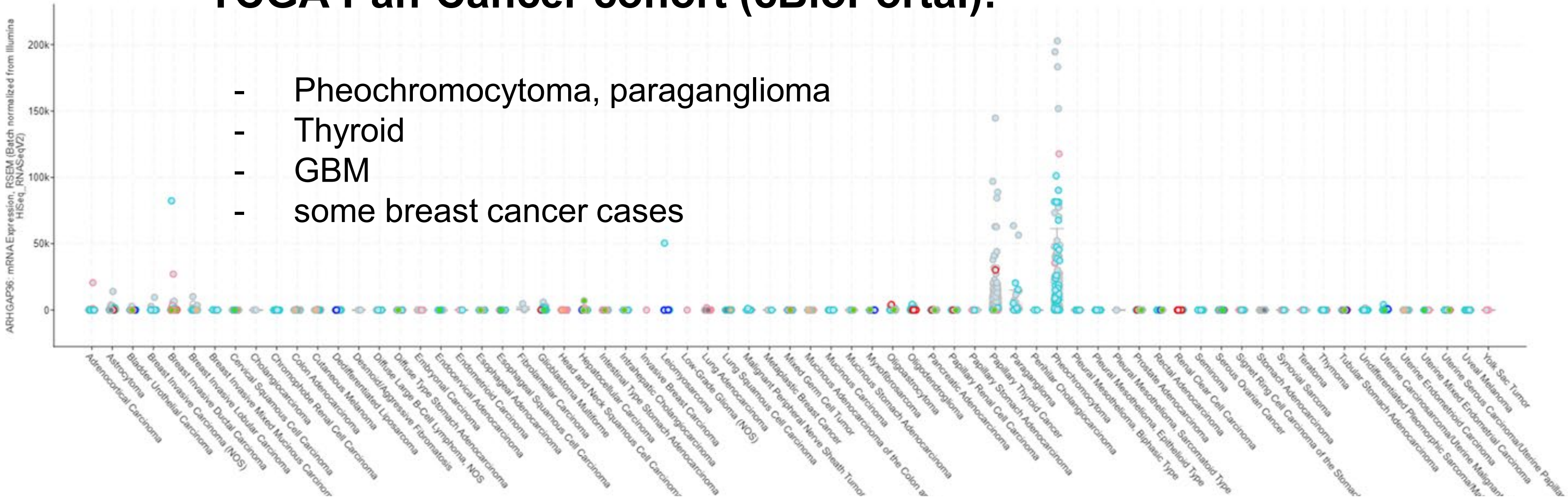


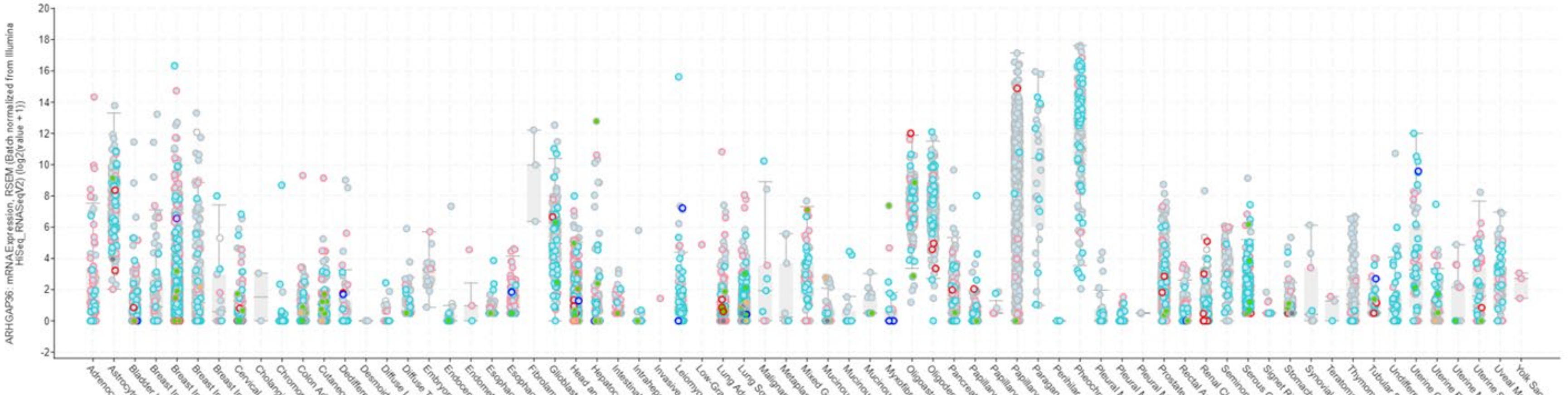

### TARGET Pediatric cohorts (cBioPortal):

- Neuroblastoma
  (large dataset = 1089 samples)

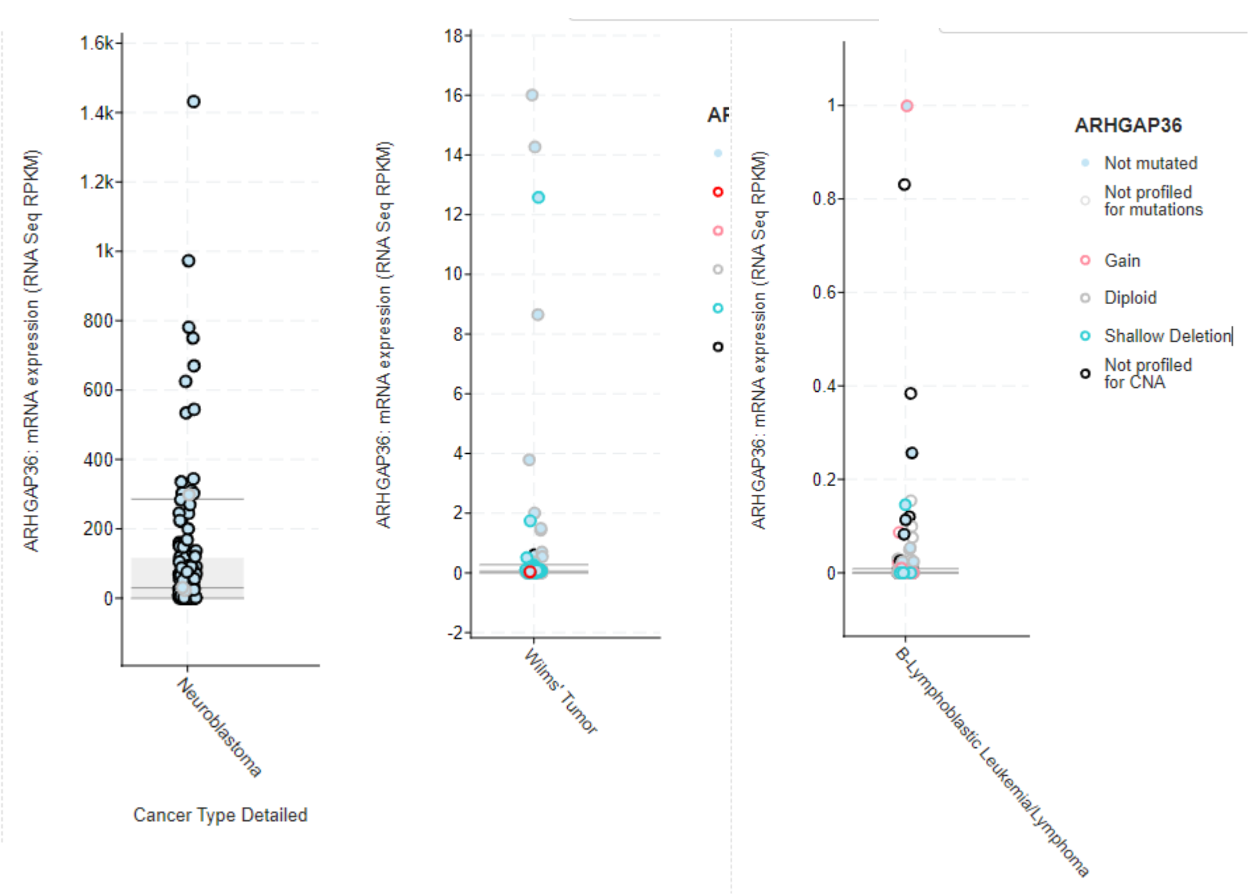

### PCAWG cohort (UCSC Xena):

- Thyroid
- Low grade glioma
- GBM
- some breast cancer cases

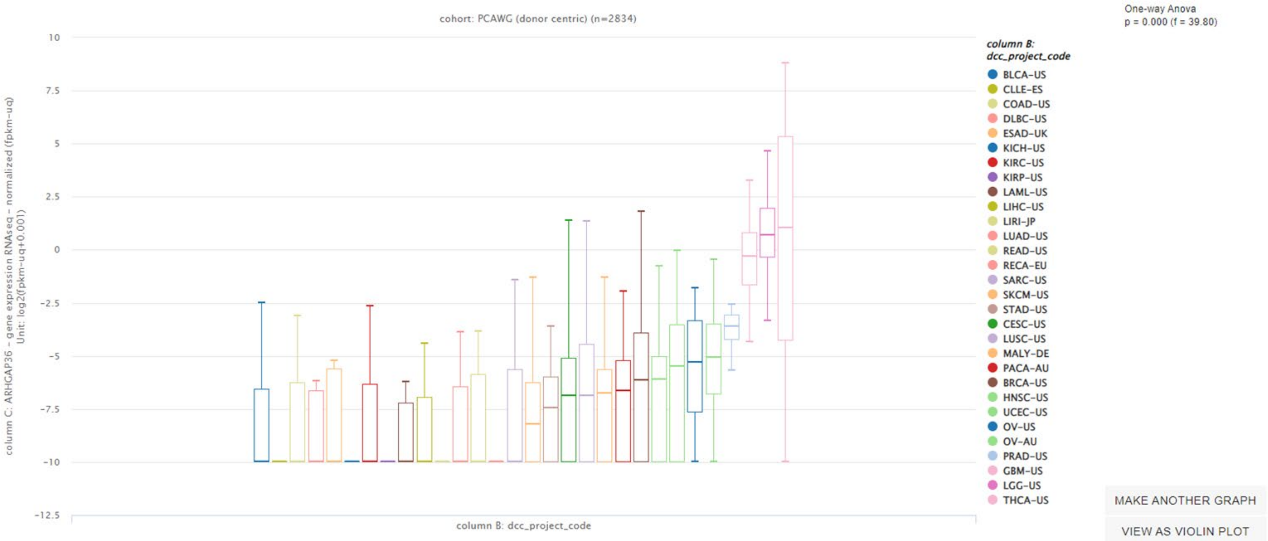


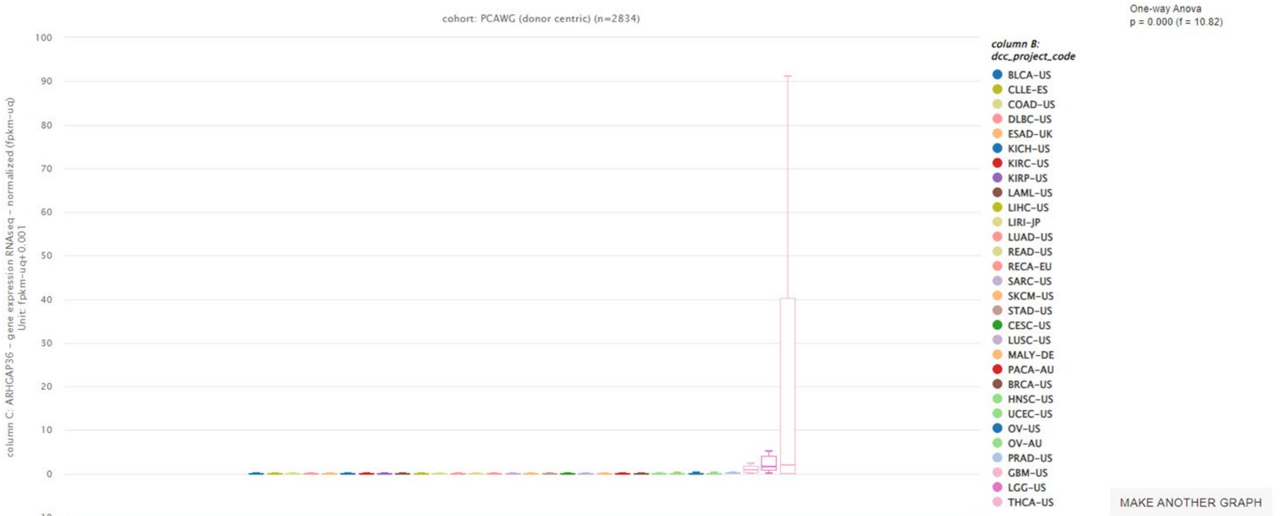

### Normal tissues (GTEx portal):

- Pituitary
- Adrenal gland
- Brain regions
- Muscle
- Breast (F)
- Cervix

\+ Bone (EMBL atlas)

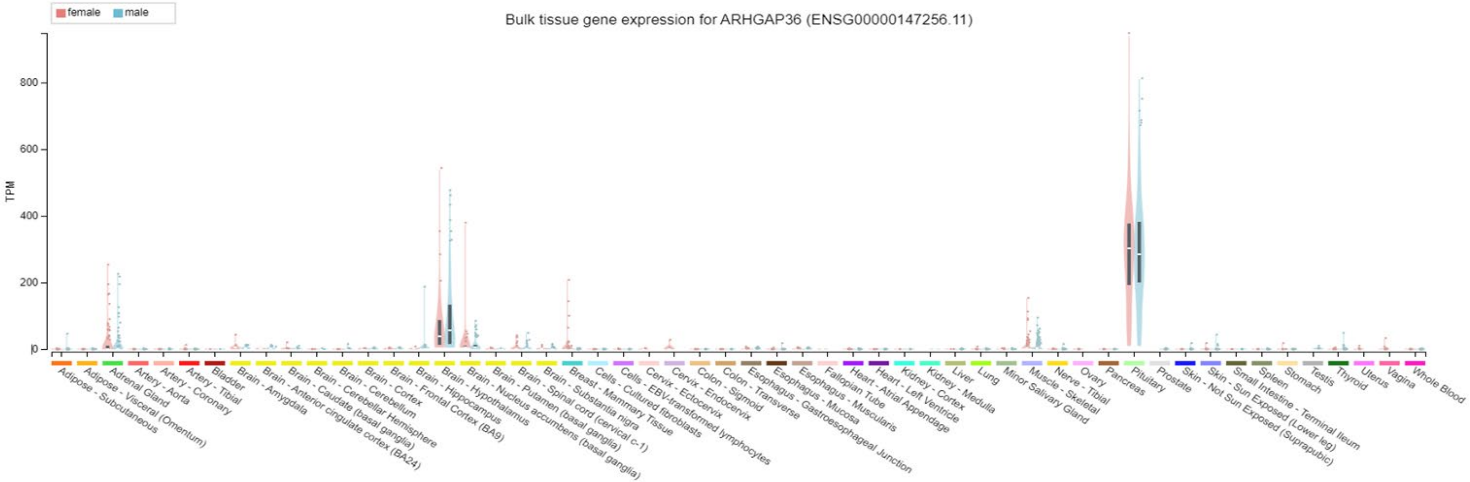


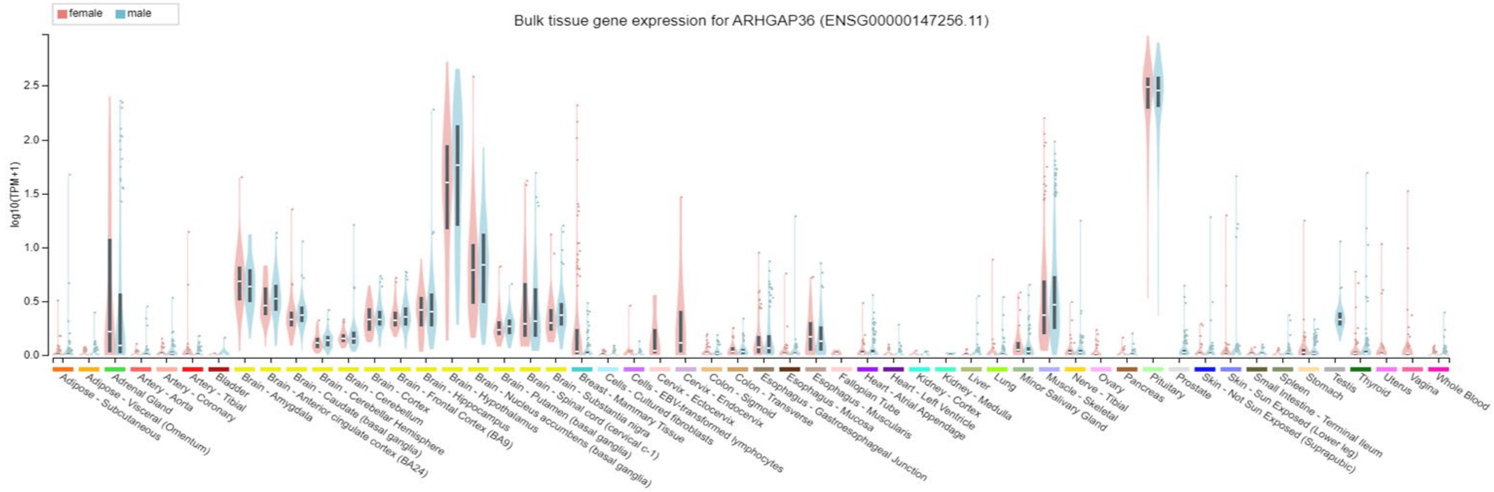
